# Expression of heat shock protein 70 is insufficient to extend *Drosophila melanogaster* longevity

**DOI:** 10.1101/753269

**Authors:** Chengfeng Xiao, Danna Hull, Shuang Qiu, Joanna Yeung, Jie Zheng, Taylor Barwell, R Meldrum Robertson, Laurent Seroude

## Abstract

It has been known for over 20 years that *Drosophila melanogaster* flies with twelve additional copies of the *hsp70* gene encoding the 70 kDa heat shock protein lives longer after a non-lethal heat treatment. Since the heat treatment also induces the expression of additional heat shock proteins, the biological effect can be due either to HSP70 acting alone or in combination. This study used the UAS/GAL4 system to determine whether *hsp70* is sufficient to affect the longevity and the resistance to thermal, oxidative or desiccation stresses of the whole organism. We observed that HSP70 expression in the nervous system or muscles has no effect on longevity or stress resistance but ubiquitous expression reduces the life span of males. We also observed that the down-regulation of *Hsp70* using RNAi did not affect longevity.

## Introduction

*Drosophila* 70 kDa inducible heat shock protein (HSP70) is synthesized in salivary glands, Malpighian tubules, brain, wing imaginal discs and many tissues after a heat shock (Tissieres *et al.* 1974; Lewis *et al.* 1975; Ashburner and Bonner 1979).The expression of *hsp70* coincides with a well-known phenomenon of heat shock-induced chromosomal puffing in *Drosophila* salivary glands (Ritossa 1962, 1963; Ritossa 1964; Ashburner 1970; Tissieres *et al.* 1974). HSP70 is the most abundant protein induced by heat shock in fly and many other organisms (Lewis *et al.* 1975; Ashburner and Bonner 1979; Velazquez and Lindquist 1984). The amino acid sequences of HSP70 are highly conserved among different species (Lindquist and Craig 1988; Daugaard *et al.* 2007). HSP70 has a nucleotide binding domain (NBD) and a substrate-binding domain (SBD) and exhibits ATPase, disaggregase, holdase, translocase and unfoldase activities (Bukau *et al.* 2006; Hartl *et al.* 2011; Saibil 2013; Mattoo and Goloubinoff 2014; Fernandez-Funez *et al.* 2016). The HSP70 protein binds preferentially to hydrophobic residues but no consensus has emerged and the molecular mechanisms responsible for substrate specificity remain unknown (Clerico *et al.* 2015).

*Drosophila hsp70* genes have undergone extensive duplication during evolution (Bettencourt and Feder 2001). There are six nearly identical *hsp70* genes at two adjacent chromosomal loci 87A and 87C (Ishhorowicz *et al.* 1977; Holmgren *et al.* 1979; Gong and Golic 2004). *Drosophila hsp70* contains no introns, presumably allowing rapid and abundant induction since it allows transcripts to circumvent the interruption of RNA splicing during heat shock (Lindquist 1986; Yost and Lindquist 1986). Upon heat shock and other types of stress, HSP70 increases and concentrates in the nuclei and nucleoli, where it is believed to bind incorrectly folded preribosome proteins and ribonucleoproteins (Kloetzel and Bautz 1983; Pelham 1984; Velazquez and Lindquist 1984; Schlesinger 1990). Heat shock has the most effect on nucleoli causing a loss of RNP granules, rRNA synthesis and processing, and ribosome assembly and export. During recovery, HSP70 leaves nuclei and is widely distributed in the cytoplasm. Without stress, cytosolic HSP70 and its cognate family members bind incorrectly folded proteins for refolding or degradation (Schlesinger 1990; Hartl *et al.* 2011; Saibil 2013; Mattoo and Goloubinoff 2014). HSP70 mainly seems to assist folding by stabilization of denatured or unfolding proteins until they reach an appropriate structural conformation.

Many studies that examined the function of HSP70 under stress in a variety of experimental systems supports a protective role. Monkey cells transfected with the *Drosophila hsp70* gene showed a better recovery of nucleolar morphology and of ribosome export after heat shock (Pelham 1984). Rat cells transfected with the human *hsp70* gene have a better thermal resistance (Li *et al.* 1991). Reciprocally, HSP70 immuno-neutralization of rat fibroblasts impairs its translocation to the nucleus and cell survival during thermal stress (Riabowol *et al.* 1988). Competitive inhibition of hsp70 expression elevates the thermosensitivity of hamster ovary cells (Johnston and Kucey 1988). Genetic and transgenic manipulations established that HSP70 alters thermotolerance in the whole organism as well. The addition of 12 extra *hsp70* copies confers greater and faster tolerance to lethal temperatures while the deletion of all *hsp70* genes show the opposite (Welte *et al.* 1993; Feder *et al.* 1996; Gong and Golic 2006; Bettencourt *et al.* 2008). Thermoprotection is also achieved by increasing hsp70 expression with the UAS/GAL4 system. Ubiquitous or motoneurons-specific expression protects larval locomotion during hyperthermia (Xiao *et al.* 2007). Mice lacking the stress-induced hsp70 genes display genomic instability, reduced body weight, sensitivity to ionizing radiations, impaired response to skeletal muscle injury and, enhanced onset, severity and progression of Huntington’s disease symptoms (Hunt *et al.* 2004; Wacker *et al.* 2009; Senf *et al.* 2013). Mice overexpressing hsp70 are less affected and recover better from muscle damage, and display a better preservation of muscle strength and slower progression of muscular dystrophy (McArdle *et al.* 2003; Gehrig *et al.* 2012).

The biochemical properties and protective effects of HSP70 raise the possibility that HSP70 is one of the determinants of the longevity of an organism. HSP70 accumulates with age in the fly muscles in correlation with increased oxidative stress and molecular damages, loss of proteostasis and induction of apoptosis (Wheeler *et al.* 1995; Zheng *et al.* 2005; Demontis and Perrimon 2010). The level of HSP70 is also increased in the skeletal muscles of old rats (Chung and Ng 2006). Tissue-specific selective HSP70 accumulation suggests that it might help prolong the life span of the organism (Wheeler *et al.* 1999). Consistent with this hypothesis, *hsp70* alleles correlate with human longevity and HSP70 level at early age is partially predictive of the remaining life span in flies (Singh *et al.* 2006; Yang and Tower 2009; Singh *et al.* 2010). Furthermore a non-lethal heat treatment extends the adult fly life span and the amplitude of the extension is higher in a strain with twelve additional *hsp70* copies (Khazaeli *et al.* 1997; Tatar *et al.* 1997). Although these observations demonstrate that HSP70 influences longevity, the heat treatment also induces the expression of additional heat shock proteins and therefore the biological effect can be due either to HSP70 acting alone or in combination. The overexpression of *hsp26, hsp27* and *hsp68* with the UAS/GAL4 system is sufficient to increase longevity and the resistance to oxidative stress (Wang *et al.* 2003; Wang *et al.* 2004; Liao *et al.* 2008). Although the overexpression of hsp70 is detrimental to growth and survival during development, several pathways influencing longevity control growth and the hyperfunction theory proposes that growth causes aging (Feder *et al.* 1992; Krebs and Feder 1997; Gems and Partridge 2013).

This study uses the UAS/GAL4 system to control the expression of *hsp70* without the need for a heat treatment and therefore be able to examine whether HSP70 is sufficient to affect organismal longevity and stress resistance.

## Materials and Methods

### Drosophila strains and cultures

Eleven UAS-hsp70 strains carrying independent insertions of pINDY5-hsp70 (pCX1) have been previously obtained (Xiao *et al.* 2007). Seven UAS strains with pCX1 inserted on the X, II or III chromosomes were used: UAS-hsp70 #2.1 (X), UAS-hsp70 #4.3 (X), UAS-hsp70 #9.1 (X), UAS-hsp70 #3.2/CyO (II), UAS-hsp70 #10.1/CyO (II), UAS-hsp70 #3.1 (III) and UAS-hsp70 #4.4 (III). They are referenced in short as #2.1, #4.3, #9.1, #3.2, #10.1, #3.1 and #4.4. The D42 (obtained from G. Boulianne), da-GAL4 (obtained from J. Merriam) and DJ694 Gal4 strains have been already described (Wodarz *et al.* 1995; Yeh *et al.* 1995; Seroude *et al.* 2002). The UAS-lacZ (#1777) strain was obtained from the Bloomington drosophila stock center. Nine UAS-hsp70-RNAi strains (R1.17[III], R1.26[II], R2.17[III], R3.12[X], R4.15[III], R5.2[III], R5.12[II], R5.12e[III] and R5.14[III]) carrying independent insertions of Sym-pUAST-hsp70 (pCX9) were obtained by P-element mediated transformation (Robertson *et al.* 1988). Standard cornmeal medium was used (0.01% molasses, 8.2% cornmeal, 3.4% killed yeast, 0.94% agar, 0.18% benzoic acid, 0.66% propionic acid).

### GAL4 expression across age

D42, da-GAL4 and w males were crossed with UAS-lacZ females. Age-synchronized cohorts of the resulting adult progenies were obtained by emptying cultures and collecting newly emerged flies within 48 h. Males and females were maintained separately at 25°C at a density of 25-30 individuals per vial. Fresh media was provided by transferring to new vials at least twice a week. Individuals were randomly removed at the selected ages by aspiration.

The temporal profiles of GAL4 expression were obtained by quantification of ß-galactosidase activity with a CPRG assay (Seroude *et al.* 2002). Individual flies were manually homogenized in 100 *µ*l extraction buffer (50 mM Na2HPO4/NaH2PO4, 1 mM MgCl2, pH 7.2, 1 cOmplete™ protease inhibitors cocktail tablet(Roche)/40ml). At least five individuals were processed per age, sex and genotype. Extracts were centrifuged for 1 min at 13 000 g. 10 *µ*l of extract supernatant was mixed with 100 *µ*l 1 mM chlorophenolred-ß-D-galactopyranoside (CPRG, Roche). Background levels from the negative control (progeny of cross with w) were averaged and subtracted from the experimental samples. Enzymatic activity was calculated as the rate of change of OD_562nm_ per minute (ΔmOD562/min/fly), and standardized to a single fly equivalent.

The anatomical location of GAL4 expression was determined by 5-bromo-4-chloro-3-indolyl-β-D-galactopyranoside (X-gal) staining (Seroude *et al.* 2002; Poirier *et al.* 2008). At least two males and two females were processed per age and genotype. 10 microns cryosections were fixed for 20 minutes with 1% glutaraldehyde in Phosphate Buffered Saline Solution (1x PBS) (137 mM NaCl, 2.7 mM KCl, 8.1 mM Na2HPO4, 1.5 mM KH2PO4, pH 7.4). Sections were washed twice for 3-5 minutes with 1x PBS before being reacted with the X-gal solution (0.2 % X-gal, 3.1 mM K3Fe(CN)6, 3.1 mM K4Fe(CN)6, 150 mM NaCl, 1 mM MgCl2, 10 mM Na2HPO4/NaH2PO4) at room temperature. Reactions were terminated with 3 3-5 min washes with 1x PBS. The sections were then mounted in 70% glycerol in PBS. Images were obtained on a Zeiss Axioplan II imaging microscope with a Leica DC500 high-resolution camera and the OpenLab imaging software (Improvision, Lexington, MA). All images were captured with the same exposure time and light intensity.

### Western blot

One male and one female were homogenized in 50 *µ*l of extraction buffer (10 mM Tris·Cl, pH 8.0, 50 mM NaCl, and 1 % NP-40) supplemented with protease inhibitors (cOmplete Cocktail Tablet, Roche). Extracts were mixed with 50*µ*l 2× SDS loading buffer (0.125 M Tris·Cl, pH 6.8, 4.6 % SDS, 20 % glycerol, 10 % β-mercaptoethanol, and 0.1 % bromophenol blue), incubated for 5 min at 100°C and centrifuged for 5 min at 10,000 g. 20 *µ*g samples were separated on a 7.5 % SDS-PAGE gel and transferred to nitrocellulose membrane. HSP70-myc was detected with the mouse 9E10 monoclonal anti-myc antibody (11667203001, Roche) at 1:5000 or the mouse 5A5 monoclonal anti-human HSP70 antibody (MA3-007, ThermoFisher Scientific) at 1:1000. α-Tubulin was used as loading control and detected with the mouse DM1A monoclonal anti-chicken α-Tubulin (Ab7291, Abcam) at 1:5000. Primary antibodies were incubated at 4°C overnight. After washes, the nitrocellulose membrane was then incubated with horseradish peroxidase (HRP)-conjugated goat anti-mouse IgG antibody (#170-6516, Bio-Rad) at 1:5000 at 4°C overnight. The ECL advance Western blotting detection kit (RPN2135, Amersham) was used for signal detection. Results were visualized and imaged with ChemiGenius bio-imaging system (Syngene, USA). ImageJ (NIH, USA) was used for the measurement of optic density of specific bands. Hsp70-myc level was normalized to α-Tubulin for quantification. Two-way ANOVA with Bonferroni post-test was performed with GraphPad™ Prism 5 to compare Hsp70-myc expression levels.

### Developmental analysis

Control animals were obtained by crossing w males with homozygous UAS-hsp70 females (UAS control) and homozygous GAL4 driver males with w females (GAL4 control). Experimental animals were generated by crossing homozygous UAS-hsp70 females with homozygous GAL4 driver males. For each cross, 20-30 males and 50 virgin females were mated for two days. At least 200 0-4 h eggs were collected from each cross and transferred with forceps onto petri plates filled with media (25 eggs per plate, 8 plates per genotype). The plates were then incubated at 25 °C. Hatching rate were assessed 26-30h after egg transfer by scoring the number of empty eggshells. Pupation and adult emergence rates were scored 5-6 days and 10-12 days after egg transfer respectively.

### Stress tests

Control animals were obtained by crossing w males with homozygous UAS-hsp70 females (UAS control) and homozygous GAL4 driver males with w females (GAL4 control). Experimental animals were generated by crossing homozygous UAS-hsp70 females with homozygous GAL4 driver males. 0-2 days old males and females were collected from control and experimental crosses and aged at 25°C in separate vials with media (20-30 flies per vial, 2 vials per replicate, vials replaced twice a week) until tested. Paraquat oxidative stress tests were performed as previously described (Lin *et al.* 1998). Dry starvation tests were done by transferring control and experimental animals to empty vial. Heat stress tests were performed by immersing vials in a 36°C water bath (water level above the bottom of the plug). The number of dead flies was scored every 4-5 hours. Kaplan-Meier (Log-rank) analysis was done with GraphPad™ Prism 5 to compare survival upon stress between control and experimental genotypes.

### Longevity measurements

Control animals were obtained by crossing w males with homozygous UAS females (UAS control) and homozygous GAL4 driver males with w females (GAL4 control). Experimental animals were generated by crossing homozygous UAS females with homozygous GAL4 driver males. 0-2 days old males and females were sorted and collected from control and experimental crosses under nitrogen anesthesia for less than three minutes. The longevity of each genotype is obtained from four vials of males and four vials of females (25-30 animals per vial) kept at 25°C. Animals were transferred to vials with fresh media and the mortality recorded at least twice a week. Four to eight replicates were measured. Each replicate used independent cultures and parents. Kaplan-Meier (Log-rank) analysis with GraphPad™ Prism 5 was used to determine the significance of experimental animals that displayed a mean longevity higher or lower than both kinds of control animals.

### Data availability

Strains and plasmids are available upon request. All data necessary for confirming the conclusions of the article are present within the article, figures and supplemental files. Supplemental Figure S1 contains western-blot data obtained with an HSP70 antibody. Supplemental Figure S2 shows the outcome of the survival data to thermal, oxidative and metabolic stresses of 3 days and 10 days old females overexpressing HSP70. Supplemental Figure S3 shows the outcome of the longevity data of females overexpressing HSP70. Supplemental Figure S4 reports the cloning steps performed to obtain the pCX9 plasmid. Supplemental Figure S5 contains western-blot data that show the down-regulation of Hsp70 by the UAS-hsp70-RNAi transgenes. Supplemental Table S1 contains the data and statistical analysis of the survival during development. Supplemental Table S2 contains the data and statistical analysis of the stress resistance experiments. Supplemental Table S3 contains the data and statistical analysis of the longevity experiments with the UAS-hsp70 transgene. Supplemental Table S4 contains the data and statistical analysis of the longevity experiments with the UAS-hsp70-RNAi transgene. The reagent table provides the information and source of transgenic strains, DNA plasmids and antibodies used in this study. Supplemental files are available at FigShare:.

## Results

### Gal4 expression across age

Three GAL4 drivers were selected for this study: da-GAL4 (Wodarz *et al.* 1995), D42 (Yeh *et al.* 1995; Parkes *et al.* 1998) and DJ694 (Seroude *et al.* 2002). DJ694 is an adult muscle-specific driver. The pattern and dynamics of GAL4 expression during aging has been described (Seroude *et al.* 2002). Newly emerged DJ694 mostly express GAL4 in the abdominal muscles. The expression level increases rapidly, predominantly in the thoracic flight muscles, to reach a two-to three-fold higher level within ten days. It remains constant until around 30 days and then gradually declines. Expression patterns of da-GAL4 and D42 from the embryo to the emergence of the adult have been reported (Wodarz *et al.* 1995; Yeh *et al.* 1995; Parkes *et al.* 1998). However, the anatomical location and dynamic of GAL4 expression during aging is missing. Therefore the spatial-temporal expression of GAL4 across age was determined by driving a UAS-lacZ reporter encoding the *E. coli* β-galactosidase (Figure 1).

**Figure 1:**
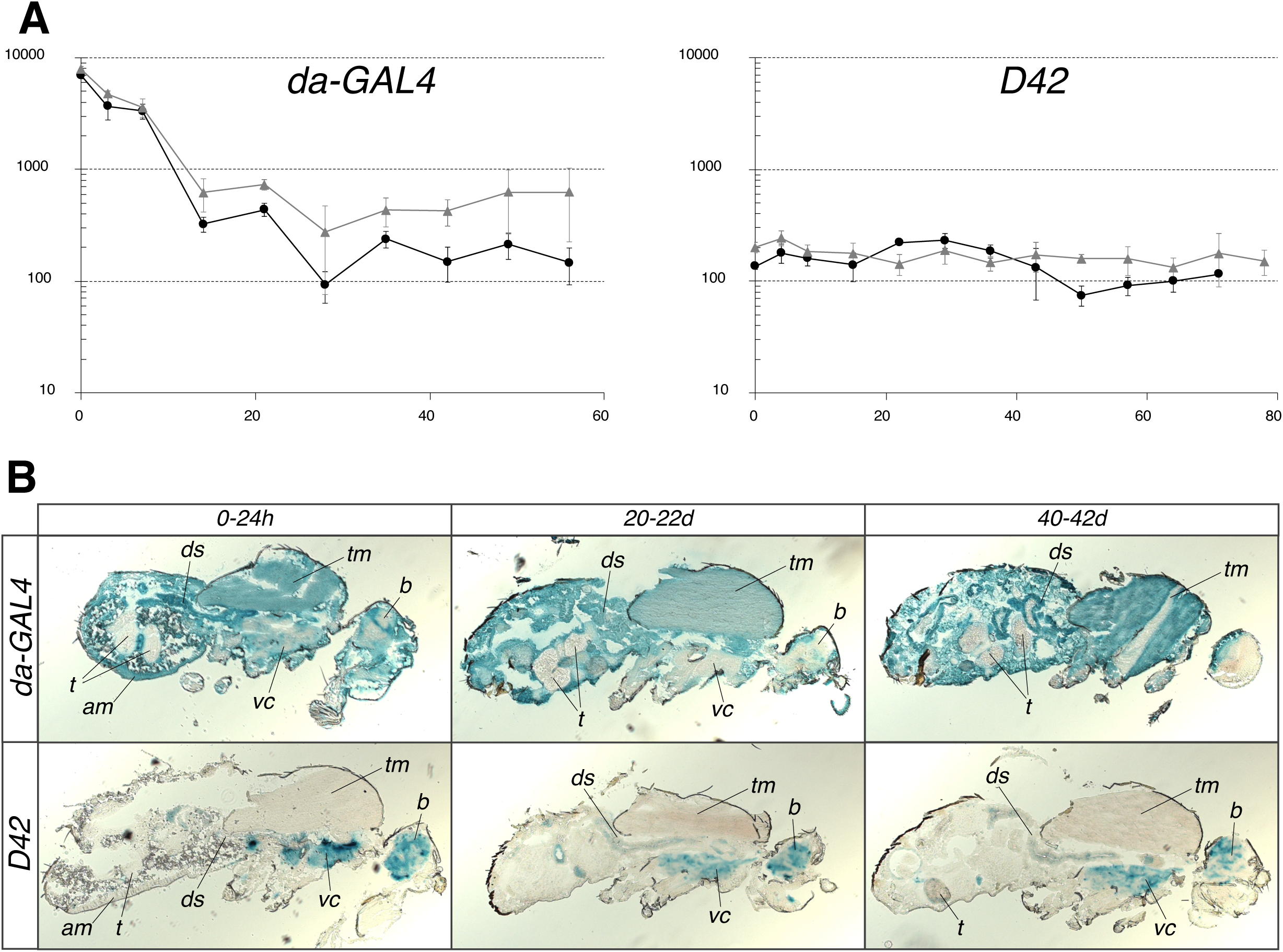
Spatial-temporal expression of da-Gal4 and D42 in adult flies. A) Quantification of ß-galactosidase activity across age in males (black circles) and females (grey triangles) with a CPRG assay. Each data point is the average of five measurements ±SD. y axis: Activity (ΔmOD/min/fly); x axis: Age (days at 25°C). B) Localization of ß-galactosidase activity across age by X-Gal staining. am: abdominal muscles; b: brain; ds: digestive system; t: testis; tm: thoracic muscles; vc: ventral chord.

The temporal expression profiles show that da-GAL4 expresses GAL4 at the highest level upon emergence but it drastically decreases by at least twenty-fold during the first two weeks and remains constant for the remainder of the life span (Figure 1A). It appears that the initial decrease is slightly less pronounced in the females. The D42 driver displays a constant level of GAL4 expression across the entire life span that is almost identical between sexes. The expression across age shows that da-GAL4 drives GAL4 expression almost ubiquitously while D42 is strictly restricted to the nervous system (Figure 1B). The da-GAL4 driver is not completely ubiquitous because no expression is detected in the testis. Furthermore it is obvious that the expression is not homogenous. In the younger animals, the signal in the central nervous system (brain and ventral cord) appears weaker than in other tissues and in the older animals is barely detectable. The signal is also hardly visible in one of the vertical thoracic flight muscle of the oldest animals. Although D42 targets specifically the central nervous system, it is not homogenous. Since both drivers drive similar level of expression in older animals, D42 result in higher overexpression in the nervous system than da-GAL4.

### hsp70 overexpression

Transgenic UAS-hsp70 fly strains that carry the Hsp70Ab cDNA under the control of a UAS promoter have already been obtained and described (Xiao *et al.* 2007). These strains express the *hsp70* coding region followed by a C-terminal EQLRSTSRTM linker and EQKLISEEDL myc epitope. The HSP70-myc fusion produced can be differentiated from the endogenous HSP70 by size and can be detected with myc and HSP70 antibodies (Figure 2 and S1). This HSP70-myc fusion is biologically active and functional since it is able to protect larval locomotor activity during hyperthermia (Xiao *et al.* 2007), to protect adult flies from viral infection (Cappucci *et al.* 2019), and to activate transposable elements (Merkling *et al.* 2015). The level of *hsp70* expression can vary between strains because the UAS-hsp70 transgene does not contain chromatin insulator elements and can therefore be affected by the genomic sequences surrounding its insertion. The effects of the genome are advantageous by providing an opportunity to test different levels of expression with the UAS/GAL4 system without using the temperature shifts that are required to modulate GAL4 activity directly or indirectly with a thermosensitive GAL80 negative regulator of GAL4 (Lee and Luo 1999; Duffy 2002). Therefore, this study used seven UAS-hsp70 strains. The level of HSP70-myc expression was determined by Western-blot with a myc antibody (Figure 2). Three and ten days old animals were processed because of the early expression changes associated with the DJ694 and da-GAL4 drivers. Three days after emergence, a specific HSP70-myc signal is clearly visible in the extracts from animals carrying the da-GAL4 transgene and one of the UAS-hsp70 transgene but not in extracts from control animals that lack one of the two transgenes (Figure 2A). At ten days, as expected from the temporal profile of GAL4 expression, the HSP70-myc signal is significantly reduced compared with three days (P < 0.0001)(Figure 2B). Although most of the variation is largely due to age factor, the genomic location of the UAS-hsp70 transgene (genotype) has a significant (P < 0.0001) but modest effect on the expression level.

**Figure 2:**
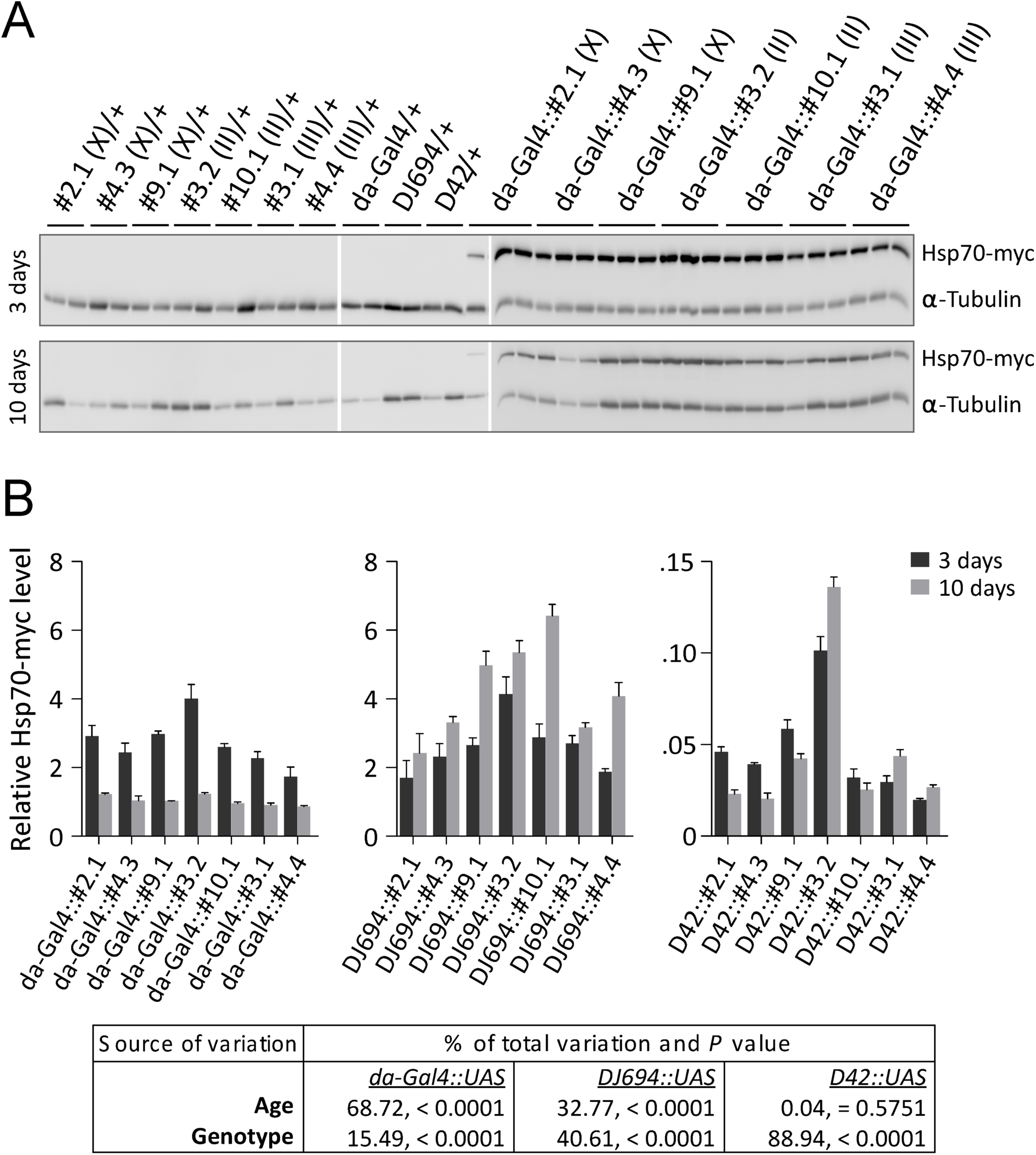
Transgenic Hsp70 expression. A) Western-blot with monoclonal 9E10 antibody of Hsp70-myc expression driven by da-Gal4 at three and ten days. (B) Quantitative analysis. The table shows the sources of variance by a two-way ANOVA.

In agreement with the GAL4 expression profile, Hsp70-myc levels increased significantly between three and ten days with the DJ694 driver (P < 0.0001) but not with the D42 driver (P = 0.5751). With both drivers, the location of the UAS transgene has a significant effect on the expression level and the range of expression obtained is wider than with the da-GAL4 driver. Between the genotypes with the lowest and highest expression level, there is a ∼40% (10 days) to ∼130% (3 days) difference with da-GAL4, a ∼135% (3 days) to ∼165% (10 days) with DJ694 and a ∼410% (3 days) to ∼560% with D42 (10 days).

### hsp70 overexpression does not cause developmental defects

Since the constitutive expression of HSP70 in the larval salivary glands at normal temperatures severely reduced cell size (Feder *et al.* 1992), a developmental analysis was first conducted to determine if the GAL4-driven expression induce developmental delay or lethality (Figure 3, Table S1). Whatever GAL4 driver is used and whatever the genomic location of the UAS-hsp70 transgene, the totality of viable individuals for any genotype (controls or experimental) were obtained 26-30 h (first instar L1 larvae), 5-6 days (pupae) and 10-12 days after egg laying. Earlier scoring underestimated the number of viable individuals while later scoring did not change the total number indicating that *hsp70* expression with these drivers does not delay development.

**Figure 3:**
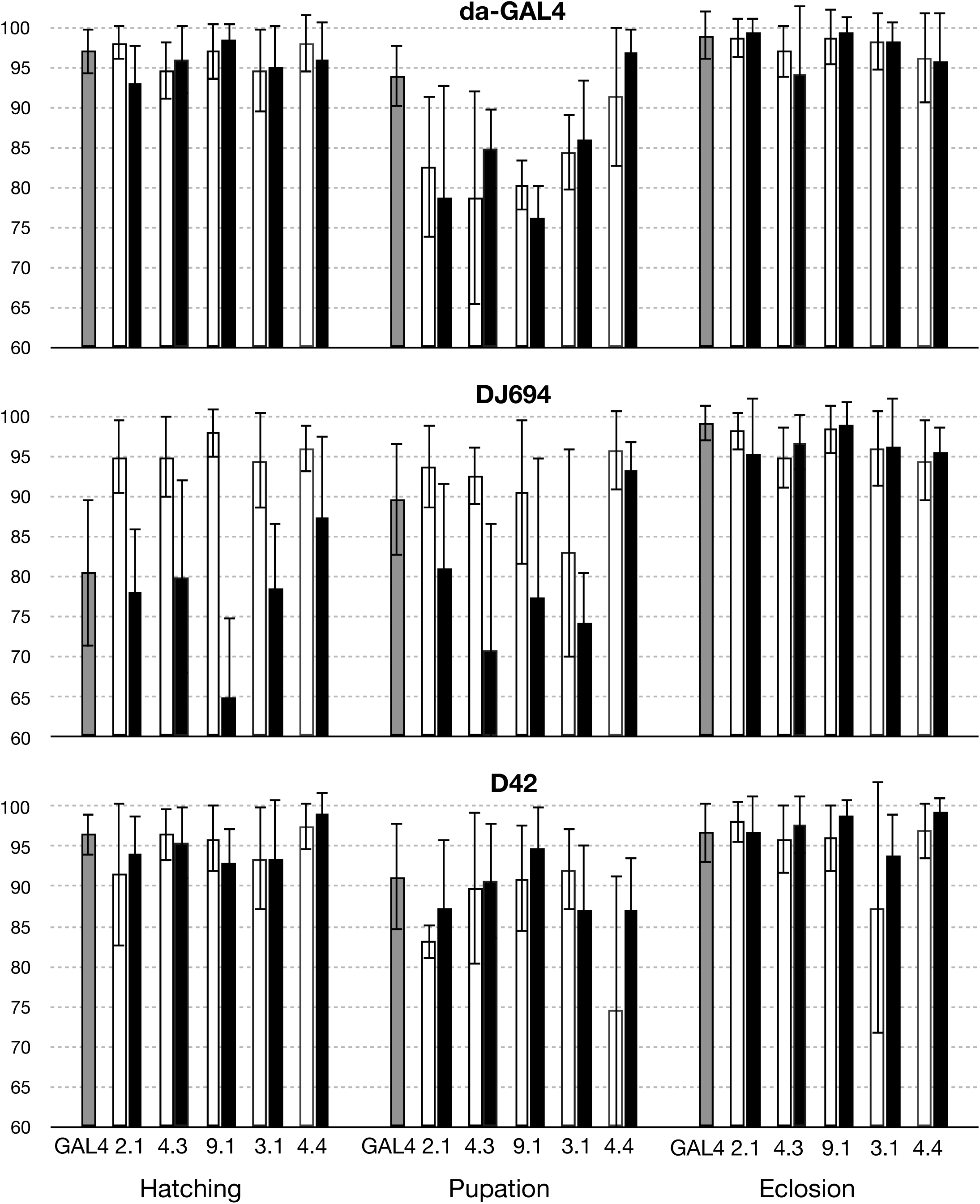
Survival during development of flies overexpressing HSP70. The mean L1 larvae (% hatching), pupae (% pupation) and adult (% eclosion) survival ±SD are plotted. Each graph compares the experimental GAL4 + UAS animals (black) to control animals missing the UAS transgene (GAL4 alone, grey) or the GAL4 driver (UAS alone, white). Statistical analysis is provided in Table S1.

No lethality was observed with the D42 driver. With the DJ694 driver, the UAS-hsp70 insertions #9.1 and #10.1 yielded significantly less L1 larvae indicating some embryonic lethality. The #4.3 insertion with the DJ694 driver and the #9.1 with the da-GAL4 driver yielded significantly less pupae indicating some larval lethality. Since most of the UAS-hsp70 insertions do not show any significant differences between experimental and control animals, the embryonic and larval lethality observed result from insertion effects, leading to the conclusion that the overexpression of hsp70 does not impair development.

### hsp70 overexpression does not affect stress resistance

The survival to three kinds of stress was measured in both sexes after three and ten days of adulthood (Table S2). High temperature stress was tested by exposure at 36°C. Oxidative stress was assessed by feeding the free radical generator paraquat. Desiccation stress was also examined with a dry starvation assay. At least three and up to 10 replicates were processed per stress, sex, age, UAS-hsp70 insertion and GAL4 driver. The results from each replicate are compiled in the figure 4 (males) and the supplemental figure S2 (females). These data do not show any consistent effect between replicates across the seven UAS strains whatever the kind of stress and location of hsp70 expression. Furthermore only very few genotypes displayed consistently a significant effect. Consistent identical effects in all the replicates is only seen in two situations in each sex. In males, the #9.1 insertion with the ubiquitous driver decreased oxidative stress survival at 10 days while the #3.1 insertion with the nervous system driver increased it at 3 days. In females, at 10 days the nervous system driver increased the survival to heat with the #3.2 insertion while it decreased the survival to starvation with the #9.1 insertion. Overall these data indicate that hsp70 expression is insufficient to affect the resistance of the organism to heat, oxidative or desiccation stresses.

**Figure 4:**
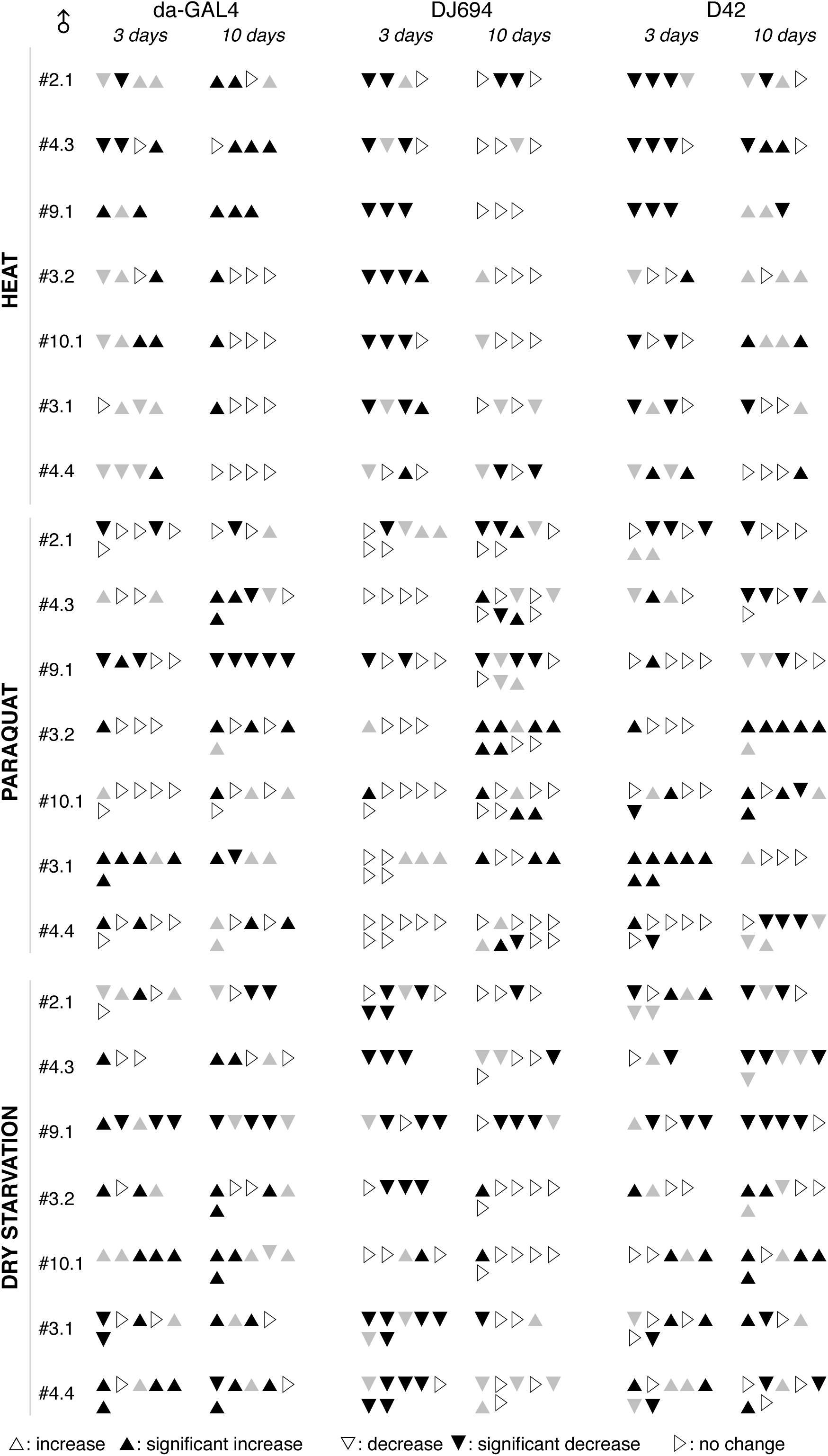
Survival to thermal, oxidative and metabolic stresses of 3 days and 10 days old males overexpressing HSP70. Each triangle reports the outcome of the statistical comparison (Table S2) between experimental and controls for an experimental replicate. ▴: significant increase ▴: increase ▾: significant decrease ▾: decrease ▹: no change.

### hsp70 is insufficient to extend longevity

The longevity of flies with both UAS-hsp70 and GAL transgenes was measured and compared to control flies lacking the GAL4 or UAS transgene (Table S3). Consistent results are only seen across all UAS-hsp70 insertions in males with the ubiquitous driver (Figure 5). All insertions resulted in a reduction of the life span in all replicates with the exception of #2.1 and #3.1 that showed this effect in 5 out of 6 and 6 out of 8 replicates respectively. No consistent effects are seen in females (Supplemental Figure S3) or with the DJ694 and D42 drivers. These results show that the overexpression of HSP70 in the muscles or nervous system is insufficient to affect longevity. However, the reduction of the life span of males with the ubiquitous driver indicate that HSP70 is detrimental in an anatomical location that remains to be identified.

**Figure 5:**
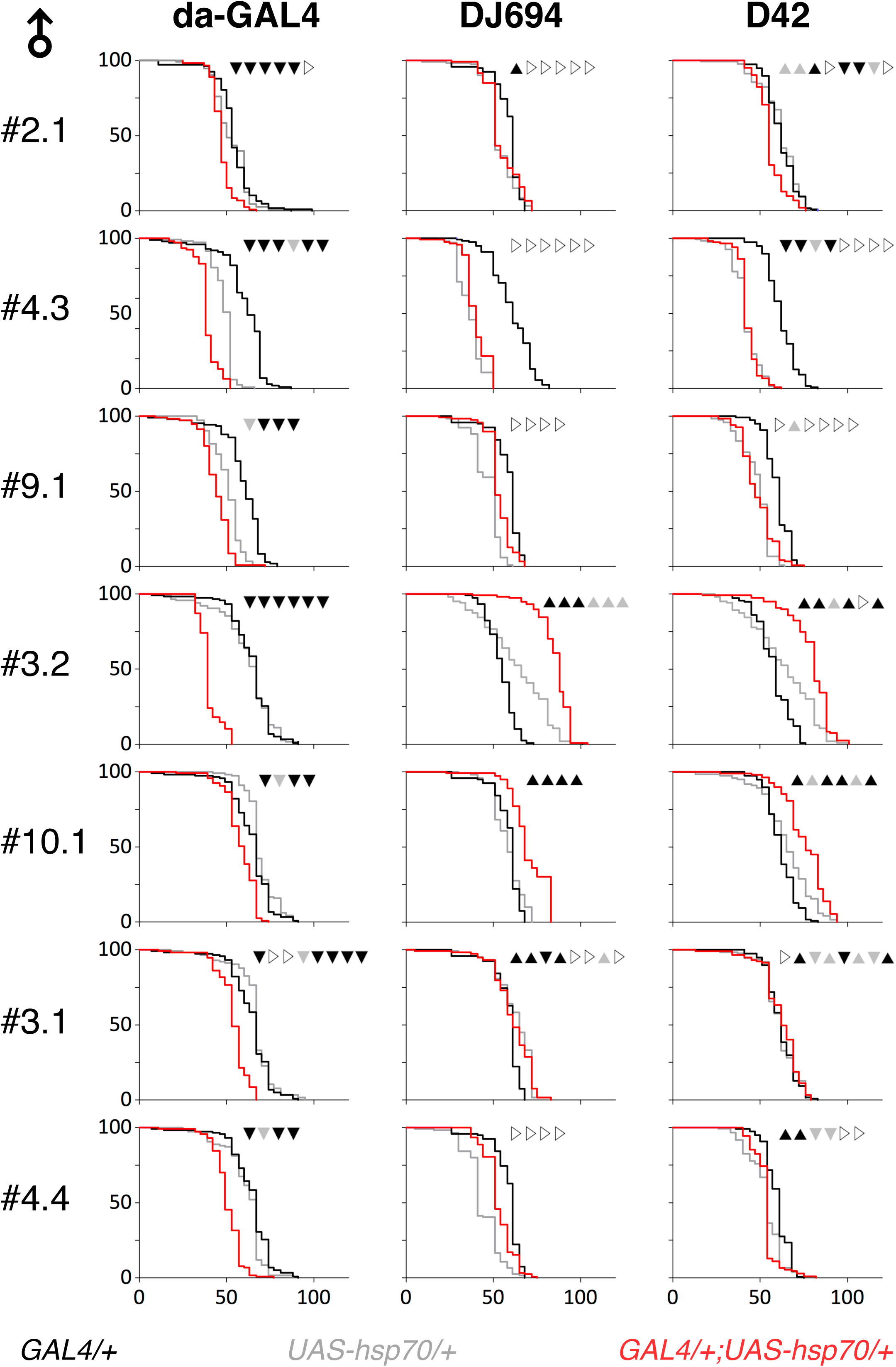
Longevity of males overexpressing HSP70. Typical longevity curves (x axis: days at 25°C; y axis: percentage alive) are shown. The outcome of the statistical comparison (Table S3) between experimental and controls for each experimental replicate is provided at the top of each graph: ▴: significant increase ▴: increase ▾: significant decrease ▾: decrease ▹: no change.

Life span extensions were consistently obtained across replicates with only two (#3.2 and #10.1) out of the seven insertions tested. Both insertions driven by DJ694 or D42 extend the longevity of males in all replicates except for #3.2 that revealed an extension in 5 out of 6 replicates with D42. The extensions associated with #3.2 appear to be also consistent in the females but with a lower magnitude. The extensions associated with #10.1 is seen in females with DJ694 but not with D42. The increased longevity with these insertions is not correlated with the level of hsp70 expression. With D42, #3.2 and #10.1 express the highest and lowest levels respectively. With DJ694, #3.2 and #10.1 express high levels that are similar to the #9.1 insertion that does not display any change in all replicates. Since the extension of longevity is not observed with all the insertions tested and is not correlated with HSP70 levels, the longevity phenotypes observed with the #3.2 and #10.1 are an artifact of the transgene insertion site.

### Down-regulation of hsp70 does not affect longevity

The *hsp70* cDNA section was excised from the pCX1 plasmid (Xiao *et al.* 2007) previously used to generate the UAS-hsp70-myc transgenic strains and inserted into the SympUAST RNAi plasmid (Supplemental Figure S4) (Giordano *et al.* 2002). The resulting pCX9 plasmid allows symmetrical transcription of *hsp70* from two convergent UAS promoters and was used to obtain nine transgenic strains. The effectiveness to reduce *hsp70* expression was first tested with the DJ694 driver by co-expressing each UAS-hsp70-RNAi insertion with the UAS-hsp70 #3.1 insertion (Supplemental Figure S5A). Seven out of the nine RNAi insertions completely prevented the detection of the transgenic HSP70-myc whereas it remains detectable with R5.2 and unaffected with R5.12. Next the effectiveness of the R3.12 insertion to reduce endogenous hsp70 expression was examined with the three GAL4 drivers (Supplemental Figure S5B). As one would expect from the anatomical location of GAL4 expression, almost complete or strong reduction of heat-induced HSP70 is observed with daGAL4 and DJ694 respectively whereas D42 expression is too localized to affect whole animal extracts. Finally, the effects of the DJ694-driven R3.12 and R5.14 insertions on endogenous HSP70 expression were compared (Figure 6A). Both insertions clearly suppress HSP70 expression but R3.12 appears to be more efficient when the level of expression is higher after heat shock.

**Figure 6:**
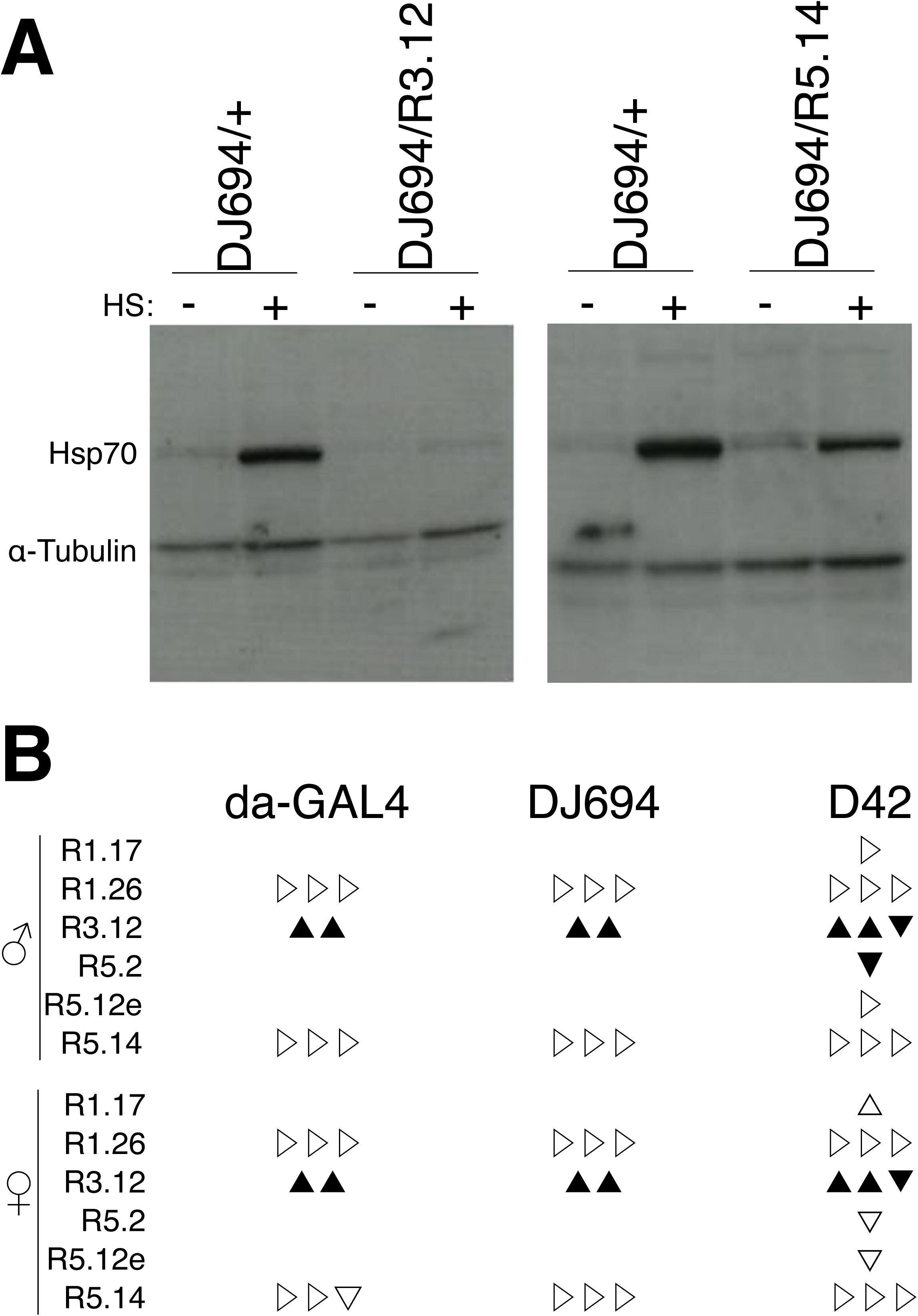
Hsp70 RNAi. A) Western-blot analysis of seven days old male extracts. HS -: no heat-shock; HS +: heat-shock. B) Longevity summary. Each triangle reports the outcome of the statistical comparison (Table S4) between experimental and controls for an experimental replicate: ▴: significant increase ▴: increase ▾: significant decrease ▾: decrease ▹: no change.

The longevity with both UAS-hsp70-RNAi and GAL transgenes was measured and compared to control flies lacking the GAL4 or UAS transgene (Table S4). Two out of the three RNAi insertions tested with all drivers did not show any effects in both sexes (Figure 6B). These results indicate that the down-regulation of hsp70 is insufficient to influence the longevity of the organism.

## Discussion

The UAS/GAL4 system allowed extensive and precise control of Hsp70 expression within a constant environment. The manipulation of HSP70 level during the pre-adult stages did not cause any visible morphological defect, lethality or developmental delays. Under normal conditions, *Hsp70* mRNA (Ireland *et al.* 1982; Welte *et al.* 1993) and protein (Feder *et al.* 1996) cannot be detected in embryos and larvae, and null mutants are morphologically normal, viable and fertile (Gong and Golic 2004). Developmental lethality has been reported when twelve extra copies of *Hsp70* are present (Krebs and Feder 1997) but the level of HSP70 expression achieved by the UAS/GAL4 system is not comparable since the DJ694 and D42 drivers are tissue-restricted and the amount of HSP70 produced with da-GAL4 is obviously lower than the amount produced by heat shock (Figure S1)(Klose *et al.* 2005). Since the analysis of null *Hsp70* mutants have demonstrated that HSP70 is required for basal and induced thermotolerance of larvae and adults (Gong and Golic 2006; Shilova *et al.* 2018), the stress resistance of animals expressing RNAi transgenes was not investigated. The overexpression of *Hsp70* with any of the three GAL4 drivers did not alter the response of adult flies to thermal, oxidative or metabolic stresses. Although it has long been known that increasing *Hsp70* expression is sufficient to increase the thermotolerance of cultured cells (Pelham 1984; Li *et al.* 1991; Solomon *et al.* 1991), it is not surprising that it is not the case *in vivo* in the whole organism (Halliwell 2003; Yazdani 2016; Place *et al.* 2017). This report adds further evidence that elevated *Hsp70* expression is sufficient to affect thermotolerance of embryos (Welte *et al.* 1993), larvae and pupae (Feder *et al.* 1996) but not adults (Feder and Krebs 1998; Jensen *et al.* 2010). The effect of *Hsp70* overexpression on the resistance to oxidative and wet starvation stresses has been investigated in the presence of additional *Hsp70* copies and did not have any consequences with or without heat shock (Minois 2001). The UAS/GAL4 system yields a range of HSP70 expression levels that are intermediate between the uninduced low level and the heat-induced high level used by Minois. This study complements the observation reported by Minois and reinforces the conclusion that the manipulation of *Hsp70* expression alone is insufficient to affect the resistance of the organism to thermal, oxidative or metabolic stress. The reduction of *Hsp70* expression with RNAi transgenes did not affect longevity. Although the western-blot analysis confirmed the effectiveness of the transgenes, it remains possible that there is residual expression. The null *Hsp70* mutant does completely prevent expression and has been recently reported to exhibit a modest reduction of the life span (Shilova *et al.* 2018) indicating that *Hsp70* is required for longevity and that only minimal amount is needed accounting for the lack of correlation between longevity and *Hsp70* RNA levels in wild-type strains (Zhao *et al.* 2005). The UAS/GAL4 system allowed for testing of the effect of *Hsp70* levels above wild-type levels without any environmental modifications that affect the expression of additional genes. Ubiquitous GAL4-mediated *Hsp70* overexpression reduced male longevity while ubiquitous overexpression by heat treatment of males carrying extra *Hsp70* copies had the opposite effect (Tatar *et al.* 1997). This result suggests that *Hsp70* has detrimental effects on longevity that are mitigated or abolished by the simultaneous expression of one or several gene products induced by the heat treatment. After extended periods of recovery from heat shock HSP70 intracellular granules are formed to control HSP70 activity by sequestration (Feder *et al.* 1992), a mechanism that may depend on gene product(s) induced by heat. Alternatively, the overexpression of *Hsp70* with the da-GAL4 driver during the pupal stage may not be detrimental enough to cause lethality and instead caused the emergence of sick animals. It is unlikely that the detrimental effect is due to overexpression in the nervous system or muscles since it is not observed with the D42 or DJ694 drivers. It will be worth investigating additional GAL4 drivers with different tissue-specificity in order to identify the anatomical location responsible for the detrimental effect in males.

The overexpression of *Hsp70* in the nervous system, muscles in both sex or ubiquitously in females had neither beneficial nor detrimental effects on life span indicating that HSP70 is insufficient to extend longevity. It is important to note that identical results were obtained in muscles with the MHC-GAL4 driver and independent UAS-hsp70 strains (Demontis and Perrimon 2010). However it cannot be excluded that higher levels of overexpression may be required to affect the life span since heat treatment of animals with extra Hsp70 copies lead to higher overexpression levels than the GAL4 drivers used in this study. The level of expression induced by the DJ694 driver is the highest that is currently available (Seroude 2002; Seroude *et al.* 2002; Barwell *et al.* 2017) and preliminary investigations using two instead of one copy of the UAS transgenes failed to further increase expression levels. It is also possible that the overexpression during the pre-adult stage is not detrimental enough to cause developmental lethality and reduced longevity but enough to prevent the extension of the life span although it is unlikely with the DJ694 driver that is not expressed in muscles at any stage of development (Barwell *et al.* 2017). If HSP70 alone is indeed insufficient to extend longevity, the extension after heat treatment of animals with extra *Hsp70* copies would suggest that HSP70 extends longevity in combination with one or several gene products induced by the heat treatment. The identification of HSP70 partners could easily be investigated by measuring the longevity of animals carrying a UAS-hsp70 transgene as well as a UAS transgene for a gene to be tested.

## Supporting information

Reagents table

## Authors contribution

CX, DH, SQ, JY and JZ performed experiments. CX, TB and LS analyzed experimental data. RMR and LS designed the study and obtained funding. LS and TB wrote the manuscript.

## Acknowledgments

The authors thank the Bloomington Stock Center (National Institutes of Health P40OD018537), Hugo Bellen, Gabrielle Boulianne and John Merriam for fly strains, and Rhonda Kristensen and Frederique Seroude for technical support. This work was supported by the Institute of Aging of the Canadian Institutes of Health Research (CIHR, MOP64248, MOP79519) and the Natural Sciences and Engineering Research Council of Canada (RGPIN/250140-2010). The sponsor(s) had no involvement in study design; in the collection, analysis and interpretation of data; in the writing of the report; and in the decision to submit the article for publication.

