## Supplementary material for "Expression of heat shock protein 70 is insufficient to extend *Drosophila melanogaster* longevity": Figure S1

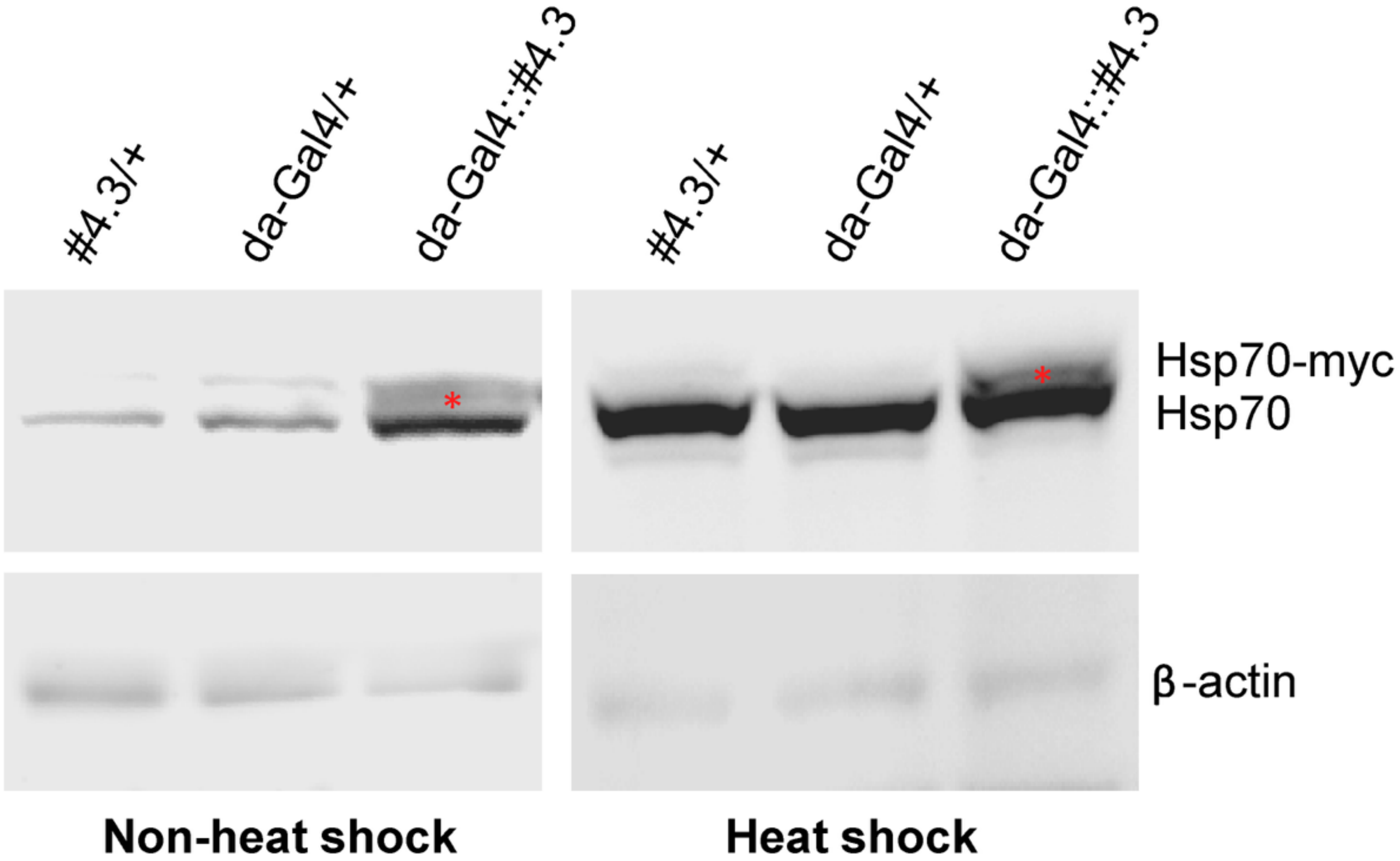


**Figure S1. Transgenic and endogenous Hsp70 expression.** Western-blot with monoclonal antibody 5A5 was performed to detect HSP70-myc (red asterisk) and endogenous HSP70 in 5-7 days old males with or without heat shock. Under both conditions, the intensity of the HSP70-myc signal is lower than the endogenous HSP70 signal. Although it remains possible that the 5A5 antibody has a lower affinity for the tagged protein, it is very likely that the level of GAL4-mediated HSP70 overexpression under non-heat shock condition is lower than the level of heat-shock induced endogenous HSP70 overexpression.
