## Supplementary material for "Expression of heat shock protein 70 is insufficient to extend *Drosophila melanogaster* longevity": Figure S2

**Supplemental Figure 2**

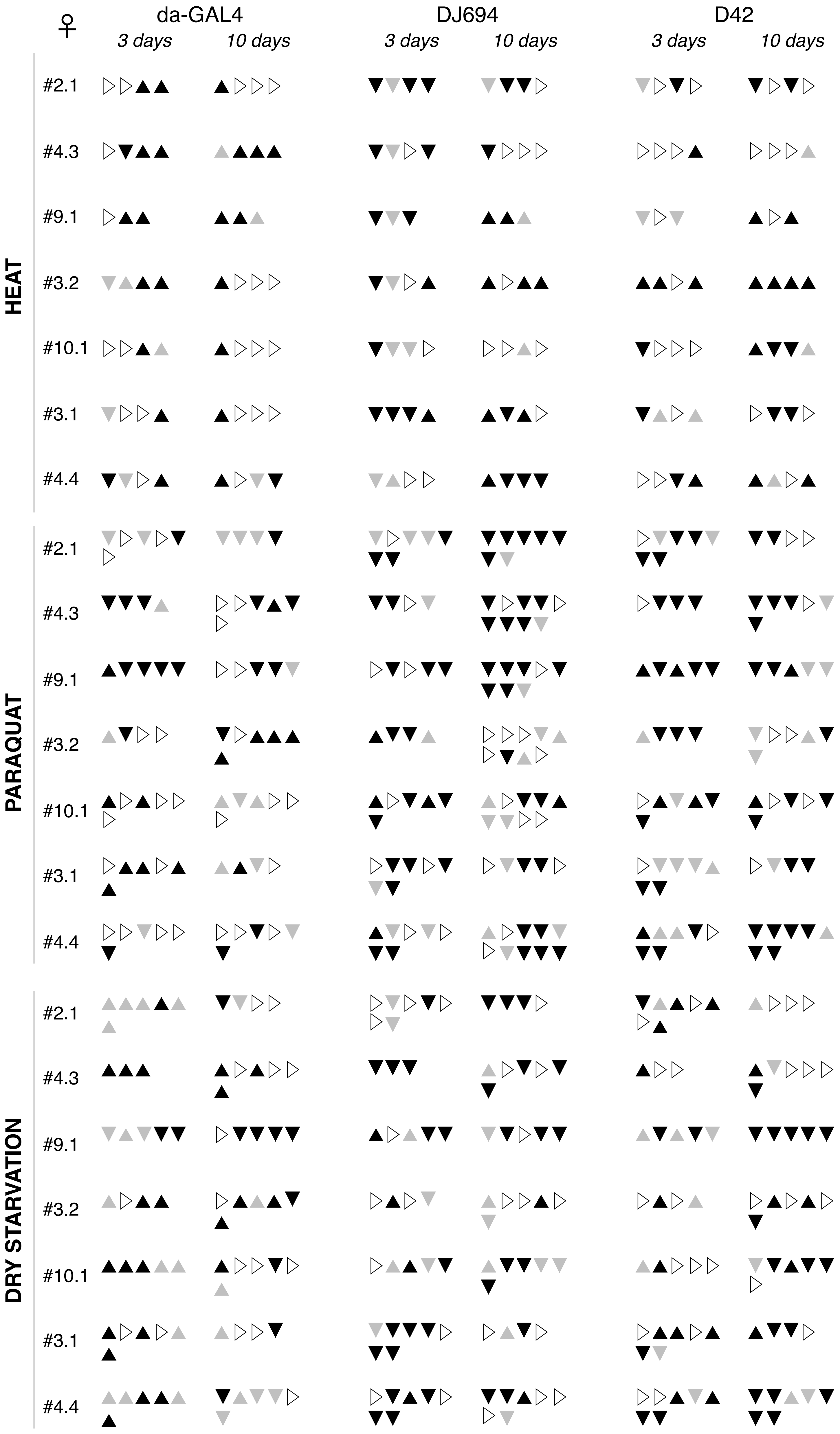


△: increase ▲: significant increase ▽: decrease ▼: significant decrease ▷: no change

**Figure S2: Survival to thermal, oxidative and metabolic stresses of 3 days and 10 days old females overexpressing HSP70.** Each triangle reports the outcome of the statistical comparison (Table S2) between experimental and controls for an experimental replicate.
