## Supplementary material for "Expression of heat shock protein 70 is insufficient to extend *Drosophila melanogaster* longevity": Figure S3

### Supplemental Figure 3

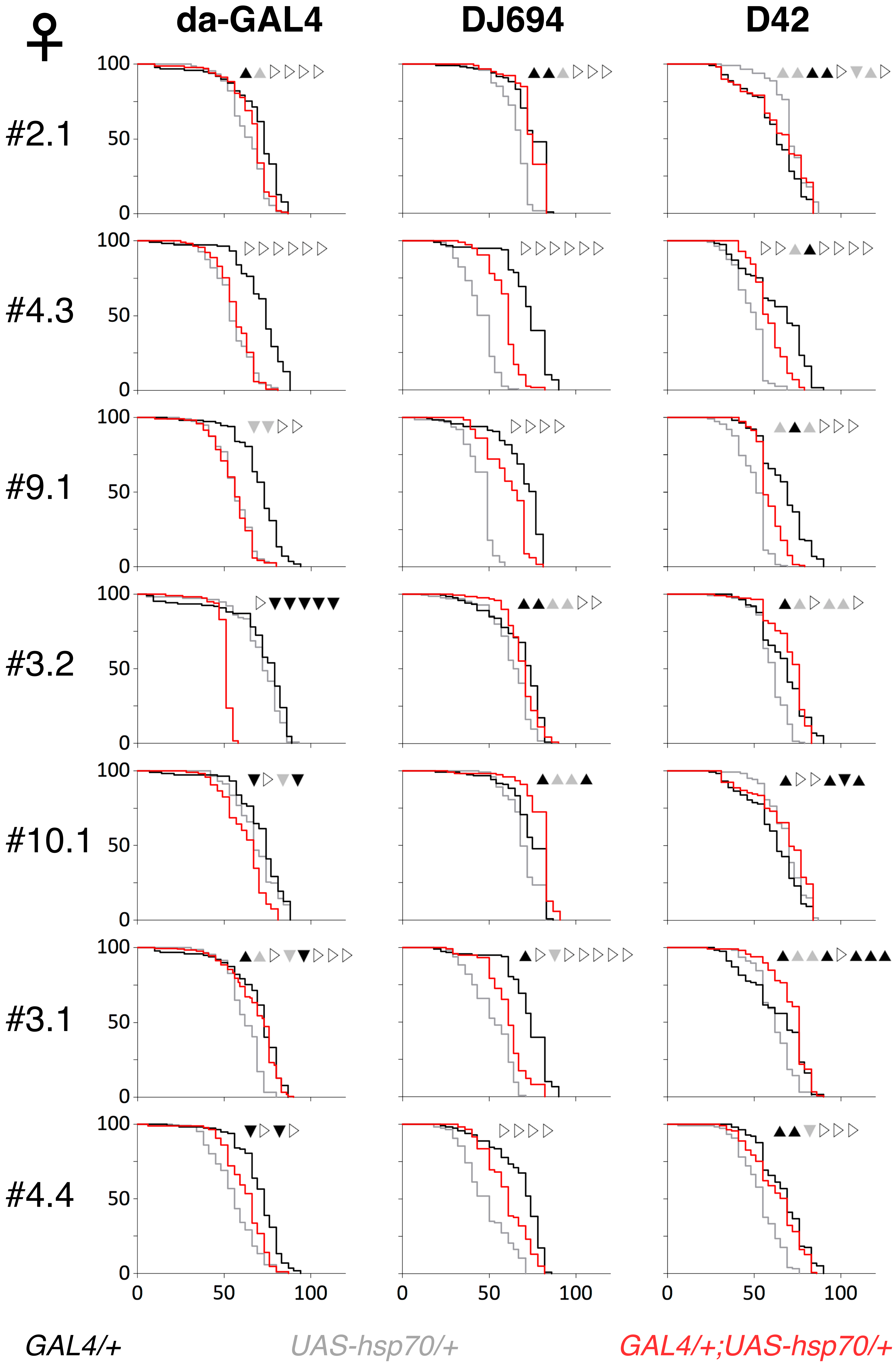


**Figure S3: Longevity of females overexpressing HSP70.** Typical longevity curves (x axis: days at 25˚C; y axis: percentage alive) are shown. The outcome of the statistical comparison (Table S3) between experimental and controls for each experimental replicate is provided at the top of each graph: ▲: significant increase ▲: increase ▼: significant decrease ▼: decrease ▷: no change.
