## Supplementary material for "Expression of heat shock protein 70 is insufficient to extend *Drosophila melanogaster* longevity": Figure S4

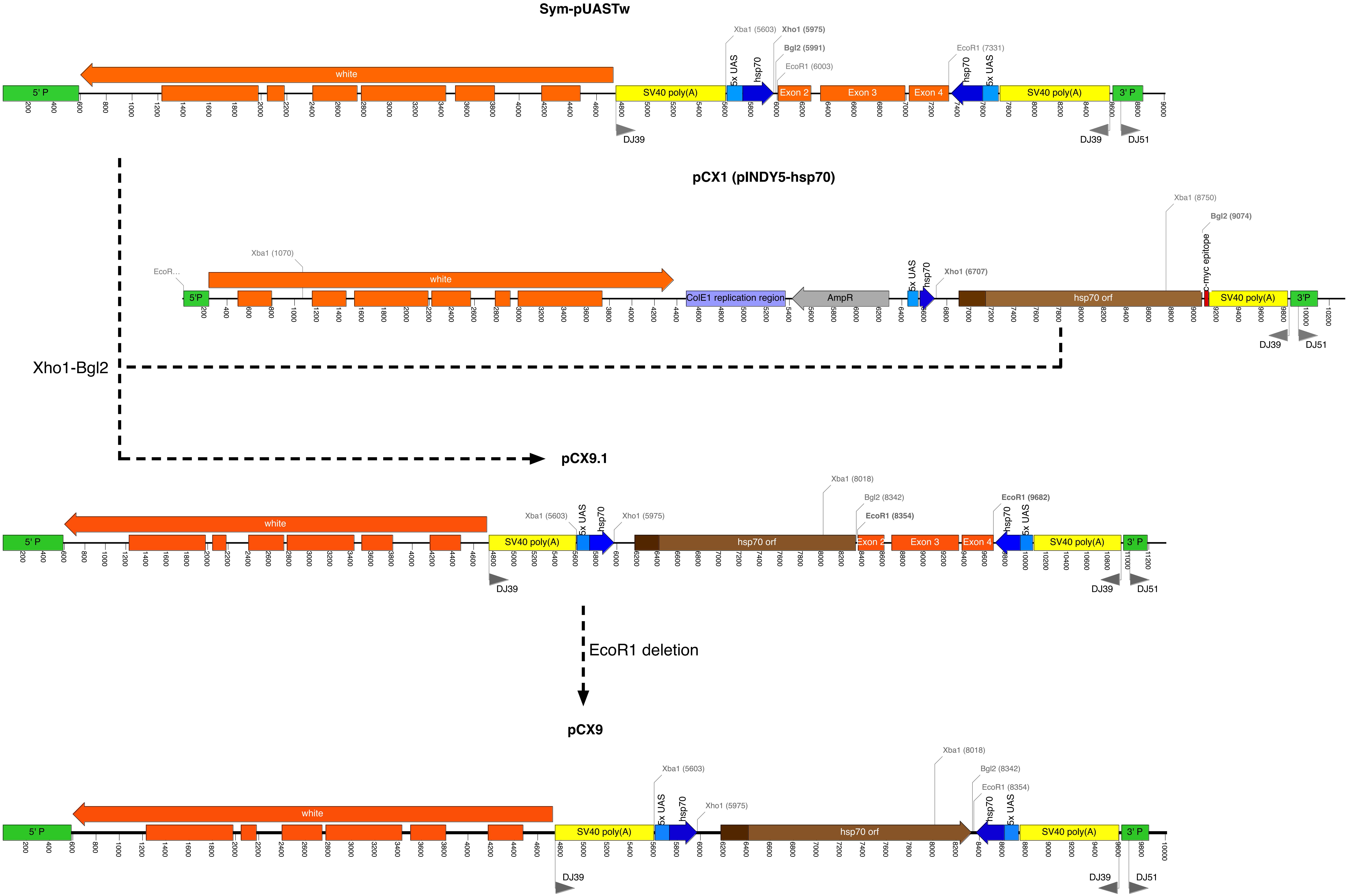


**Figure S4. Construction of pCX9 plasmid.** The *hsp70Ab* coding sequence (excised as a Xho*I*-Bgl*II* fragment from pCX1 (Xiao et al. 2007)) was first inserted into Xho*I*-Bgl*II*-opened Sym-pUASTw (Giordano et al. 2002) to obtain pCX9.1. The mini-*white* sequences from Sym-pUASTw were then removed by digestion with EcoRI and self-ligation to generate pCX9. Grey arrows show the binding sites for the oligonucleotides (DJ39, DJ51) used for inverse PCR.
