## Supplementary material for "Expression of heat shock protein 70 is insufficient to extend *Drosophila melanogaster* longevity": Figure S5

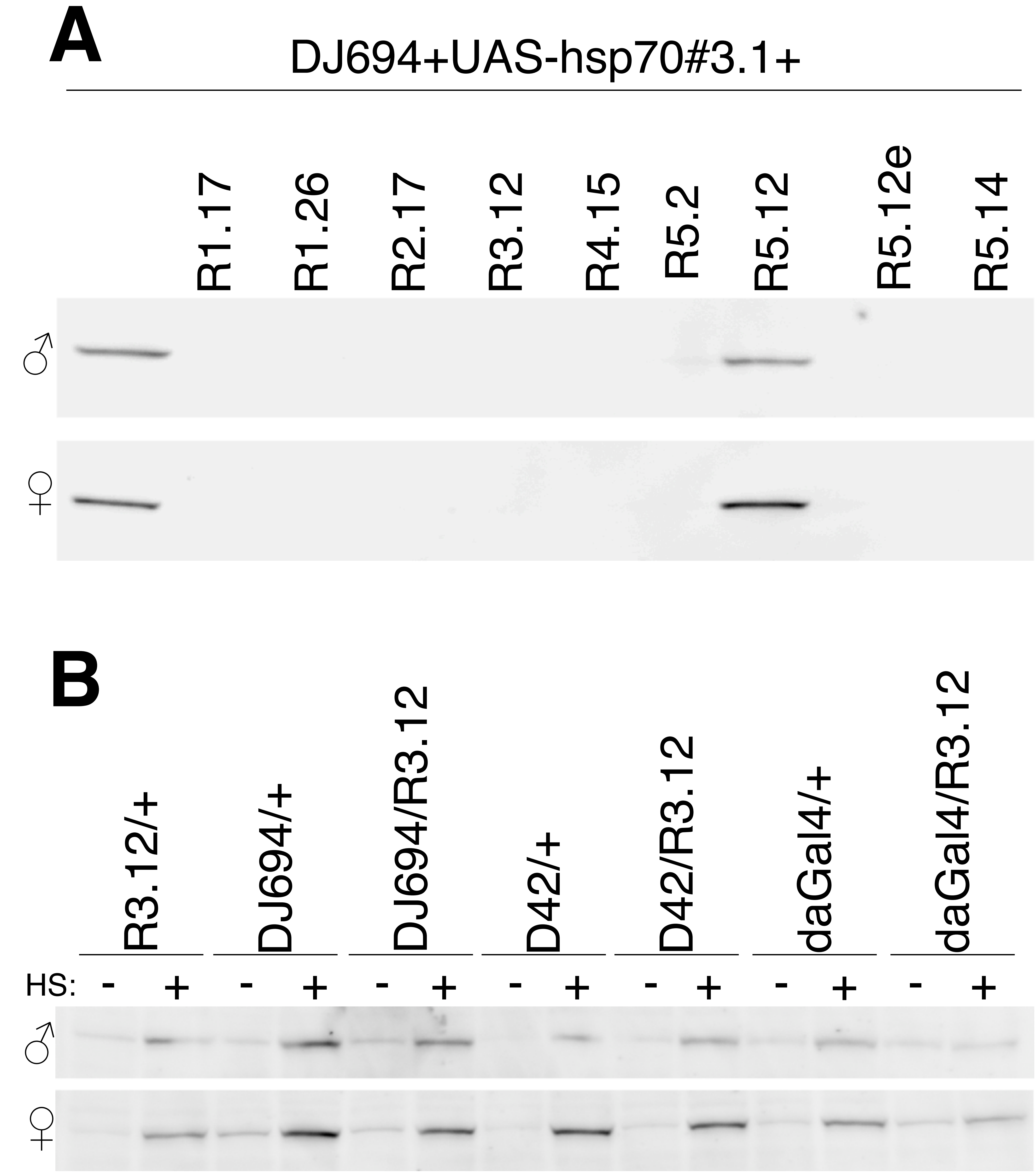


**Figure S5. Down-regulation of Hsp70 by RNAi.** (A) Inhibition of Hsp70-myc: Western-blot with monoclonal antibody 9E10 of 5-7 days old extracts. (B) Inhibition of endogenous Hsp70: Western-blot with monoclonal antibody 5A5 of 7 days old extracts. HS: heat shock, -:without, +:with.
