## Supplementary material for "Expression of heat shock protein 70 is insufficient to extend *Drosophila melanogaster* longevity": Table S1



| Gal4 | UAS | Stage | UAS | | | Gal4 | | | Gal4 + UAS | | | P | | Change |
|---|---|---|---|---|---|---|---|---|---|---|---|---|---|---|
| | | | N | Mean | STD | N | Mean | STD | N | Mean | STD | vs UAS | vs Gal4 | |
| da-GAL4 | 2.1 | L1 | 8 | 98.0 | 2.1 | 8 | 97.0 | 2.8 | 8 | 93.0 | 4.7 | 0.0154 | 0.0568 | |
| | | Pupa | 8 | 82.6 | 8.9 | 8 | 93.9 | 3.8 | 8 | 78.7 | 14.0 | 0.5202 | 0.0105 | |
| | | Adult | 8 | 98.6 | 2.5 | 8 | 98.9 | 3.1 | 8 | 99.3 | 1.9 | 0.5372 | 0.7406 | |
| da-GAL4 | 4.3 | L1 | 8 | 94.5 | 3.7 | 8 | 97.0 | 2.8 | 8 | 96.0 | 4.3 | 0.4637 | 0.5899 | |
| | | Pupa | 8 | 78.7 | 13.4 | 8 | 93.9 | 3.8 | 8 | 84.9 | 5.0 | 0.2411 | 0.0012 | |
| | | Adult | 8 | 97.0 | 3.3 | 8 | 98.9 | 3.1 | 8 | 94.1 | 8.6 | 0.3900 | 0.1545 | |
| da-GAL4 | 9.1 | L1 | 8 | 97.0 | 3.5 | 8 | 97.0 | 2.8 | 8 | 98.5 | 2.1 | 0.3190 | 0.2462 | |
| | | Pupa | 8 | 80.4 | 3.2 | 8 | 93.9 | 3.8 | 8 | **76.1** | 4.1 | 0.0359 | 0.0000 | ↓ |
| | | Adult | 8 | 98.8 | 3.5 | 8 | 98.9 | 3.1 | 8 | 99.3 | 2.0 | 0.7035 | 0.7654 | |
| da-GAL4 | 3.1 | L1 | 8 | 94.5 | 5.2 | 8 | 97.0 | 2.8 | 8 | 95.0 | 5.1 | 0.8494 | 0.3504 | |
| | | Pupa | 8 | 84.3 | 4.8 | 8 | 93.9 | 3.8 | 8 | 85.9 | 7.5 | 0.6152 | 0.0181 | |
| | | Adult | 8 | 98.2 | 3.6 | 8 | 98.9 | 3.1 | 8 | 98.2 | 2.5 | 0.9984 | 0.5996 | |
| da-GAL4 | 4.4 | L1 | 8 | 98.0 | 3.7 | 8 | 97.0 | 2.8 | 8 | 96.0 | 4.8 | 0.3654 | 0.6186 | |
| | | Pupa | 8 | 91.3 | 8.7 | 8 | 93.9 | 3.8 | 8 | 96.9 | 2.9 | 0.1070 | 0.0921 | |
| | | Adult | 8 | 96.2 | 5.7 | 8 | 98.9 | 3.1 | 8 | 95.6 | 6.2 | 0.8593 | 0.1974 | |
| da-GAL4 | 3.2/CyO | L1 | 16 | 96.8 | 3.3 | 8 | 97.0 | 2.8 | 16 | 95.8 | 3.1 | 0.3940 | 0.3476 | |
| | | Pupa | 16 | 94.8 | 4.6 | 8 | 93.9 | 3.8 | 16 | 92.0 | 6.4 | 0.2323 | 0.4569 | |
| | | Adult | 16 | 46.3 | 11.0 | 8 | 98.9 | 3.1 | 16 | 43.5 | 10.9 | 0.6573 | NA | |
| da-GAL4 | 10.1/CyO | L1 | 16 | 97.0 | 4.0 | 8 | 97.0 | 2.8 | 16 | 94.3 | 4.4 | 0.0733 | 0.1220 | |
| | | Pupa | 16 | 84.0 | 8.2 | 8 | 93.9 | 3.8 | 16 | 90.5 | 6.1 | 0.0154 | 0.1679 | |
| | | Adult | 16 | 48.6 | 10.5 | 8 | 98.9 | 3.1 | 16 | 57.5 | 18.2 | 0.1006 | NA | |
| DJ694 | 2.1 | L1 | 8 | 95.0 | 4.7 | 8 | 80.5 | 9.2 | 8 | 78.0 | 8.0 | 0.0001 | 0.5707 | |
| | | Pupa | 8 | 93.8 | 5.1 | 8 | 89.7 | 7.0 | 8 | 81.0 | 10.6 | 0.0086 | 0.0743 | |
| | | Adult | 8 | 98.2 | 2.4 | 8 | 99.2 | 2.2 | 8 | 95.4 | 6.9 | 0.2865 | 0.1563 | |
| DJ694 | 4.3 | L1 | 8 | 95.0 | 5.1 | 8 | 80.5 | 9.2 | 8 | 80.0 | 12.1 | 0.0061 | 0.9271 | |
| | | Pupa | 8 | 92.7 | 3.6 | 8 | 89.7 | 7.0 | 8 | **70.8** | 15.9 | 0.0020 | 0.0083 | ↓ |
| | | Adult | 8 | 94.8 | 3.8 | 8 | 99.2 | 2.2 | 8 | 96.8 | 3.4 | 0.2968 | 0.1170 | |
| DJ694 | 9.1 | L1 | 8 | 98.0 | 3.0 | 8 | 80.5 | 9.2 | 8 | **65.0** | 10.0 | 0.0000 | 0.0060 | ↓ |
| | | Pupa | 8 | 90.6 | 9.0 | 8 | 89.7 | 7.0 | 8 | 77.5 | 17.3 | 0.0776 | 0.0849 | |
| | | Adult | 8 | 98.4 | 3.0 | 8 | 99.2 | 2.2 | 8 | 99.0 | 2.9 | 0.7306 | 0.8444 | |
| DJ694 | 3.1 | L1 | 8 | 94.5 | 6.0 | 8 | 80.5 | 9.2 | 8 | 78.5 | 8.3 | 0.0006 | 0.6540 | |
| | | Pupa | 8 | 83.0 | 13.1 | 8 | 89.7 | 7.0 | 8 | 74.4 | 6.1 | 0.1123 | 0.0004 | |
| | | Adult | 8 | 96.0 | 4.8 | 8 | 99.2 | 2.2 | 8 | 96.3 | 6.0 | 0.9216 | 0.2168 | |
| DJ694 | 4.4 | L1 | 8 | 96.0 | 3.0 | 8 | 80.5 | 9.2 | 8 | 87.5 | 10.1 | 0.0392 | 0.1695 | |
| | | Pupa | 8 | 95.8 | 5.1 | 8 | 89.7 | 7.0 | 8 | 93.2 | 3.8 | 0.2667 | 0.2318 | |
| | | Adult | 8 | 94.5 | 5.1 | 8 | 99.2 | 2.2 | 8 | 95.6 | 3.2 | 0.6213 | 0.0182 | |
| DJ694 | 3.2/CyO | L1 | 16 | 94.8 | 5.2 | 8 | 80.5 | 9.2 | 16 | 78.0 | 7.9 | 0.0000 | 0.4943 | |
| | | Pupa | 16 | 94.5 | 4.9 | 8 | 89.7 | 7.0 | 16 | 85.9 | 11.2 | 0.0161 | 0.3970 | |
| | | Adult | 16 | 53.9 | 12.9 | 8 | 99.2 | 2.2 | 16 | 50.2 | 8.5 | 0.3201 | NA | |
| DJ694 | 10.1/CyO | L1 | 16 | 96.5 | 3.2 | 8 | 80.5 | 9.2 | 16 | **61.3** | 11.5 | 0.0000 | 0.0005 | ↓ |
| | | Pupa | 16 | 91.0 | 6.1 | 8 | 89.7 | 7.0 | 16 | 94.2 | 6.2 | 0.1452 | 0.1227 | |
| | | Adult | 16 | 56.7 | 9.1 | 8 | 99.2 | 2.2 | 16 | 56.6 | 10.1 | 0.9958 | NA | |
| D42 | 2.1 | L1 | 8 | 91.5 | 8.9 | 8 | 96.5 | 2.6 | 8 | 94.0 | 4.8 | 0.4965 | 0.2134 | |
| | | Pupa | 8 | 83.1 | 2.1 | 8 | 91.2 | 6.7 | 8 | 87.3 | 8.6 | 0.2012 | 0.3266 | |
| | | Adult | 8 | 98.1 | 2.6 | 8 | 96.7 | 3.8 | 8 | 96.9 | 4.5 | 0.5063 | 0.9423 | |
| D42 | 4.3 | L1 | 8 | 96.5 | 3.3 | 8 | 96.5 | 2.6 | 8 | 95.5 | 4.5 | 0.6217 | 0.5938 | |
| | | Pupa | 8 | 89.7 | 9.5 | 8 | 91.2 | 6.7 | 8 | 90.7 | 7.3 | 0.8189 | 0.8912 | |
| | | Adult | 8 | 95.9 | 4.4 | 8 | 96.7 | 3.8 | 8 | 97.7 | 3.7 | 0.4073 | 0.6179 | |

**Table S1**

| Gal4 | UAS | Stage | UAS | | | Gal4 | | | Gal4 + UAS | | | P | | Change |
|---|---|---|---|---|---|---|---|---|---|---|---|---|---|---|
| | | | N | Mean | STD | N | Mean | STD | N | Mean | STD | vs UAS | vs Gal4 | |
| D42 | 9.1 | L1 | 8 | 96.0 | 4.3 | 8 | 96.5 | 2.6 | 8 | 93.0 | 4.1 | 0.1759 | 0.0615 | |
| | | Pupa | 8 | 91.0 | 6.7 | 8 | 91.2 | 6.7 | 8 | 94.7 | 5.3 | 0.2444 | 0.2664 | |
| | | Adult | 8 | 96.1 | 4.2 | 8 | 96.7 | 3.8 | 8 | 98.9 | 2.0 | 0.1086 | 0.1675 | |
| D42 | 3.1 | L1 | 8 | 93.5 | 6.4 | 8 | 96.5 | 2.6 | 8 | 93.5 | 7.4 | 1.0000 | 0.2962 | |
| | | Pupa | 8 | 92.1 | 5.1 | 8 | 91.2 | 6.7 | 8 | 87.1 | 8.1 | 0.1540 | 0.2859 | |
| | | Adult | 8 | 87.3 | 15.8 | 8 | 96.7 | 3.8 | 8 | 93.8 | 5.3 | 0.2900 | 0.2227 | |
| D42 | 4.4 | L1 | 8 | 97.5 | 3.0 | 8 | 96.5 | 2.6 | 8 | 99.0 | 2.8 | 0.3190 | 0.0852 | |
| | | Pupa | 8 | 74.7 | 16.8 | 8 | 91.2 | 6.7 | 8 | 87.0 | 6.6 | 0.0745 | 0.2258 | |
| | | Adult | 8 | 96.9 | 3.5 | 8 | 96.7 | 3.8 | 8 | 99.4 | 1.8 | 0.1029 | 0.0922 | |
| D42 | 3.2/CyO | L1 | 16 | 96.3 | 5.4 | 8 | 96.5 | 2.6 | 16 | 93.0 | 3.7 | 0.0690 | 0.0265 | |
| | | Pupa | 16 | 87.1 | 7.8 | 8 | 91.2 | 6.7 | 16 | 88.4 | 7.2 | 0.5426 | 0.3694 | |
| | | Adult | 16 | 48.2 | 6.5 | 8 | 96.7 | 3.8 | 16 | 46.7 | 6.2 | 0.4143 | NA | |
| D42 | 10.1/CyO | L1 | 16 | 97.3 | 3.5 | 8 | 96.5 | 2.6 | 16 | 98.0 | 2.9 | 0.5150 | 0.2310 | |
| | | Pupa | 16 | 89.0 | 6.4 | 8 | 91.2 | 6.7 | 16 | 94.9 | 5.1 | 0.0069 | 0.1423 | |
| | | Adult | 16 | 39.1 | 6.6 | 8 | 96.7 | 3.8 | 16 | 39.0 | 5.1 | 0.9748 | NA | |

**Table S1. Developmental analysis of Hsp70 overexpression.** Percentage hatching (L1), pupation (Pupa) and eclosion (Adult) are compared between experimental animals (Gal4+UAS) and control animals missing the driver (UAS) or UAS (Gal4) transgene (P: Kruskal-Wallis test). The first columns indicate the genotype of the mother while the second column indicate the genotype of the father used to generate the experimental and control animals. The change column indicates experimental animals that are significantly ($P<0.05$) lower than both controls. N: number of independent measurements, STD: Standard deviation.
