## Supplementary material for "Expression of heat shock protein 70 is insufficient to extend *Drosophila melanogaster* longevity": Table S2

| Stress | Age | Gal4 | UAS | Sex | Rep[1] | UAS | | | Gal4 | | | Gal4 + UAS | | | % Extension | | P |
|---|---|---|---|---|---|---|---|---|---|---|---|---|---|---|---|---|---|
| | | | | | | n | Mean | Max[2] | n | Mean | Max[2] | n | Mean | Max[2] | Mean | Max[2] | |
| Heat | 3 | daGal4 | 2.1 | ♀ | 1 | 37 | 72.0 | 84 | 39 | 61.7 | 84 | 39 | 63.8 | 93 | 0.0 | 10.7 | |
| Heat | 3 | daGal4 | 2.1 | ♀ | 2 | 41 | 64.9 | 76 | 39 | 60.1 | 84 | 40 | 60.6 | 84 | 0.0 | 0.0 | |
| Heat | 3 | daGal4 | 2.1 | ♀ | 3 | 42 | 59.0 | 76 | 38 | 54.1 | 76 | 39 | 59.7 | 76 | 1.3 | 0.0 | **0.0048** |
| Heat | 3 | daGal4 | 2.1 | ♀ | 4 | 40 | 24.9 | 41 | 40 | 22.3 | 41 | 40 | 27.8 | 45 | 11.9 | 9.8 | **0.0139** |
| Heat | 3 | daGal4 | 2.1 | ♂ | 1 | 38 | 61.4 | 84 | 39 | 60.8 | 84 | 40 | 57.8 | 76 | -5.0 | -9.5 | 0.0974 |
| Heat | 3 | daGal4 | 2.1 | ♂ | 2 | 40 | 58.9 | 68 | 38 | 53.9 | 84 | 40 | 52.5 | 68 | -2.6 | 0.0 | **0.0082** |
| Heat | 3 | daGal4 | 2.1 | ♂ | 3 | 40 | 51.4 | 76 | 38 | 52.9 | 76 | 40 | 53.0 | 76 | 0.1 | 0.0 | 0.6036 |
| Heat | 3 | daGal4 | 2.1 | ♂ | 4 | 40 | 23.5 | 34 | 40 | 23.6 | 45 | 40 | 25.8 | 41 | 9.7 | 0.0 | 0.1872 |
| Heat | 3 | daGal4 | 4.3 | ♀ | 1 | 40 | 68.0 | 84 | 39 | 61.7 | 84 | 38 | 66.8 | 93 | 0.0 | 10.7 | |
| Heat | 3 | daGal4 | 4.3 | ♀ | 2 | 40 | 60.7 | 76 | 39 | 60.1 | 84 | 39 | 56.3 | 68 | -6.3 | -10.5 | **0.0030** |
| Heat | 3 | daGal4 | 4.3 | ♀ | 3 | 41 | 51.1 | 56 | 38 | 54.1 | 76 | 38 | 57.9 | 76 | 7.0 | 0.0 | **0.0450** |
| Heat | 3 | daGal4 | 4.3 | ♀ | 4 | 39 | 28.1 | 36 | 40 | 27.9 | 36 | 40 | 33.6 | 46 | 19.5 | 27.8 | **0.0005** |
| Heat | 3 | daGal4 | 4.3 | ♂ | 1 | 41 | 63.7 | 76 | 39 | 60.8 | 84 | 45 | 58.6 | 76 | -3.7 | 0.0 | **0.0175** |
| Heat | 3 | daGal4 | 4.3 | ♂ | 2 | 42 | 55.5 | 76 | 38 | 53.9 | 84 | 46 | 51.3 | 64 | -4.8 | -15.8 | **0.0332** |
| Heat | 3 | daGal4 | 4.3 | ♂ | 3 | 41 | 51.9 | 76 | 38 | 52.9 | 76 | 40 | 52.1 | 68 | 0.0 | -10.5 | |
| Heat | 3 | daGal4 | 4.3 | ♂ | 4 | 40 | 25.5 | 36 | 40 | 29.5 | 41 | 40 | 29.8 | 41 | 1.1 | 0.0 | **0.0017** |
| Heat | 3 | daGal4 | 9.1 | ♀ | 1 | 39 | 69.1 | 93 | 39 | 61.7 | 84 | 40 | 67.0 | 84 | 0.0 | 0.0 | |
| Heat | 3 | daGal4 | 9.1 | ♀ | 2 | 41 | 61.6 | 76 | 39 | 60.1 | 84 | 38 | 65.7 | 76 | 6.7 | 0.0 | **0.0375** |
| Heat | 3 | daGal4 | 9.1 | ♀ | 3 | 43 | 60.7 | 76 | 38 | 54.1 | 76 | 39 | 62.2 | 84 | 2.3 | 10.5 | **0.0002** |
| Heat | 3 | daGal4 | 9.1 | ♂ | 1 | 39 | 68.7 | 84 | 39 | 60.8 | 84 | 38 | 69.7 | 93 | 1.5 | 10.7 | **0.0079** |
| Heat | 3 | daGal4 | 9.1 | ♂ | 2 | 40 | 57.7 | 76 | 38 | 53.9 | 84 | 34 | 60.8 | 76 | 5.4 | 0.0 | 0.1188 |
| Heat | 3 | daGal4 | 9.1 | ♂ | 3 | 40 | 55.4 | 76 | 38 | 52.9 | 76 | 40 | 58.3 | 84 | 5.3 | 10.5 | **0.0352** |
| Heat | 3 | daGal4 | 3.2 | ♀ | 1 | 40 | 60.1 | 76 | 39 | 61.7 | 84 | 42 | 60.0 | 76 | -0.2 | 0.0 | 0.3989 |
| Heat | 3 | daGal4 | 3.2 | ♀ | 2 | 39 | 52.8 | 68 | 39 | 60.1 | 84 | 38 | 61.8 | 76 | 2.8 | 0.0 | 0.6183 |
| Heat | 3 | daGal4 | 3.2 | ♀ | 3 | 40 | 50.2 | 56 | 38 | 54.1 | 76 | 39 | 54.3 | 76 | 0.3 | 0.0 | **0.0010** |
| Heat | 3 | daGal4 | 3.2 | ♀ | 4 | 39 | 31.2 | 42 | 38 | 23.3 | 38 | 40 | 40.4 | 46 | 29.7 | 9.5 | **<0.0001** |
| Heat | 3 | daGal4 | 3.2 | ♂ | 1 | 40 | 60.1 | 93 | 39 | 60.8 | 84 | 39 | 58.2 | 76 | -3.2 | -9.5 | 0.3003 |
| Heat | 3 | daGal4 | 3.2 | ♂ | 2 | 45 | 46.5 | 64 | 38 | 53.9 | 84 | 40 | 57.3 | 76 | 6.3 | 0.0 | 0.4225 |
| Heat | 3 | daGal4 | 3.2 | ♂ | 3 | 40 | 47.4 | 60 | 38 | 52.9 | 76 | 40 | 49.4 | 76 | 0.0 | 0.0 | |
| Heat | 3 | daGal4 | 3.2 | ♂ | 4 | 36 | 21.3 | 30 | 40 | 19.5 | 30 | 40 | 27.0 | 34 | 26.6 | 13.3 | **<0.0001** |
| Heat | 3 | daGal4 | 10.1 | ♀ | 1 | 40 | 61.5 | 84 | 39 | 61.7 | 84 | 40 | 61.5 | 76 | 0.0 | -9.5 | |
| Heat | 3 | daGal4 | 10.1 | ♀ | 2 | 40 | 50.7 | 56 | 39 | 60.1 | 84 | 38 | 58.4 | 76 | 0.0 | 0.0 | |
| Heat | 3 | daGal4 | 10.1 | ♀ | 3 | 40 | 50.4 | 56 | 38 | 54.1 | 76 | 40 | 63.5 | 93 | 17.3 | 22.4 | **<0.0001** |
| Heat | 3 | daGal4 | 10.1 | ♀ | 4 | 39 | 32.5 | 41 | 40 | 28.4 | 36 | 40 | 33.5 | 41 | 3.2 | 0.0 | 0.7430 |
| Heat | 3 | daGal4 | 10.1 | ♂ | 1 | 29 | 61.8 | 84 | 39 | 60.8 | 84 | 40 | 60.4 | 84 | -0.7 | 0.0 | 0.4536 |
| Heat | 3 | daGal4 | 10.1 | ♂ | 2 | 40 | 46.4 | 52 | 38 | 53.9 | 84 | 39 | 56.4 | 76 | 4.7 | 0.0 | 0.4541 |
| Heat | 3 | daGal4 | 10.1 | ♂ | 3 | 39 | 48.3 | 60 | 38 | 52.9 | 76 | 39 | 53.9 | 76 | 1.9 | 0.0 | **0.0027** |
| Heat | 3 | daGal4 | 10.1 | ♂ | 4 | 40 | 26.2 | 36 | 40 | 22.0 | 26 | 40 | 28.8 | 36 | 9.9 | 0.0 | **0.0218** |
| Heat | 3 | daGal4 | 3.1 | ♀ | 1 | 40 | 59.0 | 76 | 39 | 61.7 | 84 | 43 | 58.0 | 76 | -1.8 | 0.0 | 0.0565 |
| Heat | 3 | daGal4 | 3.1 | ♀ | 2 | 41 | 50.4 | 60 | 39 | 60.1 | 84 | 39 | 53.8 | 76 | 0.0 | 0.0 | |
| Heat | 3 | daGal4 | 3.1 | ♀ | 3 | 38 | 46.2 | 56 | 38 | 54.1 | 76 | 41 | 52.0 | 64 | 0.0 | 0.0 | |
| Heat | 3 | daGal4 | 3.1 | ♀ | 4 | 38 | 22.6 | 34 | 38 | 23.3 | 38 | 40 | 25.9 | 38 | 11.4 | 0.0 | **0.0067** |
| Heat | 3 | daGal4 | 3.1 | ♂ | 1 | 39 | 61.9 | 76 | 39 | 60.8 | 84 | 40 | 61.0 | 76 | 0.0 | 0.0 | |
| Heat | 3 | daGal4 | 3.1 | ♂ | 2 | 40 | 42.6 | 56 | 38 | 53.9 | 84 | 40 | 54.9 | 76 | 1.9 | 0.0 | 0.9906 |
| Heat | 3 | daGal4 | 3.1 | ♂ | 3 | 18 | 52.2 | 76 | 38 | 52.9 | 76 | 39 | 51.6 | 68 | -1.2 | -10.5 | 0.3324 |
| Heat | 3 | daGal4 | 3.1 | ♂ | 4 | 40 | 16.5 | 26 | 40 | 19.5 | 30 | 40 | 20.1 | 26 | 2.8 | 0.0 | 0.4985 |
| Heat | 3 | daGal4 | 4.4 | ♀ | 1 | 41 | 67.1 | 84 | 39 | 61.7 | 84 | 38 | 60.3 | 76 | -2.2 | -9.5 | **0.0014** |
| Heat | 3 | daGal4 | 4.4 | ♀ | 2 | 33 | 60.8 | 76 | 39 | 60.1 | 84 | 44 | 59.7 | 93 | -0.7 | 10.7 | 0.6783 |
| Heat | 3 | daGal4 | 4.4 | ♀ | 3 | 40 | 59.3 | 76 | 38 | 54.1 | 76 | 40 | 55.6 | 76 | 0.0 | 0.0 | |

### Table S2



| Stress | Age | Gal4 | UAS | Sex | Rep[1] | UAS | | | Gal4 | | | Gal4 + UAS | | | % Extension | | *P* |
|---|---|---|---|---|---|---|---|---|---|---|---|---|---|---|---|---|---|
| | | | | | | n | Mean | Max[2] | n | Mean | Max[2] | n | Mean | Max[2] | Mean | Max[2] | |
| Heat | 3 | daGal4 | 4.4 | ♀ | 4 | 40 | 32.2 | 41 | 40 | 27.9 | 36 | 40 | 38.0 | 46 | 18.0 | 12.2 | **0.0002** |
| Heat | 3 | daGal4 | 4.4 | ♂ | 1 | 43 | 69.9 | 84 | 39 | 60.8 | 84 | 42 | 60.5 | 76 | -0.6 | -9.5 | 0.3669 |
| Heat | 3 | daGal4 | 4.4 | ♂ | 2 | 40 | 56.2 | 76 | 38 | 53.9 | 84 | 41 | 53.8 | 84 | -0.2 | 0.0 | 0.5348 |
| Heat | 3 | daGal4 | 4.4 | ♂ | 3 | 38 | 55.2 | 76 | 38 | 52.9 | 76 | 40 | 52.7 | 68 | -0.5 | -10.5 | 0.2226 |
| Heat | 3 | daGal4 | 4.4 | ♂ | 4 | 40 | 31.3 | 36 | 40 | 29.5 | 41 | 40 | 34.9 | 41 | 11.5 | 0.0 | **0.0003** |
| Heat | 3 | DJ694 | 2.1 | ♀ | 1 | 37 | 72.0 | 84 | 39 | 69.4 | 93 | 39 | 67.3 | 93 | -3.0 | 0.0 | **0.0434** |
| Heat | 3 | DJ694 | 2.1 | ♀ | 2 | 41 | 64.9 | 76 | 31 | 60.9 | 76 | 46 | 60.4 | 84 | -0.9 | 10.5 | 0.0525 |
| Heat | 3 | DJ694 | 2.1 | ♀ | 3 | 42 | 59.0 | 76 | 37 | 65.3 | 84 | 43 | 56.8 | 76 | -3.6 | 0.0 | **0.0028** |
| Heat | 3 | DJ694 | 2.1 | ♀ | 4 | 40 | 24.9 | 41 | 39 | 24.5 | 41 | 40 | 18.7 | 30 | -23.7 | -26.8 | **<0.0001** |
| Heat | 3 | DJ694 | 2.1 | ♂ | 1 | 38 | 61.4 | 84 | 37 | 63.2 | 84 | 40 | 52.4 | 76 | -14.6 | -9.5 | **<0.0001** |
| Heat | 3 | DJ694 | 2.1 | ♂ | 2 | 40 | 58.9 | 68 | 33 | 58.4 | 76 | 40 | 52.3 | 76 | -10.5 | 0.0 | **0.0039** |
| Heat | 3 | DJ694 | 2.1 | ♂ | 3 | 40 | 51.4 | 76 | 35 | 54.1 | 76 | 42 | 55.1 | 76 | 2.0 | 0.0 | 0.1223 |
| Heat | 3 | DJ694 | 2.1 | ♂ | 4 | 40 | 23.5 | 34 | 40 | 21.4 | 34 | 40 | 21.9 | 41 | 0.0 | 20.6 | |
| Heat | 3 | DJ694 | 4.3 | ♀ | 1 | 40 | 68.0 | 84 | 39 | 69.4 | 93 | 40 | 63.5 | 76 | -6.6 | -9.5 | **0.0141** |
| Heat | 3 | DJ694 | 4.3 | ♀ | 2 | 40 | 60.7 | 76 | 31 | 60.9 | 76 | 36 | 60.6 | 76 | -0.2 | 0.0 | 0.4922 |
| Heat | 3 | DJ694 | 4.3 | ♀ | 3 | 41 | 51.1 | 56 | 37 | 65.3 | 84 | 41 | 52.2 | 76 | 0.0 | 0.0 | |
| Heat | 3 | DJ694 | 4.3 | ♀ | 4 | 39 | 28.1 | 36 | 40 | 24.9 | 36 | 40 | 22.6 | 41 | -9.3 | 13.9 | **0.0106** |
| Heat | 3 | DJ694 | 4.3 | ♂ | 1 | 41 | 63.7 | 76 | 37 | 63.2 | 84 | 38 | 55.6 | 76 | -12.0 | 0.0 | **0.0026** |
| Heat | 3 | DJ694 | 4.3 | ♂ | 2 | 42 | 55.5 | 76 | 33 | 58.4 | 76 | 39 | 54.1 | 76 | -2.7 | 0.0 | 0.0694 |
| Heat | 3 | DJ694 | 4.3 | ♂ | 3 | 41 | 51.9 | 76 | 35 | 54.1 | 76 | 41 | 43.6 | 48 | -16.0 | -36.8 | **<0.0001** |
| Heat | 3 | DJ694 | 4.3 | ♂ | 4 | 40 | 25.5 | 36 | 39 | 29.6 | 36 | 40 | 27.1 | 36 | 0.0 | 0.0 | |
| Heat | 3 | DJ694 | 9.1 | ♀ | 1 | 39 | 69.1 | 93 | 39 | 69.4 | 93 | 41 | 61.1 | 84 | -11.6 | -9.7 | **0.0056** |
| Heat | 3 | DJ694 | 9.1 | ♀ | 2 | 41 | 61.6 | 76 | 31 | 60.9 | 76 | 40 | 58.8 | 76 | -3.5 | 0.0 | 0.2002 |
| Heat | 3 | DJ694 | 9.1 | ♀ | 3 | 43 | 60.7 | 76 | 37 | 65.3 | 84 | 39 | 46.3 | 52 | -23.9 | -31.6 | **<0.0001** |
| Heat | 3 | DJ694 | 9.1 | ♂ | 1 | 39 | 68.7 | 84 | 37 | 63.2 | 84 | 40 | 52.1 | 64 | -17.5 | -23.8 | **<0.0001** |
| Heat | 3 | DJ694 | 9.1 | ♂ | 2 | 40 | 57.7 | 76 | 33 | 58.4 | 76 | 38 | 44.2 | 68 | -23.4 | -10.5 | **<0.0001** |
| Heat | 3 | DJ694 | 9.1 | ♂ | 3 | 40 | 55.4 | 76 | 35 | 54.1 | 76 | 30 | 48.4 | 64 | -10.5 | -15.8 | **0.0051** |
| Heat | 3 | DJ694 | 3.2 | ♀ | 1 | 40 | 60.1 | 76 | 39 | 69.4 | 93 | 40 | 54.5 | 60 | -9.3 | -21.1 | **<0.0001** |
| Heat | 3 | DJ694 | 3.2 | ♀ | 2 | 39 | 52.8 | 68 | 31 | 60.9 | 76 | 40 | 52.5 | 56 | -0.6 | -17.6 | 0.9298 |
| Heat | 3 | DJ694 | 3.2 | ♀ | 3 | 40 | 50.2 | 56 | 37 | 65.3 | 84 | 33 | 51.9 | 56 | 0.0 | 0.0 | |
| Heat | 3 | DJ694 | 3.2 | ♀ | 4 | 39 | 31.2 | 42 | 39 | 17.7 | 22 | 40 | 37.4 | 46 | 20.0 | 9.5 | **<0.0001** |
| Heat | 3 | DJ694 | 3.2 | ♂ | 1 | 40 | 60.1 | 93 | 37 | 63.2 | 84 | 39 | 47.8 | 64 | -20.4 | -23.8 | **<0.0001** |
| Heat | 3 | DJ694 | 3.2 | ♂ | 2 | 45 | 46.5 | 64 | 33 | 58.4 | 76 | 39 | 40.4 | 48 | -13.1 | -25.0 | **<0.0001** |
| Heat | 3 | DJ694 | 3.2 | ♂ | 3 | 40 | 47.4 | 60 | 35 | 54.1 | 76 | 39 | 45.4 | 56 | -4.1 | -6.7 | **0.0004** |
| Heat | 3 | DJ694 | 3.2 | ♂ | 4 | 36 | 21.3 | 30 | 40 | 18.1 | 30 | 40 | 21.6 | 30 | 1.4 | 0.0 | **0.0002** |
| Heat | 3 | DJ694 | 10.1 | ♀ | 1 | 40 | 61.5 | 84 | 39 | 69.4 | 93 | 40 | 55.6 | 64 | -9.6 | -23.8 | **0.0010** |
| Heat | 3 | DJ694 | 10.1 | ♀ | 2 | 40 | 50.7 | 56 | 31 | 60.9 | 76 | 40 | 49.9 | 56 | -1.6 | 0.0 | 0.2561 |
| Heat | 3 | DJ694 | 10.1 | ♀ | 3 | 40 | 50.4 | 56 | 37 | 65.3 | 84 | 40 | 48.8 | 56 | -3.2 | 0.0 | 0.1880 |
| Heat | 3 | DJ694 | 10.1 | ♀ | 4 | 39 | 32.5 | 41 | 40 | 27.3 | 31 | 39 | 32.2 | 36 | 0.0 | 0.0 | |
| Heat | 3 | DJ694 | 10.1 | ♂ | 1 | 29 | 61.8 | 84 | 37 | 63.2 | 84 | 39 | 48.2 | 60 | -22.0 | -28.6 | **<0.0001** |
| Heat | 3 | DJ694 | 10.1 | ♂ | 2 | 40 | 46.4 | 52 | 33 | 58.4 | 76 | 40 | 43.3 | 52 | -6.7 | 0.0 | **0.0010** |
| Heat | 3 | DJ694 | 10.1 | ♂ | 3 | 39 | 48.3 | 60 | 35 | 54.1 | 76 | 30 | 47.1 | 52 | -2.6 | -13.3 | **0.0033** |
| Heat | 3 | DJ694 | 10.1 | ♂ | 4 | 40 | 26.2 | 36 | 41 | 22.8 | 31 | 40 | 24.5 | 31 | 0.0 | 0.0 | |
| Heat | 3 | DJ694 | 3.1 | ♀ | 1 | 40 | 59.0 | 76 | 39 | 69.4 | 93 | 41 | 51.6 | 56 | -12.5 | -26.3 | **0.0003** |
| Heat | 3 | DJ694 | 3.1 | ♀ | 2 | 41 | 50.4 | 60 | 31 | 60.9 | 76 | 40 | 46.4 | 52 | -8.0 | -13.3 | **0.0006** |
| Heat | 3 | DJ694 | 3.1 | ♀ | 3 | 38 | 46.2 | 56 | 37 | 65.3 | 84 | 40 | 44.3 | 48 | -4.1 | -14.3 | **0.0063** |
| Heat | 3 | DJ694 | 3.1 | ♀ | 4 | 38 | 22.6 | 34 | 39 | 17.7 | 22 | 40 | 26.7 | 38 | 18.3 | 11.8 | **0.0012** |
| Heat | 3 | DJ694 | 3.1 | ♂ | 1 | 39 | 61.9 | 76 | 37 | 63.2 | 84 | 41 | 49.1 | 52 | -20.8 | -31.6 | **<0.0001** |
| Heat | 3 | DJ694 | 3.1 | ♂ | 2 | 40 | 42.6 | 56 | 33 | 58.4 | 76 | 40 | 41.8 | 44 | -1.9 | -21.4 | 0.6872 |

### Table S2



| Stress | Age | Gal4 | UAS | Sex | Rep[1] | UAS | | | Gal4 | | | Gal4 + UAS | | | % Extension | | *P* |
|---|---|---|---|---|---|---|---|---|---|---|---|---|---|---|---|---|---|
| | | | | | | n | Mean | Max[2] | n | Mean | Max[2] | n | Mean | Max[2] | Mean | Max[2] | |
| Heat | 3 | DJ694 | 3.1 | ♂ | 3 | 18 | 52.2 | 76 | 35 | 54.1 | 76 | 44 | 42.1 | 48 | -19.4 | -36.8 | **0.0004** |
| Heat | 3 | DJ694 | 3.1 | ♂ | 4 | 40 | 16.5 | 26 | 40 | 18.1 | 30 | 40 | 18.3 | 22 | 0.8 | -15.4 | **0.0111** |
| Heat | 3 | DJ694 | 4.4 | ♀ | 1 | 41 | 67.1 | 84 | 39 | 69.4 | 93 | 38 | 66.7 | 84 | -0.7 | 0.0 | 0.2066 |
| Heat | 3 | DJ694 | 4.4 | ♀ | 2 | 33 | 60.8 | 76 | 31 | 60.9 | 76 | 41 | 64.6 | 93 | 6.1 | 22.4 | 0.1122 |
| Heat | 3 | DJ694 | 4.4 | ♀ | 3 | 40 | 59.3 | 76 | 37 | 65.3 | 84 | 39 | 64.7 | 93 | 0.0 | 10.7 | |
| Heat | 3 | DJ694 | 4.4 | ♀ | 4 | 40 | 32.2 | 41 | 40 | 24.9 | 36 | 40 | 26.3 | 36 | 0.0 | 0.0 | |
| Heat | 3 | DJ694 | 4.4 | ♂ | 1 | 43 | 69.9 | 84 | 37 | 63.2 | 84 | 40 | 59.2 | 76 | -6.3 | -9.5 | 0.0892 |
| Heat | 3 | DJ694 | 4.4 | ♂ | 2 | 40 | 56.2 | 76 | 33 | 58.4 | 76 | 43 | 57.5 | 76 | 0.0 | 0.0 | |
| Heat | 3 | DJ694 | 4.4 | ♂ | 3 | 38 | 55.2 | 76 | 35 | 54.1 | 76 | 44 | 60.1 | 76 | 8.9 | 0.0 | **0.0149** |
| Heat | 3 | DJ694 | 4.4 | ♂ | 4 | 40 | 31.3 | 36 | 39 | 29.6 | 36 | 40 | 30.8 | 36 | 0.0 | 0.0 | |
| Heat | 3 | D42 | 2.1 | ♀ | 1 | 37 | 72.0 | 84 | 30 | 63.0 | 93 | 39 | 58.7 | 68 | -6.9 | -19.0 | 0.0854 |
| Heat | 3 | D42 | 2.1 | ♀ | 2 | 41 | 64.9 | 76 | 36 | 54.3 | 76 | 43 | 56.0 | 84 | 0.0 | 10.5 | |
| Heat | 3 | D42 | 2.1 | ♀ | 3 | 42 | 59.0 | 76 | 40 | 62.4 | 93 | 38 | 56.2 | 76 | -4.7 | 0.0 | **0.0024** |
| Heat | 3 | D42 | 2.1 | ♀ | 4 | 40 | 24.9 | 41 | 40 | 19.1 | 30 | 41 | 22.0 | 41 | 0.0 | 0.0 | |
| Heat | 3 | D42 | 2.1 | ♂ | 1 | 38 | 61.4 | 84 | 37 | 58.8 | 76 | 39 | 49.1 | 64 | -16.5 | -15.8 | **<0.0001** |
| Heat | 3 | D42 | 2.1 | ♂ | 2 | 40 | 58.9 | 68 | 32 | 52.0 | 76 | 39 | 45.8 | 64 | -11.8 | -5.9 | **0.0265** |
| Heat | 3 | D42 | 2.1 | ♂ | 3 | 40 | 51.4 | 76 | 39 | 53.9 | 76 | 39 | 49.0 | 64 | -4.6 | -15.8 | **0.0461** |
| Heat | 3 | D42 | 2.1 | ♂ | 4 | 40 | 23.5 | 34 | 39 | 21.0 | 41 | 40 | 20.9 | 41 | -0.8 | 0.0 | 0.1893 |
| Heat | 3 | D42 | 4.3 | ♀ | 1 | 40 | 68.0 | 84 | 30 | 63.0 | 93 | 36 | 66.5 | 84 | 0.0 | 0.0 | |
| Heat | 3 | D42 | 4.3 | ♀ | 2 | 40 | 60.7 | 76 | 36 | 54.3 | 76 | 35 | 59.1 | 76 | 0.0 | 0.0 | |
| Heat | 3 | D42 | 4.3 | ♀ | 3 | 41 | 51.1 | 56 | 40 | 62.4 | 93 | 39 | 61.9 | 76 | 0.0 | 0.0 | |
| Heat | 3 | D42 | 4.3 | ♀ | 4 | 39 | 28.1 | 36 | 40 | 27.5 | 36 | 40 | 30.9 | 46 | 10.0 | 27.8 | **0.0205** |
| Heat | 3 | D42 | 4.3 | ♂ | 1 | 41 | 63.7 | 76 | 37 | 58.8 | 76 | 39 | 52.3 | 64 | -11.1 | -15.8 | **0.0067** |
| Heat | 3 | D42 | 4.3 | ♂ | 2 | 42 | 55.5 | 76 | 32 | 52.0 | 76 | 40 | 50.1 | 68 | -3.7 | -10.5 | **0.0266** |
| Heat | 3 | D42 | 4.3 | ♂ | 3 | 41 | 51.9 | 76 | 39 | 53.9 | 76 | 42 | 49.0 | 56 | -5.7 | -26.3 | **0.0082** |
| Heat | 3 | D42 | 4.3 | ♂ | 4 | 40 | 25.5 | 36 | 40 | 27.9 | 41 | 40 | 27.7 | 36 | 0.0 | 0.0 | |
| Heat | 3 | D42 | 9.1 | ♀ | 1 | 39 | 69.1 | 93 | 30 | 63.0 | 93 | 39 | 61.3 | 76 | -2.7 | -18.3 | 0.4739 |
| Heat | 3 | D42 | 9.1 | ♀ | 2 | 41 | 61.6 | 76 | 36 | 54.3 | 76 | 46 | 59.4 | 84 | 0.0 | 10.5 | |
| Heat | 3 | D42 | 9.1 | ♀ | 3 | 43 | 60.7 | 76 | 40 | 62.4 | 93 | 39 | 59.9 | 76 | -1.4 | 0.0 | 0.1615 |
| Heat | 3 | D42 | 9.1 | ♂ | 1 | 39 | 68.7 | 84 | 37 | 58.8 | 76 | 41 | 53.4 | 76 | -9.3 | 0.0 | **0.0103** |
| Heat | 3 | D42 | 9.1 | ♂ | 2 | 40 | 57.7 | 76 | 32 | 52.0 | 76 | 42 | 45.5 | 56 | -12.5 | -26.3 | **0.0004** |
| Heat | 3 | D42 | 9.1 | ♂ | 3 | 40 | 55.4 | 76 | 39 | 53.9 | 76 | 42 | 50.3 | 76 | -6.8 | 0.0 | **0.0081** |
| Heat | 3 | D42 | 3.2 | ♀ | 1 | 40 | 60.1 | 76 | 30 | 63.0 | 93 | 40 | 65.0 | 93 | 3.0 | 0.0 | **0.0197** |
| Heat | 3 | D42 | 3.2 | ♀ | 2 | 39 | 52.8 | 68 | 36 | 54.3 | 76 | 37 | 55.9 | 68 | 2.9 | 0.0 | **0.0180** |
| Heat | 3 | D42 | 3.2 | ♀ | 3 | 40 | 50.2 | 56 | 40 | 62.4 | 93 | 39 | 55.0 | 76 | 0.0 | 0.0 | |
| Heat | 3 | D42 | 3.2 | ♀ | 4 | 39 | 31.2 | 42 | 39 | 22.6 | 30 | 40 | 34.5 | 42 | 10.6 | 0.0 | **0.0208** |
| Heat | 3 | D42 | 3.2 | ♂ | 1 | 40 | 60.1 | 93 | 37 | 58.8 | 76 | 39 | 56.5 | 76 | -3.9 | 0.0 | 0.1721 |
| Heat | 3 | D42 | 3.2 | ♂ | 2 | 45 | 46.5 | 64 | 32 | 52.0 | 76 | 47 | 46.8 | 64 | 0.0 | 0.0 | |
| Heat | 3 | D42 | 3.2 | ♂ | 3 | 40 | 47.4 | 60 | 39 | 53.9 | 76 | 40 | 48.0 | 68 | 0.0 | 0.0 | |
| Heat | 3 | D42 | 3.2 | ♂ | 4 | 36 | 21.3 | 30 | 39 | 21.3 | 38 | 40 | 25.7 | 34 | 20.4 | 0.0 | **0.0989** |
| Heat | 3 | D42 | 10.1 | ♀ | 1 | 40 | 61.5 | 84 | 30 | 63.0 | 93 | 41 | 52.0 | 60 | -15.4 | -28.6 | **<0.0001** |
| Heat | 3 | D42 | 10.1 | ♀ | 2 | 40 | 50.7 | 56 | 36 | 54.3 | 76 | 41 | 53.3 | 56 | 0.0 | 0.0 | |
| Heat | 3 | D42 | 10.1 | ♀ | 3 | 40 | 50.4 | 56 | 40 | 62.4 | 93 | 40 | 52.7 | 60 | 0.0 | 0.0 | |
| Heat | 3 | D42 | 10.1 | ♀ | 4 | 39 | 32.5 | 41 | 39 | 25.0 | 36 | 40 | 31.9 | 36 | 0.0 | 0.0 | |
| Heat | 3 | D42 | 10.1 | ♂ | 1 | 29 | 61.8 | 84 | 37 | 58.8 | 76 | 39 | 49.6 | 64 | -15.6 | -15.8 | **<0.0001** |
| Heat | 3 | D42 | 10.1 | ♂ | 2 | 40 | 46.4 | 52 | 32 | 52.0 | 76 | 40 | 49.7 | 56 | 0.0 | 0.0 | |
| Heat | 3 | D42 | 10.1 | ♂ | 3 | 39 | 48.3 | 60 | 39 | 53.9 | 76 | 39 | 45.0 | 52 | -6.8 | -13.3 | **0.0127** |
| Heat | 3 | D42 | 10.1 | ♂ | 4 | 40 | 26.2 | 36 | 40 | 21.7 | 26 | 40 | 25.8 | 31 | 0.0 | 0.0 | |
| Heat | 3 | D42 | 3.1 | ♀ | 1 | 40 | 59.0 | 76 | 30 | 63.0 | 93 | 52 | 57.9 | 76 | -1.8 | 0.0 | **0.0290** |

### Table S2



| Stress | Age | Gal4 | UAS | Sex | Rep[1] | UAS | | | Gal4 | | | Gal4 + UAS | | | % Extension | | *P* |
|---|---|---|---|---|---|---|---|---|---|---|---|---|---|---|---|---|---|
| | | | | | | n | Mean | Max[2] | n | Mean | Max[2] | n | Mean | Max[2] | Mean | Max[2] | |
| Heat | 3 | D42 | 3.1 | ♀ | 2 | 41 | 50.4 | 60 | 36 | 54.3 | 76 | 42 | 58.7 | 93 | 8.1 | 22.4 | 0.1511 |
| Heat | 3 | D42 | 3.1 | ♀ | 3 | 38 | 46.2 | 56 | 40 | 62.4 | 93 | 41 | 52.7 | 76 | 0.0 | 0.0 | |
| Heat | 3 | D42 | 3.1 | ♀ | 4 | 38 | 22.6 | 34 | 39 | 22.6 | 30 | 40 | 23.7 | 34 | 5.0 | 0.0 | 0.1976 |
| Heat | 3 | D42 | 3.1 | ♂ | 1 | 39 | 61.9 | 76 | 37 | 58.8 | 76 | 40 | 54.7 | 76 | -7.0 | 0.0 | **0.0006** |
| Heat | 3 | D42 | 3.1 | ♂ | 2 | 40 | 42.6 | 56 | 32 | 52.0 | 76 | 40 | 52.9 | 76 | 1.7 | 0.0 | 0.7850 |
| Heat | 3 | D42 | 3.1 | ♂ | 3 | 18 | 52.2 | 76 | 39 | 53.9 | 76 | 42 | 49.8 | 68 | -4.6 | -10.5 | **0.0430** |
| Heat | 3 | D42 | 3.1 | ♂ | 4 | 40 | 16.5 | 26 | 39 | 21.3 | 38 | 40 | 17.5 | 22 | 0.0 | -15.4 | 0.0041 |
| Heat | 3 | D42 | 4.4 | ♀ | 1 | 41 | 67.1 | 84 | 30 | 63.0 | 93 | 38 | 65.1 | 93 | 0.0 | 0.0 | |
| Heat | 3 | D42 | 4.4 | ♀ | 2 | 33 | 60.8 | 76 | 36 | 54.3 | 76 | 47 | 60.6 | 93 | 0.0 | 22.4 | |
| Heat | 3 | D42 | 4.4 | ♀ | 3 | 40 | 59.3 | 76 | 40 | 62.4 | 93 | 46 | 55.4 | 76 | -6.6 | 0.0 | **0.0004** |
| Heat | 3 | D42 | 4.4 | ♀ | 4 | 40 | 32.2 | 41 | 40 | 27.5 | 36 | 40 | 36.5 | 46 | 13.4 | 12.2 | **0.0011** |
| Heat | 3 | D42 | 4.4 | ♂ | 1 | 43 | 69.9 | 84 | 37 | 58.8 | 76 | 41 | 57.5 | 76 | -2.3 | 0.0 | 0.6225 |
| Heat | 3 | D42 | 4.4 | ♂ | 2 | 40 | 56.2 | 76 | 32 | 52.0 | 76 | 45 | 58.9 | 84 | 4.9 | 10.5 | **0.0091** |
| Heat | 3 | D42 | 4.4 | ♂ | 3 | 38 | 55.2 | 76 | 39 | 53.9 | 76 | 40 | 53.6 | 76 | -0.6 | 0.0 | 0.5432 |
| Heat | 3 | D42 | 4.4 | ♂ | 4 | 40 | 31.3 | 36 | 41 | 28.0 | 41 | 40 | 35.8 | 46 | 14.4 | 12.2 | **<0.0001** |
| Heat | 10 | daGal4 | 2.1 | ♀ | 1 | 40 | 20.8 | 30 | 38 | 18.5 | 30 | 40 | 21.1 | 30 | 1.4 | 0.0 | **0.0164** |
| Heat | 10 | daGal4 | 2.1 | ♀ | 2 | 39 | 31.5 | 36 | 41 | 30.0 | 33 | 39 | 30.4 | 33 | 0.0 | 0.0 | |
| Heat | 10 | daGal4 | 2.1 | ♀ | 3 | 39 | 33.5 | 40 | 40 | 30.7 | 36 | 40 | 32.1 | 40 | 0.0 | 0.0 | |
| Heat | 10 | daGal4 | 2.1 | ♀ | 4 | 36 | 30.1 | 33 | 37 | 28.5 | 33 | 40 | 29.0 | 33 | 0.0 | 0.0 | |
| Heat | 10 | daGal4 | 2.1 | ♂ | 1 | 39 | 16.4 | 21 | 37 | 15.9 | 21 | 40 | 17.9 | 21 | 9.1 | 0.0 | **0.0027** |
| Heat | 10 | daGal4 | 2.1 | ♂ | 2 | 39 | 27.0 | 28 | 38 | 25.3 | 28 | 39 | 27.7 | 28 | 2.7 | 0.0 | **0.0346** |
| Heat | 10 | daGal4 | 2.1 | ♂ | 3 | 38 | 27.4 | 33 | 40 | 25.7 | 28 | 41 | 27.3 | 33 | 0.0 | 0.0 | |
| Heat | 10 | daGal4 | 2.1 | ♂ | 4 | 40 | 24.7 | 28 | 40 | 24.9 | 28 | 39 | 25.0 | 28 | 0.5 | 0.0 | 0.2057 |
| Heat | 10 | daGal4 | 4.3 | ♀ | 1 | 26 | 17.5 | 21 | 39 | 17.8 | 21 | 40 | 18.7 | 21 | 4.9 | 0.0 | 0.0697 |
| Heat | 10 | daGal4 | 4.3 | ♀ | 2 | 39 | 31.2 | 36 | 41 | 30.0 | 33 | 40 | 33.4 | 40 | 7.0 | 11.1 | **0.0005** |
| Heat | 10 | daGal4 | 4.3 | ♀ | 3 | 41 | 32.2 | 36 | 40 | 30.7 | 36 | 42 | 33.0 | 36 | 2.4 | 0.0 | **0.0008** |
| Heat | 10 | daGal4 | 4.3 | ♀ | 4 | 38 | 29.1 | 33 | 37 | 28.5 | 33 | 40 | 31.7 | 33 | 9.1 | 0.0 | **0.0243** |
| Heat | 10 | daGal4 | 4.3 | ♂ | 1 | 27 | 14.6 | 17 | 39 | 16.9 | 21 | 40 | 15.1 | 17 | 0.0 | 0.0 | |
| Heat | 10 | daGal4 | 4.3 | ♂ | 2 | 31 | 25.7 | 28 | 38 | 25.3 | 28 | 39 | 28.4 | 33 | 10.7 | 17.9 | **0.0013** |
| Heat | 10 | daGal4 | 4.3 | ♂ | 3 | 41 | 26.2 | 28 | 40 | 25.7 | 28 | 39 | 27.8 | 33 | 6.4 | 17.9 | **0.0004** |
| Heat | 10 | daGal4 | 4.3 | ♂ | 4 | 41 | 24.2 | 28 | 40 | 24.9 | 28 | 35 | 27.1 | 28 | 8.9 | 0.0 | **<0.0001** |
| Heat | 10 | daGal4 | 9.1 | ♀ | 1 | 39 | 31.8 | 36 | 41 | 30.0 | 33 | 41 | 32.0 | 40 | 0.6 | 11.1 | **0.0027** |
| Heat | 10 | daGal4 | 9.1 | ♀ | 2 | 40 | 32.8 | 40 | 40 | 30.7 | 36 | 41 | 35.0 | 40 | 6.9 | 0.0 | **0.0051** |
| Heat | 10 | daGal4 | 9.1 | ♀ | 3 | 39 | 29.3 | 33 | 37 | 28.5 | 33 | 39 | 29.5 | 36 | 0.7 | 9.1 | 0.0638 |
| Heat | 10 | daGal4 | 9.1 | ♂ | 1 | 40 | 25.8 | 28 | 38 | 25.3 | 28 | 39 | 29.0 | 33 | 12.5 | 17.9 | **<0.0001** |
| Heat | 10 | daGal4 | 9.1 | ♂ | 2 | 41 | 27.6 | 33 | 40 | 25.7 | 28 | 38 | 30.5 | 33 | 10.6 | 0.0 | **<0.0001** |
| Heat | 10 | daGal4 | 9.1 | ♂ | 3 | 43 | 25.1 | 28 | 40 | 24.9 | 28 | 41 | 27.5 | 28 | 9.6 | 0.0 | **<0.0001** |
| Heat | 10 | daGal4 | 3.2 | ♀ | 1 | 32 | 19.0 | 27 | 40 | 18.0 | 24 | 40 | 30.6 | 40 | 61.2 | 48.1 | **<0.0001** |
| Heat | 10 | daGal4 | 3.2 | ♀ | 2 | 39 | 35.3 | 40 | 41 | 30.0 | 33 | 40 | 32.0 | 36 | 0.0 | 0.0 | |
| Heat | 10 | daGal4 | 3.2 | ♀ | 3 | 40 | 36.6 | 40 | 40 | 30.7 | 36 | 38 | 32.3 | 36 | 0.0 | 0.0 | |
| Heat | 10 | daGal4 | 3.2 | ♀ | 4 | 37 | 32.0 | 40 | 37 | 28.5 | 33 | 40 | 29.4 | 33 | 0.0 | 0.0 | |
| Heat | 10 | daGal4 | 3.2 | ♂ | 1 | 36 | 16.1 | 24 | 40 | 17.1 | 21 | 40 | 17.6 | 21 | 2.6 | 0.0 | **0.0426** |
| Heat | 10 | daGal4 | 3.2 | ♂ | 2 | 40 | 33.5 | 40 | 38 | 25.3 | 28 | 42 | 29.4 | 33 | 0.0 | 0.0 | |
| Heat | 10 | daGal4 | 3.2 | ♂ | 3 | 41 | 32.4 | 40 | 40 | 25.7 | 28 | 42 | 31.8 | 40 | 0.0 | 0.0 | |
| Heat | 10 | daGal4 | 3.2 | ♂ | 4 | 43 | 30.3 | 36 | 40 | 24.9 | 28 | 46 | 28.1 | 33 | 0.0 | 0.0 | |
| Heat | 10 | daGal4 | 10.1 | ♀ | 1 | 36 | 19.3 | 21 | 39 | 17.8 | 21 | 40 | 21.4 | 30 | 10.4 | 42.9 | **0.0019** |
| Heat | 10 | daGal4 | 10.1 | ♀ | 2 | 41 | 34.1 | 40 | 41 | 30.0 | 33 | 40 | 30.4 | 33 | 0.0 | 0.0 | |
| Heat | 10 | daGal4 | 10.1 | ♀ | 3 | 41 | 36.1 | 45 | 40 | 30.7 | 36 | 40 | 31.7 | 36 | 0.0 | 0.0 | |
| Heat | 10 | daGal4 | 10.1 | ♀ | 4 | 39 | 33.4 | 40 | 37 | 28.5 | 33 | 38 | 28.5 | 33 | 0.0 | 0.0 | |

### Table S2



| Stress | Age | Gal4 | UAS | Sex | Rep[1] | UAS | | | Gal4 | | | Gal4 + UAS | | | % Extension | | *P* |
|---|---|---|---|---|---|---|---|---|---|---|---|---|---|---|---|---|---|
| | | | | | | n | Mean | Max[2] | n | Mean | Max[2] | n | Mean | Max[2] | Mean | Max[2] | |
| Heat | 10 | daGal4 | 10.1 | ♂ | 1 | 40 | 16.6 | 21 | 39 | 16.9 | 21 | 38 | 17.9 | 21 | 6.2 | 0.0 | **0.0296** |
| Heat | 10 | daGal4 | 10.1 | ♂ | 2 | 40 | 31.9 | 40 | 38 | 25.3 | 28 | 39 | 30.4 | 33 | 0.0 | 0.0 | |
| Heat | 10 | daGal4 | 10.1 | ♂ | 3 | 42 | 32.5 | 40 | 40 | 25.7 | 28 | 39 | 31.9 | 36 | 0.0 | 0.0 | |
| Heat | 10 | daGal4 | 10.1 | ♂ | 4 | 45 | 29.1 | 36 | 40 | 24.9 | 28 | 40 | 28.5 | 33 | 0.0 | 0.0 | |
| Heat | 10 | daGal4 | 3.1 | ♀ | 1 | 40 | 15.5 | 21 | 40 | 18.0 | 24 | 40 | 21.6 | 32 | 19.7 | 33.3 | **0.0006** |
| Heat | 10 | daGal4 | 3.1 | ♀ | 2 | 41 | 35.6 | 45 | 41 | 30.0 | 33 | 40 | 32.1 | 33 | 0.0 | 0.0 | |
| Heat | 10 | daGal4 | 3.1 | ♀ | 3 | 35 | 36.0 | 40 | 40 | 30.7 | 36 | 41 | 31.0 | 36 | 0.0 | 0.0 | |
| Heat | 10 | daGal4 | 3.1 | ♀ | 4 | 40 | 32.8 | 40 | 37 | 28.5 | 33 | 42 | 30.7 | 36 | 0.0 | 0.0 | |
| Heat | 10 | daGal4 | 3.1 | ♂ | 1 | 39 | 13.3 | 21 | 40 | 17.1 | 21 | 40 | 18.8 | 24 | 9.6 | 14.3 | **0.0068** |
| Heat | 10 | daGal4 | 3.1 | ♂ | 2 | 40 | 30.8 | 40 | 38 | 25.3 | 28 | 39 | 30.4 | 33 | 0.0 | 0.0 | |
| Heat | 10 | daGal4 | 3.1 | ♂ | 3 | 40 | 32.6 | 40 | 40 | 25.7 | 28 | 40 | 29.7 | 33 | 0.0 | 0.0 | |
| Heat | 10 | daGal4 | 3.1 | ♂ | 4 | 39 | 29.1 | 36 | 40 | 24.9 | 28 | 41 | 27.9 | 33 | 0.0 | 0.0 | |
| Heat | 10 | daGal4 | 4.4 | ♀ | 1 | 39 | 17.6 | 21 | 39 | 17.8 | 21 | 39 | 18.6 | 21 | 4.6 | 0.0 | **0.0338** |
| Heat | 10 | daGal4 | 4.4 | ♀ | 2 | 40 | 32.1 | 36 | 41 | 30.0 | 33 | 41 | 31.0 | 33 | 0.0 | 0.0 | |
| Heat | 10 | daGal4 | 4.4 | ♀ | 3 | 40 | 34.5 | 40 | 40 | 30.7 | 36 | 42 | 29.6 | 36 | -3.4 | 0.0 | 0.1136 |
| Heat | 10 | daGal4 | 4.4 | ♀ | 4 | 38 | 29.8 | 36 | 37 | 28.5 | 33 | 41 | 28.1 | 33 | -1.4 | 0.0 | **0.0027** |
| Heat | 10 | daGal4 | 4.4 | ♂ | 1 | 39 | 13.9 | 17 | 39 | 16.9 | 21 | 40 | 15.8 | 17 | 0.0 | 0.0 | |
| Heat | 10 | daGal4 | 4.4 | ♂ | 2 | 39 | 29.7 | 33 | 38 | 25.3 | 28 | 40 | 27.9 | 28 | 0.0 | 0.0 | |
| Heat | 10 | daGal4 | 4.4 | ♂ | 3 | 39 | 29.9 | 33 | 40 | 25.7 | 28 | 41 | 29.0 | 33 | 0.0 | 0.0 | |
| Heat | 10 | daGal4 | 4.4 | ♂ | 4 | 41 | 27.2 | 28 | 40 | 24.9 | 28 | 38 | 25.6 | 28 | 0.0 | 0.0 | |
| Heat | 10 | DJ694 | 2.1 | ♀ | 1 | 40 | 20.8 | 30 | 38 | 20.4 | 30 | 40 | 20.1 | 30 | -1.8 | 0.0 | 0.1930 |
| Heat | 10 | DJ694 | 2.1 | ♀ | 2 | 39 | 31.5 | 36 | 40 | 33.3 | 36 | 39 | 30.6 | 36 | -2.7 | 0.0 | **0.0002** |
| Heat | 10 | DJ694 | 2.1 | ♀ | 3 | 39 | 33.5 | 40 | 41 | 35.3 | 40 | 40 | 32.6 | 40 | -2.8 | 0.0 | **0.0037** |
| Heat | 10 | DJ694 | 2.1 | ♀ | 4 | 36 | 30.1 | 33 | 39 | 31.3 | 33 | 39 | 31.1 | 33 | 0.0 | 0.0 | |
| Heat | 10 | DJ694 | 2.1 | ♂ | 1 | 39 | 16.4 | 21 | 37 | 14.9 | 21 | 40 | 16.1 | 21 | 0.0 | 0.0 | |
| Heat | 10 | DJ694 | 2.1 | ♂ | 2 | 39 | 27.0 | 28 | 40 | 27.9 | 28 | 41 | 26.6 | 28 | -1.3 | 0.0 | **0.0003** |
| Heat | 10 | DJ694 | 2.1 | ♂ | 3 | 38 | 27.4 | 33 | 40 | 26.3 | 28 | 40 | 24.6 | 28 | -6.5 | 0.0 | **<0.0001** |
| Heat | 10 | DJ694 | 2.1 | ♂ | 4 | 40 | 24.7 | 28 | 38 | 27.4 | 33 | 39 | 24.9 | 28 | 0.0 | 0.0 | |
| Heat | 10 | DJ694 | 4.3 | ♀ | 1 | 26 | 17.5 | 21 | 40 | 18.9 | 21 | 37 | 17.3 | 21 | -0.8 | 0.0 | **0.0176** |
| Heat | 10 | DJ694 | 4.3 | ♀ | 2 | 39 | 31.2 | 36 | 40 | 33.3 | 36 | 38 | 31.3 | 36 | 0.0 | 0.0 | |
| Heat | 10 | DJ694 | 4.3 | ♀ | 3 | 41 | 32.2 | 36 | 41 | 35.3 | 40 | 40 | 34.3 | 40 | 0.0 | 0.0 | |
| Heat | 10 | DJ694 | 4.3 | ♀ | 4 | 38 | 29.1 | 33 | 39 | 31.3 | 33 | 41 | 30.7 | 33 | 0.0 | 0.0 | |
| Heat | 10 | DJ694 | 4.3 | ♂ | 1 | 27 | 14.6 | 17 | 40 | 16.0 | 17 | 38 | 14.9 | 17 | 0.0 | 0.0 | |
| Heat | 10 | DJ694 | 4.3 | ♂ | 2 | 31 | 25.7 | 28 | 40 | 27.9 | 28 | 42 | 27.3 | 28 | 0.0 | 0.0 | |
| Heat | 10 | DJ694 | 4.3 | ♂ | 3 | 41 | 26.2 | 28 | 40 | 26.3 | 28 | 38 | 25.9 | 28 | -1.1 | 0.0 | 0.2365 |
| Heat | 10 | DJ694 | 4.3 | ♂ | 4 | 41 | 24.2 | 28 | 38 | 27.4 | 33 | 40 | 26.1 | 28 | 0.0 | 0.0 | |
| Heat | 10 | DJ694 | 9.1 | ♀ | 1 | 39 | 31.8 | 36 | 40 | 33.3 | 36 | 39 | 33.6 | 36 | 1.0 | 0.0 | **0.0298** |
| Heat | 10 | DJ694 | 9.1 | ♀ | 2 | 40 | 32.8 | 40 | 41 | 35.3 | 40 | 41 | 37.4 | 40 | 5.9 | 0.0 | **0.0035** |
| Heat | 10 | DJ694 | 9.1 | ♀ | 3 | 39 | 29.3 | 33 | 39 | 31.3 | 33 | 40 | 31.8 | 33 | 1.6 | 0.0 | 0.9057 |
| Heat | 10 | DJ694 | 9.1 | ♂ | 1 | 40 | 25.8 | 28 | 40 | 27.9 | 28 | 41 | 27.6 | 28 | 0.0 | 0.0 | |
| Heat | 10 | DJ694 | 9.1 | ♂ | 2 | 41 | 27.6 | 33 | 40 | 26.3 | 28 | 40 | 27.4 | 28 | 0.0 | 0.0 | |
| Heat | 10 | DJ694 | 9.1 | ♂ | 3 | 43 | 25.1 | 28 | 38 | 27.4 | 33 | 43 | 26.0 | 28 | 0.0 | 0.0 | |
| Heat | 10 | DJ694 | 3.2 | ♀ | 1 | 32 | 19.0 | 27 | 40 | 14.9 | 21 | 40 | 26.9 | 36 | 41.6 | 33.3 | **<0.0001** |
| Heat | 10 | DJ694 | 3.2 | ♀ | 2 | 39 | 35.3 | 40 | 40 | 33.3 | 36 | 40 | 35.2 | 40 | 0.0 | 0.0 | |
| Heat | 10 | DJ694 | 3.2 | ♀ | 3 | 40 | 36.6 | 40 | 41 | 35.3 | 40 | 40 | 41.1 | 45 | 12.4 | 12.5 | **<0.0001** |
| Heat | 10 | DJ694 | 3.2 | ♀ | 4 | 37 | 32.0 | 40 | 39 | 31.3 | 33 | 40 | 36.7 | 45 | 14.6 | 12.5 | **<0.0001** |
| Heat | 10 | DJ694 | 3.2 | ♂ | 1 | 36 | 16.1 | 24 | 40 | 15.7 | 21 | 40 | 16.2 | 18 | 0.7 | -14.3 | 0.5306 |
| Heat | 10 | DJ694 | 3.2 | ♂ | 2 | 40 | 33.5 | 40 | 40 | 27.9 | 28 | 40 | 28.5 | 33 | 0.0 | 0.0 | |
| Heat | 10 | DJ694 | 3.2 | ♂ | 3 | 41 | 32.4 | 40 | 40 | 26.3 | 28 | 40 | 29.9 | 33 | 0.0 | 0.0 | |

### Table S2



| Stress | Age | Gal4 | UAS | Sex | Rep[1] | UAS | | | Gal4 | | | Gal4 + UAS | | | % Extension | | *P* |
|---|---|---|---|---|---|---|---|---|---|---|---|---|---|---|---|---|---|
| | | | | | | n | Mean | Max[2] | n | Mean | Max[2] | n | Mean | Max[2] | Mean | Max[2] | |
| Heat | 10 | DJ694 | 3.2 | ♂ | 4 | 43 | 30.3 | 36 | 38 | 27.4 | 33 | 39 | 27.8 | 33 | 0.0 | 0.0 | |
| Heat | 10 | DJ694 | 10.1 | ♀ | 1 | 36 | 19.3 | 21 | 40 | 18.9 | 21 | 42 | 18.9 | 24 | 0.0 | 14.3 | |
| Heat | 10 | DJ694 | 10.1 | ♀ | 2 | 41 | 34.1 | 40 | 40 | 33.3 | 36 | 40 | 33.8 | 40 | 0.0 | 0.0 | |
| Heat | 10 | DJ694 | 10.1 | ♀ | 3 | 41 | 36.1 | 45 | 41 | 35.3 | 40 | 40 | 36.6 | 45 | 1.3 | 0.0 | 0.1489 |
| Heat | 10 | DJ694 | 10.1 | ♀ | 4 | 39 | 33.4 | 40 | 39 | 31.3 | 33 | 40 | 32.7 | 36 | 0.0 | 0.0 | |
| Heat | 10 | DJ694 | 10.1 | ♂ | 1 | 40 | 16.6 | 21 | 40 | 16.0 | 17 | 25 | 15.2 | 17 | -4.8 | 0.0 | 0.0319 |
| Heat | 10 | DJ694 | 10.1 | ♂ | 2 | 40 | 31.9 | 40 | 40 | 27.9 | 28 | 39 | 31.6 | 33 | 0.0 | 0.0 | |
| Heat | 10 | DJ694 | 10.1 | ♂ | 3 | 42 | 32.5 | 40 | 40 | 26.3 | 28 | 41 | 30.1 | 36 | 0.0 | 0.0 | |
| Heat | 10 | DJ694 | 10.1 | ♂ | 4 | 45 | 29.1 | 36 | 38 | 27.4 | 0.37 | 43 | 28.8 | 33 | 0.0 | 0.0 | |
| Heat | 10 | DJ694 | 3.1 | ♀ | 1 | 40 | 15.5 | 21 | 40 | 14.9 | 21 | 40 | 19.1 | 24 | 22.7 | 14.3 | **<0.0001** |
| Heat | 10 | DJ694 | 3.1 | ♀ | 2 | 41 | 35.6 | 45 | 40 | 33.3 | 36 | 40 | 32.0 | 33 | -3.9 | -8.3 | **0.0025** |
| Heat | 10 | DJ694 | 3.1 | ♀ | 3 | 35 | 36.0 | 40 | 41 | 35.3 | 40 | 41 | 37.2 | 40 | 3.4 | 0.0 | **0.0079** |
| Heat | 10 | DJ694 | 3.1 | ♀ | 4 | 40 | 32.8 | 40 | 39 | 31.3 | 33 | 38 | 31.9 | 36 | 0.0 | 0.0 | |
| Heat | 10 | DJ694 | 3.1 | ♂ | 1 | 39 | 13.3 | 21 | 40 | 15.7 | 21 | 40 | 15.7 | 18 | 0.0 | -14.3 | |
| Heat | 10 | DJ694 | 3.1 | ♂ | 2 | 40 | 30.8 | 40 | 40 | 27.9 | 28 | 39 | 27.6 | 28 | -1.1 | 0.0 | 0.1595 |
| Heat | 10 | DJ694 | 3.1 | ♂ | 3 | 40 | 32.6 | 40 | 40 | 26.3 | 28 | 38 | 29.2 | 33 | 0.0 | 0.0 | |
| Heat | 10 | DJ694 | 3.1 | ♂ | 4 | 39 | 29.1 | 36 | 38 | 27.4 | 33 | 39 | 26.7 | 28 | -2.8 | -15.2 | 0.0795 |
| Heat | 10 | DJ694 | 4.4 | ♀ | 1 | 39 | 17.6 | 21 | 40 | 18.9 | 21 | 40 | 20.4 | 21 | 7.9 | 0.0 | **0.0016** |
| Heat | 10 | DJ694 | 4.4 | ♀ | 2 | 40 | 32.1 | 36 | 40 | 33.3 | 36 | 40 | 30.1 | 33 | -6.0 | -8.3 | **0.0009** |
| Heat | 10 | DJ694 | 4.4 | ♀ | 3 | 40 | 34.5 | 40 | 41 | 35.3 | 40 | 40 | 31.1 | 33 | -9.9 | -17.5 | **<0.0001** |
| Heat | 10 | DJ694 | 4.4 | ♀ | 4 | 38 | 29.8 | 36 | 39 | 31.3 | 33 | 41 | 27.3 | 33 | -8.3 | 0.0 | **0.0014** |
| Heat | 10 | DJ694 | 4.4 | ♂ | 1 | 39 | 13.9 | 17 | 40 | 16.0 | 17 | 38 | 13.8 | 17 | -0.6 | 0.0 | 0.9654 |
| Heat | 10 | DJ694 | 4.4 | ♂ | 2 | 39 | 29.7 | 33 | 40 | 27.9 | 28 | 40 | 27.4 | 28 | -1.8 | 0.0 | **0.0493** |
| Heat | 10 | DJ694 | 4.4 | ♂ | 3 | 39 | 29.9 | 33 | 40 | 26.3 | 28 | 40 | 27.0 | 28 | 0.0 | 0.0 | |
| Heat | 10 | DJ694 | 4.4 | ♂ | 4 | 41 | 27.2 | 28 | 38 | 27.4 | 33 | 41 | 23.8 | 24 | -12.6 | -14.3 | **<0.0001** |
| Heat | 10 | D42 | 2.1 | ♀ | 1 | 40 | 20.8 | 30 | 38 | 22.7 | 30 | 40 | 20.3 | 25 | -2.2 | -16.7 | **0.0065** |
| Heat | 10 | D42 | 2.1 | ♀ | 2 | 39 | 31.5 | 36 | 38 | 33.6 | 40 | 40 | 32.8 | 36 | 0.0 | 0.0 | |
| Heat | 10 | D42 | 2.1 | ♀ | 3 | 39 | 33.5 | 40 | 40 | 35.4 | 40 | 31 | 32.5 | 36 | -2.8 | -10.0 | **0.0001** |
| Heat | 10 | D42 | 2.1 | ♀ | 4 | 36 | 30.1 | 33 | 40 | 28.8 | 36 | 40 | 28.9 | 36 | 0.0 | 0.0 | |
| Heat | 10 | D42 | 2.1 | ♂ | 1 | 39 | 16.4 | 21 | 36 | 13.7 | 17 | 40 | 13.5 | 17 | -1.6 | 0.0 | 0.4803 |
| Heat | 10 | D42 | 2.1 | ♂ | 2 | 39 | 27.0 | 28 | 42 | 27.8 | 33 | 50 | 25.1 | 33 | -6.8 | 0.0 | **0.0003** |
| Heat | 10 | D42 | 2.1 | ♂ | 3 | 38 | 27.4 | 33 | 40 | 27.6 | 33 | 40 | 28.1 | 33 | 1.8 | 0.0 | 0.2207 |
| Heat | 10 | D42 | 2.1 | ♂ | 4 | 40 | 24.7 | 28 | 40 | 25.8 | 33 | 40 | 25.0 | 28 | 0.0 | 0.0 | |
| Heat | 10 | D42 | 4.3 | ♀ | 1 | 26 | 17.5 | 21 | 39 | 21.3 | 30 | 40 | 19.3 | 21 | 0.0 | 0.0 | |
| Heat | 10 | D42 | 4.3 | ♀ | 2 | 39 | 31.2 | 36 | 38 | 33.6 | 40 | 40 | 32.0 | 36 | 0.0 | 0.0 | |
| Heat | 10 | D42 | 4.3 | ♀ | 3 | 41 | 32.2 | 36 | 40 | 35.4 | 40 | 39 | 34.7 | 40 | 0.0 | 0.0 | |
| Heat | 10 | D42 | 4.3 | ♀ | 4 | 38 | 29.1 | 33 | 40 | 28.8 | 36 | 40 | 29.3 | 33 | 0.9 | 0.0 | 0.2837 |
| Heat | 10 | D42 | 4.3 | ♂ | 1 | 27 | 14.6 | 17 | 39 | 18.5 | 21 | 40 | 12.6 | 13 | -13.9 | -23.5 | **<0.0001** |
| Heat | 10 | D42 | 4.3 | ♂ | 2 | 31 | 25.7 | 28 | 42 | 27.8 | 33 | 40 | 30.2 | 33 | 8.6 | 0.0 | **0.0004** |
| Heat | 10 | D42 | 4.3 | ♂ | 3 | 41 | 26.2 | 28 | 40 | 27.6 | 33 | 42 | 28.7 | 33 | 4.2 | 0.0 | **0.0002** |
| Heat | 10 | D42 | 4.3 | ♂ | 4 | 41 | 24.2 | 28 | 40 | 25.8 | 33 | 40 | 25.1 | 28 | 0.0 | 0.0 | |
| Heat | 10 | D42 | 9.1 | ♀ | 1 | 39 | 31.8 | 36 | 38 | 33.6 | 40 | 40 | 34.1 | 40 | 1.5 | 0.0 | **0.0047** |
| Heat | 10 | D42 | 9.1 | ♀ | 2 | 40 | 32.8 | 40 | 40 | 35.4 | 40 | 40 | 34.7 | 40 | 0.0 | 0.0 | |
| Heat | 10 | D42 | 9.1 | ♀ | 3 | 39 | 29.3 | 33 | 40 | 28.8 | 36 | 39 | 31.5 | 33 | 7.3 | 0.0 | **0.0003** |
| Heat | 10 | D42 | 9.1 | ♂ | 1 | 40 | 25.8 | 28 | 42 | 27.8 | 33 | 39 | 27.9 | 33 | 0.5 | 0.0 | 0.6270 |
| Heat | 10 | D42 | 9.1 | ♂ | 2 | 41 | 27.6 | 33 | 40 | 27.6 | 33 | 41 | 28.5 | 33 | 3.4 | 0.0 | 0.0760 |
| Heat | 10 | D42 | 9.1 | ♂ | 3 | 43 | 25.1 | 28 | 40 | 25.8 | 33 | 38 | 23.8 | 28 | -5.3 | 0.0 | **0.0182** |
| Heat | 10 | D42 | 3.2 | ♀ | 1 | 32 | 19.0 | 27 | 37 | 18.6 | 27 | 40 | 30.8 | 36 | 62.1 | 33.3 | **<0.0001** |
| Heat | 10 | D42 | 3.2 | ♀ | 2 | 39 | 35.3 | 40 | 38 | 33.6 | 40 | 40 | 36.3 | 40 | 3.0 | 0.0 | **0.0011** |

### Table S2



| Stress | Age | Gal4 | UAS | Sex | Rep[1] | UAS | | | Gal4 | | | Gal4 + UAS | | | % Extension | | *P* |
|---|---|---|---|---|---|---|---|---|---|---|---|---|---|---|---|---|---|
| | | | | | | n | Mean | Max[2] | n | Mean | Max[2] | n | Mean | Max[2] | Mean | Max[2] | |
| Heat | 10 | D42 | 3.2 | ♀ | 3 | 40 | 36.6 | 40 | 40 | 35.4 | 40 | 40 | 37.7 | 40 | 3.2 | 0.0 | **0.0006** |
| Heat | 10 | D42 | 3.2 | ♀ | 4 | 37 | 32.0 | 40 | 40 | 28.8 | 36 | 44 | 36.3 | 45 | 13.4 | 12.5 | **<0.0001** |
| Heat | 10 | D42 | 3.2 | ♂ | 1 | 36 | 16.1 | 24 | 38 | 15.9 | 21 | 40 | 16.3 | 24 | 1.2 | 0.0 | 0.4681 |
| Heat | 10 | D42 | 3.2 | ♂ | 2 | 40 | 33.5 | 40 | 42 | 27.8 | 33 | 41 | 31.9 | 36 | 0.0 | 0.0 | |
| Heat | 10 | D42 | 3.2 | ♂ | 3 | 41 | 32.4 | 40 | 40 | 27.6 | 33 | 43 | 33.4 | 40 | 3.0 | 0.0 | 0.1824 |
| Heat | 10 | D42 | 3.2 | ♂ | 4 | 43 | 30.3 | 36 | 40 | 25.8 | 33 | 37 | 31.1 | 36 | 2.5 | 0.0 | 0.4366 |
| Heat | 10 | D42 | 10.1 | ♀ | 1 | 36 | 19.3 | 21 | 39 | 21.3 | 30 | 31 | 26.0 | 0.79 | 22.2 | 0.0 | **<0.0001** |
| Heat | 10 | D42 | 10.1 | ♀ | 2 | 41 | 34.1 | 40 | 38 | 33.6 | 40 | 39 | 31.7 | 0.68 | -5.5 | 12.5 | **0.0149** |
| Heat | 10 | D42 | 10.1 | ♀ | 3 | 41 | 36.1 | 45 | 40 | 35.4 | 40 | 40 | 29.1 | 1.33 | -17.7 | 0.0 | **0.0022** |
| Heat | 10 | D42 | 10.1 | ♀ | 4 | 39 | 33.4 | 40 | 40 | 28.8 | 36 | 41 | 34.3 | 0.5 | 2.9 | 0.0 | 0.2401 |
| Heat | 10 | D42 | 10.1 | ♂ | 1 | 40 | 16.6 | 21 | 39 | 18.5 | 21 | 36 | 19.2 | 0.47 | 3.4 | 14.3 | **0.0002** |
| Heat | 10 | D42 | 10.1 | ♂ | 2 | 40 | 31.9 | 40 | 42 | 27.8 | 33 | 40 | 32.4 | 0.36 | 1.6 | 0.0 | 0.4031 |
| Heat | 10 | D42 | 10.1 | ♂ | 3 | 42 | 32.5 | 40 | 40 | 27.6 | 33 | 47 | 33.3 | 0.41 | 2.7 | 0.0 | 0.4520 |
| Heat | 10 | D42 | 10.1 | ♂ | 4 | 45 | 29.1 | 36 | 40 | 25.8 | 33 | 37 | 32.8 | 0.39 | 12.5 | 0.0 | **<0.0001** |
| Heat | 10 | D42 | 3.1 | ♀ | 1 | 40 | 15.5 | 21 | 37 | 18.6 | 27 | 40 | 18.4 | 24 | 0.0 | 0.0 | |
| Heat | 10 | D42 | 3.1 | ♀ | 2 | 41 | 35.6 | 45 | 38 | 33.6 | 40 | 30 | 32.0 | 33 | -4.8 | -17.5 | **0.0371** |
| Heat | 10 | D42 | 3.1 | ♀ | 3 | 35 | 36.0 | 40 | 40 | 35.4 | 40 | 40 | 28.3 | 36 | -19.9 | -10.0 | **<0.0001** |
| Heat | 10 | D42 | 3.1 | ♀ | 4 | 40 | 32.8 | 40 | 40 | 28.8 | 36 | 40 | 31.6 | 36 | 0.0 | 0.0 | |
| Heat | 10 | D42 | 3.1 | ♂ | 1 | 39 | 13.3 | 21 | 38 | 15.9 | 21 | 40 | 11.6 | 15 | -12.6 | -28.6 | **0.0115** |
| Heat | 10 | D42 | 3.1 | ♂ | 2 | 40 | 30.8 | 40 | 42 | 27.8 | 33 | 40 | 28.4 | 33 | 0.0 | 0.0 | |
| Heat | 10 | D42 | 3.1 | ♂ | 3 | 40 | 32.6 | 40 | 40 | 27.6 | 33 | 40 | 28.6 | 33 | 0.0 | 0.0 | |
| Heat | 10 | D42 | 3.1 | ♂ | 4 | 39 | 29.1 | 36 | 40 | 25.8 | 33 | 30 | 29.6 | 33 | 1.9 | 0.0 | 0.8362 |
| Heat | 10 | D42 | 4.4 | ♀ | 1 | 39 | 17.6 | 21 | 39 | 21.3 | 30 | 40 | 23.3 | 30 | 9.4 | 0.0 | **0.0109** |
| Heat | 10 | D42 | 4.4 | ♀ | 2 | 40 | 32.1 | 36 | 38 | 33.6 | 40 | 38 | 34.6 | 40 | 3.0 | 0.0 | 0.2075 |
| Heat | 10 | D42 | 4.4 | ♀ | 3 | 40 | 34.5 | 40 | 40 | 35.4 | 40 | 40 | 35.1 | 40 | 0.0 | 0.0 | 0.2551 |
| Heat | 10 | D42 | 4.4 | ♀ | 4 | 38 | 29.8 | 36 | 40 | 28.8 | 36 | 39 | 33.6 | 36 | 13.0 | 0.0 | **<0.0001** |
| Heat | 10 | D42 | 4.4 | ♂ | 1 | 39 | 13.9 | 17 | 39 | 18.5 | 21 | 39 | 14.4 | 17 | 0.0 | 0.0 | |
| Heat | 10 | D42 | 4.4 | ♂ | 2 | 39 | 29.7 | 33 | 42 | 27.8 | 33 | 40 | 29.4 | 33 | 0.0 | 0.0 | |
| Heat | 10 | D42 | 4.4 | ♂ | 3 | 39 | 29.9 | 33 | 40 | 27.6 | 33 | 41 | 29.8 | 33 | 0.0 | 0.0 | |
| Heat | 10 | D42 | 4.4 | ♂ | 4 | 41 | 27.2 | 28 | 40 | 25.8 | 33 | 39 | 31.0 | 33 | 13.7 | 0.0 | **<0.0001** |
| Paraquat | 3 | daGal4 | 2.1 | ♀ | 1 | 40 | 29.4 | 47 | 40 | 25.0 | 42 | 40 | 24.6 | 52 | -1.5 | 10.6 | 0.0802 |
| Paraquat | 3 | daGal4 | 2.1 | ♀ | 2 | 40 | 18.9 | 30 | 40 | 14.4 | 20 | 40 | 15.3 | 20 | 0.0 | 0.0 | |
| Paraquat | 3 | daGal4 | 2.1 | ♀ | 3 | 40 | 55.9 | 90 | 38 | 51.3 | 90 | 38 | 49.9 | 90 | -2.6 | 0.0 | 0.7017 |
| Paraquat | 3 | daGal4 | 2.1 | ♀ | 4 | 40 | 17.8 | 25 | 40 | 13.6 | 20 | 40 | 15.5 | 20 | 0.0 | 0.0 | |
| Paraquat | 3 | daGal4 | 2.1 | ♀ | 5 | 37 | 56.4 | 84 | 40 | 43.6 | 90 | 39 | 42.1 | 90 | -3.4 | 0.0 | **0.0348** |
| Paraquat | 3 | daGal4 | 2.1 | ♀ | 6 | 42 | 50.9 | 84 | 39 | 41.6 | 74 | 39 | 45.3 | 90 | 0.0 | 7.1 | |
| Paraquat | 3 | daGal4 | 2.1 | ♂ | 1 | 40 | 27.5 | 57 | 40 | 22.5 | 42 | 40 | 18.6 | 32 | -17.2 | -23.8 | **0.0199** |
| Paraquat | 3 | daGal4 | 2.1 | ♂ | 2 | 40 | 17.9 | 25 | 40 | 14.1 | 20 | 40 | 14.8 | 25 | 0.0 | 0.0 | |
| Paraquat | 3 | daGal4 | 2.1 | ♂ | 3 | 40 | 35.5 | 66 | 41 | 22.8 | 34 | 41 | 28.2 | 66 | 0.0 | 0.0 | |
| Paraquat | 3 | daGal4 | 2.1 | ♂ | 4 | 40 | 17.8 | 25 | 40 | 14.1 | 20 | 40 | 13.8 | 20 | -2.7 | 0.0 | **0.0001** |
| Paraquat | 3 | daGal4 | 2.1 | ♂ | 5 | 34 | 40.2 | 84 | 40 | 26.6 | 50 | 40 | 27.6 | 50 | 0.0 | 0.0 | |
| Paraquat | 3 | daGal4 | 2.1 | ♂ | 6 | 41 | 39.0 | 84 | 42 | 23.1 | 38 | 40 | 27.2 | 50 | 0.0 | 0.0 | |
| Paraquat | 3 | daGal4 | 4.3 | ♀ | 1 | 40 | 30.8 | 43 | 40 | 38.8 | 56 | 40 | 28.5 | 56 | -7.5 | 0.0 | **0.0012** |
| Paraquat | 3 | daGal4 | 4.3 | ♀ | 2 | 42 | 49.1 | 84 | 38 | 51.3 | 90 | 40 | 40.9 | 74 | -16.9 | -11.9 | **0.0135** |
| Paraquat | 3 | daGal4 | 4.3 | ♀ | 3 | 41 | 39.3 | 74 | 40 | 43.6 | 90 | 37 | 34.5 | 58 | -12.0 | -21.6 | **0.0263** |
| Paraquat | 3 | daGal4 | 4.3 | ♀ | 4 | 40 | 43.2 | 74 | 39 | 41.6 | 74 | 41 | 45.1 | 74 | 4.3 | 0.0 | 0.2569 |
| Paraquat | 3 | daGal4 | 4.3 | ♂ | 1 | 40 | 27.8 | 56 | 40 | 29.4 | 69 | 40 | 32.2 | 69 | 9.4 | 0.0 | 0.2330 |
| Paraquat | 3 | daGal4 | 4.3 | ♂ | 2 | 50 | 42.9 | 84 | 41 | 22.8 | 34 | 40 | 30.5 | 74 | 0.0 | 0.0 | |
| Paraquat | 3 | daGal4 | 4.3 | ♂ | 3 | 41 | 41.6 | 110 | 40 | 26.6 | 50 | 40 | 39.7 | 84 | 0.0 | 0.0 | |

### Table S2



| Stress | Age | Gal4 | UAS | Sex | Rep[1] | UAS n | UAS Mean | UAS Max[2] | Gal4 n | Gal4 Mean | Gal4 Max[2] | Gal4 + UAS n | Gal4 + UAS Mean | Gal4 + UAS Max[2] | % Extension Mean | % Extension Max[2] | *P* |
|---|---|---|---|---|---|---|---|---|---|---|---|---|---|---|---|---|---|
| Paraquat | 3 | daGal4 | 4.3 | ♂ | 4 | 41 | 46.7 | 84 | 42 | 23.1 | 38 | 36 | 48.2 | 90 | 3.2 | 7.1 | 0.8014 |
| Paraquat | 3 | daGal4 | 9.1 | ♀ | 1 | 39 | 21.2 | 32 | 36 | 16.9 | 32 | 39 | 23.0 | 41 | 8.6 | 28.1 | **0.0002** |
| Paraquat | 3 | daGal4 | 9.1 | ♀ | 2 | 39 | 54.4 | 90 | 38 | 51.3 | 90 | 40 | 40.7 | 84 | -20.7 | -6.7 | **0.0063** |
| Paraquat | 3 | daGal4 | 9.1 | ♀ | 3 | 40 | 21.5 | 32 | 37 | 18.5 | 32 | 40 | 17.8 | 23 | -3.6 | -28.1 | **0.0162** |
| Paraquat | 3 | daGal4 | 9.1 | ♀ | 4 | 39 | 55.1 | 84 | 40 | 43.6 | 90 | 42 | 34.2 | 58 | -21.5 | -31.0 | **0.0262** |
| Paraquat | 3 | daGal4 | 9.1 | ♀ | 5 | 46 | 49.4 | 90 | 39 | 41.6 | 74 | 40 | 41.2 | 84 | -0.9 | 0.0 | **0.0313** |
| Paraquat | 3 | daGal4 | 9.1 | ♂ | 1 | 40 | 14.7 | 23 | 39 | 17.6 | 32 | 40 | 14.3 | 23 | -3.1 | 0.0 | **0.0088** |
| Paraquat | 3 | daGal4 | 9.1 | ♂ | 2 | 42 | 31.0 | 66 | 41 | 22.8 | 34 | 40 | 31.1 | 58 | 0.5 | 0.0 | **0.0004** |
| Paraquat | 3 | daGal4 | 9.1 | ♂ | 3 | 40 | 15.7 | 23 | 40 | 15.1 | 23 | 38 | 13.6 | 23 | -9.7 | 0.0 | **0.0249** |
| Paraquat | 3 | daGal4 | 9.1 | ♂ | 4 | 34 | 38.2 | 74 | 40 | 26.6 | 50 | 43 | 26.8 | 50 | 0.0 | 0.0 | |
| Paraquat | 3 | daGal4 | 9.1 | ♂ | 5 | 39 | 31.2 | 66 | 42 | 23.1 | 38 | 39 | 25.7 | 46 | 0.0 | 0.0 | |
| Paraquat | 3 | daGal4 | 3.2 | ♀ | 1 | 40 | 30.7 | 80 | 40 | 27.8 | 65 | 40 | 31.1 | 75 | 1.3 | 0.0 | 0.2841 |
| Paraquat | 3 | daGal4 | 3.2 | ♀ | 2 | 36 | 69.0 | 118 | 38 | 51.3 | 90 | 40 | 50.8 | 110 | -1.0 | 0.0 | **0.0110** |
| Paraquat | 3 | daGal4 | 3.2 | ♀ | 3 | 38 | 67.4 | 110 | 40 | 43.6 | 90 | 42 | 54.8 | 90 | 0.0 | 0.0 | |
| Paraquat | 3 | daGal4 | 3.2 | ♀ | 4 | 40 | 60.2 | 110 | 39 | 41.6 | 74 | 40 | 41.9 | 110 | 0.0 | 0.0 | |
| Paraquat | 3 | daGal4 | 3.2 | ♂ | 1 | 40 | 33.9 | 80 | 40 | 25.1 | 51 | 40 | 42.1 | 94 | 24.0 | 17.5 | **0.0005** |
| Paraquat | 3 | daGal4 | 3.2 | ♂ | 2 | 39 | 54.8 | 90 | 41 | 22.8 | 34 | 41 | 51.2 | 110 | 0.0 | 22.2 | |
| Paraquat | 3 | daGal4 | 3.2 | ♂ | 3 | 40 | 62.8 | 110 | 40 | 26.6 | 50 | 41 | 41.6 | 110 | 0.0 | 0.0 | |
| Paraquat | 3 | daGal4 | 3.2 | ♂ | 4 | 37 | 64.3 | 110 | 42 | 23.1 | 38 | 40 | 52.2 | 118 | 0.0 | 7.3 | |
| Paraquat | 3 | daGal4 | 10.1 | ♀ | 1 | 40 | 41.6 | 117 | 40 | 42.5 | 81 | 39 | 54.3 | 105 | 27.6 | 0.0 | **0.0252** |
| Paraquat | 3 | daGal4 | 10.1 | ♀ | 2 | 40 | 23.7 | 41 | 36 | 16.9 | 32 | 40 | 22.3 | 41 | 0.0 | 0.0 | |
| Paraquat | 3 | daGal4 | 10.1 | ♀ | 3 | 28 | 66.1 | 90 | 38 | 51.3 | 90 | 42 | 67.0 | 110 | 1.5 | 22.2 | **0.0101** |
| Paraquat | 3 | daGal4 | 10.1 | ♀ | 4 | 40 | 24.3 | 59 | 37 | 18.5 | 32 | 40 | 23.1 | 41 | 0.0 | 0.0 | |
| Paraquat | 3 | daGal4 | 10.1 | ♀ | 5 | 39 | 88.3 | 118 | 40 | 43.6 | 90 | 41 | 60.0 | 110 | 0.0 | 0.0 | |
| Paraquat | 3 | daGal4 | 10.1 | ♀ | 6 | 37 | 78.3 | 110 | 39 | 41.6 | 74 | 40 | 59.6 | 110 | 0.0 | 0.0 | |
| Paraquat | 3 | daGal4 | 10.1 | ♂ | 1 | 40 | 63.7 | 117 | 40 | 34.0 | 69 | 40 | 69.0 | 129 | 8.4 | 10.3 | 0.2277 |
| Paraquat | 3 | daGal4 | 10.1 | ♂ | 2 | 40 | 23.3 | 41 | 39 | 17.6 | 32 | 39 | 21.6 | 41 | 0.0 | 0.0 | |
| Paraquat | 3 | daGal4 | 10.1 | ♂ | 3 | 39 | 47.5 | 90 | 41 | 22.8 | 34 | 43 | 42.5 | 110 | 0.0 | 22.2 | |
| Paraquat | 3 | daGal4 | 10.1 | ♂ | 4 | 40 | 28.3 | 41 | 40 | 15.1 | 23 | 38 | 18.7 | 32 | 0.0 | 0.0 | |
| Paraquat | 3 | daGal4 | 10.1 | ♂ | 5 | 39 | 73.1 | 130 | 40 | 26.6 | 50 | 41 | 42.6 | 110 | 0.0 | 0.0 | |
| Paraquat | 3 | daGal4 | 10.1 | ♂ | 6 | 42 | 59.0 | 110 | 42 | 23.1 | 38 | 40 | 38.9 | 90 | 0.0 | 0.0 | |
| Paraquat | 3 | daGal4 | 3.1 | ♀ | 1 | 40 | 33.4 | 64 | 40 | 36.4 | 64 | 40 | 35.0 | 64 | 0.0 | 0.0 | |
| Paraquat | 3 | daGal4 | 3.1 | ♀ | 2 | 40 | 16.0 | 25 | 40 | 14.4 | 20 | 40 | 17.4 | 25 | 8.6 | 0.0 | **0.0011** |
| Paraquat | 3 | daGal4 | 3.1 | ♀ | 3 | 40 | 46.1 | 74 | 38 | 51.3 | 90 | 40 | 55.4 | 90 | 8.1 | 0.0 | **0.0016** |
| Paraquat | 3 | daGal4 | 3.1 | ♀ | 4 | 40 | 18.0 | 30 | 40 | 13.6 | 20 | 40 | 16.5 | 25 | 0.0 | 0.0 | |
| Paraquat | 3 | daGal4 | 3.1 | ♀ | 5 | 36 | 50.6 | 84 | 40 | 43.6 | 90 | 41 | 58.1 | 84 | 14.8 | 0.0 | **0.0242** |
| Paraquat | 3 | daGal4 | 3.1 | ♀ | 6 | 44 | 49.4 | 74 | 39 | 41.6 | 74 | 39 | 59.9 | 110 | 21.4 | 48.6 | **0.0048** |
| Paraquat | 3 | daGal4 | 3.1 | ♂ | 1 | 40 | 23.3 | 39 | 40 | 26.8 | 55 | 40 | 38.8 | 82 | 44.8 | 49.1 | **0.0002** |
| Paraquat | 3 | daGal4 | 3.1 | ♂ | 2 | 40 | 18.8 | 40 | 40 | 14.1 | 20 | 40 | 24.1 | 64 | 28.7 | 60.0 | **0.0373** |
| Paraquat | 3 | daGal4 | 3.1 | ♂ | 3 | 45 | 41.4 | 90 | 41 | 22.8 | 34 | 41 | 59.2 | 110 | 42.8 | 22.2 | **0.0009** |
| Paraquat | 3 | daGal4 | 3.1 | ♂ | 4 | 40 | 19.3 | 25 | 40 | 14.1 | 20 | 40 | 21.6 | 49 | 12.1 | 96.0 | 0.2077 |
| Paraquat | 3 | daGal4 | 3.1 | ♂ | 5 | 39 | 41.6 | 84 | 40 | 26.6 | 50 | 41 | 53.8 | 110 | 29.1 | 31.0 | **0.0115** |
| Paraquat | 3 | daGal4 | 3.1 | ♂ | 6 | 41 | 44.3 | 84 | 42 | 23.1 | 38 | 42 | 57.3 | 110 | 29.3 | 31.0 | **0.0259** |
| Paraquat | 3 | daGal4 | 4.4 | ♀ | 1 | 40 | 45.4 | 69 | 40 | 38.8 | 56 | 40 | 45.0 | 81 | 0.0 | 17.4 | |
| Paraquat | 3 | daGal4 | 4.4 | ♀ | 2 | 40 | 26.5 | 41 | 36 | 16.9 | 32 | 39 | 22.0 | 41 | 0.0 | 0.0 | |
| Paraquat | 3 | daGal4 | 4.4 | ♀ | 3 | 36 | 53.7 | 90 | 38 | 51.3 | 90 | 41 | 46.9 | 110 | -8.5 | 22.2 | 0.3531 |
| Paraquat | 3 | daGal4 | 4.4 | ♀ | 4 | 40 | 28.2 | 48 | 37 | 18.5 | 32 | 43 | 20.0 | 41 | 0.0 | 0.0 | |
| Paraquat | 3 | daGal4 | 4.4 | ♀ | 5 | 40 | 67.1 | 110 | 40 | 43.6 | 90 | 39 | 43.9 | 90 | 0.0 | 0.0 | |
| Paraquat | 3 | daGal4 | 4.4 | ♀ | 6 | 41 | 57.2 | 90 | 39 | 41.6 | 74 | 41 | 40.6 | 84 | -2.4 | 0.0 | **0.0023** |

### Table S2

| Stress | Age | Gal4 | UAS | Sex | Rep[1] | UAS | | | Gal4 | | | Gal4 + UAS | | | % Extension | | *P* |
|---|---|---|---|---|---|---|---|---|---|---|---|---|---|---|---|---|---|
| | | | | | | n | Mean | Max[2] | n | Mean | Max[2] | n | Mean | Max[2] | Mean | Max[2] | |
| Paraquat | 3 | daGal4 | 4.4 | ♂ | 1 | 40 | 39.2 | 81 | 40 | 29.4 | 69 | 40 | 46.8 | 105 | 19.3 | 29.6 | **0.0002** |
| Paraquat | 3 | daGal4 | 4.4 | ♂ | 2 | 39 | 19.1 | 32 | 39 | 17.6 | 32 | 39 | 17.7 | 59 | 0.0 | 84.4 | |
| Paraquat | 3 | daGal4 | 4.4 | ♂ | 3 | 41 | 37.7 | 84 | 41 | 22.8 | 34 | 41 | 49.6 | 110 | 31.6 | 31.0 | **0.0444** |
| Paraquat | 3 | daGal4 | 4.4 | ♂ | 4 | 40 | 19.3 | 32 | 40 | 15.1 | 23 | 41 | 16.7 | 23 | 0.0 | 0.0 | |
| Paraquat | 3 | daGal4 | 4.4 | ♂ | 5 | 40 | 57.9 | 110 | 40 | 26.6 | 50 | 42 | 40.4 | 84 | 0.0 | 0.0 | |
| Paraquat | 3 | daGal4 | 4.4 | ♂ | 6 | 40 | 43.0 | 110 | 42 | 23.1 | 38 | 40 | 32.1 | 84 | 0.0 | 0.0 | |
| Paraquat | 3 | DJ694 | 2.1 | ♀ | 1 | 40 | 29.5 | 51 | 40 | 20.7 | 35 | 40 | 20.2 | 40 | -2.5 | 0.0 | 0.7691 |
| Paraquat | 3 | DJ694 | 2.1 | ♀ | 2 | 40 | 29.4 | 47 | 40 | 18.6 | 27 | 40 | 19.9 | 32 | 0.0 | 0.0 | |
| Paraquat | 3 | DJ694 | 2.1 | ♀ | 3 | 40 | 18.9 | 30 | 40 | 15.6 | 25 | 40 | 14.1 | 20 | -9.6 | -20.0 | 0.0938 |
| Paraquat | 3 | DJ694 | 2.1 | ♀ | 4 | 40 | 55.9 | 90 | 24 | 56.7 | 84 | 39 | 45.8 | 84 | -17.9 | 0.0 | 0.1047 |
| Paraquat | 3 | DJ694 | 2.1 | ♀ | 5 | 40 | 17.8 | 25 | 40 | 15.6 | 20 | 40 | 14.5 | 20 | -7.2 | 0.0 | **0.0002** |
| Paraquat | 3 | DJ694 | 2.1 | ♀ | 6 | 37 | 56.4 | 84 | 39 | 77.6 | 130 | 39 | 41.4 | 90 | -26.6 | 0.0 | **0.0100** |
| Paraquat | 3 | DJ694 | 2.1 | ♀ | 7 | 42 | 50.9 | 84 | 42 | 65.7 | 118 | 40 | 46.2 | 110 | -9.2 | 0.0 | **0.0027** |
| Paraquat | 3 | DJ694 | 2.1 | ♂ | 1 | 40 | 25.2 | 40 | 40 | 16.6 | 22 | 40 | 19.3 | 30 | 0.0 | 0.0 | |
| Paraquat | 3 | DJ694 | 2.1 | ♂ | 2 | 40 | 27.5 | 57 | 38 | 19.1 | 22 | 40 | 17.4 | 22 | -9.1 | 0.0 | **0.0294** |
| Paraquat | 3 | DJ694 | 2.1 | ♂ | 3 | 40 | 17.9 | 25 | 40 | 18.0 | 30 | 40 | 17.6 | 30 | -1.4 | 0.0 | 0.7459 |
| Paraquat | 3 | DJ694 | 2.1 | ♂ | 4 | 40 | 35.5 | 66 | 45 | 21.5 | 38 | 50 | 36.6 | 74 | 3.0 | 12.1 | 0.4456 |
| Paraquat | 3 | DJ694 | 2.1 | ♂ | 5 | 40 | 17.8 | 25 | 40 | 17.5 | 25 | 40 | 19.1 | 30 | 7.7 | 20.0 | 0.1967 |
| Paraquat | 3 | DJ694 | 2.1 | ♂ | 6 | 34 | 40.2 | 84 | 41 | 20.0 | 34 | 38 | 24.1 | 42 | 0.0 | 0.0 | |
| Paraquat | 3 | DJ694 | 2.1 | ♂ | 7 | 41 | 39.0 | 84 | 41 | 22.5 | 34 | 40 | 28.7 | 66 | 0.0 | 0.0 | |
| Paraquat | 3 | DJ694 | 4.3 | ♀ | 1 | 40 | 30.8 | 43 | 40 | 30.9 | 56 | 40 | 20.9 | 31 | -32.3 | -27.9 | **<0.0001** |
| Paraquat | 3 | DJ694 | 4.3 | ♀ | 2 | 42 | 49.1 | 84 | 24 | 56.7 | 84 | 35 | 39.9 | 66 | -18.7 | -21.4 | **0.0006** |
| Paraquat | 3 | DJ694 | 4.3 | ♀ | 3 | 41 | 39.3 | 74 | 39 | 77.6 | 130 | 37 | 54.2 | 84 | 0.0 | 0.0 | |
| Paraquat | 3 | DJ694 | 4.3 | ♀ | 4 | 40 | 43.2 | 74 | 42 | 65.7 | 118 | 39 | 40.1 | 66 | -7.2 | -10.8 | 0.3893 |
| Paraquat | 3 | DJ694 | 4.3 | ♂ | 1 | 40 | 27.8 | 56 | 40 | 15.3 | 22 | 40 | 18.0 | 43 | 0.0 | 0.0 | |
| Paraquat | 3 | DJ694 | 4.3 | ♂ | 2 | 50 | 42.9 | 84 | 45 | 21.5 | 38 | 41 | 25.4 | 38 | 0.0 | 0.0 | |
| Paraquat | 3 | DJ694 | 4.3 | ♂ | 3 | 41 | 41.6 | 110 | 41 | 20.0 | 34 | 40 | 26.5 | 34 | 0.0 | 0.0 | |
| Paraquat | 3 | DJ694 | 4.3 | ♂ | 4 | 41 | 46.7 | 84 | 41 | 22.5 | 34 | 39 | 28.8 | 50 | 0.0 | 0.0 | |
| Paraquat | 3 | DJ694 | 9.1 | ♀ | 1 | 39 | 21.2 | 32 | 40 | 16.6 | 23 | 40 | 16.6 | 32 | 0.0 | 0.0 | |
| Paraquat | 3 | DJ694 | 9.1 | ♀ | 2 | 39 | 54.4 | 90 | 24 | 56.7 | 84 | 38 | 43.9 | 66 | -19.3 | -21.4 | **0.0006** |
| Paraquat | 3 | DJ694 | 9.1 | ♀ | 3 | 40 | 21.5 | 32 | 39 | 18.5 | 32 | 39 | 21.2 | 32 | 0.0 | 0.0 | |
| Paraquat | 3 | DJ694 | 9.1 | ♀ | 4 | 39 | 55.1 | 84 | 39 | 77.6 | 130 | 40 | 45.3 | 66 | -17.8 | -21.4 | **0.0064** |
| Paraquat | 3 | DJ694 | 9.1 | ♀ | 5 | 46 | 49.4 | 90 | 42 | 65.7 | 118 | 41 | 46.7 | 74 | -5.4 | -17.8 | **0.0002** |
| Paraquat | 3 | DJ694 | 9.1 | ♂ | 1 | 40 | 14.7 | 23 | 40 | 13.1 | 16 | 40 | 12.9 | 16 | -1.3 | 0.0 | **0.0042** |
| Paraquat | 3 | DJ694 | 9.1 | ♂ | 2 | 42 | 31.0 | 66 | 45 | 21.5 | 38 | 41 | 22.5 | 30 | 0.0 | -21.1 | |
| Paraquat | 3 | DJ694 | 9.1 | ♂ | 3 | 40 | 15.7 | 23 | 40 | 15.1 | 16 | 39 | 13.4 | 16 | -10.7 | 0.0 | **0.0061** |
| Paraquat | 3 | DJ694 | 9.1 | ♂ | 4 | 34 | 38.2 | 74 | 41 | 20.0 | 34 | 42 | 20.3 | 42 | 0.0 | 0.0 | |
| Paraquat | 3 | DJ694 | 9.1 | ♂ | 5 | 39 | 31.2 | 66 | 41 | 22.5 | 34 | 40 | 25.3 | 42 | 0.0 | 0.0 | |
| Paraquat | 3 | DJ694 | 3.2 | ♀ | 1 | 40 | 30.7 | 80 | 40 | 25.7 | 45 | 40 | 43.0 | 94 | 40.4 | 17.5 | **0.0049** |
| Paraquat | 3 | DJ694 | 3.2 | ♀ | 2 | 36 | 69.0 | 118 | 24 | 56.7 | 84 | 29 | 43.8 | 110 | -22.7 | 0.0 | **0.0006** |
| Paraquat | 3 | DJ694 | 3.2 | ♀ | 3 | 38 | 67.4 | 110 | 39 | 77.6 | 130 | 42 | 57.3 | 110 | -15.0 | 0.0 | **0.0135** |
| Paraquat | 3 | DJ694 | 3.2 | ♀ | 4 | 40 | 60.2 | 110 | 42 | 65.7 | 118 | 42 | 69.3 | 118 | 5.6 | 0.0 | 0.0540 |
| Paraquat | 3 | DJ694 | 3.2 | ♂ | 1 | 40 | 33.9 | 80 | 40 | 21.6 | 35 | 40 | 40.7 | 94 | 20.0 | 17.5 | 0.1150 |
| Paraquat | 3 | DJ694 | 3.2 | ♂ | 2 | 39 | 54.8 | 90 | 45 | 21.5 | 38 | 44 | 34.4 | 74 | 0.0 | 0.0 | |
| Paraquat | 3 | DJ694 | 3.2 | ♂ | 3 | 40 | 62.8 | 110 | 41 | 20.0 | 34 | 40 | 35.5 | 84 | 0.0 | 0.0 | |
| Paraquat | 3 | DJ694 | 3.2 | ♂ | 4 | 37 | 64.3 | 110 | 41 | 22.5 | 34 | 39 | 40.4 | 90 | 0.0 | 0.0 | |
| Paraquat | 3 | DJ694 | 10.1 | ♀ | 1 | 40 | 41.6 | 117 | 40 | 60.1 | 117 | 40 | 89.8 | 129 | 49.5 | 10.3 | **<0.0001** |
| Paraquat | 3 | DJ694 | 10.1 | ♀ | 2 | 40 | 23.7 | 41 | 40 | 16.6 | 23 | 40 | 23.3 | 41 | 0.0 | 0.0 | |
| Paraquat | 3 | DJ694 | 10.1 | ♀ | 3 | 28 | 66.1 | 90 | 24 | 56.7 | 84 | 34 | 43.9 | 110 | -22.5 | 22.2 | **0.0158** |

**Table S2**



| Stress | Age | Gal4 | UAS | Sex | Rep[1] | UAS n | UAS Mean | UAS Max[2] | Gal4 n | Gal4 Mean | Gal4 Max[2] | Gal4 + UAS n | Gal4 + UAS Mean | Gal4 + UAS Max[2] | % Extension Mean | % Extension Max[2] | *P* |
|---|---|---|---|---|---|---|---|---|---|---|---|---|---|---|---|---|---|
| Paraquat | 3 | DJ694 | 10.1 | ♀ | 4 | 40 | 24.3 | 59 | 39 | 18.5 | 32 | 39 | 26.0 | 59 | 6.9 | 0.0 | **0.0003** |
| Paraquat | 3 | DJ694 | 10.1 | ♀ | 5 | 39 | 88.3 | 118 | 39 | 77.6 | 130 | 46 | 50.8 | 110 | -34.5 | -6.8 | **0.0005** |
| Paraquat | 3 | DJ694 | 10.1 | ♀ | 6 | 37 | 78.3 | 110 | 42 | 65.7 | 118 | 30 | 57.5 | 110 | -12.4 | 0.0 | **0.0142** |
| Paraquat | 3 | DJ694 | 10.1 | ♂ | 1 | 40 | 63.7 | 117 | 39 | 33.1 | 81 | 40 | 79.0 | 141 | 24.0 | 20.5 | **0.0160** |
| Paraquat | 3 | DJ694 | 10.1 | ♂ | 2 | 40 | 23.3 | 41 | 40 | 13.1 | 16 | 34 | 18.7 | 32 | 0.0 | 0.0 | |
| Paraquat | 3 | DJ694 | 10.1 | ♂ | 3 | 39 | 47.5 | 90 | 45 | 21.5 | 38 | 46 | 35.3 | 74 | 0.0 | 0.0 | |
| Paraquat | 3 | DJ694 | 10.1 | ♂ | 4 | 40 | 28.3 | 41 | 40 | 15.1 | 16 | 37 | 22.1 | 41 | 0.0 | 0.0 | |
| Paraquat | 3 | DJ694 | 10.1 | ♂ | 5 | 39 | 73.1 | 130 | 41 | 20.0 | 34 | 26 | 40.2 | 84 | 0.0 | 0.0 | |
| Paraquat | 3 | DJ694 | 10.1 | ♂ | 6 | 42 | 59.0 | 110 | 41 | 22.5 | 34 | 39 | 34.7 | 90 | 0.0 | 0.0 | |
| Paraquat | 3 | DJ694 | 3.1 | ♀ | 1 | 40 | 31.4 | 45 | 40 | 20.7 | 35 | 40 | 22.6 | 35 | 0.0 | 0.0 | |
| Paraquat | 3 | DJ694 | 3.1 | ♀ | 2 | 40 | 33.4 | 64 | 40 | 40.1 | 76 | 40 | 28.5 | 64 | -14.7 | 0.0 | **0.0009** |
| Paraquat | 3 | DJ694 | 3.1 | ♀ | 3 | 40 | 16.0 | 25 | 40 | 15.6 | 25 | 40 | 13.0 | 15 | -16.8 | -40.0 | **0.0002** |
| Paraquat | 3 | DJ694 | 3.1 | ♀ | 4 | 40 | 46.1 | 74 | 24 | 56.7 | 84 | 37 | 53.7 | 90 | 0.0 | 7.1 | |
| Paraquat | 3 | DJ694 | 3.1 | ♀ | 5 | 40 | 18.0 | 30 | 40 | 15.6 | 20 | 40 | 13.6 | 20 | -12.8 | 0.0 | **0.0076** |
| Paraquat | 3 | DJ694 | 3.1 | ♀ | 6 | 36 | 50.6 | 84 | 39 | 77.6 | 130 | 41 | 42.1 | 110 | -16.8 | 0.0 | 0.1365 |
| Paraquat | 3 | DJ694 | 3.1 | ♀ | 7 | 44 | 49.4 | 74 | 42 | 65.7 | 118 | 40 | 47.1 | 90 | -4.7 | 0.0 | **0.0008** |
| Paraquat | 3 | DJ694 | 3.1 | ♂ | 1 | 40 | 30.5 | 57 | 40 | 16.6 | 22 | 40 | 27.1 | 63 | 0.0 | 10.5 | |
| Paraquat | 3 | DJ694 | 3.1 | ♂ | 2 | 40 | 23.3 | 39 | 40 | 18.9 | 25 | 40 | 21.5 | 35 | 0.0 | 0.0 | |
| Paraquat | 3 | DJ694 | 3.1 | ♂ | 3 | 40 | 18.8 | 40 | 40 | 18.0 | 30 | 40 | 20.1 | 40 | 7.2 | 0.0 | 0.1637 |
| Paraquat | 3 | DJ694 | 3.1 | ♂ | 4 | 45 | 41.4 | 90 | 45 | 21.5 | 38 | 54 | 42.6 | 84 | 2.7 | 0.0 | 0.7185 |
| Paraquat | 3 | DJ694 | 3.1 | ♂ | 5 | 40 | 19.3 | 25 | 40 | 17.5 | 25 | 40 | 19.9 | 30 | 3.2 | 20.0 | 0.0502 |
| Paraquat | 3 | DJ694 | 3.1 | ♂ | 6 | 39 | 41.6 | 84 | 41 | 20.0 | 34 | 39 | 33.5 | 90 | 0.0 | 7.1 | |
| Paraquat | 3 | DJ694 | 3.1 | ♂ | 7 | 41 | 44.3 | 84 | 41 | 22.5 | 34 | 44 | 37.1 | 58 | 0.0 | 0.0 | |
| Paraquat | 3 | DJ694 | 4.4 | ♀ | 1 | 40 | 35.5 | 58 | 40 | 45.1 | 67 | 40 | 55.8 | 97 | 23.8 | 44.8 | **0.0047** |
| Paraquat | 3 | DJ694 | 4.4 | ♀ | 2 | 40 | 45.4 | 69 | 40 | 30.9 | 56 | 40 | 28.1 | 56 | -9.0 | 0.0 | 0.3708 |
| Paraquat | 3 | DJ694 | 4.4 | ♀ | 3 | 40 | 26.5 | 41 | 40 | 16.6 | 23 | 40 | 21.0 | 32 | 0.0 | 0.0 | |
| Paraquat | 3 | DJ694 | 4.4 | ♀ | 4 | 36 | 53.7 | 90 | 24 | 56.7 | 84 | 40 | 52.8 | 110 | -1.7 | 22.2 | 0.5266 |
| Paraquat | 3 | DJ694 | 4.4 | ♀ | 5 | 40 | 28.2 | 48 | 39 | 18.5 | 32 | 40 | 23.0 | 32 | 0.0 | 0.0 | |
| Paraquat | 3 | DJ694 | 4.4 | ♀ | 6 | 40 | 67.1 | 110 | 39 | 77.6 | 130 | 39 | 46.5 | 74 | -30.7 | -32.7 | **<0.0001** |
| Paraquat | 3 | DJ694 | 4.4 | ♀ | 7 | 41 | 57.2 | 90 | 42 | 65.7 | 118 | 46 | 50.6 | 110 | -11.5 | 0.0 | **0.0091** |
| Paraquat | 3 | DJ694 | 4.4 | ♂ | 1 | 40 | 38.7 | 67 | 40 | 25.5 | 34 | 40 | 34.2 | 77 | 0.0 | 14.9 | |
| Paraquat | 3 | DJ694 | 4.4 | ♂ | 2 | 40 | 39.2 | 81 | 40 | 15.3 | 22 | 40 | 20.1 | 31 | 0.0 | 0.0 | |
| Paraquat | 3 | DJ694 | 4.4 | ♂ | 3 | 39 | 19.1 | 32 | 40 | 13.1 | 16 | 40 | 17.2 | 32 | 0.0 | 0.0 | |
| Paraquat | 3 | DJ694 | 4.4 | ♂ | 4 | 41 | 37.7 | 84 | 45 | 21.5 | 38 | 42 | 29.3 | 42 | 0.0 | 0.0 | |
| Paraquat | 3 | DJ694 | 4.4 | ♂ | 5 | 40 | 19.3 | 32 | 40 | 15.1 | 16 | 40 | 16.5 | 23 | 0.0 | 0.0 | |
| Paraquat | 3 | DJ694 | 4.4 | ♂ | 6 | 40 | 57.9 | 110 | 41 | 20.0 | 34 | 39 | 24.2 | 34 | 0.0 | 0.0 | |
| Paraquat | 3 | DJ694 | 4.4 | ♂ | 7 | 40 | 43.0 | 110 | 41 | 22.5 | 34 | 41 | 26.7 | 58 | 0.0 | 0.0 | |
| Paraquat | 3 | D42 | 2.1 | ♀ | 1 | 40 | 29.5 | 51 | 40 | 23.3 | 35 | 40 | 24.5 | 45 | 0.0 | 0.0 | |
| Paraquat | 3 | D42 | 2.1 | ♀ | 2 | 40 | 29.4 | 47 | 40 | 28.1 | 52 | 40 | 27.3 | 42 | -3.1 | -10.6 | 0.2322 |
| Paraquat | 3 | D42 | 2.1 | ♀ | 3 | 40 | 18.9 | 30 | 40 | 16.6 | 20 | 40 | 15.8 | 20 | -5.3 | 0.0 | **0.0010** |
| Paraquat | 3 | D42 | 2.1 | ♀ | 4 | 40 | 55.9 | 90 | 40 | 52.6 | 110 | 41 | 24.3 | 42 | -53.7 | -53.3 | **<0.0001** |
| Paraquat | 3 | D42 | 2.1 | ♀ | 5 | 40 | 17.8 | 25 | 40 | 17.5 | 25 | 40 | 16.9 | 25 | -3.6 | 0.0 | 0.2549 |
| Paraquat | 3 | D42 | 2.1 | ♀ | 6 | 37 | 56.4 | 84 | 41 | 61.9 | 110 | 41 | 24.8 | 46 | -56.0 | -45.2 | **<0.0001** |
| Paraquat | 3 | D42 | 2.1 | ♀ | 7 | 42 | 50.9 | 84 | 41 | 58.4 | 110 | 40 | 21.7 | 38 | -57.3 | -54.8 | **<0.0001** |
| Paraquat | 3 | D42 | 2.1 | ♂ | 1 | 40 | 25.2 | 40 | 40 | 22.8 | 30 | 40 | 23.5 | 40 | 0.0 | 0.0 | |
| Paraquat | 3 | D42 | 2.1 | ♂ | 2 | 40 | 27.5 | 57 | 40 | 26.3 | 47 | 40 | 20.6 | 37 | -21.4 | -21.3 | **0.0004** |
| Paraquat | 3 | D42 | 2.1 | ♂ | 3 | 40 | 17.9 | 25 | 40 | 16.3 | 20 | 40 | 14.6 | 20 | -10.0 | 0.0 | **0.0002** |
| Paraquat | 3 | D42 | 2.1 | ♂ | 4 | 40 | 35.5 | 66 | 40 | 33.0 | 74 | 38 | 33.4 | 58 | 0.0 | -12.1 | |
| Paraquat | 3 | D42 | 2.1 | ♂ | 5 | 40 | 17.8 | 25 | 40 | 15.9 | 25 | 40 | 14.4 | 20 | -9.4 | -20.0 | **0.0003** |

### Table S2



| Stress | Age | Gal4 | UAS | Sex | Rep[1] | UAS n | UAS Mean | UAS Max[2] | Gal4 n | Gal4 Mean | Gal4 Max[2] | Gal4 + UAS n | Gal4 + UAS Mean | Gal4 + UAS Max[2] | % Extension Mean | % Extension Max[2] | *P* |
|---|---|---|---|---|---|---|---|---|---|---|---|---|---|---|---|---|---|
| Paraquat | 3 | D42 | 2.1 | ♂ | 6 | 34 | 40.2 | 84 | 40 | 28.7 | 58 | 40 | 44.7 | 74 | 11.1 | 0.0 | 0.4630 |
| Paraquat | 3 | D42 | 2.1 | ♂ | 7 | 41 | 39.0 | 84 | 40 | 38.6 | 84 | 31 | 45.6 | 84 | 17.0 | 0.0 | 0.1017 |
| Paraquat | 3 | D42 | 4.3 | ♀ | 1 | 40 | 30.8 | 43 | 40 | 43.8 | 69 | 40 | 33.2 | 69 | 0.0 | 0.0 | |
| Paraquat | 3 | D42 | 4.3 | ♀ | 2 | 42 | 49.1 | 84 | 40 | 52.6 | 110 | 40 | 35.3 | 66 | -28.3 | -21.4 | **0.0003** |
| Paraquat | 3 | D42 | 4.3 | ♀ | 3 | 41 | 39.3 | 74 | 41 | 61.9 | 110 | 40 | 31.4 | 50 | -20.0 | -32.4 | **0.0098** |
| Paraquat | 3 | D42 | 4.3 | ♀ | 4 | 40 | 43.2 | 74 | 41 | 58.4 | 110 | 40 | 27.8 | 58 | -35.8 | -21.6 | **0.0001** |
| Paraquat | 3 | D42 | 4.3 | ♂ | 1 | 40 | 27.8 | 56 | 40 | 31.1 | 56 | 40 | 27.1 | 56 | -2.5 | 0.0 | 0.2014 |
| Paraquat | 3 | D42 | 4.3 | ♂ | 2 | 50 | 42.9 | 84 | 40 | 33.0 | 74 | 41 | 44.1 | 84 | 2.9 | 0.0 | **0.0011** |
| Paraquat | 3 | D42 | 4.3 | ♂ | 3 | 41 | 41.6 | 110 | 40 | 28.7 | 58 | 39 | 44.1 | 90 | 6.0 | 0.0 | 0.5716 |
| Paraquat | 3 | D42 | 4.3 | ♂ | 4 | 41 | 46.7 | 84 | 40 | 38.6 | 84 | 41 | 39.4 | 74 | 0.0 | -11.9 | |
| Paraquat | 3 | D42 | 9.1 | ♀ | 1 | 39 | 21.2 | 32 | 40 | 24.2 | 41 | 42 | 27.1 | 48 | 12.1 | 17.1 | **0.0028** |
| Paraquat | 3 | D42 | 9.1 | ♀ | 2 | 39 | 54.4 | 90 | 40 | 52.6 | 110 | 40 | 28.3 | 58 | -46.1 | -35.6 | **<0.0001** |
| Paraquat | 3 | D42 | 9.1 | ♀ | 3 | 40 | 21.5 | 32 | 40 | 25.8 | 41 | 41 | 30.0 | 48 | 16.3 | 17.1 | **0.0495** |
| Paraquat | 3 | D42 | 9.1 | ♀ | 4 | 39 | 55.1 | 84 | 41 | 61.9 | 110 | 40 | 30.4 | 58 | -44.9 | -31.0 | **<0.0001** |
| Paraquat | 3 | D42 | 9.1 | ♀ | 5 | 46 | 49.4 | 90 | 41 | 58.4 | 110 | 42 | 25.4 | 42 | -48.5 | -53.3 | **<0.0001** |
| Paraquat | 3 | D42 | 9.1 | ♂ | 1 | 40 | 14.7 | 23 | 40 | 16.3 | 23 | 40 | 14.9 | 23 | 0.0 | 0.0 | |
| Paraquat | 3 | D42 | 9.1 | ♂ | 2 | 42 | 31.0 | 66 | 40 | 33.0 | 74 | 43 | 37.6 | 74 | 14.0 | 0.0 | **0.0066** |
| Paraquat | 3 | D42 | 9.1 | ♂ | 3 | 40 | 15.7 | 23 | 40 | 20.2 | 32 | 40 | 17.4 | 32 | 0.0 | 0.0 | |
| Paraquat | 3 | D42 | 9.1 | ♂ | 4 | 34 | 38.2 | 74 | 40 | 28.7 | 58 | 40 | 34.5 | 74 | 0.0 | 0.0 | |
| Paraquat | 3 | D42 | 9.1 | ♂ | 5 | 39 | 31.2 | 66 | 40 | 38.6 | 84 | 39 | 33.2 | 58 | 0.0 | -12.1 | |
| Paraquat | 3 | D42 | 3.2 | ♀ | 1 | 40 | 30.7 | 80 | 40 | 30.1 | 55 | 40 | 32.0 | 75 | 4.3 | 0.0 | 0.3500 |
| Paraquat | 3 | D42 | 3.2 | ♀ | 2 | 36 | 69.0 | 118 | 40 | 52.6 | 110 | 40 | 25.0 | 42 | -52.4 | -61.8 | **<0.0001** |
| Paraquat | 3 | D42 | 3.2 | ♀ | 3 | 38 | 67.4 | 110 | 41 | 61.9 | 110 | 40 | 20.6 | 30 | -66.7 | -72.7 | **<0.0001** |
| Paraquat | 3 | D42 | 3.2 | ♀ | 4 | 40 | 60.2 | 110 | 41 | 58.4 | 110 | 40 | 19.7 | 38 | -66.3 | -65.5 | **<0.0001** |
| Paraquat | 3 | D42 | 3.2 | ♂ | 1 | 40 | 33.9 | 80 | 40 | 25.6 | 45 | 40 | 36.6 | 88 | 7.9 | 10.0 | **0.0031** |
| Paraquat | 3 | D42 | 3.2 | ♂ | 2 | 39 | 54.8 | 90 | 40 | 33.0 | 74 | 42 | 45.4 | 110 | 0.0 | 22.2 | |
| Paraquat | 3 | D42 | 3.2 | ♂ | 3 | 40 | 62.8 | 110 | 40 | 28.7 | 58 | 39 | 36.3 | 74 | 0.0 | 0.0 | |
| Paraquat | 3 | D42 | 3.2 | ♂ | 4 | 37 | 64.3 | 110 | 40 | 38.6 | 84 | 41 | 49.9 | 110 | 0.0 | 0.0 | |
| Paraquat | 3 | D42 | 10.1 | ♀ | 1 | 40 | 41.6 | 117 | 40 | 53.0 | 117 | 40 | 45.2 | 105 | 0.0 | -10.3 | |
| Paraquat | 3 | D42 | 10.1 | ♀ | 2 | 40 | 23.7 | 41 | 40 | 24.2 | 41 | 39 | 31.3 | 59 | 29.7 | 43.9 | **0.0008** |
| Paraquat | 3 | D42 | 10.1 | ♀ | 3 | 28 | 66.1 | 90 | 40 | 52.6 | 110 | 44 | 47.2 | 110 | -10.1 | 0.0 | 0.1142 |
| Paraquat | 3 | D42 | 10.1 | ♀ | 4 | 40 | 24.3 | 59 | 40 | 25.8 | 41 | 40 | 32.2 | 48 | 24.6 | 0.0 | **0.0014** |
| Paraquat | 3 | D42 | 10.1 | ♀ | 5 | 39 | 88.3 | 118 | 41 | 61.9 | 110 | 40 | 28.9 | 66 | -53.3 | -40.0 | **<0.0001** |
| Paraquat | 3 | D42 | 10.1 | ♀ | 6 | 37 | 78.3 | 110 | 41 | 58.4 | 110 | 40 | 31.1 | 84 | -46.9 | -23.6 | **<0.0001** |
| Paraquat | 3 | D42 | 10.1 | ♂ | 1 | 40 | 63.7 | 117 | 40 | 54.4 | 105 | 40 | 58.7 | 117 | 0.0 | 0.0 | |
| Paraquat | 3 | D42 | 10.1 | ♂ | 2 | 40 | 23.3 | 41 | 40 | 16.3 | 23 | 41 | 25.9 | 59 | 10.8 | 43.9 | 0.2396 |
| Paraquat | 3 | D42 | 10.1 | ♂ | 3 | 39 | 47.5 | 90 | 40 | 33.0 | 74 | 46 | 49.8 | 110 | 4.7 | 22.2 | **0.0005** |
| Paraquat | 3 | D42 | 10.1 | ♂ | 4 | 40 | 28.3 | 41 | 40 | 20.2 | 32 | 40 | 24.6 | 41 | 0.0 | 0.0 | |
| Paraquat | 3 | D42 | 10.1 | ♂ | 5 | 39 | 73.1 | 130 | 40 | 28.7 | 58 | 41 | 46.9 | 110 | 0.0 | 0.0 | |
| Paraquat | 3 | D42 | 10.1 | ♂ | 6 | 42 | 59.0 | 110 | 40 | 38.6 | 84 | 39 | 35.5 | 90 | -7.9 | 0.0 | **0.0001** |
| Paraquat | 3 | D42 | 3.1 | ♀ | 1 | 40 | 31.4 | 45 | 40 | 23.3 | 35 | 40 | 29.6 | 51 | 0.0 | 13.3 | |
| Paraquat | 3 | D42 | 3.1 | ♀ | 2 | 40 | 33.4 | 64 | 38 | 37.2 | 76 | 40 | 30.6 | 64 | -8.4 | 0.0 | 0.0687 |
| Paraquat | 3 | D42 | 3.1 | ♀ | 3 | 40 | 16.0 | 25 | 40 | 16.6 | 20 | 40 | 15.6 | 25 | -2.3 | 0.0 | 0.2561 |
| Paraquat | 3 | D42 | 3.1 | ♀ | 4 | 40 | 46.1 | 74 | 40 | 52.6 | 110 | 50 | 40.9 | 74 | -11.3 | 0.0 | 0.0505 |
| Paraquat | 3 | D42 | 3.1 | ♀ | 5 | 40 | 18.0 | 30 | 40 | 17.5 | 25 | 40 | 18.9 | 30 | 4.9 | 0.0 | 0.1826 |
| Paraquat | 3 | D42 | 3.1 | ♀ | 6 | 36 | 50.6 | 84 | 41 | 61.9 | 110 | 40 | 30.7 | 58 | -39.3 | -31.0 | **<0.0001** |
| Paraquat | 3 | D42 | 3.1 | ♀ | 7 | 44 | 49.4 | 74 | 41 | 58.4 | 110 | 39 | 26.9 | 50 | -45.5 | -32.4 | **<0.0001** |
| Paraquat | 3 | D42 | 3.1 | ♂ | 1 | 40 | 30.5 | 57 | 40 | 22.8 | 30 | 40 | 39.7 | 69 | 30.1 | 21.1 | **0.0080** |
| Paraquat | 3 | D42 | 3.1 | ♂ | 2 | 40 | 23.3 | 39 | 38 | 35.0 | 70 | 40 | 44.6 | 82 | 27.3 | 17.1 | **0.0195** |

### Table S2

| Stress | Age | Gal4 | UAS | Sex | Rep[1] | UAS | | | Gal4 | | | Gal4 + UAS | | | % Extension | | *P* |
|---|---|---|---|---|---|---|---|---|---|---|---|---|---|---|---|---|---|
| | | | | | | n | Mean | Max[2] | n | Mean | Max[2] | n | Mean | Max[2] | Mean | Max[2] | |
| Paraquat | 3 | D42 | 3.1 | ♂ | 3 | 40 | 18.8 | 40 | 40 | 16.3 | 20 | 40 | 24.5 | 49 | 30.4 | 22.5 | **0.0013** |
| Paraquat | 3 | D42 | 3.1 | ♂ | 4 | 45 | 41.4 | 90 | 40 | 33.0 | 74 | 40 | 51.3 | 110 | 23.7 | 22.2 | **0.0004** |
| Paraquat | 3 | D42 | 3.1 | ♂ | 5 | 40 | 19.3 | 25 | 40 | 15.9 | 25 | 40 | 24.5 | 40 | 27.1 | 60.0 | **0.0010** |
| Paraquat | 3 | D42 | 3.1 | ♂ | 6 | 39 | 41.6 | 84 | 40 | 28.7 | 58 | 37 | 55.5 | 110 | 33.2 | 31.0 | **0.0106** |
| Paraquat | 3 | D42 | 3.1 | ♂ | 7 | 41 | 44.3 | 84 | 40 | 38.6 | 84 | 44 | 52.6 | 110 | 18.6 | 31.0 | **0.0059** |
| Paraquat | 3 | D42 | 4.4 | ♀ | 1 | 40 | 35.5 | 58 | 40 | 37.7 | 58 | 40 | 49.5 | 72 | 31.4 | 24.1 | **0.0001** |
| Paraquat | 3 | D42 | 4.4 | ♀ | 2 | 40 | 45.4 | 69 | 40 | 43.8 | 69 | 40 | 46.1 | 69 | 1.4 | 0.0 | 0.6113 |
| Paraquat | 3 | D42 | 4.4 | ♀ | 3 | 40 | 26.5 | 41 | 40 | 24.2 | 41 | 46 | 27.6 | 41 | 4.4 | 0.0 | 0.1040 |
| Paraquat | 3 | D42 | 4.4 | ♀ | 4 | 36 | 53.7 | 90 | 40 | 52.6 | 110 | 42 | 44.8 | 84 | -14.8 | -6.7 | **0.0320** |
| Paraquat | 3 | D42 | 4.4 | ♀ | 5 | 40 | 28.2 | 48 | 40 | 25.8 | 41 | 47 | 27.2 | 41 | 0.0 | 0.0 | |
| Paraquat | 3 | D42 | 4.4 | ♀ | 6 | 40 | 67.1 | 110 | 41 | 61.9 | 110 | 41 | 46.8 | 74 | -24.4 | -32.7 | **0.0005** |
| Paraquat | 3 | D42 | 4.4 | ♀ | 7 | 41 | 57.2 | 90 | 41 | 58.4 | 110 | 40 | 46.1 | 84 | -19.5 | -6.7 | **0.0075** |
| Paraquat | 3 | D42 | 4.4 | ♂ | 1 | 40 | 38.7 | 67 | 40 | 33.3 | 58 | 40 | 49.2 | 87 | 27.1 | 29.9 | **0.0038** |
| Paraquat | 3 | D42 | 4.4 | ♂ | 2 | 40 | 39.2 | 81 | 40 | 31.1 | 56 | 40 | 36.3 | 69 | 0.0 | 0.0 | |
| Paraquat | 3 | D42 | 4.4 | ♂ | 3 | 39 | 19.1 | 32 | 40 | 16.3 | 23 | 42 | 18.5 | 32 | 0.0 | 0.0 | |
| Paraquat | 3 | D42 | 4.4 | ♂ | 4 | 41 | 37.7 | 84 | 40 | 33.0 | 74 | 39 | 33.0 | 74 | 0.0 | 0.0 | |
| Paraquat | 3 | D42 | 4.4 | ♂ | 5 | 40 | 19.3 | 32 | 40 | 20.2 | 32 | 41 | 19.5 | 32 | 0.0 | 0.0 | |
| Paraquat | 3 | D42 | 4.4 | ♂ | 6 | 40 | 57.9 | 110 | 40 | 28.7 | 58 | 40 | 37.8 | 74 | 0.0 | 0.0 | |
| Paraquat | 3 | D42 | 4.4 | ♂ | 7 | 40 | 43.0 | 110 | 40 | 38.6 | 84 | 38 | 33.4 | 58 | -13.4 | -31.0 | **0.0268** |
| Paraquat | 10 | daGal4 | 2.1 | ♀ | 1 | 40 | 25.3 | 52 | 36 | 15.8 | 32 | 40 | 14.3 | 22 | -9.5 | -31.3 | 0.3221 |
| Paraquat | 10 | daGal4 | 2.1 | ♀ | 2 | 40 | 17.9 | 26 | 40 | 14.7 | 22 | 42 | 13.1 | 26 | -10.6 | 0.0 | 0.1758 |
| Paraquat | 10 | daGal4 | 2.1 | ♀ | 3 | 36 | 18.6 | 26 | 40 | 19.1 | 30 | 41 | 17.4 | 30 | -6.1 | 0.0 | 0.2041 |
| Paraquat | 10 | daGal4 | 2.1 | ♀ | 4 | 41 | 19.3 | 22 | 45 | 17.3 | 26 | 44 | 15.8 | 22 | -8.5 | 0.0 | **0.0001** |
| Paraquat | 10 | daGal4 | 2.1 | ♂ | 1 | 40 | 24.5 | 47 | 38 | 20.2 | 42 | 40 | 21.8 | 47 | 0.0 | 0.0 | |
| Paraquat | 10 | daGal4 | 2.1 | ♂ | 2 | 41 | 17.6 | 26 | 40 | 17.5 | 30 | 42 | 13.7 | 22 | -21.5 | -15.4 | **0.0009** |
| Paraquat | 10 | daGal4 | 2.1 | ♂ | 3 | 40 | 15.4 | 22 | 45 | 13.3 | 18 | 42 | 13.4 | 22 | 0.0 | 0.0 | |
| Paraquat | 10 | daGal4 | 2.1 | ♂ | 4 | 40 | 15.9 | 30 | 45 | 14.4 | 18 | 43 | 16.2 | 42 | 1.8 | 40.0 | 0.4360 |
| Paraquat | 10 | daGal4 | 4.3 | ♀ | 1 | 40 | 20.8 | 31 | 35 | 15.4 | 22 | 40 | 17.4 | 31 | 0.0 | 0.0 | |
| Paraquat | 10 | daGal4 | 4.3 | ♀ | 2 | 40 | 18.2 | 33 | 40 | 13.2 | 25 | 40 | 16.0 | 25 | 0.0 | 0.0 | |
| Paraquat | 10 | daGal4 | 4.3 | ♀ | 3 | 40 | 18.5 | 26 | 40 | 14.7 | 22 | 40 | 14.1 | 30 | -4.1 | 15.4 | **0.0015** |
| Paraquat | 10 | daGal4 | 4.3 | ♀ | 4 | 40 | 15.8 | 25 | 40 | 13.9 | 25 | 40 | 16.1 | 25 | 2.2 | 0.0 | **0.0225** |
| Paraquat | 10 | daGal4 | 4.3 | ♀ | 5 | 42 | 24.3 | 42 | 40 | 19.1 | 30 | 40 | 18.9 | 35 | -1.3 | 0.0 | **0.0010** |
| Paraquat | 10 | daGal4 | 4.3 | ♀ | 6 | 42 | 19.5 | 26 | 45 | 17.3 | 26 | 41 | 19.1 | 26 | 0.0 | 0.0 | |
| Paraquat | 10 | daGal4 | 4.3 | ♂ | 1 | 40 | 20.5 | 31 | 40 | 18.2 | 31 | 40 | 23.6 | 43 | 15.1 | 38.7 | **0.0010** |
| Paraquat | 10 | daGal4 | 4.3 | ♂ | 2 | 40 | 13.4 | 20 | 40 | 12.7 | 16 | 40 | 14.8 | 25 | 10.1 | 25.0 | **0.0115** |
| Paraquat | 10 | daGal4 | 4.3 | ♂ | 3 | 58 | 18.7 | 26 | 40 | 17.5 | 30 | 44 | 13.6 | 22 | -22.0 | -15.4 | **0.0015** |
| Paraquat | 10 | daGal4 | 4.3 | ♂ | 4 | 40 | 14.0 | 20 | 40 | 13.7 | 25 | 40 | 13.4 | 20 | -1.6 | 0.0 | 0.5721 |
| Paraquat | 10 | daGal4 | 4.3 | ♂ | 5 | 42 | 16.0 | 35 | 45 | 13.3 | 18 | 14 | 15.7 | 18 | 0.0 | 0.0 | |
| Paraquat | 10 | daGal4 | 4.3 | ♂ | 6 | 40 | 13.6 | 22 | 45 | 14.4 | 18 | 48 | 16.6 | 30 | 16.0 | 36.4 | **0.0136** |
| Paraquat | 10 | daGal4 | 9.1 | ♀ | 1 | 40 | 19.0 | 27 | 39 | 14.3 | 21 | 40 | 16.5 | 40 | 0.0 | 48.1 | |
| Paraquat | 10 | daGal4 | 9.1 | ♀ | 2 | 41 | 18.7 | 30 | 40 | 14.7 | 22 | 41 | 15.6 | 26 | 0.0 | 0.0 | |
| Paraquat | 10 | daGal4 | 9.1 | ♀ | 3 | 41 | 18.2 | 27 | 40 | 14.9 | 21 | 30 | 12.5 | 21 | -15.7 | 0.0 | **0.0072** |
| Paraquat | 10 | daGal4 | 9.1 | ♀ | 4 | 42 | 22.8 | 35 | 40 | 19.1 | 30 | 41 | 18.6 | 38 | -2.7 | 8.6 | **0.0050** |
| Paraquat | 10 | daGal4 | 9.1 | ♀ | 5 | 42 | 21.0 | 30 | 45 | 17.3 | 26 | 41 | 16.0 | 18 | -7.7 | -30.8 | 0.1465 |
| Paraquat | 10 | daGal4 | 9.1 | ♂ | 1 | 40 | 15.6 | 21 | 41 | 11.7 | 15 | 40 | 10.5 | 15 | -10.3 | 0.0 | **0.0094** |
| Paraquat | 10 | daGal4 | 9.1 | ♂ | 2 | 38 | 16.3 | 30 | 40 | 17.5 | 30 | 42 | 13.7 | 22 | -16.1 | -26.7 | **0.0017** |
| Paraquat | 10 | daGal4 | 9.1 | ♂ | 3 | 40 | 17.7 | 27 | 39 | 13.4 | 21 | 40 | 10.9 | 15 | -18.7 | -28.6 | **0.0017** |
| Paraquat | 10 | daGal4 | 9.1 | ♂ | 4 | 40 | 14.1 | 22 | 45 | 13.3 | 18 | 40 | 12.5 | 14 | -6.1 | -22.2 | **0.0059** |
| Paraquat | 10 | daGal4 | 9.1 | ♂ | 5 | 43 | 13.1 | 22 | 45 | 14.4 | 18 | 44 | 11.5 | 18 | -12.4 | 0.0 | **0.0096** |

### Table S2



| Stress | Age | Gal4 | UAS | Sex | Rep[1] | UAS n | UAS Mean | UAS Max[2] | Gal4 n | Gal4 Mean | Gal4 Max[2] | Gal4 + UAS n | Gal4 + UAS Mean | Gal4 + UAS Max[2] | % Extension Mean | % Extension Max[2] | P |
|---|---|---|---|---|---|---|---|---|---|---|---|---|---|---|---|---|---|
| Paraquat | 10 | daGal4 | 3.2 | ♀ | 1 | 38 | 19.1 | 40 | 40 | 15.3 | 26 | 40 | 15.1 | 26 | -1.0 | 0.0 | **0.0198** |
| Paraquat | 10 | daGal4 | 3.2 | ♀ | 2 | 40 | 19.2 | 33 | 40 | 13.2 | 25 | 40 | 16.6 | 25 | 0.0 | 0.0 | |
| Paraquat | 10 | daGal4 | 3.2 | ♀ | 3 | 35 | 17.1 | 30 | 40 | 14.7 | 22 | 40 | 20.5 | 35 | 19.7 | 16.7 | **0.0387** |
| Paraquat | 10 | daGal4 | 3.2 | ♀ | 4 | 40 | 17.3 | 25 | 40 | 13.9 | 25 | 40 | 17.8 | 33 | 3.0 | 32.0 | **0.0011** |
| Paraquat | 10 | daGal4 | 3.2 | ♀ | 5 | 40 | 22.2 | 42 | 40 | 19.1 | 30 | 43 | 24.5 | 35 | 10.4 | 0.0 | **0.0002** |
| Paraquat | 10 | daGal4 | 3.2 | ♀ | 6 | 39 | 19.9 | 35 | 45 | 17.3 | 26 | 33 | 23.1 | 42 | 16.2 | 20.0 | **0.0020** |
| Paraquat | 10 | daGal4 | 3.2 | ♂ | 1 | 38 | 24.3 | 45 | 40 | 19.9 | 45 | 40 | 44.7 | 75 | 83.4 | 66.7 | **<0.0001** |
| Paraquat | 10 | daGal4 | 3.2 | ♂ | 2 | 40 | 18.3 | 25 | 40 | 12.7 | 16 | 40 | 17.5 | 48 | 0.0 | 92.0 | |
| Paraquat | 10 | daGal4 | 3.2 | ♂ | 3 | 38 | 25.0 | 42 | 40 | 17.5 | 30 | 37 | 25.1 | 61 | 0.4 | 45.2 | **0.0014** |
| Paraquat | 10 | daGal4 | 3.2 | ♂ | 4 | 40 | 18.4 | 33 | 40 | 13.7 | 25 | 40 | 18.2 | 33 | 0.0 | 0.0 | |
| Paraquat | 10 | daGal4 | 3.2 | ♂ | 5 | 38 | 24.3 | 61 | 45 | 13.3 | 18 | 37 | 33.8 | 70 | 38.9 | 14.8 | **0.0082** |
| Paraquat | 10 | daGal4 | 3.2 | ♂ | 6 | 39 | 29.8 | 61 | 45 | 14.4 | 18 | 37 | 34.4 | 70 | 15.4 | 14.8 | 0.1981 |
| Paraquat | 10 | daGal4 | 10.1 | ♀ | 1 | 29 | 17.3 | 22 | 31 | 17.2 | 17 | 40 | 18.2 | 28 | 4.8 | 27.3 | 0.2535 |
| Paraquat | 10 | daGal4 | 10.1 | ♀ | 2 | 40 | 18.9 | 27 | 39 | 14.3 | 21 | 40 | 14.2 | 27 | -1.3 | 0.0 | 0.7502 |
| Paraquat | 10 | daGal4 | 10.1 | ♀ | 3 | 36 | 22.9 | 38 | 40 | 14.7 | 22 | 38 | 25.3 | 61 | 10.1 | 60.5 | 0.4944 |
| Paraquat | 10 | daGal4 | 10.1 | ♀ | 4 | 40 | 18.8 | 27 | 40 | 14.9 | 21 | 36 | 17.2 | 27 | 0.0 | 0.0 | |
| Paraquat | 10 | daGal4 | 10.1 | ♀ | 5 | 36 | 27.9 | 61 | 40 | 19.1 | 30 | 40 | 23.4 | 42 | 0.0 | 0.0 | |
| Paraquat | 10 | daGal4 | 10.1 | ♀ | 6 | 37 | 24.2 | 38 | 45 | 17.3 | 26 | 43 | 23.9 | 61 | 0.0 | 60.5 | |
| Paraquat | 10 | daGal4 | 10.1 | ♂ | 1 | 37 | 18.8 | 28 | 31 | 20.1 | 42 | 38 | 39.7 | 89 | 97.0 | 111.9 | **<0.0001** |
| Paraquat | 10 | daGal4 | 10.1 | ♂ | 2 | 40 | 25.9 | 40 | 41 | 11.7 | 15 | 40 | 14.8 | 21 | 0.0 | 0.0 | |
| Paraquat | 10 | daGal4 | 10.1 | ♂ | 3 | 38 | 25.9 | 61 | 40 | 17.5 | 30 | 40 | 27.2 | 70 | 5.1 | 14.8 | 0.8843 |
| Paraquat | 10 | daGal4 | 10.1 | ♂ | 4 | 40 | 20.6 | 40 | 39 | 13.4 | 21 | 39 | 19.7 | 40 | 0.0 | 0.0 | |
| Paraquat | 10 | daGal4 | 10.1 | ♂ | 5 | 37 | 25.4 | 42 | 45 | 13.3 | 18 | 41 | 28.5 | 70 | 12.3 | 66.7 | 0.6356 |
| Paraquat | 10 | daGal4 | 10.1 | ♂ | 6 | 39 | 24.7 | 42 | 45 | 14.4 | 18 | 40 | 20.4 | 38 | 0.0 | 0.0 | |
| Paraquat | 10 | daGal4 | 3.1 | ♀ | 1 | 40 | 12.5 | 22 | 40 | 12.7 | 22 | 40 | 14.1 | 30 | 10.8 | 36.4 | 0.2253 |
| Paraquat | 10 | daGal4 | 3.1 | ♀ | 2 | 43 | 14.0 | 22 | 40 | 14.7 | 22 | 39 | 16.1 | 22 | 9.2 | 0.0 | **0.0228** |
| Paraquat | 10 | daGal4 | 3.1 | ♀ | 3 | 40 | 17.9 | 30 | 40 | 19.1 | 30 | 44 | 17.7 | 30 | -1.0 | 0.0 | 0.2319 |
| Paraquat | 10 | daGal4 | 3.1 | ♀ | 4 | 44 | 18.2 | 35 | 45 | 17.3 | 26 | 42 | 17.5 | 38 | 0.0 | 8.6 | |
| Paraquat | 10 | daGal4 | 3.1 | ♂ | 1 | 33 | 23.5 | 34 | 39 | 23.6 | 61 | 40 | 52.9 | 93 | 123.9 | 52.5 | **<0.0001** |
| Paraquat | 10 | daGal4 | 3.1 | ♂ | 2 | 41 | 19.1 | 35 | 40 | 17.5 | 30 | 39 | 14.6 | 22 | -16.5 | -26.7 | **0.0019** |
| Paraquat | 10 | daGal4 | 3.1 | ♂ | 3 | 42 | 16.6 | 30 | 45 | 13.3 | 18 | 40 | 17.3 | 30 | 4.5 | 0.0 | 0.5164 |
| Paraquat | 10 | daGal4 | 3.1 | ♂ | 4 | 42 | 15.1 | 26 | 45 | 14.4 | 18 | 43 | 15.2 | 22 | 0.4 | 0.0 | 0.2269 |
| Paraquat | 10 | daGal4 | 4.4 | ♀ | 1 | 40 | 26.2 | 43 | 35 | 15.4 | 22 | 40 | 17.2 | 31 | 0.0 | 0.0 | |
| Paraquat | 10 | daGal4 | 4.4 | ♀ | 2 | 40 | 19.8 | 27 | 39 | 14.3 | 21 | 69 | 16.0 | 27 | 0.0 | 0.0 | |
| Paraquat | 10 | daGal4 | 4.4 | ♀ | 3 | 40 | 17.0 | 26 | 40 | 14.7 | 22 | 39 | 13.2 | 26 | -10.3 | 0.0 | **0.0009** |
| Paraquat | 10 | daGal4 | 4.4 | ♀ | 4 | 40 | 19.8 | 40 | 40 | 14.9 | 21 | 40 | 16.5 | 40 | 0.0 | 0.0 | |
| Paraquat | 10 | daGal4 | 4.4 | ♀ | 5 | 41 | 23.9 | 30 | 40 | 19.1 | 30 | 43 | 17.7 | 30 | -7.2 | 0.0 | 0.2879 |
| Paraquat | 10 | daGal4 | 4.4 | ♀ | 6 | 34 | 20.5 | 35 | 45 | 17.3 | 26 | 42 | 15.3 | 22 | -11.3 | -15.4 | **0.0005** |
| Paraquat | 10 | daGal4 | 4.4 | ♂ | 1 | 39 | 29.7 | 43 | 40 | 18.2 | 31 | 40 | 30.8 | 56 | 3.5 | 30.2 | 0.6732 |
| Paraquat | 10 | daGal4 | 4.4 | ♂ | 2 | 40 | 24.8 | 40 | 41 | 11.7 | 15 | 56 | 17.0 | 40 | 0.0 | 0.0 | |
| Paraquat | 10 | daGal4 | 4.4 | ♂ | 3 | 41 | 17.9 | 26 | 40 | 17.5 | 30 | 41 | 28.7 | 70 | 60.0 | 133.3 | **<0.0001** |
| Paraquat | 10 | daGal4 | 4.4 | ♂ | 4 | 40 | 18.4 | 27 | 39 | 13.4 | 21 | 40 | 14.7 | 21 | 0.0 | 0.0 | |
| Paraquat | 10 | daGal4 | 4.4 | ♂ | 5 | 40 | 17.0 | 26 | 45 | 13.3 | 18 | 40 | 22.9 | 42 | 34.4 | 61.5 | **0.0005** |
| Paraquat | 10 | daGal4 | 4.4 | ♂ | 6 | 39 | 15.2 | 22 | 45 | 14.4 | 18 | 41 | 15.9 | 35 | 4.1 | 59.1 | 0.7465 |
| Paraquat | 10 | DJ694 | 2.1 | ♀ | 1 | 40 | 25.3 | 52 | 40 | 17.0 | 27 | 40 | 14.6 | 22 | -14.0 | -18.5 | **0.0141** |
| Paraquat | 10 | DJ694 | 2.1 | ♀ | 2 | 40 | 17.9 | 26 | 39 | 20.9 | 38 | 43 | 17.8 | 26 | -0.5 | 0.0 | **0.0195** |
| Paraquat | 10 | DJ694 | 2.1 | ♀ | 3 | 40 | 12.1 | 16 | 40 | 13.0 | 16 | 40 | 12.0 | 16 | -0.8 | 0.0 | **0.0411** |
| Paraquat | 10 | DJ694 | 2.1 | ♀ | 4 | 36 | 18.6 | 26 | 34 | 16.9 | 35 | 32 | 13.3 | 22 | -21.7 | -15.4 | **0.0119** |
| Paraquat | 10 | DJ694 | 2.1 | ♀ | 5 | 40 | 14.2 | 20 | 40 | 13.6 | 24 | 40 | 10.8 | 12 | -20.6 | -40.0 | **<0.0001** |

### Table S2

| Stress | Age | Gal4 | UAS | Sex | Rep[1] | UAS | | | Gal4 | | | Gal4 + UAS | | | % Extension | | *P* |
|---|---|---|---|---|---|---|---|---|---|---|---|---|---|---|---|---|---|
| | | | | | | n | Mean | Max[2] | n | Mean | Max[2] | n | Mean | Max[2] | Mean | Max[2] | |
| Paraquat | 10 | DJ694 | 2.1 | ♀ | 6 | 41 | 19.3 | 22 | 38 | 16.1 | 26 | 40 | 12.8 | 18 | -20.5 | -18.2 | **0.0002** |
| Paraquat | 10 | DJ694 | 2.1 | ♀ | 7 | 40 | 13.2 | 20 | 40 | 11.4 | 16 | 40 | 10.9 | 12 | -4.4 | -25.0 | 0.3596 |
| Paraquat | 10 | DJ694 | 2.1 | ♂ | 1 | 40 | 24.5 | 47 | 40 | 29.6 | 57 | 40 | 21.5 | 42 | -12.2 | -10.6 | **0.0006** |
| Paraquat | 10 | DJ694 | 2.1 | ♂ | 2 | 41 | 17.6 | 26 | 44 | 14.4 | 18 | 40 | 12.9 | 18 | -10.2 | 0.0 | **0.0131** |
| Paraquat | 10 | DJ694 | 2.1 | ♂ | 3 | 40 | 9.5 | 12 | 40 | 10.6 | 16 | 40 | 10.8 | 12 | 1.9 | 0.0 | **0.0038** |
| Paraquat | 10 | DJ694 | 2.1 | ♂ | 4 | 40 | 15.4 | 22 | 38 | 12.4 | 18 | 27 | 10.9 | 14 | -12.5 | -22.2 | 0.0792 |
| Paraquat | 10 | DJ694 | 2.1 | ♂ | 5 | 40 | 10.5 | 12 | 40 | 12.4 | 16 | 40 | 10.5 | 12 | 0.0 | 0.0 | |
| Paraquat | 10 | DJ694 | 2.1 | ♂ | 6 | 40 | 15.9 | 30 | 42 | 13.3 | 18 | 40 | 13.3 | 18 | 0.0 | 0.0 | |
| Paraquat | 10 | DJ694 | 2.1 | ♂ | 7 | 40 | 10.2 | 12 | 40 | 9.5 | 12 | 40 | 10.1 | 12 | 0.0 | 0.0 | |
| Paraquat | 10 | DJ694 | 4.3 | ♀ | 1 | 40 | 20.8 | 31 | 40 | 16.8 | 31 | 40 | 15.9 | 31 | -5.2 | 0.0 | **0.0018** |
| Paraquat | 10 | DJ694 | 4.3 | ♀ | 2 | 40 | 18.2 | 33 | 40 | 12.4 | 16 | 40 | 13.8 | 16 | 0.0 | 0.0 | |
| Paraquat | 10 | DJ694 | 4.3 | ♀ | 3 | 40 | 18.5 | 26 | 39 | 20.9 | 38 | 41 | 17.1 | 30 | -7.4 | 0.0 | **0.0024** |
| Paraquat | 10 | DJ694 | 4.3 | ♀ | 4 | 40 | 12.2 | 16 | 40 | 13.0 | 16 | 40 | 10.3 | 12 | -16.0 | -25.0 | **<0.0001** |
| Paraquat | 10 | DJ694 | 4.3 | ♀ | 5 | 40 | 15.8 | 25 | 40 | 13.2 | 16 | 40 | 15.2 | 20 | 0.0 | 0.0 | |
| Paraquat | 10 | DJ694 | 4.3 | ♀ | 6 | 42 | 24.3 | 42 | 34 | 16.9 | 35 | 28 | 13.9 | 26 | -18.1 | -25.7 | **0.0368** |
| Paraquat | 10 | DJ694 | 4.3 | ♀ | 7 | 40 | 12.8 | 16 | 40 | 13.6 | 24 | 30 | 11.3 | 12 | -11.5 | -25.0 | **0.0011** |
| Paraquat | 10 | DJ694 | 4.3 | ♀ | 8 | 42 | 19.5 | 26 | 38 | 16.1 | 26 | 40 | 13.8 | 18 | -14.2 | -30.8 | **0.0158** |
| Paraquat | 10 | DJ694 | 4.3 | ♀ | 9 | 40 | 13.5 | 20 | 40 | 11.4 | 16 | 40 | 10.9 | 12 | -4.4 | -25.0 | 0.3375 |
| Paraquat | 10 | DJ694 | 4.3 | ♂ | 1 | 40 | 20.5 | 31 | 40 | 19.5 | 43 | 41 | 24.9 | 43 | 21.3 | 0.0 | **0.0103** |
| Paraquat | 10 | DJ694 | 4.3 | ♂ | 2 | 40 | 13.4 | 20 | 40 | 15.1 | 20 | 40 | 13.9 | 20 | 0.0 | 0.0 | |
| Paraquat | 10 | DJ694 | 4.3 | ♂ | 3 | 58 | 18.7 | 26 | 44 | 14.4 | 18 | 42 | 13.9 | 18 | -3.2 | 0.0 | 0.5680 |
| Paraquat | 10 | DJ694 | 4.3 | ♂ | 4 | 40 | 11.4 | 16 | 40 | 10.6 | 16 | 50 | 11.0 | 16 | 0.0 | 0.0 | |
| Paraquat | 10 | DJ694 | 4.3 | ♂ | 5 | 40 | 14.0 | 20 | 40 | 14.2 | 20 | 40 | 12.9 | 20 | -8.1 | 0.0 | 0.2208 |
| Paraquat | 10 | DJ694 | 4.3 | ♂ | 6 | 42 | 16.0 | 35 | 38 | 12.4 | 18 | 33 | 15.3 | 38 | 0.0 | 8.6 | |
| Paraquat | 10 | DJ694 | 4.3 | ♂ | 7 | 40 | 11.1 | 12 | 40 | 12.4 | 16 | 40 | 10.7 | 12 | -3.6 | 0.0 | **0.0021** |
| Paraquat | 10 | DJ694 | 4.3 | ♂ | 8 | 40 | 13.6 | 22 | 42 | 13.3 | 18 | 36 | 15.8 | 26 | 16.0 | 18.2 | **0.0066** |
| Paraquat | 10 | DJ694 | 4.3 | ♂ | 9 | 40 | 11.3 | 16 | 40 | 9.5 | 12 | 40 | 11.0 | 12 | 0.0 | 0.0 | |
| Paraquat | 10 | DJ694 | 9.1 | ♀ | 1 | 40 | 19.0 | 27 | 39 | 15.9 | 21 | 40 | 15.4 | 27 | -3.4 | 0.0 | **0.0008** |
| Paraquat | 10 | DJ694 | 9.1 | ♀ | 2 | 41 | 18.7 | 30 | 39 | 20.9 | 38 | 40 | 14.8 | 22 | -20.8 | -26.7 | **0.0015** |
| Paraquat | 10 | DJ694 | 9.1 | ♀ | 3 | 40 | 13.0 | 20 | 40 | 13.0 | 16 | 40 | 11.5 | 12 | -11.5 | -25.0 | **0.0009** |
| Paraquat | 10 | DJ694 | 9.1 | ♀ | 4 | 41 | 18.2 | 27 | 40 | 14.4 | 21 | 38 | 18.1 | 27 | 0.0 | 0.0 | |
| Paraquat | 10 | DJ694 | 9.1 | ♀ | 5 | 42 | 22.8 | 35 | 34 | 16.9 | 35 | 37 | 12.1 | 18 | -28.7 | -48.6 | **0.0003** |
| Paraquat | 10 | DJ694 | 9.1 | ♀ | 6 | 40 | 15.1 | 20 | 40 | 13.6 | 24 | 40 | 10.8 | 12 | -20.6 | -40.0 | **<0.0001** |
| Paraquat | 10 | DJ694 | 9.1 | ♀ | 7 | 42 | 21.0 | 30 | 38 | 16.1 | 26 | 39 | 11.2 | 14 | -30.3 | -46.2 | **<0.0001** |
| Paraquat | 10 | DJ694 | 9.1 | ♀ | 8 | 40 | 11.7 | 16 | 40 | 11.4 | 16 | 40 | 10.8 | 12 | -5.3 | -25.0 | 0.0642 |
| Paraquat | 10 | DJ694 | 9.1 | ♂ | 1 | 40 | 15.6 | 21 | 40 | 15.0 | 21 | 39 | 13.2 | 15 | -12.0 | -28.6 | **0.0011** |
| Paraquat | 10 | DJ694 | 9.1 | ♂ | 2 | 38 | 16.3 | 30 | 44 | 14.4 | 18 | 40 | 14.3 | 26 | -0.4 | 0.0 | 0.0911 |
| Paraquat | 10 | DJ694 | 9.1 | ♂ | 3 | 40 | 11.0 | 16 | 40 | 10.6 | 16 | 40 | 9.4 | 12 | -11.3 | -25.0 | **0.0011** |
| Paraquat | 10 | DJ694 | 9.1 | ♂ | 4 | 40 | 17.7 | 27 | 39 | 14.7 | 21 | 40 | 11.8 | 15 | -20.2 | -28.6 | **<0.0001** |
| Paraquat | 10 | DJ694 | 9.1 | ♂ | 5 | 40 | 14.1 | 22 | 38 | 12.4 | 18 | 37 | 12.8 | 18 | 0.0 | 0.0 | |
| Paraquat | 10 | DJ694 | 9.1 | ♂ | 6 | 40 | 10.8 | 12 | 40 | 12.4 | 16 | 40 | 11.0 | 12 | 0.0 | 0.0 | |
| Paraquat | 10 | DJ694 | 9.1 | ♂ | 7 | 43 | 13.1 | 22 | 42 | 13.3 | 18 | 41 | 12.8 | 18 | -1.8 | 0.0 | 0.4324 |
| Paraquat | 10 | DJ694 | 9.1 | ♂ | 8 | 40 | 11.4 | 16 | 40 | 9.5 | 12 | 40 | 11.7 | 16 | 2.6 | 0.0 | 0.5051 |
| Paraquat | 10 | DJ694 | 3.2 | ♀ | 1 | 38 | 19.1 | 40 | 35 | 19.7 | 35 | 40 | 19.6 | 40 | 0.0 | 0.0 | |
| Paraquat | 10 | DJ694 | 3.2 | ♀ | 2 | 40 | 19.2 | 33 | 40 | 12.4 | 16 | 40 | 15.8 | 20 | 0.0 | 0.0 | |
| Paraquat | 10 | DJ694 | 3.2 | ♀ | 3 | 35 | 17.1 | 30 | 39 | 20.9 | 38 | 43 | 19.7 | 42 | 0.0 | 10.5 | |
| Paraquat | 10 | DJ694 | 3.2 | ♀ | 4 | 40 | 14.0 | 24 | 40 | 13.0 | 16 | 40 | 12.8 | 34 | -1.5 | 41.7 | 0.3058 |
| Paraquat | 10 | DJ694 | 3.2 | ♀ | 5 | 40 | 17.3 | 25 | 40 | 13.2 | 16 | 40 | 18.0 | 33 | 4.1 | 32.0 | 0.5341 |
| Paraquat | 10 | DJ694 | 3.2 | ♀ | 6 | 40 | 22.2 | 42 | 34 | 16.9 | 35 | 38 | 20.6 | 61 | 0.0 | 45.2 | |

### Table S2



| Stress | Age | Gal4 | UAS | Sex | Rep[1] | UAS | | | Gal4 | | | Gal4 + UAS | | | % Extension | | *P* |
|---|---|---|---|---|---|---|---|---|---|---|---|---|---|---|---|---|---|
| | | | | | | n | Mean | Max[2] | n | Mean | Max[2] | n | Mean | Max[2] | Mean | Max[2] | |
| Paraquat | 10 | DJ694 | 3.2 | ♀ | 7 | 30 | 14.7 | 24 | 40 | 13.6 | 24 | 40 | 12.3 | 20 | -9.6 | -16.7 | **0.0287** |
| Paraquat | 10 | DJ694 | 3.2 | ♀ | 8 | 39 | 19.9 | 35 | 38 | 16.1 | 26 | 34 | 23.6 | 42 | 18.5 | 20.0 | 0.0589 |
| Paraquat | 10 | DJ694 | 3.2 | ♀ | 9 | 40 | 15.6 | 34 | 40 | 11.4 | 16 | 40 | 13.0 | 20 | 0.0 | 0.0 | |
| Paraquat | 10 | DJ694 | 3.2 | ♂ | 1 | 38 | 24.3 | 45 | 39 | 24.0 | 63 | 40 | 47.4 | 75 | 94.6 | 19.0 | **<0.0001** |
| Paraquat | 10 | DJ694 | 3.2 | ♂ | 2 | 40 | 18.3 | 25 | 40 | 15.1 | 20 | 40 | 25.1 | 48 | 37.0 | 92.0 | **0.0016** |
| Paraquat | 10 | DJ694 | 3.2 | ♂ | 3 | 38 | 25.0 | 42 | 44 | 14.4 | 18 | 41 | 29.2 | 61 | 17.0 | 45.2 | 0.4007 |
| Paraquat | 10 | DJ694 | 3.2 | ♂ | 4 | 40 | 14.0 | 20 | 40 | 10.6 | 16 | 40 | 16.1 | 39 | 15.2 | 95.0 | **0.0009** |
| Paraquat | 10 | DJ694 | 3.2 | ♂ | 5 | 40 | 18.4 | 33 | 40 | 14.2 | 20 | 40 | 24.9 | 58 | 35.1 | 75.8 | **0.0019** |
| Paraquat | 10 | DJ694 | 3.2 | ♂ | 6 | 38 | 24.3 | 61 | 38 | 12.4 | 18 | 40 | 34.8 | 61 | 43.1 | 0.0 | **<0.0001** |
| Paraquat | 10 | DJ694 | 3.2 | ♂ | 7 | 40 | 14.9 | 20 | 40 | 12.4 | 16 | 40 | 17.3 | 63 | 16.1 | 215.0 | **0.0057** |
| Paraquat | 10 | DJ694 | 3.2 | ♂ | 8 | 39 | 29.8 | 61 | 42 | 13.3 | 18 | 44 | 27.1 | 70 | 0.0 | 14.8 | |
| Paraquat | 10 | DJ694 | 3.2 | ♂ | 9 | 40 | 15.2 | 20 | 40 | 9.5 | 12 | 40 | 14.1 | 24 | 0.0 | 20.0 | |
| Paraquat | 10 | DJ694 | 10.1 | ♀ | 1 | 29 | 17.3 | 22 | 24 | 17.0 | 17 | 39 | 17.4 | 22 | 0.2 | 0.0 | 0.1672 |
| Paraquat | 10 | DJ694 | 10.1 | ♀ | 2 | 40 | 18.9 | 27 | 39 | 15.9 | 21 | 40 | 16.0 | 27 | 0.0 | 0.0 | |
| Paraquat | 10 | DJ694 | 10.1 | ♀ | 3 | 36 | 22.9 | 38 | 39 | 20.9 | 38 | 40 | 16.7 | 35 | -20.5 | -7.9 | **0.0011** |
| Paraquat | 10 | DJ694 | 10.1 | ♀ | 4 | 40 | 15.4 | 24 | 40 | 13.0 | 16 | 40 | 9.9 | 12 | -23.8 | -25.0 | **<0.0001** |
| Paraquat | 10 | DJ694 | 10.1 | ♀ | 5 | 40 | 18.8 | 27 | 40 | 14.4 | 21 | 40 | 18.8 | 40 | 0.4 | 48.1 | **0.0004** |
| Paraquat | 10 | DJ694 | 10.1 | ♀ | 6 | 36 | 27.9 | 61 | 34 | 16.9 | 35 | 31 | 16.6 | 30 | -2.0 | -14.3 | 0.9156 |
| Paraquat | 10 | DJ694 | 10.1 | ♀ | 7 | 40 | 17.6 | 34 | 40 | 13.6 | 24 | 40 | 12.3 | 20 | -9.6 | -16.7 | 0.0900 |
| Paraquat | 10 | DJ694 | 10.1 | ♀ | 8 | 37 | 24.2 | 38 | 38 | 16.1 | 26 | 40 | 23.0 | 38 | 0.0 | 0.0 | |
| Paraquat | 10 | DJ694 | 10.1 | ♀ | 9 | 40 | 15.3 | 20 | 40 | 11.4 | 16 | 40 | 11.7 | 20 | 0.0 | 0.0 | |
| Paraquat | 10 | DJ694 | 10.1 | ♂ | 1 | 37 | 18.8 | 28 | 29 | 24.5 | 48 | 35 | 42.3 | 73 | 72.6 | 52.1 | **0.0002** |
| Paraquat | 10 | DJ694 | 10.1 | ♂ | 2 | 40 | 25.9 | 40 | 40 | 15.0 | 21 | 40 | 22.9 | 40 | 0.0 | 0.0 | |
| Paraquat | 10 | DJ694 | 10.1 | ♂ | 3 | 38 | 25.9 | 61 | 44 | 14.4 | 18 | 43 | 28.8 | 70 | 11.1 | 14.8 | 0.3863 |
| Paraquat | 10 | DJ694 | 10.1 | ♂ | 4 | 40 | 13.3 | 20 | 40 | 10.6 | 16 | 40 | 12.7 | 34 | 0.0 | 70.0 | |
| Paraquat | 10 | DJ694 | 10.1 | ♂ | 5 | 40 | 20.6 | 40 | 39 | 14.7 | 21 | 39 | 15.9 | 21 | 0.0 | 0.0 | |
| Paraquat | 10 | DJ694 | 10.1 | ♂ | 6 | 37 | 25.4 | 42 | 38 | 12.4 | 18 | 32 | 19.6 | 42 | 0.0 | 0.0 | |
| Paraquat | 10 | DJ694 | 10.1 | ♂ | 7 | 40 | 17.2 | 46 | 40 | 12.4 | 16 | 40 | 14.6 | 39 | 0.0 | 0.0 | |
| Paraquat | 10 | DJ694 | 10.1 | ♂ | 8 | 39 | 24.7 | 42 | 42 | 13.3 | 18 | 42 | 34.0 | 70 | 37.9 | 66.7 | **0.0053** |
| Paraquat | 10 | DJ694 | 10.1 | ♂ | 9 | 40 | 14.9 | 24 | 40 | 9.5 | 12 | 38 | 24.6 | 63 | 65.5 | 162.5 | **<0.0001** |
| Paraquat | 10 | DJ694 | 3.1 | ♀ | 1 | 40 | 12.5 | 22 | 40 | 15.9 | 26 | 40 | 14.4 | 22 | 0.0 | 0.0 | |
| Paraquat | 10 | DJ694 | 3.1 | ♀ | 2 | 43 | 14.0 | 22 | 39 | 20.9 | 38 | 38 | 13.3 | 26 | -5.3 | 0.0 | 0.2802 |
| Paraquat | 10 | DJ694 | 3.1 | ♀ | 3 | 40 | 13.8 | 20 | 40 | 13.0 | 16 | 40 | 9.9 | 12 | -23.8 | -25.0 | **<0.0001** |
| Paraquat | 10 | DJ694 | 3.1 | ♀ | 4 | 40 | 17.9 | 30 | 34 | 16.9 | 35 | 35 | 12.6 | 18 | -25.3 | -40.0 | **0.0036** |
| Paraquat | 10 | DJ694 | 3.1 | ♀ | 5 | 44 | 18.2 | 35 | 38 | 16.1 | 26 | 47 | 16.1 | 26 | 0.0 | 0.0 | |
| Paraquat | 10 | DJ694 | 3.1 | ♂ | 1 | 33 | 23.5 | 34 | 38 | 30.0 | 61 | 39 | 42.1 | 79 | 40.2 | 29.5 | **0.0110** |
| Paraquat | 10 | DJ694 | 3.1 | ♂ | 2 | 41 | 19.1 | 35 | 44 | 14.4 | 18 | 40 | 18.1 | 35 | 0.0 | 0.0 | |
| Paraquat | 10 | DJ694 | 3.1 | ♂ | 3 | 40 | 12.4 | 20 | 40 | 10.6 | 16 | 40 | 12.4 | 16 | 0.0 | 0.0 | |
| Paraquat | 10 | DJ694 | 3.1 | ♂ | 4 | 42 | 16.6 | 30 | 38 | 12.4 | 18 | 35 | 17.7 | 38 | 6.9 | 26.7 | **0.0026** |
| Paraquat | 10 | DJ694 | 3.1 | ♂ | 5 | 42 | 15.1 | 26 | 42 | 13.3 | 18 | 33 | 17.5 | 35 | 15.5 | 34.6 | **0.0002** |
| Paraquat | 10 | DJ694 | 4.4 | ♀ | 1 | 40 | 22.0 | 45 | 40 | 22.4 | 40 | 40 | 22.5 | 35 | 0.6 | -12.5 | 0.5867 |
| Paraquat | 10 | DJ694 | 4.4 | ♀ | 2 | 40 | 26.2 | 43 | 40 | 16.8 | 31 | 40 | 17.6 | 31 | 0.0 | 0.0 | |
| Paraquat | 10 | DJ694 | 4.4 | ♀ | 3 | 40 | 19.8 | 27 | 39 | 15.9 | 21 | 40 | 15.5 | 21 | -2.7 | 0.0 | **0.0001** |
| Paraquat | 10 | DJ694 | 4.4 | ♀ | 4 | 40 | 17.0 | 26 | 39 | 20.9 | 38 | 41 | 16.0 | 30 | -6.0 | 0.0 | **0.0002** |
| Paraquat | 10 | DJ694 | 4.4 | ♀ | 5 | 40 | 13.0 | 20 | 40 | 13.0 | 16 | 40 | 12.7 | 20 | -2.3 | 0.0 | 0.5633 |
| Paraquat | 10 | DJ694 | 4.4 | ♀ | 6 | 40 | 19.8 | 40 | 40 | 14.4 | 21 | 40 | 15.9 | 21 | 0.0 | 0.0 | |
| Paraquat | 10 | DJ694 | 4.4 | ♀ | 7 | 41 | 23.9 | 30 | 34 | 16.9 | 35 | 43 | 14.7 | 22 | -13.4 | -26.7 | 0.0999 |
| Paraquat | 10 | DJ694 | 4.4 | ♀ | 8 | 40 | 16.0 | 20 | 40 | 13.6 | 24 | 40 | 11.7 | 16 | -14.0 | -20.0 | **0.0036** |
| Paraquat | 10 | DJ694 | 4.4 | ♀ | 9 | 34 | 20.5 | 35 | 38 | 16.1 | 26 | 42 | 12.7 | 18 | -21.2 | -30.8 | **0.0009** |

**Table S2**

| Stress | Age | Gal4 | UAS | Sex | Rep[1] | UAS | | | Gal4 | | | Gal4 + UAS | | | % Extension | | *P* |
|---|---|---|---|---|---|---|---|---|---|---|---|---|---|---|---|---|---|
| | | | | | | n | Mean | Max[2] | n | Mean | Max[2] | n | Mean | Max[2] | Mean | Max[2] | |
| Paraquat | 10 | DJ694 | 4.4 | ♀ | 10 | 40 | 13.0 | 20 | 40 | 11.4 | 16 | 40 | 9.7 | 12 | -14.9 | -25.0 | **0.0013** |
| Paraquat | 10 | DJ694 | 4.4 | ♂ | 1 | 40 | 25.9 | 55 | 40 | 19.5 | 30 | 45 | 23.8 | 35 | 0.0 | 0.0 | |
| Paraquat | 10 | DJ694 | 4.4 | ♂ | 2 | 39 | 29.7 | 43 | 40 | 19.5 | 43 | 39 | 31.9 | 69 | 7.3 | 60.5 | 0.5747 |
| Paraquat | 10 | DJ694 | 4.4 | ♂ | 3 | 40 | 24.8 | 40 | 40 | 15.0 | 21 | 40 | 20.5 | 40 | 0.0 | 0.0 | |
| Paraquat | 10 | DJ694 | 4.4 | ♂ | 4 | 41 | 17.9 | 26 | 44 | 14.4 | 18 | 40 | 16.8 | 30 | 0.0 | 15.4 | |
| Paraquat | 10 | DJ694 | 4.4 | ♂ | 5 | 40 | 13.2 | 20 | 40 | 10.6 | 16 | 40 | 11.9 | 16 | 0.0 | 0.0 | |
| Paraquat | 10 | DJ694 | 4.4 | ♂ | 6 | 40 | 18.4 | 27 | 39 | 14.7 | 21 | 40 | 20.6 | 40 | 12.4 | 48.1 | 0.1945 |
| Paraquat | 10 | DJ694 | 4.4 | ♂ | 7 | 40 | 17.0 | 26 | 38 | 12.4 | 18 | 40 | 17.1 | 30 | 0.6 | 15.4 | **0.0001** |
| Paraquat | 10 | DJ694 | 4.4 | ♂ | 8 | 39 | 14.1 | 24 | 40 | 12.4 | 16 | 40 | 11.6 | 16 | -6.5 | 0.0 | **0.0001** |
| Paraquat | 10 | DJ694 | 4.4 | ♂ | 9 | 39 | 15.2 | 22 | 42 | 13.3 | 18 | 37 | 14.8 | 22 | 0.0 | 0.0 | |
| Paraquat | 10 | DJ694 | 4.4 | ♂ | 10 | 40 | 12.4 | 20 | 40 | 9.5 | 12 | 40 | 12.3 | 20 | 0.0 | 0.0 | |
| Paraquat | 10 | D42 | 2.1 | ♀ | 1 | 40 | 25.3 | 52 | 40 | 24.3 | 47 | 40 | 20.1 | 42 | -17.0 | -10.6 | **0.0236** |
| Paraquat | 10 | D42 | 2.1 | ♀ | 2 | 40 | 17.9 | 26 | 40 | 24.5 | 61 | 40 | 14.6 | 26 | -18.4 | 0.0 | **0.0014** |
| Paraquat | 10 | D42 | 2.1 | ♀ | 3 | 36 | 18.6 | 26 | 41 | 36.7 | 61 | 40 | 22.4 | 30 | 0.0 | 0.0 | |
| Paraquat | 10 | D42 | 2.1 | ♀ | 4 | 41 | 19.3 | 22 | 58 | 28.5 | 61 | 44 | 19.9 | 42 | 0.0 | 0.0 | |
| Paraquat | 10 | D42 | 2.1 | ♂ | 1 | 40 | 24.5 | 47 | 40 | 25.9 | 52 | 40 | 21.3 | 37 | -13.3 | -21.3 | **0.0211** |
| Paraquat | 10 | D42 | 2.1 | ♂ | 2 | 41 | 17.6 | 26 | 41 | 23.5 | 38 | 38 | 21.8 | 30 | 0.0 | 0.0 | |
| Paraquat | 10 | D42 | 2.1 | ♂ | 3 | 40 | 15.4 | 22 | 40 | 19.9 | 26 | 40 | 17.4 | 26 | 0.0 | 0.0 | |
| Paraquat | 10 | D42 | 2.1 | ♂ | 4 | 40 | 15.9 | 30 | 44 | 20.7 | 38 | 41 | 16.8 | 30 | 0.0 | 0.0 | |
| Paraquat | 10 | D42 | 4.3 | ♀ | 1 | 40 | 20.8 | 31 | 39 | 22.9 | 43 | 40 | 19.5 | 31 | -6.1 | 0.0 | **0.0494** |
| Paraquat | 10 | D42 | 4.3 | ♀ | 2 | 40 | 18.2 | 33 | 40 | 14.5 | 25 | 40 | 14.5 | 16 | -0.2 | -36.0 | **0.0006** |
| Paraquat | 10 | D42 | 4.3 | ♀ | 3 | 40 | 18.5 | 26 | 40 | 24.5 | 61 | 42 | 15.5 | 26 | -16.5 | 0.0 | **0.0234** |
| Paraquat | 10 | D42 | 4.3 | ♀ | 4 | 40 | 15.8 | 25 | 40 | 14.7 | 20 | 40 | 14.7 | 20 | 0.0 | 0.0 | |
| Paraquat | 10 | D42 | 4.3 | ♀ | 5 | 42 | 24.3 | 42 | 41 | 36.7 | 61 | 44 | 24.0 | 38 | -1.2 | -9.5 | 0.8692 |
| Paraquat | 10 | D42 | 4.3 | ♀ | 6 | 42 | 19.5 | 26 | 58 | 28.5 | 61 | 39 | 16.5 | 26 | -15.6 | 0.0 | **0.0036** |
| Paraquat | 10 | D42 | 4.3 | ♂ | 1 | 40 | 20.5 | 31 | 40 | 26.8 | 43 | 40 | 17.2 | 22 | -16.1 | -29.0 | **0.0032** |
| Paraquat | 10 | D42 | 4.3 | ♂ | 2 | 40 | 13.4 | 20 | 40 | 13.2 | 20 | 40 | 10.7 | 16 | -18.6 | -20.0 | **0.0013** |
| Paraquat | 10 | D42 | 4.3 | ♂ | 3 | 58 | 18.7 | 26 | 41 | 23.5 | 38 | 40 | 20.9 | 35 | 0.0 | 0.0 | |
| Paraquat | 10 | D42 | 4.3 | ♂ | 4 | 40 | 14.0 | 20 | 40 | 14.4 | 72 | 40 | 10.5 | 20 | -25.0 | 0.0 | **0.0002** |
| Paraquat | 10 | D42 | 4.3 | ♂ | 5 | 42 | 16.0 | 35 | 40 | 19.9 | 26 | 42 | 22.2 | 61 | 11.5 | 74.3 | 0.2516 |
| Paraquat | 10 | D42 | 4.3 | ♂ | 6 | 40 | 13.6 | 22 | 44 | 20.7 | 38 | 42 | 18.6 | 30 | 0.0 | 0.0 | |
| Paraquat | 10 | D42 | 9.1 | ♀ | 1 | 40 | 19.0 | 27 | 40 | 21.8 | 27 | 40 | 17.5 | 27 | -7.7 | 0.0 | **0.0002** |
| Paraquat | 10 | D42 | 9.1 | ♀ | 2 | 41 | 18.7 | 30 | 40 | 24.5 | 61 | 41 | 12.7 | 18 | -31.9 | -40.0 | **<0.0001** |
| Paraquat | 10 | D42 | 9.1 | ♀ | 3 | 41 | 18.2 | 27 | 40 | 20.2 | 40 | 39 | 21.7 | 40 | 7.4 | 0.0 | **0.0261** |
| Paraquat | 10 | D42 | 9.1 | ♀ | 4 | 42 | 22.8 | 35 | 41 | 36.7 | 61 | 38 | 21.4 | 35 | -6.2 | 0.0 | 0.3870 |
| Paraquat | 10 | D42 | 9.1 | ♀ | 5 | 42 | 21.0 | 30 | 58 | 28.5 | 61 | 45 | 18.8 | 35 | -10.2 | 0.0 | 0.0604 |
| Paraquat | 10 | D42 | 9.1 | ♂ | 1 | 40 | 15.6 | 21 | 40 | 19.0 | 40 | 39 | 14.3 | 21 | -8.4 | 0.0 | 0.0759 |
| Paraquat | 10 | D42 | 9.1 | ♂ | 2 | 38 | 16.3 | 30 | 41 | 23.5 | 38 | 39 | 15.0 | 22 | -8.1 | -26.7 | 0.2309 |
| Paraquat | 10 | D42 | 9.1 | ♂ | 3 | 40 | 17.7 | 27 | 40 | 18.3 | 40 | 40 | 14.7 | 21 | -17.0 | -22.2 | **0.0016** |
| Paraquat | 10 | D42 | 9.1 | ♂ | 4 | 40 | 14.1 | 22 | 40 | 19.9 | 26 | 39 | 18.0 | 30 | 0.0 | 15.4 | |
| Paraquat | 10 | D42 | 9.1 | ♂ | 5 | 43 | 13.1 | 22 | 44 | 20.7 | 38 | 41 | 15.7 | 22 | 0.0 | 0.0 | |
| Paraquat | 10 | D42 | 3.2 | ♀ | 1 | 38 | 19.1 | 40 | 38 | 20.3 | 40 | 40 | 18.5 | 40 | -2.9 | 0.0 | 0.3700 |
| Paraquat | 10 | D42 | 3.2 | ♀ | 2 | 40 | 19.2 | 33 | 40 | 14.5 | 25 | 40 | 17.1 | 33 | 0.0 | 0.0 | |
| Paraquat | 10 | D42 | 3.2 | ♀ | 3 | 35 | 17.1 | 30 | 40 | 24.5 | 61 | 41 | 17.5 | 35 | 0.0 | 0.0 | |
| Paraquat | 10 | D42 | 3.2 | ♀ | 4 | 40 | 17.3 | 25 | 40 | 14.7 | 20 | 40 | 18.9 | 33 | 9.6 | 32.0 | 0.1735 |
| Paraquat | 10 | D42 | 3.2 | ♀ | 5 | 40 | 22.2 | 42 | 41 | 36.7 | 61 | 41 | 18.0 | 30 | -18.9 | -28.6 | **0.0357** |
| Paraquat | 10 | D42 | 3.2 | ♀ | 6 | 39 | 19.9 | 35 | 58 | 28.5 | 61 | 36 | 19.3 | 30 | -2.8 | -14.3 | 0.8258 |
| Paraquat | 10 | D42 | 3.2 | ♂ | 1 | 38 | 24.3 | 45 | 37 | 24.5 | 45 | 40 | 42.2 | 69 | 72.3 | 53.3 | **<0.0001** |
| Paraquat | 10 | D42 | 3.2 | ♂ | 2 | 40 | 18.3 | 25 | 40 | 13.2 | 20 | 40 | 22.9 | 58 | 25.1 | 132.0 | **0.0309** |

### Table S2



| Stress | Age | Gal4 | UAS | Sex | Rep[1] | UAS | | | Gal4 | | | Gal4 + UAS | | | % Extension | | *P* |
|---|---|---|---|---|---|---|---|---|---|---|---|---|---|---|---|---|---|
| | | | | | | n | Mean | Max[2] | n | Mean | Max[2] | n | Mean | Max[2] | Mean | Max[2] | |
| Paraquat | 10 | D42 | 3.2 | ♂ | 3 | 38 | 25.0 | 42 | 41 | 23.5 | 38 | 38 | 32.0 | 61 | 28.0 | 45.2 | **0.0035** |
| Paraquat | 10 | D42 | 3.2 | ♂ | 4 | 40 | 18.4 | 33 | 40 | 14.4 | 72 | 40 | 21.8 | 38 | 18.2 | 0.0 | **0.0002** |
| Paraquat | 10 | D42 | 3.2 | ♂ | 5 | 38 | 24.3 | 61 | 40 | 19.9 | 26 | 40 | 32.3 | 82 | 32.9 | 34.4 | **0.0038** |
| Paraquat | 10 | D42 | 3.2 | ♂ | 6 | 39 | 29.8 | 61 | 44 | 20.7 | 38 | 45 | 33.6 | 70 | 12.6 | 14.8 | 0.1980 |
| Paraquat | 10 | D42 | 10.1 | ♀ | 1 | 29 | 17.3 | 22 | 31 | 17.3 | 22 | 40 | 20.5 | 42 | 18.2 | 90.9 | **0.0133** |
| Paraquat | 10 | D42 | 10.1 | ♀ | 2 | 40 | 18.9 | 27 | 40 | 21.8 | 27 | 39 | 19.8 | 27 | 0.0 | 0.0 | |
| Paraquat | 10 | D42 | 10.1 | ♀ | 3 | 36 | 22.9 | 38 | 40 | 24.5 | 61 | 41 | 18.8 | 30 | -18.1 | -21.1 | **0.0089** |
| Paraquat | 10 | D42 | 10.1 | ♀ | 4 | 40 | 18.8 | 27 | 40 | 20.2 | 40 | 40 | 19.1 | 40 | 0.0 | 0.0 | |
| Paraquat | 10 | D42 | 10.1 | ♀ | 5 | 36 | 27.9 | 61 | 41 | 36.7 | 61 | 41 | 16.9 | 38 | -39.4 | -37.7 | **<0.0001** |
| Paraquat | 10 | D42 | 10.1 | ♀ | 6 | 37 | 24.2 | 38 | 58 | 28.5 | 61 | 36 | 17.0 | 38 | -29.9 | 0.0 | **0.0003** |
| Paraquat | 10 | D42 | 10.1 | ♂ | 1 | 37 | 18.8 | 28 | 36 | 19.9 | 42 | 35 | 55.1 | 100 | 177.4 | 138.1 | **<0.0001** |
| Paraquat | 10 | D42 | 10.1 | ♂ | 2 | 40 | 25.9 | 40 | 40 | 19.0 | 40 | 40 | 20.2 | 27 | 0.0 | -32.5 | |
| Paraquat | 10 | D42 | 10.1 | ♂ | 3 | 38 | 25.9 | 61 | 41 | 23.5 | 38 | 37 | 29.4 | 82 | 13.3 | 34.4 | **0.0414** |
| Paraquat | 10 | D42 | 10.1 | ♂ | 4 | 40 | 20.6 | 40 | 40 | 18.3 | 40 | 40 | 17.5 | 27 | -4.5 | -32.5 | **0.0066** |
| Paraquat | 10 | D42 | 10.1 | ♂ | 5 | 37 | 25.4 | 42 | 40 | 19.9 | 26 | 38 | 26.7 | 42 | 5.2 | 0.0 | 0.5106 |
| Paraquat | 10 | D42 | 10.1 | ♂ | 6 | 39 | 24.7 | 42 | 44 | 20.7 | 38 | 49 | 26.7 | 61 | 8.4 | 45.2 | **0.0115** |
| Paraquat | 10 | D42 | 3.1 | ♀ | 1 | 40 | 12.5 | 22 | 39 | 16.9 | 50 | 40 | 16.5 | 34 | 0.0 | 0.0 | |
| Paraquat | 10 | D42 | 3.1 | ♀ | 2 | 43 | 14.0 | 22 | 40 | 24.5 | 61 | 43 | 13.8 | 22 | -1.3 | 0.0 | 0.7175 |
| Paraquat | 10 | D42 | 3.1 | ♀ | 3 | 40 | 17.9 | 30 | 41 | 36.7 | 61 | 28 | 12.9 | 26 | -28.2 | -13.3 | **0.0011** |
| Paraquat | 10 | D42 | 3.1 | ♀ | 4 | 44 | 18.2 | 35 | 58 | 28.5 | 61 | 39 | 13.0 | 18 | -28.8 | -48.6 | **<0.0001** |
| Paraquat | 10 | D42 | 3.1 | ♂ | 1 | 33 | 23.5 | 34 | 37 | 35.9 | 79 | 38 | 42.3 | 79 | 17.7 | 0.0 | 0.2826 |
| Paraquat | 10 | D42 | 3.1 | ♂ | 2 | 41 | 19.1 | 35 | 41 | 23.5 | 38 | 40 | 22.1 | 35 | 0.0 | 0.0 | |
| Paraquat | 10 | D42 | 3.1 | ♂ | 3 | 42 | 16.6 | 30 | 40 | 19.9 | 26 | 37 | 18.4 | 38 | 0.0 | 26.7 | |
| Paraquat | 10 | D42 | 3.1 | ♂ | 4 | 42 | 15.1 | 26 | 44 | 20.7 | 38 | 39 | 18.5 | 35 | 0.0 | 0.0 | |
| Paraquat | 10 | D42 | 4.4 | ♀ | 1 | 40 | 22.0 | 45 | 40 | 24.8 | 40 | 40 | 21.6 | 30 | -1.7 | -25.0 | **0.0119** |
| Paraquat | 10 | D42 | 4.4 | ♀ | 2 | 40 | 26.2 | 43 | 39 | 22.9 | 43 | 40 | 22.0 | 43 | -3.8 | 0.0 | **0.0465** |
| Paraquat | 10 | D42 | 4.4 | ♀ | 3 | 40 | 19.8 | 27 | 40 | 21.8 | 27 | 40 | 19.0 | 40 | -3.9 | 48.1 | **0.0138** |
| Paraquat | 10 | D42 | 4.4 | ♀ | 4 | 40 | 17.0 | 26 | 40 | 24.5 | 61 | 44 | 14.3 | 26 | -16.0 | 0.0 | **0.0218** |
| Paraquat | 10 | D42 | 4.4 | ♀ | 5 | 40 | 19.8 | 40 | 40 | 20.2 | 40 | 40 | 20.3 | 40 | 0.2 | 0.0 | 0.7853 |
| Paraquat | 10 | D42 | 4.4 | ♀ | 6 | 41 | 23.9 | 30 | 41 | 36.7 | 61 | 28 | 11.1 | 14 | -53.3 | -53.3 | **<0.0001** |
| Paraquat | 10 | D42 | 4.4 | ♀ | 7 | 34 | 20.5 | 35 | 58 | 28.5 | 61 | 41 | 12.8 | 18 | -37.4 | -48.6 | **<0.0001** |
| Paraquat | 10 | D42 | 4.4 | ♂ | 1 | 40 | 25.9 | 55 | 40 | 29.1 | 55 | 40 | 28.4 | 60 | 0.0 | 9.1 | |
| Paraquat | 10 | D42 | 4.4 | ♂ | 2 | 39 | 29.7 | 43 | 40 | 26.8 | 43 | 40 | 22.9 | 31 | -14.6 | -27.9 | **0.0001** |
| Paraquat | 10 | D42 | 4.4 | ♂ | 3 | 40 | 24.8 | 40 | 40 | 19.0 | 40 | 40 | 17.9 | 40 | -5.8 | 0.0 | **0.0007** |
| Paraquat | 10 | D42 | 4.4 | ♂ | 4 | 41 | 17.9 | 26 | 41 | 23.5 | 38 | 40 | 15.6 | 26 | -13.0 | 0.0 | **0.0311** |
| Paraquat | 10 | D42 | 4.4 | ♂ | 5 | 40 | 18.4 | 27 | 40 | 18.3 | 40 | 39 | 16.9 | 27 | -7.5 | 0.0 | 0.1100 |
| Paraquat | 10 | D42 | 4.4 | ♂ | 6 | 40 | 17.0 | 26 | 40 | 19.9 | 26 | 35 | 16.1 | 30 | -5.4 | 15.4 | 0.0763 |
| Paraquat | 10 | D42 | 4.4 | ♂ | 7 | 39 | 15.2 | 22 | 44 | 20.7 | 38 | 43 | 22.3 | 35 | 7.7 | 0.0 | 0.6678 |
| Starvation | 3 | daGal4 | 2.1 | ♀ | 1 | 40 | 54.3 | 73 | 40 | 54.6 | 68 | 40 | 56.9 | 73 | 4.2 | 0.0 | 0.2560 |
| Starvation | 3 | daGal4 | 2.1 | ♀ | 2 | 40 | 28.7 | 40 | 40 | 28.6 | 40 | 40 | 31.2 | 40 | 8.7 | 0.0 | 0.0992 |
| Starvation | 3 | daGal4 | 2.1 | ♀ | 3 | 40 | 29.5 | 34 | 40 | 29.7 | 38 | 40 | 31.1 | 42 | 4.7 | 10.5 | 0.0574 |
| Starvation | 3 | daGal4 | 2.1 | ♀ | 4 | 40 | 29.0 | 40 | 40 | 23.3 | 32 | 40 | 32.1 | 48 | 10.8 | 20.0 | **0.0412** |
| Starvation | 3 | daGal4 | 2.1 | ♀ | 5 | 40 | 29.2 | 34 | 40 | 32.5 | 42 | 41 | 35.2 | 42 | 8.2 | 0.0 | 0.2027 |
| Starvation | 3 | daGal4 | 2.1 | ♀ | 6 | 40 | 30.2 | 34 | 41 | 32.0 | 38 | 40 | 34.1 | 38 | 6.7 | 0.0 | 0.1151 |
| Starvation | 3 | daGal4 | 2.1 | ♂ | 1 | 40 | 37.9 | 53 | 40 | 38.4 | 53 | 40 | 36.7 | 48 | -3.1 | -9.4 | 0.2710 |
| Starvation | 3 | daGal4 | 2.1 | ♂ | 2 | 40 | 17.0 | 20 | 40 | 16.5 | 20 | 40 | 17.5 | 20 | 3.2 | 0.0 | 0.1174 |
| Starvation | 3 | daGal4 | 2.1 | ♂ | 3 | 40 | 21.6 | 26 | 40 | 21.9 | 26 | 41 | 23.8 | 46 | 8.5 | 76.9 | **0.0355** |
| Starvation | 3 | daGal4 | 2.1 | ♂ | 4 | 40 | 17.1 | 20 | 40 | 18.5 | 27 | 40 | 17.1 | 27 | 0.0 | 0.0 | |
| Starvation | 3 | daGal4 | 2.1 | ♂ | 5 | 40 | 22.8 | 30 | 40 | 23.1 | 30 | 40 | 23.6 | 30 | 2.2 | 0.0 | 0.2027 |

### Table S2



| Stress | Age | Gal4 | UAS | Sex | Rep[1] | UAS | | | Gal4 | | | Gal4 + UAS | | | % Extension | | *P* |
|---|---|---|---|---|---|---|---|---|---|---|---|---|---|---|---|---|---|
| | | | | | | n | Mean | Max[2] | n | Mean | Max[2] | n | Mean | Max[2] | Mean | Max[2] | |
| Starvation | 3 | daGal4 | 2.1 | ♂ | 6 | 40 | 23.4 | 30 | 40 | 24.3 | 34 | 40 | 24.3 | 30 | 0.0 | 0.0 | |
| Starvation | 3 | daGal4 | 4.3 | ♀ | 1 | 40 | 28.5 | 34 | 40 | 29.7 | 38 | 40 | 32.7 | 38 | 10.1 | 0.0 | **0.0034** |
| Starvation | 3 | daGal4 | 4.3 | ♀ | 2 | 40 | 29.6 | 38 | 40 | 32.5 | 42 | 40 | 33.9 | 38 | 4.3 | 0.0 | **0.0004** |
| Starvation | 3 | daGal4 | 4.3 | ♀ | 3 | 40 | 30.5 | 42 | 41 | 32.0 | 38 | 30 | 32.9 | 42 | 3.1 | 0.0 | **0.0306** |
| Starvation | 3 | daGal4 | 4.3 | ♂ | 1 | 40 | 23.6 | 26 | 40 | 21.9 | 26 | 40 | 25.5 | 30 | 8.1 | 15.4 | **0.0123** |
| Starvation | 3 | daGal4 | 4.3 | ♂ | 2 | 40 | 22.2 | 26 | 40 | 23.1 | 30 | 40 | 22.8 | 30 | 0.0 | 0.0 | |
| Starvation | 3 | daGal4 | 4.3 | ♂ | 3 | 40 | 22.8 | 30 | 40 | 24.3 | 34 | 40 | 24.2 | 34 | 0.0 | 0.0 | |
| Starvation | 3 | daGal4 | 9.1 | ♀ | 1 | 41 | 30.7 | 41 | 39 | 29.0 | 41 | 40 | 28.1 | 41 | -3.3 | 0.0 | 0.3129 |
| Starvation | 3 | daGal4 | 9.1 | ♀ | 2 | 40 | 30.2 | 34 | 40 | 29.7 | 38 | 40 | 30.3 | 34 | 0.3 | 0.0 | 0.6507 |
| Starvation | 3 | daGal4 | 9.1 | ♀ | 3 | 40 | 30.9 | 32 | 38 | 30.1 | 41 | 39 | 28.6 | 41 | -5.2 | 0.0 | 0.4383 |
| Starvation | 3 | daGal4 | 9.1 | ♀ | 4 | 40 | 33.5 | 42 | 40 | 32.5 | 42 | 40 | 29.9 | 38 | -8.0 | -9.5 | **0.0007** |
| Starvation | 3 | daGal4 | 9.1 | ♀ | 5 | 40 | 32.8 | 42 | 41 | 32.0 | 38 | 41 | 30.7 | 38 | -4.0 | 0.0 | **0.0115** |
| Starvation | 3 | daGal4 | 9.1 | ♂ | 1 | 40 | 13.8 | 16 | 39 | 14.4 | 23 | 39 | 16.3 | 23 | 13.2 | 0.0 | **0.0049** |
| Starvation | 3 | daGal4 | 9.1 | ♂ | 2 | 40 | 23.7 | 30 | 40 | 21.9 | 26 | 40 | 20.2 | 22 | -7.8 | -15.4 | **0.0028** |
| Starvation | 3 | daGal4 | 9.1 | ♂ | 3 | 40 | 15.0 | 23 | 38 | 14.4 | 23 | 39 | 15.6 | 23 | 4.1 | 0.0 | 0.2116 |
| Starvation | 3 | daGal4 | 9.1 | ♂ | 4 | 40 | 23.7 | 30 | 40 | 23.1 | 30 | 40 | 21.2 | 22 | -8.2 | -26.7 | **0.0002** |
| Starvation | 3 | daGal4 | 9.1 | ♂ | 5 | 40 | 25.1 | 30 | 40 | 24.3 | 34 | 40 | 21.2 | 26 | -12.8 | -13.3 | **<0.0001** |
| Starvation | 3 | daGal4 | 3.2 | ♀ | 1 | 35 | 34.0 | 52 | 40 | 55.3 | 72 | 40 | 56.4 | 77 | 2.0 | 6.9 | 0.2976 |
| Starvation | 3 | daGal4 | 3.2 | ♀ | 2 | 42 | 32.7 | 42 | 40 | 29.7 | 38 | 41 | 32.6 | 42 | 0.0 | 0.0 | |
| Starvation | 3 | daGal4 | 3.2 | ♀ | 3 | 40 | 32.5 | 38 | 40 | 32.5 | 42 | 40 | 37.0 | 46 | 13.8 | 9.5 | **0.0014** |
| Starvation | 3 | daGal4 | 3.2 | ♀ | 4 | 40 | 35.1 | 42 | 41 | 32.0 | 38 | 41 | 37.9 | 46 | 8.0 | 9.5 | **0.0054** |
| Starvation | 3 | daGal4 | 3.2 | ♂ | 1 | 40 | 27.7 | 37 | 40 | 30.6 | 42 | 40 | 38.0 | 52 | 24.1 | 23.8 | **<0.0001** |
| Starvation | 3 | daGal4 | 3.2 | ♂ | 2 | 40 | 23.9 | 30 | 40 | 21.9 | 26 | 40 | 23.3 | 30 | 0.0 | 0.0 | |
| Starvation | 3 | daGal4 | 3.2 | ♂ | 3 | 40 | 24.0 | 38 | 40 | 23.1 | 30 | 40 | 25.0 | 30 | 4.2 | 0.0 | **0.0186** |
| Starvation | 3 | daGal4 | 3.2 | ♂ | 4 | 40 | 25.0 | 34 | 40 | 24.3 | 34 | 20 | 25.2 | 30 | 0.8 | -11.8 | 0.4527 |
| Starvation | 3 | daGal4 | 10.1 | ♀ | 1 | 40 | 68.7 | 84 | 40 | 58.1 | 72 | 40 | 76.6 | 120 | 11.5 | 42.9 | **0.0146** |
| Starvation | 3 | daGal4 | 10.1 | ♀ | 2 | 40 | 30.5 | 41 | 39 | 29.0 | 41 | 40 | 34.3 | 41 | 12.4 | 0.0 | **0.0046** |
| Starvation | 3 | daGal4 | 10.1 | ♀ | 3 | 40 | 33.0 | 38 | 40 | 29.7 | 38 | 39 | 35.2 | 46 | 6.8 | 21.1 | **0.0042** |
| Starvation | 3 | daGal4 | 10.1 | ♀ | 4 | 41 | 36.7 | 50 | 40 | 32.5 | 42 | 38 | 38.5 | 50 | 4.9 | 0.0 | 0.1078 |
| Starvation | 3 | daGal4 | 10.1 | ♀ | 5 | 40 | 37.9 | 50 | 41 | 32.0 | 38 | 41 | 39.3 | 50 | 3.8 | 0.0 | 0.0693 |
| Starvation | 3 | daGal4 | 10.1 | ♂ | 1 | 40 | 54.9 | 84 | 40 | 44.8 | 59 | 40 | 59.6 | 84 | 8.6 | 0.0 | 0.1305 |
| Starvation | 3 | daGal4 | 10.1 | ♂ | 2 | 40 | 18.0 | 23 | 39 | 14.4 | 23 | 39 | 19.5 | 23 | 8.4 | 0.0 | 0.1063 |
| Starvation | 3 | daGal4 | 10.1 | ♂ | 3 | 40 | 24.7 | 30 | 40 | 21.9 | 26 | 40 | 27.4 | 34 | 10.9 | 13.3 | **0.0017** |
| Starvation | 3 | daGal4 | 10.1 | ♂ | 4 | 40 | 24.0 | 30 | 40 | 23.1 | 30 | 39 | 31.2 | 42 | 30.1 | 40.0 | **<0.0001** |
| Starvation | 3 | daGal4 | 10.1 | ♂ | 5 | 40 | 26.0 | 34 | 40 | 24.3 | 34 | 41 | 28.4 | 42 | 9.4 | 23.5 | **0.0002** |
| Starvation | 3 | daGal4 | 3.1 | ♀ | 1 | 40 | 43.8 | 54 | 40 | 41.1 | 58 | 40 | 47.4 | 58 | 8.3 | 0.0 | **0.0014** |
| Starvation | 3 | daGal4 | 3.1 | ♀ | 2 | 40 | 26.7 | 32 | 40 | 28.6 | 40 | 40 | 27.1 | 40 | 0.0 | 0.0 | |
| Starvation | 3 | daGal4 | 3.1 | ♀ | 3 | 40 | 27.8 | 34 | 40 | 29.7 | 38 | 41 | 32.1 | 38 | 8.2 | 0.0 | **0.0040** |
| Starvation | 3 | daGal4 | 3.1 | ♀ | 4 | 40 | 23.4 | 32 | 40 | 23.3 | 32 | 40 | 23.4 | 40 | 0.0 | 25.0 | |
| Starvation | 3 | daGal4 | 3.1 | ♀ | 5 | 40 | 29.8 | 34 | 40 | 32.5 | 42 | 40 | 34.5 | 42 | 6.2 | 0.0 | 0.3871 |
| Starvation | 3 | daGal4 | 3.1 | ♀ | 6 | 40 | 31.4 | 34 | 41 | 32.0 | 38 | 38 | 34.0 | 46 | 6.4 | 21.1 | **0.0006** |
| Starvation | 3 | daGal4 | 3.1 | ♂ | 1 | 40 | 28.8 | 38 | 39 | 29.7 | 38 | 40 | 28.0 | 34 | -2.6 | -10.5 | **0.0351** |
| Starvation | 3 | daGal4 | 3.1 | ♂ | 2 | 40 | 15.9 | 20 | 40 | 16.5 | 20 | 40 | 16.2 | 20 | 0.0 | 0.0 | |
| Starvation | 3 | daGal4 | 3.1 | ♂ | 3 | 40 | 21.4 | 22 | 40 | 21.9 | 26 | 40 | 23.3 | 30 | 6.4 | 15.4 | **0.0006** |
| Starvation | 3 | daGal4 | 3.1 | ♂ | 4 | 40 | 15.3 | 20 | 40 | 18.5 | 27 | 40 | 16.5 | 20 | 0.0 | 0.0 | |
| Starvation | 3 | daGal4 | 3.1 | ♂ | 5 | 40 | 23.0 | 30 | 40 | 23.1 | 30 | 40 | 24.2 | 30 | 4.8 | 0.0 | 0.0856 |
| Starvation | 3 | daGal4 | 3.1 | ♂ | 6 | 40 | 22.7 | 26 | 40 | 24.3 | 34 | 40 | 22.4 | 26 | -1.3 | 0.0 | **0.0040** |
| Starvation | 3 | daGal4 | 4.4 | ♀ | 1 | 40 | 64.8 | 84 | 40 | 62.8 | 84 | 39 | 67.5 | 84 | 4.1 | 0.0 | 0.1475 |
| Starvation | 3 | daGal4 | 4.4 | ♀ | 2 | 40 | 30.3 | 32 | 39 | 29.0 | 41 | 40 | 32.3 | 41 | 6.5 | 0.0 | 0.0642 |

### Table S2

| Stress | Age | Gal4 | UAS | Sex | Rep[1] | UAS | | | Gal4 | | | Gal4 + UAS | | | % Extension | | *P* |
|---|---|---|---|---|---|---|---|---|---|---|---|---|---|---|---|---|---|
| | | | | | | n | Mean | Max[2] | n | Mean | Max[2] | n | Mean | Max[2] | Mean | Max[2] | |
| Starvation | 3 | daGal4 | 4.4 | ♀ | 3 | 40 | 29.9 | 34 | 40 | 29.7 | 38 | 41 | 32.6 | 42 | 9.1 | 10.5 | **0.0032** |
| Starvation | 3 | daGal4 | 4.4 | ♀ | 4 | 40 | 30.9 | 32 | 38 | 30.1 | 41 | 41 | 41.0 | 48 | 33.0 | 17.1 | **<0.0001** |
| Starvation | 3 | daGal4 | 4.4 | ♀ | 5 | 41 | 34.7 | 42 | 40 | 32.5 | 42 | 40 | 34.8 | 42 | 0.3 | 0.0 | 0.1886 |
| Starvation | 3 | daGal4 | 4.4 | ♀ | 6 | 40 | 34.7 | 42 | 41 | 32.0 | 38 | 40 | 35.7 | 42 | 2.9 | 0.0 | **0.0012** |
| Starvation | 3 | daGal4 | 4.4 | ♂ | 1 | 40 | 47.5 | 59 | 40 | 49.7 | 59 | 37 | 51.9 | 72 | 4.4 | 22.0 | **0.0443** |
| Starvation | 3 | daGal4 | 4.4 | ♂ | 2 | 40 | 15.4 | 23 | 39 | 14.4 | 23 | 44 | 15.2 | 23 | 0.0 | 0.0 | |
| Starvation | 3 | daGal4 | 4.4 | ♂ | 3 | 40 | 21.8 | 26 | 40 | 21.9 | 26 | 40 | 22.6 | 30 | 3.2 | 15.4 | 0.0908 |
| Starvation | 3 | daGal4 | 4.4 | ♂ | 4 | 40 | 15.5 | 23 | 38 | 14.4 | 23 | 41 | 21.0 | 32 | 35.3 | 39.1 | **<0.0001** |
| Starvation | 3 | daGal4 | 4.4 | ♂ | 5 | 40 | 23.1 | 30 | 40 | 23.1 | 30 | 39 | 25.2 | 30 | 9.0 | 0.0 | **0.0053** |
| Starvation | 3 | daGal4 | 4.4 | ♂ | 6 | 41 | 23.3 | 26 | 40 | 24.3 | 34 | 40 | 26.1 | 34 | 7.4 | 0.0 | **0.0010** |
| Starvation | 3 | DJ694 | 2.1 | ♀ | 1 | 40 | 44.3 | 63 | 40 | 34.6 | 40 | 40 | 36.4 | 45 | 0.0 | 0.0 | |
| Starvation | 3 | DJ694 | 2.1 | ♀ | 2 | 40 | 54.3 | 73 | 40 | 54.9 | 68 | 40 | 52.9 | 63 | -2.5 | -7.4 | 0.1174 |
| Starvation | 3 | DJ694 | 2.1 | ♀ | 3 | 40 | 28.7 | 40 | 40 | 24.8 | 32 | 40 | 27.5 | 32 | 0.0 | 0.0 | |
| Starvation | 3 | DJ694 | 2.1 | ♀ | 4 | 41 | 26.0 | 34 | 41 | 25.6 | 34 | 40 | 22.6 | 34 | -11.9 | 0.0 | **0.0019** |
| Starvation | 3 | DJ694 | 2.1 | ♀ | 5 | 40 | 29.0 | 40 | 40 | 22.5 | 32 | 40 | 27.2 | 40 | 0.0 | 0.0 | |
| Starvation | 3 | DJ694 | 2.1 | ♀ | 6 | 40 | 26.4 | 34 | 42 | 28.0 | 38 | 42 | 26.4 | 34 | 0.0 | 0.0 | |
| Starvation | 3 | DJ694 | 2.1 | ♀ | 7 | 28 | 27.6 | 34 | 40 | 29.1 | 34 | 40 | 27.0 | 34 | -2.2 | 0.0 | 0.0515 |
| Starvation | 3 | DJ694 | 2.1 | ♂ | 1 | 40 | 30.2 | 45 | 40 | 24.0 | 35 | 40 | 29.6 | 45 | 0.0 | 0.0 | |
| Starvation | 3 | DJ694 | 2.1 | ♂ | 2 | 40 | 37.9 | 53 | 39 | 34.0 | 43 | 39 | 33.5 | 43 | -1.5 | 0.0 | **0.0004** |
| Starvation | 3 | DJ694 | 2.1 | ♂ | 3 | 40 | 17.0 | 20 | 40 | 16.7 | 20 | 40 | 15.9 | 20 | -4.9 | 0.0 | 0.3501 |
| Starvation | 3 | DJ694 | 2.1 | ♂ | 4 | 40 | 20.8 | 34 | 40 | 18.5 | 26 | 39 | 16.8 | 22 | -9.0 | -15.4 | **0.0254** |
| Starvation | 3 | DJ694 | 2.1 | ♂ | 5 | 40 | 17.1 | 20 | 40 | 15.6 | 27 | 40 | 17.1 | 20 | 0.0 | 0.0 | |
| Starvation | 3 | DJ694 | 2.1 | ♂ | 6 | 40 | 20.8 | 34 | 41 | 18.9 | 22 | 38 | 16.0 | 22 | -15.4 | 0.0 | **0.0001** |
| Starvation | 3 | DJ694 | 2.1 | ♂ | 7 | 39 | 20.1 | 34 | 40 | 17.9 | 22 | 37 | 17.6 | 22 | -1.4 | 0.0 | **0.0189** |
| Starvation | 3 | DJ694 | 4.3 | ♀ | 1 | 40 | 25.7 | 34 | 41 | 25.6 | 34 | 39 | 23.1 | 34 | -9.8 | 0.0 | **0.0118** |
| Starvation | 3 | DJ694 | 4.3 | ♀ | 2 | 42 | 25.7 | 34 | 42 | 28.0 | 38 | 40 | 22.0 | 26 | -14.5 | -23.5 | **0.0001** |
| Starvation | 3 | DJ694 | 4.3 | ♀ | 3 | 40 | 28.1 | 34 | 40 | 29.1 | 34 | 40 | 24.7 | 34 | -12.1 | 0.0 | **0.0041** |
| Starvation | 3 | DJ694 | 4.3 | ♂ | 1 | 39 | 18.6 | 26 | 40 | 18.5 | 26 | 41 | 15.0 | 22 | -18.7 | -15.4 | **<0.0001** |
| Starvation | 3 | DJ694 | 4.3 | ♂ | 2 | 40 | 19.2 | 26 | 41 | 18.9 | 22 | 40 | 15.2 | 22 | -19.5 | 0.0 | **<0.0001** |
| Starvation | 3 | DJ694 | 4.3 | ♂ | 3 | 41 | 18.6 | 26 | 40 | 17.9 | 22 | 37 | 16.1 | 22 | -10.1 | 0.0 | **0.0041** |
| Starvation | 3 | DJ694 | 9.1 | ♀ | 1 | 41 | 30.7 | 41 | 39 | 30.0 | 32 | 40 | 31.4 | 32 | 2.3 | 0.0 | **0.0437** |
| Starvation | 3 | DJ694 | 9.1 | ♀ | 2 | 40 | 27.9 | 34 | 41 | 25.6 | 34 | 39 | 27.1 | 34 | 0.0 | 0.0 | |
| Starvation | 3 | DJ694 | 9.1 | ♀ | 3 | 40 | 30.9 | 32 | 40 | 27.6 | 32 | 40 | 32.0 | 41 | 3.6 | 28.1 | 0.1301 |
| Starvation | 3 | DJ694 | 9.1 | ♀ | 4 | 49 | 27.8 | 34 | 42 | 28.0 | 38 | 40 | 23.1 | 34 | -16.9 | 0.0 | **<0.0001** |
| Starvation | 3 | DJ694 | 9.1 | ♀ | 5 | 41 | 29.1 | 34 | 40 | 29.1 | 34 | 40 | 24.1 | 34 | -17.2 | 0.0 | **<0.0001** |
| Starvation | 3 | DJ694 | 9.1 | ♂ | 1 | 40 | 13.8 | 16 | 39 | 14.3 | 23 | 40 | 13.4 | 16 | -2.7 | 0.0 | 0.1750 |
| Starvation | 3 | DJ694 | 9.1 | ♂ | 2 | 40 | 18.9 | 26 | 40 | 18.5 | 26 | 40 | 14.3 | 22 | -22.4 | -15.4 | **<0.0001** |
| Starvation | 3 | DJ694 | 9.1 | ♂ | 3 | 40 | 15.0 | 23 | 38 | 13.6 | 16 | 40 | 14.1 | 23 | 0.0 | 0.0 | |
| Starvation | 3 | DJ694 | 9.1 | ♂ | 4 | 50 | 20.0 | 34 | 41 | 18.9 | 22 | 41 | 14.6 | 18 | -22.6 | -18.2 | **<0.0001** |
| Starvation | 3 | DJ694 | 9.1 | ♂ | 5 | 40 | 19.3 | 26 | 40 | 17.9 | 22 | 37 | 14.2 | 18 | -20.5 | -18.2 | **<0.0001** |
| Starvation | 3 | DJ694 | 3.2 | ♀ | 1 | 35 | 34.0 | 52 | 40 | 51.9 | 62 | 40 | 42.0 | 57 | 0.0 | 0.0 | |
| Starvation | 3 | DJ694 | 3.2 | ♀ | 2 | 39 | 26.5 | 34 | 41 | 25.6 | 34 | 41 | 28.1 | 34 | 6.3 | 0.0 | **0.0265** |
| Starvation | 3 | DJ694 | 3.2 | ♀ | 3 | 39 | 30.1 | 38 | 42 | 28.0 | 38 | 40 | 28.9 | 38 | 0.0 | 0.0 | |
| Starvation | 3 | DJ694 | 3.2 | ♀ | 4 | 37 | 27.8 | 38 | 40 | 29.1 | 34 | 37 | 26.9 | 38 | -3.5 | 0.0 | 0.0638 |
| Starvation | 3 | DJ694 | 3.2 | ♂ | 1 | 40 | 27.7 | 37 | 40 | 31.1 | 42 | 40 | 30.0 | 37 | 0.0 | 0.0 | |
| Starvation | 3 | DJ694 | 3.2 | ♂ | 2 | 36 | 21.3 | 34 | 40 | 18.5 | 26 | 39 | 18.2 | 34 | -1.5 | 0.0 | **0.0349** |
| Starvation | 3 | DJ694 | 3.2 | ♂ | 3 | 40 | 21.4 | 34 | 41 | 18.9 | 22 | 41 | 17.7 | 34 | -6.2 | 0.0 | **0.0153** |
| Starvation | 3 | DJ694 | 3.2 | ♂ | 4 | 40 | 21.2 | 34 | 40 | 17.9 | 22 | 41 | 16.9 | 26 | -5.4 | 0.0 | **0.0026** |
| Starvation | 3 | DJ694 | 10.1 | ♀ | 1 | 40 | 68.7 | 84 | 37 | 49.9 | 72 | 40 | 63.7 | 84 | 0.0 | 0.0 | |

**Table S2**



| Stress | Age | Gal4 | UAS | Sex | Rep[1] | UAS | | | Gal4 | | | Gal4 + UAS | | | % Extension | | *P* |
|---|---|---|---|---|---|---|---|---|---|---|---|---|---|---|---|---|---|
| | | | | | | n | Mean | Max[2] | n | Mean | Max[2] | n | Mean | Max[2] | Mean | Max[2] | |
| Starvation | 3 | DJ694 | 10.1 | ♀ | 2 | 40 | 30.5 | 41 | 39 | 30.0 | 32 | 40 | 30.7 | 41 | 0.7 | 0.0 | 0.4364 |
| Starvation | 3 | DJ694 | 10.1 | ♀ | 3 | 39 | 27.7 | 34 | 41 | 25.6 | 34 | 41 | 30.1 | 38 | 8.6 | 11.8 | **0.0002** |
| Starvation | 3 | DJ694 | 10.1 | ♀ | 4 | 35 | 29.2 | 34 | 42 | 28.0 | 38 | 20 | 27.2 | 34 | -2.9 | 0.0 | 0.0906 |
| Starvation | 3 | DJ694 | 10.1 | ♀ | 5 | 25 | 30.6 | 38 | 40 | 29.1 | 34 | 39 | 26.2 | 38 | -10.1 | 0.0 | **0.0090** |
| Starvation | 3 | DJ694 | 10.1 | ♂ | 1 | 40 | 54.9 | 84 | 40 | 36.9 | 47 | 40 | 48.7 | 72 | 0.0 | 0.0 | |
| Starvation | 3 | DJ694 | 10.1 | ♂ | 2 | 40 | 18.0 | 23 | 39 | 14.3 | 23 | 40 | 15.0 | 16 | 0.0 | -30.4 | |
| Starvation | 3 | DJ694 | 10.1 | ♂ | 3 | 37 | 21.4 | 34 | 40 | 18.5 | 26 | 42 | 23.5 | 34 | 9.9 | 0.0 | 0.0610 |
| Starvation | 3 | DJ694 | 10.1 | ♂ | 4 | 41 | 20.0 | 34 | 41 | 18.9 | 22 | 40 | 20.7 | 26 | 3.2 | 0.0 | **0.0134** |
| Starvation | 3 | DJ694 | 10.1 | ♂ | 5 | 41 | 23.3 | 34 | 40 | 17.9 | 22 | 42 | 22.2 | 34 | 0.0 | 0.0 | |
| Starvation | 3 | DJ694 | 3.1 | ♀ | 1 | 40 | 32.2 | 51 | 40 | 34.6 | 40 | 40 | 32.0 | 57 | -0.8 | 11.8 | 0.2482 |
| Starvation | 3 | DJ694 | 3.1 | ♀ | 2 | 40 | 43.8 | 54 | 40 | 43.1 | 50 | 40 | 40.1 | 46 | -7.0 | -8.0 | **0.0004** |
| Starvation | 3 | DJ694 | 3.1 | ♀ | 3 | 40 | 26.7 | 32 | 40 | 24.8 | 32 | 40 | 24.6 | 32 | -0.8 | 0.0 | **0.0166** |
| Starvation | 3 | DJ694 | 3.1 | ♀ | 4 | 39 | 30.4 | 34 | 41 | 25.6 | 34 | 35 | 23.5 | 34 | -8.2 | 0.0 | **0.0226** |
| Starvation | 3 | DJ694 | 3.1 | ♀ | 5 | 40 | 23.4 | 32 | 40 | 22.5 | 32 | 40 | 22.9 | 27 | 0.0 | -15.6 | |
| Starvation | 3 | DJ694 | 3.1 | ♀ | 6 | 44 | 31.2 | 38 | 42 | 28.0 | 38 | 40 | 23.3 | 26 | -16.8 | -31.6 | **<0.0001** |
| Starvation | 3 | DJ694 | 3.1 | ♀ | 7 | 39 | 29.6 | 34 | 40 | 29.1 | 34 | 38 | 25.8 | 34 | -11.5 | 0.0 | **0.0009** |
| Starvation | 3 | DJ694 | 3.1 | ♂ | 1 | 40 | 26.8 | 45 | 40 | 24.0 | 35 | 40 | 20.3 | 30 | -15.2 | -14.3 | **0.0025** |
| Starvation | 3 | DJ694 | 3.1 | ♂ | 2 | 40 | 28.8 | 38 | 40 | 29.0 | 38 | 40 | 25.4 | 34 | -11.7 | -10.5 | **0.0008** |
| Starvation | 3 | DJ694 | 3.1 | ♂ | 3 | 40 | 15.9 | 20 | 40 | 16.7 | 20 | 40 | 15.4 | 20 | -3.5 | 0.0 | 0.1466 |
| Starvation | 3 | DJ694 | 3.1 | ♂ | 4 | 40 | 19.3 | 22 | 40 | 18.5 | 26 | 37 | 16.8 | 22 | -8.7 | 0.0 | **0.0002** |
| Starvation | 3 | DJ694 | 3.1 | ♂ | 5 | 40 | 15.3 | 20 | 40 | 15.6 | 27 | 40 | 14.4 | 17 | -6.0 | -15.0 | **0.0377** |
| Starvation | 3 | DJ694 | 3.1 | ♂ | 6 | 42 | 19.4 | 22 | 41 | 18.9 | 22 | 40 | 18.3 | 26 | -2.9 | 18.2 | 0.1782 |
| Starvation | 3 | DJ694 | 3.1 | ♂ | 7 | 40 | 19.3 | 26 | 40 | 17.9 | 22 | 41 | 17.3 | 22 | -2.8 | 0.0 | **0.0056** |
| Starvation | 3 | DJ694 | 4.4 | ♀ | 1 | 40 | 35.8 | 43 | 40 | 29.6 | 34 | 40 | 32.1 | 43 | 0.0 | 0.0 | |
| Starvation | 3 | DJ694 | 4.4 | ♀ | 2 | 40 | 64.8 | 84 | 40 | 61.6 | 84 | 40 | 53.9 | 72 | -12.5 | -14.3 | **0.0009** |
| Starvation | 3 | DJ694 | 4.4 | ♀ | 3 | 40 | 30.3 | 32 | 39 | 30.0 | 32 | 40 | 31.8 | 41 | 4.9 | 28.1 | **0.0388** |
| Starvation | 3 | DJ694 | 4.4 | ♀ | 4 | 39 | 27.5 | 34 | 41 | 25.6 | 34 | 41 | 22.6 | 34 | -11.7 | 0.0 | **0.0019** |
| Starvation | 3 | DJ694 | 4.4 | ♀ | 5 | 40 | 30.9 | 32 | 40 | 27.6 | 32 | 39 | 30.4 | 32 | 0.0 | 0.0 | |
| Starvation | 3 | DJ694 | 4.4 | ♀ | 6 | 48 | 30.0 | 34 | 42 | 28.0 | 38 | 41 | 24.1 | 34 | -13.8 | 0.0 | **0.0003** |
| Starvation | 3 | DJ694 | 4.4 | ♀ | 7 | 38 | 29.9 | 34 | 40 | 29.1 | 34 | 37 | 24.8 | 34 | -14.8 | 0.0 | **0.0002** |
| Starvation | 3 | DJ694 | 4.4 | ♂ | 1 | 40 | 21.3 | 24 | 40 | 26.4 | 29 | 40 | 20.3 | 29 | -4.8 | 0.0 | 0.0786 |
| Starvation | 3 | DJ694 | 4.4 | ♂ | 2 | 40 | 47.5 | 59 | 40 | 44.1 | 59 | 40 | 40.4 | 46 | -8.3 | -22.0 | **0.0001** |
| Starvation | 3 | DJ694 | 4.4 | ♂ | 3 | 40 | 15.4 | 23 | 39 | 14.3 | 23 | 40 | 13.7 | 23 | -4.1 | 0.0 | **0.0145** |
| Starvation | 3 | DJ694 | 4.4 | ♂ | 4 | 39 | 18.9 | 26 | 40 | 18.5 | 26 | 40 | 15.8 | 22 | -14.4 | -15.4 | **0.0010** |
| Starvation | 3 | DJ694 | 4.4 | ♂ | 5 | 40 | 15.5 | 23 | 38 | 13.6 | 16 | 40 | 14.2 | 23 | 0.0 | 0.0 | |
| Starvation | 3 | DJ694 | 4.4 | ♂ | 6 | 39 | 17.5 | 22 | 41 | 18.9 | 22 | 42 | 16.0 | 26 | -8.8 | 18.2 | **0.0004** |
| Starvation | 3 | DJ694 | 4.4 | ♂ | 7 | 39 | 18.9 | 26 | 40 | 17.9 | 22 | 40 | 14.3 | 22 | -19.7 | 0.0 | **<0.0001** |
| Starvation | 3 | D42 | 2.1 | ♀ | 1 | 40 | 44.3 | 63 | 40 | 38.0 | 45 | 40 | 32.0 | 51 | -15.7 | 0.0 | **0.0002** |
| Starvation | 3 | D42 | 2.1 | ♀ | 2 | 40 | 54.3 | 73 | 40 | 53.1 | 63 | 50 | 55.7 | 63 | 2.7 | 0.0 | 0.2767 |
| Starvation | 3 | D42 | 2.1 | ♀ | 3 | 40 | 28.7 | 40 | 40 | 24.9 | 32 | 40 | 33.9 | 40 | 18.2 | 0.0 | **0.0053** |
| Starvation | 3 | D42 | 2.1 | ♀ | 4 | 40 | 36.3 | 46 | 40 | 39.0 | 50 | 42 | 37.1 | 46 | 0.0 | 0.0 | |
| Starvation | 3 | D42 | 2.1 | ♀ | 5 | 40 | 29.0 | 40 | 40 | 22.5 | 32 | 40 | 33.8 | 40 | 16.8 | 0.0 | **0.0008** |
| Starvation | 3 | D42 | 2.1 | ♀ | 6 | 40 | 37.0 | 42 | 40 | 39.9 | 50 | 40 | 38.3 | 46 | 0.0 | 0.0 | |
| Starvation | 3 | D42 | 2.1 | ♀ | 7 | 40 | 36.7 | 42 | 42 | 39.5 | 50 | 40 | 40.1 | 50 | 1.5 | 0.0 | **0.0005** |
| Starvation | 3 | D42 | 2.1 | ♂ | 1 | 40 | 30.2 | 45 | 40 | 27.6 | 35 | 40 | 26.5 | 35 | -4.1 | 0.0 | **0.0014** |
| Starvation | 3 | D42 | 2.1 | ♂ | 2 | 40 | 37.9 | 53 | 40 | 36.4 | 48 | 40 | 36.6 | 48 | 0.0 | 0.0 | |
| Starvation | 3 | D42 | 2.1 | ♂ | 3 | 40 | 17.0 | 20 | 40 | 17.1 | 20 | 40 | 18.2 | 20 | 6.3 | 0.0 | **0.0011** |
| Starvation | 3 | D42 | 2.1 | ♂ | 4 | 40 | 26.7 | 34 | 40 | 27.7 | 42 | 40 | 28.3 | 38 | 2.2 | 0.0 | 0.0840 |
| Starvation | 3 | D42 | 2.1 | ♂ | 5 | 40 | 17.1 | 20 | 40 | 14.0 | 17 | 40 | 18.9 | 20 | 10.4 | 0.0 | **<0.0001** |

**Table S2**



| Stress | Age | Gal4 | UAS | Sex | Rep[1] | UAS | | | Gal4 | | | Gal4 + UAS | | | % Extension | | *P* |
|---|---|---|---|---|---|---|---|---|---|---|---|---|---|---|---|---|---|
| | | | | | | n | Mean | Max[2] | n | Mean | Max[2] | n | Mean | Max[2] | Mean | Max[2] | |
| Starvation | 3 | D42 | 2.1 | ♂ | 6 | 40 | 28.9 | 38 | 40 | 27.9 | 38 | 40 | 27.7 | 34 | -0.7 | -10.5 | 0.3159 |
| Starvation | 3 | D42 | 2.1 | ♂ | 7 | 40 | 26.7 | 34 | 40 | 28.3 | 38 | 40 | 26.4 | 38 | -1.1 | 0.0 | 0.1402 |
| Starvation | 3 | D42 | 4.3 | ♀ | 1 | 40 | 36.1 | 42 | 40 | 39.0 | 50 | 40 | 39.2 | 46 | 0.5 | 0.0 | **0.0005** |
| Starvation | 3 | D42 | 4.3 | ♀ | 2 | 40 | 37.8 | 46 | 40 | 39.9 | 50 | 40 | 38.6 | 46 | 0.0 | 0.0 | |
| Starvation | 3 | D42 | 4.3 | ♀ | 3 | 43 | 36.0 | 42 | 42 | 39.5 | 50 | 40 | 38.0 | 46 | 0.0 | 0.0 | |
| Starvation | 3 | D42 | 4.3 | ♂ | 1 | 40 | 30.2 | 42 | 40 | 27.7 | 42 | 40 | 28.4 | 34 | 0.0 | -19.0 | |
| Starvation | 3 | D42 | 4.3 | ♂ | 2 | 41 | 28.0 | 34 | 40 | 27.9 | 38 | 40 | 28.2 | 34 | 0.9 | 0.0 | 0.6510 |
| Starvation | 3 | D42 | 4.3 | ♂ | 3 | 40 | 27.1 | 38 | 40 | 28.3 | 38 | 40 | 23.8 | 30 | -12.2 | -21.1 | **0.0002** |
| Starvation | 3 | D42 | 9.1 | ♀ | 1 | 41 | 30.7 | 41 | 40 | 31.8 | 41 | 43 | 32.2 | 41 | 1.2 | 0.0 | 0.0692 |
| Starvation | 3 | D42 | 9.1 | ♀ | 2 | 40 | 38.0 | 42 | 40 | 39.0 | 50 | 40 | 36.0 | 46 | -5.3 | 0.0 | **0.0165** |
| Starvation | 3 | D42 | 9.1 | ♀ | 3 | 40 | 30.9 | 32 | 38 | 30.3 | 41 | 41 | 31.0 | 32 | 0.6 | 0.0 | 0.3354 |
| Starvation | 3 | D42 | 9.1 | ♀ | 4 | 42 | 39.4 | 46 | 40 | 39.9 | 50 | 40 | 34.9 | 42 | -11.5 | -8.7 | **<0.0001** |
| Starvation | 3 | D42 | 9.1 | ♀ | 5 | 40 | 39.2 | 50 | 42 | 39.5 | 50 | 40 | 38.3 | 42 | -2.3 | -16.0 | 0.1892 |
| Starvation | 3 | D42 | 9.1 | ♂ | 1 | 40 | 13.8 | 16 | 40 | 12.8 | 16 | 44 | 14.1 | 23 | 2.8 | 43.8 | 0.0564 |
| Starvation | 3 | D42 | 9.1 | ♂ | 2 | 40 | 29.8 | 42 | 40 | 27.7 | 42 | 40 | 24.9 | 34 | -10.1 | -19.0 | **0.0109** |
| Starvation | 3 | D42 | 9.1 | ♂ | 3 | 40 | 15.0 | 23 | 40 | 13.3 | 16 | 43 | 13.7 | 16 | 0.0 | 0.0 | |
| Starvation | 3 | D42 | 9.1 | ♂ | 4 | 40 | 27.1 | 38 | 40 | 27.9 | 38 | 40 | 24.5 | 34 | -9.6 | -10.5 | **0.0047** |
| Starvation | 3 | D42 | 9.1 | ♂ | 5 | 40 | 28.6 | 42 | 40 | 28.3 | 38 | 40 | 23.5 | 30 | -17.0 | -21.1 | **<0.0001** |
| Starvation | 3 | D42 | 3.2 | ♀ | 1 | 35 | 34.0 | 52 | 42 | 51.5 | 67 | 40 | 51.4 | 62 | 0.0 | 0.0 | |
| Starvation | 3 | D42 | 3.2 | ♀ | 2 | 40 | 40.5 | 50 | 40 | 39.0 | 50 | 40 | 41.5 | 50 | 2.5 | 0.0 | **0.0279** |
| Starvation | 3 | D42 | 3.2 | ♀ | 3 | 40 | 41.1 | 50 | 40 | 39.9 | 50 | 40 | 41.0 | 50 | 0.0 | 0.0 | |
| Starvation | 3 | D42 | 3.2 | ♀ | 4 | 40 | 41.3 | 50 | 42 | 39.5 | 50 | 40 | 42.1 | 50 | 1.9 | 0.0 | 0.1144 |
| Starvation | 3 | D42 | 3.2 | ♂ | 1 | 40 | 27.7 | 37 | 40 | 31.1 | 37 | 40 | 31.9 | 47 | 2.4 | 27.0 | **0.0137** |
| Starvation | 3 | D42 | 3.2 | ♂ | 2 | 40 | 28.9 | 38 | 40 | 27.7 | 42 | 40 | 29.1 | 38 | 0.7 | 0.0 | 0.3563 |
| Starvation | 3 | D42 | 3.2 | ♂ | 3 | 40 | 30.3 | 42 | 40 | 27.9 | 38 | 40 | 28.6 | 38 | 0.0 | 0.0 | |
| Starvation | 3 | D42 | 3.2 | ♂ | 4 | 40 | 33.2 | 42 | 40 | 28.3 | 38 | 40 | 29.2 | 38 | 0.0 | 0.0 | |
| Starvation | 3 | D42 | 10.1 | ♀ | 1 | 40 | 68.7 | 84 | 40 | 54.0 | 84 | 40 | 72.7 | 96 | 5.8 | 14.3 | 0.1433 |
| Starvation | 3 | D42 | 10.1 | ♀ | 2 | 40 | 30.5 | 41 | 40 | 31.8 | 41 | 39 | 34.5 | 41 | 8.4 | 0.0 | **0.0035** |
| Starvation | 3 | D42 | 10.1 | ♀ | 3 | 40 | 41.8 | 46 | 40 | 39.0 | 50 | 40 | 39.2 | 46 | 0.0 | 0.0 | |
| Starvation | 3 | D42 | 10.1 | ♀ | 4 | 39 | 45.5 | 59 | 40 | 39.9 | 50 | 40 | 40.6 | 46 | 0.0 | -8.0 | |
| Starvation | 3 | D42 | 10.1 | ♀ | 5 | 40 | 42.5 | 50 | 42 | 39.5 | 50 | 41 | 41.4 | 50 | 0.0 | 0.0 | |
| Starvation | 3 | D42 | 10.1 | ♂ | 1 | 40 | 54.9 | 84 | 40 | 46.2 | 59 | 40 | 53.7 | 72 | 0.0 | 0.0 | |
| Starvation | 3 | D42 | 10.1 | ♂ | 2 | 40 | 18.0 | 23 | 40 | 12.8 | 16 | 40 | 17.4 | 23 | 0.0 | 0.0 | |
| Starvation | 3 | D42 | 10.1 | ♂ | 3 | 40 | 31.7 | 42 | 40 | 27.7 | 42 | 40 | 33.1 | 42 | 4.4 | 0.0 | **0.0002** |
| Starvation | 3 | D42 | 10.1 | ♂ | 4 | 41 | 33.6 | 42 | 40 | 27.9 | 38 | 42 | 34.7 | 42 | 3.1 | 0.0 | 0.4653 |
| Starvation | 3 | D42 | 10.1 | ♂ | 5 | 40 | 30.5 | 38 | 40 | 28.3 | 38 | 40 | 34.5 | 42 | 13.1 | 10.5 | **0.0005** |
| Starvation | 3 | D42 | 3.1 | ♀ | 1 | 40 | 32.2 | 51 | 40 | 38.0 | 45 | 40 | 36.5 | 57 | 0.0 | 11.8 | 0.0982 |
| Starvation | 3 | D42 | 3.1 | ♀ | 2 | 40 | 43.8 | 54 | 37 | 41.1 | 50 | 40 | 44.7 | 54 | 2.1 | 0.0 | **0.0012** |
| Starvation | 3 | D42 | 3.1 | ♀ | 3 | 40 | 26.7 | 32 | 40 | 24.9 | 32 | 40 | 29.2 | 40 | 9.4 | 25.0 | **0.0003** |
| Starvation | 3 | D42 | 3.1 | ♀ | 4 | 40 | 36.8 | 42 | 40 | 39.0 | 50 | 42 | 37.1 | 46 | 0.0 | 0.0 | |
| Starvation | 3 | D42 | 3.1 | ♀ | 5 | 40 | 23.4 | 32 | 40 | 22.5 | 32 | 40 | 29.2 | 32 | 24.6 | 0.0 | **<0.0001** |
| Starvation | 3 | D42 | 3.1 | ♀ | 6 | 40 | 38.4 | 46 | 40 | 39.9 | 50 | 42 | 37.2 | 46 | -3.0 | 0.0 | **0.0213** |
| Starvation | 3 | D42 | 3.1 | ♀ | 7 | 40 | 37.7 | 46 | 42 | 39.5 | 50 | 40 | 37.5 | 42 | -0.5 | -8.7 | 0.0568 |
| Starvation | 3 | D42 | 3.1 | ♂ | 1 | 40 | 26.8 | 45 | 40 | 27.6 | 35 | 40 | 25.9 | 35 | -3.5 | 0.0 | 0.0850 |
| Starvation | 3 | D42 | 3.1 | ♂ | 2 | 40 | 28.8 | 38 | 40 | 25.0 | 34 | 40 | 27.3 | 31 | 0.0 | -8.8 | |
| Starvation | 3 | D42 | 3.1 | ♂ | 3 | 40 | 15.9 | 20 | 40 | 17.1 | 20 | 40 | 18.7 | 27 | 8.9 | 35.0 | **0.0001** |
| Starvation | 3 | D42 | 3.1 | ♂ | 4 | 40 | 24.7 | 34 | 40 | 27.7 | 42 | 40 | 26.9 | 34 | 0.0 | 0.0 | |
| Starvation | 3 | D42 | 3.1 | ♂ | 5 | 40 | 15.3 | 20 | 40 | 14.0 | 17 | 40 | 18.5 | 27 | 20.8 | 35.0 | **<0.0001** |
| Starvation | 3 | D42 | 3.1 | ♂ | 6 | 40 | 25.9 | 34 | 40 | 27.9 | 38 | 40 | 26.8 | 34 | 0.0 | 0.0 | |

### Table S2



| Stress | Age | Gal4 | UAS | Sex | Rep[1] | UAS | | | Gal4 | | | Gal4 + UAS | | | % Extension | | *P* |
|---|---|---|---|---|---|---|---|---|---|---|---|---|---|---|---|---|---|
| | | | | | | n | Mean | Max[2] | n | Mean | Max[2] | n | Mean | Max[2] | Mean | Max[2] | |
| Starvation | 3 | D42 | 3.1 | ♂ | 7 | 40 | 27.0 | 34 | 40 | 28.3 | 38 | 40 | 25.6 | 34 | -5.2 | 0.0 | **0.0147** |
| Starvation | 3 | D42 | 4.4 | ♀ | 1 | 40 | 35.8 | 43 | 40 | 29.7 | 34 | 40 | 31.3 | 34 | 0.0 | 0.0 | |
| Starvation | 3 | D42 | 4.4 | ♀ | 2 | 40 | 64.8 | 84 | 39 | 72.6 | 84 | 40 | 68.8 | 84 | 0.0 | 0.0 | |
| Starvation | 3 | D42 | 4.4 | ♀ | 3 | 40 | 30.3 | 32 | 40 | 31.8 | 41 | 42 | 33.9 | 48 | 6.6 | 17.1 | **0.0015** |
| Starvation | 3 | D42 | 4.4 | ♀ | 4 | 40 | 39.5 | 50 | 40 | 39.0 | 50 | 40 | 37.4 | 42 | -4.1 | -16.0 | 0.1325 |
| Starvation | 3 | D42 | 4.4 | ♀ | 5 | 40 | 30.9 | 32 | 38 | 30.3 | 41 | 41 | 39.2 | 41 | 26.9 | 0.0 | **<0.0001** |
| Starvation | 3 | D42 | 4.4 | ♀ | 6 | 40 | 41.4 | 46 | 40 | 39.9 | 50 | 40 | 36.1 | 42 | -9.5 | -8.7 | **0.0015** |
| Starvation | 3 | D42 | 4.4 | ♀ | 7 | 40 | 39.2 | 50 | 42 | 39.5 | 50 | 40 | 34.4 | 42 | -12.2 | -16.0 | **<0.0001** |
| Starvation | 3 | D42 | 4.4 | ♂ | 1 | 40 | 21.3 | 24 | 40 | 22.4 | 29 | 40 | 23.5 | 29 | 5.0 | 0.0 | **0.0033** |
| Starvation | 3 | D42 | 4.4 | ♂ | 2 | 40 | 47.5 | 59 | 34 | 44.5 | 59 | 40 | 47.2 | 72 | 0.0 | 22.0 | |
| Starvation | 3 | D42 | 4.4 | ♂ | 3 | 40 | 15.4 | 23 | 40 | 12.8 | 16 | 40 | 15.8 | 23 | 2.6 | 0.0 | 0.6505 |
| Starvation | 3 | D42 | 4.4 | ♂ | 4 | 40 | 27.0 | 38 | 40 | 27.7 | 42 | 40 | 27.8 | 34 | 0.4 | -10.5 | 0.4817 |
| Starvation | 3 | D42 | 4.4 | ♂ | 5 | 40 | 15.5 | 23 | 40 | 13.3 | 16 | 41 | 22.7 | 32 | 46.2 | 39.1 | **<0.0001** |
| Starvation | 3 | D42 | 4.4 | ♂ | 6 | 40 | 27.9 | 38 | 40 | 27.9 | 38 | 40 | 26.5 | 38 | -5.0 | 0.0 | 0.3560 |
| Starvation | 3 | D42 | 4.4 | ♂ | 7 | 40 | 27.6 | 34 | 40 | 28.3 | 38 | 40 | 25.5 | 38 | -7.6 | 0.0 | **0.0402** |
| Starvation | 10 | daGal4 | 2.1 | ♀ | 1 | 40 | 49.5 | 63 | 40 | 48.8 | 63 | 40 | 46.2 | 58 | -5.4 | -7.9 | **0.0208** |
| Starvation | 10 | daGal4 | 2.1 | ♀ | 2 | 40 | 31.0 | 38 | 40 | 33.3 | 38 | 41 | 29.2 | 38 | -5.7 | 0.0 | 0.0995 |
| Starvation | 10 | daGal4 | 2.1 | ♀ | 3 | 40 | 29.3 | 34 | 41 | 36.3 | 42 | 43 | 31.2 | 38 | 0.0 | 0.0 | |
| Starvation | 10 | daGal4 | 2.1 | ♀ | 4 | 40 | 30.2 | 34 | 40 | 35.5 | 42 | 40 | 31.2 | 38 | 0.0 | 0.0 | |
| Starvation | 10 | daGal4 | 2.1 | ♂ | 1 | 39 | 27.9 | 38 | 36 | 28.3 | 33 | 40 | 27.5 | 38 | -1.8 | 0.0 | 0.4315 |
| Starvation | 10 | daGal4 | 2.1 | ♂ | 2 | 41 | 15.7 | 18 | 38 | 16.5 | 18 | 40 | 15.8 | 22 | 0.0 | 22.2 | |
| Starvation | 10 | daGal4 | 2.1 | ♂ | 3 | 40 | 17.9 | 22 | 39 | 17.3 | 18 | 40 | 16.3 | 18 | -5.7 | 0.0 | **0.0012** |
| Starvation | 10 | daGal4 | 2.1 | ♂ | 4 | 40 | 16.7 | 18 | 39 | 17.6 | 22 | 40 | 16.0 | 22 | -4.2 | 0.0 | **0.0114** |
| Starvation | 10 | daGal4 | 4.3 | ♀ | 1 | 40 | 48.6 | 59 | 38 | 51.1 | 72 | 40 | 60.8 | 84 | 18.9 | 16.7 | **0.0001** |
| Starvation | 10 | daGal4 | 4.3 | ♀ | 2 | 40 | 29.6 | 38 | 40 | 25.5 | 38 | 40 | 27.8 | 48 | 0.0 | 26.3 | |
| Starvation | 10 | daGal4 | 4.3 | ♀ | 3 | 40 | 30.7 | 38 | 40 | 33.3 | 38 | 40 | 33.5 | 38 | 0.6 | 0.0 | **0.0489** |
| Starvation | 10 | daGal4 | 4.3 | ♀ | 4 | 40 | 32.6 | 43 | 40 | 24.8 | 33 | 40 | 29.2 | 48 | 0.0 | 11.6 | |
| Starvation | 10 | daGal4 | 4.3 | ♀ | 5 | 40 | 30.0 | 38 | 41 | 36.3 | 42 | 40 | 35.1 | 42 | 0.0 | 0.0 | |
| Starvation | 10 | daGal4 | 4.3 | ♀ | 6 | 39 | 31.5 | 38 | 40 | 35.5 | 42 | 40 | 35.9 | 46 | 1.1 | 9.5 | **0.0002** |
| Starvation | 10 | daGal4 | 4.3 | ♂ | 1 | 37 | 26.6 | 34 | 40 | 23.6 | 34 | 40 | 34.3 | 47 | 28.9 | 38.2 | **<0.0001** |
| Starvation | 10 | daGal4 | 4.3 | ♂ | 2 | 40 | 11.6 | 16 | 40 | 10.4 | 16 | 40 | 13.9 | 33 | 19.6 | 106.3 | **0.0147** |
| Starvation | 10 | daGal4 | 4.3 | ♂ | 3 | 40 | 17.3 | 22 | 38 | 16.5 | 18 | 41 | 17.1 | 18 | 0.0 | 0.0 | |
| Starvation | 10 | daGal4 | 4.3 | ♂ | 4 | 40 | 17.2 | 18 | 39 | 17.3 | 18 | 39 | 17.7 | 22 | 2.4 | 22.2 | 0.5355 |
| Starvation | 10 | daGal4 | 4.3 | ♂ | 5 | 39 | 16.1 | 18 | 39 | 17.6 | 22 | 39 | 17.6 | 22 | 0.0 | 0.0 | |
| Starvation | 10 | daGal4 | 9.1 | ♀ | 1 | 39 | 32.7 | 40 | 39 | 25.6 | 40 | 39 | 27.3 | 40 | 0.0 | 0.0 | |
| Starvation | 10 | daGal4 | 9.1 | ♀ | 2 | 39 | 31.6 | 38 | 40 | 33.3 | 38 | 40 | 30.8 | 38 | -2.7 | 0.0 | **0.0004** |
| Starvation | 10 | daGal4 | 9.1 | ♀ | 3 | 40 | 31.3 | 40 | 40 | 26.6 | 40 | 34 | 22.7 | 40 | -14.6 | 0.0 | **0.0176** |
| Starvation | 10 | daGal4 | 9.1 | ♀ | 4 | 40 | 33.3 | 42 | 41 | 36.3 | 42 | 40 | 30.2 | 38 | -9.3 | -9.5 | **0.0003** |
| Starvation | 10 | daGal4 | 9.1 | ♀ | 5 | 40 | 34.2 | 42 | 40 | 35.5 | 42 | 40 | 29.1 | 34 | -14.9 | -19.0 | **<0.0001** |
| Starvation | 10 | daGal4 | 9.1 | ♂ | 1 | 39 | 11.3 | 15 | 39 | 11.2 | 15 | 37 | 10.1 | 10 | -9.3 | -33.3 | **0.0087** |
| Starvation | 10 | daGal4 | 9.1 | ♂ | 2 | 40 | 16.4 | 18 | 38 | 16.5 | 18 | 40 | 16.0 | 18 | -2.4 | 0.0 | 0.3321 |
| Starvation | 10 | daGal4 | 9.1 | ♂ | 3 | 40 | 11.8 | 15 | 39 | 10.9 | 15 | 40 | 10.0 | 10 | -8.2 | -33.3 | **0.0053** |
| Starvation | 10 | daGal4 | 9.1 | ♂ | 4 | 40 | 15.5 | 18 | 39 | 17.3 | 18 | 40 | 14.7 | 18 | -5.2 | 0.0 | **0.0465** |
| Starvation | 10 | daGal4 | 9.1 | ♂ | 5 | 40 | 16.0 | 18 | 39 | 17.6 | 22 | 39 | 15.1 | 18 | -5.4 | 0.0 | 0.1044 |
| Starvation | 10 | daGal4 | 3.2 | ♀ | 1 | 35 | 39.2 | 48 | 40 | 47.3 | 63 | 40 | 46.5 | 63 | 0.0 | 0.0 | |
| Starvation | 10 | daGal4 | 3.2 | ♀ | 2 | 40 | 28.8 | 33 | 40 | 25.5 | 38 | 40 | 30.1 | 43 | 4.4 | 13.2 | **0.0038** |
| Starvation | 10 | daGal4 | 3.2 | ♀ | 3 | 40 | 32.8 | 38 | 40 | 33.3 | 38 | 39 | 33.6 | 38 | 0.9 | 0.0 | 0.1294 |
| Starvation | 10 | daGal4 | 3.2 | ♀ | 4 | 40 | 24.7 | 33 | 40 | 24.8 | 33 | 40 | 29.4 | 38 | 18.9 | 15.2 | **0.0005** |
| Starvation | 10 | daGal4 | 3.2 | ♀ | 5 | 40 | 34.2 | 42 | 41 | 36.3 | 42 | 40 | 34.0 | 42 | -0.6 | 0.0 | **0.0463** |

### Table S2



| Stress | Age | Gal4 | UAS | Sex | Rep[1] | UAS | | | Gal4 | | | Gal4 + UAS | | | % Extension | | *P* |
|---|---|---|---|---|---|---|---|---|---|---|---|---|---|---|---|---|---|
| | | | | | | n | Mean | Max[2] | n | Mean | Max[2] | n | Mean | Max[2] | Mean | Max[2] | |
| Starvation | 10 | daGal4 | 3.2 | ♀ | 6 | 40 | 33.9 | 42 | 40 | 35.5 | 42 | 40 | 36.3 | 46 | 2.3 | 9.5 | **0.0197** |
| Starvation | 10 | daGal4 | 3.2 | ♂ | 1 | 40 | 20.9 | 29 | 40 | 24.9 | 33 | 40 | 28.8 | 38 | 15.8 | 15.2 | **0.0013** |
| Starvation | 10 | daGal4 | 3.2 | ♂ | 2 | 40 | 11.7 | 16 | 40 | 10.4 | 16 | 40 | 11.3 | 20 | 0.0 | 25.0 | |
| Starvation | 10 | daGal4 | 3.2 | ♂ | 3 | 40 | 19.6 | 26 | 38 | 16.5 | 18 | 40 | 19.1 | 26 | 0.0 | 0.0 | |
| Starvation | 10 | daGal4 | 3.2 | ♂ | 4 | 40 | 13.1 | 20 | 40 | 10.7 | 16 | 40 | 14.1 | 25 | 8.0 | 25.0 | **0.0015** |
| Starvation | 10 | daGal4 | 3.2 | ♂ | 5 | 40 | 18.7 | 22 | 39 | 17.3 | 18 | 39 | 19.7 | 26 | 5.6 | 18.2 | 0.0764 |
| Starvation | 10 | daGal4 | 3.2 | ♂ | 6 | 40 | 19.1 | 22 | 39 | 17.6 | 22 | 40 | 21.4 | 34 | 12.0 | 54.5 | **0.0322** |
| Starvation | 10 | daGal4 | 10.1 | ♀ | 1 | 36 | 26.9 | 47 | 34 | 25.1 | 33 | 40 | 31.5 | 47 | 16.9 | 0.0 | **0.0005** |
| Starvation | 10 | daGal4 | 10.1 | ♀ | 2 | 40 | 33.6 | 40 | 39 | 25.6 | 40 | 40 | 25.6 | 40 | 0.0 | 0.0 | |
| Starvation | 10 | daGal4 | 10.1 | ♀ | 3 | 44 | 35.2 | 42 | 40 | 33.3 | 38 | 40 | 34.7 | 42 | 0.0 | 0.0 | |
| Starvation | 10 | daGal4 | 10.1 | ♀ | 4 | 39 | 28.2 | 40 | 40 | 26.6 | 40 | 32 | 22.6 | 40 | -15.2 | 0.0 | **0.0049** |
| Starvation | 10 | daGal4 | 10.1 | ♀ | 5 | 40 | 35.1 | 42 | 41 | 36.3 | 42 | 40 | 36.3 | 42 | 0.0 | 0.0 | |
| Starvation | 10 | daGal4 | 10.1 | ♀ | 6 | 40 | 36.0 | 42 | 40 | 35.5 | 42 | 40 | 36.9 | 42 | 2.5 | 0.0 | 0.1098 |
| Starvation | 10 | daGal4 | 10.1 | ♂ | 1 | 36 | 19.8 | 22 | 36 | 21.6 | 22 | 39 | 24.6 | 33 | 13.9 | 50.0 | **0.0004** |
| Starvation | 10 | daGal4 | 10.1 | ♂ | 2 | 40 | 11.6 | 15 | 39 | 11.2 | 15 | 40 | 13.7 | 21 | 17.4 | 40.0 | **0.0014** |
| Starvation | 10 | daGal4 | 10.1 | ♂ | 3 | 40 | 19.1 | 22 | 38 | 16.5 | 18 | 43 | 19.9 | 26 | 4.0 | 18.2 | 0.1526 |
| Starvation | 10 | daGal4 | 10.1 | ♂ | 4 | 40 | 13.3 | 21 | 39 | 10.9 | 15 | 39 | 10.3 | 15 | -5.9 | 0.0 | 0.0783 |
| Starvation | 10 | daGal4 | 10.1 | ♂ | 5 | 40 | 18.7 | 26 | 39 | 17.3 | 18 | 42 | 20.2 | 30 | 8.0 | 15.4 | 0.0615 |
| Starvation | 10 | daGal4 | 10.1 | ♂ | 6 | 37 | 18.4 | 22 | 39 | 17.6 | 22 | 40 | 20.3 | 26 | 10.1 | 18.2 | **0.0170** |
| Starvation | 10 | daGal4 | 3.1 | ♀ | 1 | 37 | 32.7 | 46 | 33 | 34.7 | 46 | 40 | 35.9 | 54 | 3.4 | 17.4 | 0.0574 |
| Starvation | 10 | daGal4 | 3.1 | ♀ | 2 | 40 | 30.5 | 38 | 40 | 33.3 | 38 | 40 | 31.3 | 38 | 0.0 | 0.0 | |
| Starvation | 10 | daGal4 | 3.1 | ♀ | 3 | 40 | 31.4 | 38 | 41 | 36.3 | 42 | 40 | 34.4 | 42 | 0.0 | 0.0 | |
| Starvation | 10 | daGal4 | 3.1 | ♀ | 4 | 40 | 34.3 | 46 | 40 | 35.5 | 42 | 39 | 33.4 | 42 | -2.7 | 0.0 | **0.0087** |
| Starvation | 10 | daGal4 | 3.1 | ♂ | 1 | 40 | 18.6 | 26 | 40 | 18.5 | 30 | 40 | 21.4 | 30 | 15.1 | 0.0 | **0.0082** |
| Starvation | 10 | daGal4 | 3.1 | ♂ | 2 | 41 | 18.1 | 22 | 38 | 16.5 | 18 | 40 | 18.5 | 22 | 2.2 | 0.0 | 0.2871 |
| Starvation | 10 | daGal4 | 3.1 | ♂ | 3 | 40 | 16.5 | 22 | 39 | 17.3 | 18 | 41 | 18.3 | 26 | 5.8 | 18.2 | **0.0031** |
| Starvation | 10 | daGal4 | 3.1 | ♂ | 4 | 40 | 16.6 | 22 | 39 | 17.6 | 22 | 38 | 16.7 | 18 | 0.0 | -18.2 | |
| Starvation | 10 | daGal4 | 4.4 | ♀ | 1 | 40 | 30.5 | 40 | 39 | 25.6 | 40 | 36 | 24.9 | 40 | -2.5 | 0.0 | **0.0005** |
| Starvation | 10 | daGal4 | 4.4 | ♀ | 2 | 36 | 53.6 | 59 | 38 | 51.1 | 72 | 39 | 54.5 | 72 | 1.6 | 0.0 | 0.1343 |
| Starvation | 10 | daGal4 | 4.4 | ♀ | 3 | 40 | 34.1 | 38 | 40 | 33.3 | 38 | 41 | 32.1 | 42 | -3.5 | 10.5 | 0.0820 |
| Starvation | 10 | daGal4 | 4.4 | ♀ | 4 | 41 | 32.9 | 40 | 40 | 26.6 | 40 | 40 | 25.4 | 40 | -4.5 | 0.0 | 0.3065 |
| Starvation | 10 | daGal4 | 4.4 | ♀ | 5 | 40 | 34.7 | 42 | 41 | 36.3 | 42 | 31 | 35.8 | 46 | 0.0 | 9.5 | |
| Starvation | 10 | daGal4 | 4.4 | ♀ | 6 | 40 | 36.5 | 42 | 40 | 35.5 | 42 | 37 | 35.0 | 42 | -1.5 | 0.0 | 0.2074 |
| Starvation | 10 | daGal4 | 4.4 | ♂ | 1 | 40 | 11.6 | 15 | 39 | 11.2 | 15 | 42 | 10.4 | 15 | -7.4 | 0.0 | **0.0040** |
| Starvation | 10 | daGal4 | 4.4 | ♂ | 2 | 37 | 27.2 | 34 | 40 | 23.6 | 34 | 40 | 30.7 | 47 | 12.8 | 38.2 | **0.0286** |
| Starvation | 10 | daGal4 | 4.4 | ♂ | 3 | 40 | 16.7 | 18 | 38 | 16.5 | 18 | 39 | 17.2 | 22 | 2.9 | 22.2 | 0.1972 |
| Starvation | 10 | daGal4 | 4.4 | ♂ | 4 | 40 | 12.4 | 15 | 39 | 10.9 | 15 | 40 | 13.2 | 21 | 6.5 | 40.0 | **0.0004** |
| Starvation | 10 | daGal4 | 4.4 | ♂ | 5 | 40 | 17.1 | 22 | 39 | 17.3 | 18 | 39 | 17.3 | 22 | 0.0 | 0.0 | |
| Starvation | 10 | daGal4 | 4.4 | ♂ | 6 | 41 | 16.0 | 18 | 39 | 17.6 | 22 | 39 | 18.9 | 26 | 7.6 | 18.2 | **0.0228** |
| Starvation | 10 | DJ694 | 2.1 | ♀ | 1 | 40 | 49.5 | 63 | 40 | 49.0 | 58 | 40 | 47.5 | 58 | -3.1 | 0.0 | **0.0256** |
| Starvation | 10 | DJ694 | 2.1 | ♀ | 2 | 40 | 23.9 | 30 | 40 | 22.4 | 26 | 40 | 20.5 | 22 | -8.5 | -15.4 | **0.0009** |
| Starvation | 10 | DJ694 | 2.1 | ♀ | 3 | 40 | 23.8 | 30 | 41 | 24.7 | 30 | 40 | 22.9 | 26 | -3.8 | -13.3 | **0.0008** |
| Starvation | 10 | DJ694 | 2.1 | ♀ | 4 | 40 | 23.5 | 30 | 40 | 26.0 | 30 | 49 | 24.2 | 30 | 0.0 | 0.0 | |
| Starvation | 10 | DJ694 | 2.1 | ♂ | 1 | 39 | 27.9 | 38 | 38 | 26.9 | 33 | 39 | 27.3 | 33 | 0.0 | 0.0 | |
| Starvation | 10 | DJ694 | 2.1 | ♂ | 2 | 40 | 15.3 | 18 | 40 | 11.8 | 14 | 40 | 12.1 | 14 | 0.0 | 0.0 | |
| Starvation | 10 | DJ694 | 2.1 | ♂ | 3 | 40 | 14.3 | 18 | 40 | 15.7 | 18 | 40 | 13.3 | 18 | -7.0 | 0.0 | **0.0199** |
| Starvation | 10 | DJ694 | 2.1 | ♂ | 4 | 40 | 14.5 | 18 | 40 | 12.9 | 14 | 40 | 13.6 | 18 | 0.0 | 0.0 | |
| Starvation | 10 | DJ694 | 4.3 | ♀ | 1 | 40 | 48.6 | 59 | 38 | 50.3 | 72 | 40 | 52.8 | 72 | 4.9 | 0.0 | 0.4071 |
| Starvation | 10 | DJ694 | 4.3 | ♀ | 2 | 40 | 29.6 | 38 | 40 | 20.6 | 33 | 40 | 25.1 | 33 | 0.0 | 0.0 | |

### Table S2



| Stress | Age | Gal4 | UAS | Sex | Rep[1] | UAS | | | Gal4 | | | Gal4 + UAS | | | % Extension | | *P* |
|---|---|---|---|---|---|---|---|---|---|---|---|---|---|---|---|---|---|
| | | | | | | n | Mean | Max[2] | n | Mean | Max[2] | n | Mean | Max[2] | Mean | Max[2] | |
| Starvation | 10 | DJ694 | 4.3 | ♀ | 3 | 38 | 24.4 | 30 | 40 | 22.4 | 26 | 40 | 20.0 | 22 | -10.7 | -15.4 | **<0.0001** |
| Starvation | 10 | DJ694 | 4.3 | ♀ | 4 | 40 | 32.6 | 43 | 40 | 21.6 | 33 | 40 | 23.1 | 33 | 0.0 | 0.0 | |
| Starvation | 10 | DJ694 | 4.3 | ♀ | 5 | 40 | 25.0 | 30 | 41 | 24.7 | 30 | 30 | 21.3 | 22 | -13.7 | -26.7 | **<0.0001** |
| Starvation | 10 | DJ694 | 4.3 | ♀ | 6 | 40 | 26.7 | 34 | 40 | 26.0 | 30 | 40 | 22.5 | 26 | -13.5 | -13.3 | **<0.0001** |
| Starvation | 10 | DJ694 | 4.3 | ♂ | 1 | 37 | 26.6 | 34 | 40 | 25.9 | 34 | 40 | 25.6 | 34 | -1.1 | 0.0 | 0.4436 |
| Starvation | 10 | DJ694 | 4.3 | ♂ | 2 | 40 | 11.6 | 16 | 40 | 11.0 | 16 | 40 | 10.0 | 20 | -8.7 | 25.0 | 0.0795 |
| Starvation | 10 | DJ694 | 4.3 | ♂ | 3 | 40 | 16.4 | 18 | 40 | 11.8 | 14 | 40 | 13.7 | 14 | 0.0 | 0.0 | |
| Starvation | 10 | DJ694 | 4.3 | ♂ | 4 | 40 | 10.5 | 16 | 40 | 9.4 | 16 | 40 | 9.5 | 16 | 0.0 | 0.0 | |
| Starvation | 10 | DJ694 | 4.3 | ♂ | 5 | 40 | 16.1 | 18 | 40 | 15.7 | 18 | 40 | 14.1 | 18 | -10.2 | 0.0 | **0.0003** |
| Starvation | 10 | DJ694 | 4.3 | ♂ | 6 | 40 | 15.0 | 18 | 40 | 12.9 | 14 | 40 | 13.4 | 14 | 0.0 | 0.0 | |
| Starvation | 10 | DJ694 | 9.1 | ♀ | 1 | 39 | 32.7 | 40 | 39 | 26.3 | 40 | 40 | 24.8 | 40 | -5.8 | 0.0 | 0.1103 |
| Starvation | 10 | DJ694 | 9.1 | ♀ | 2 | 39 | 26.3 | 30 | 40 | 22.4 | 26 | 40 | 19.9 | 22 | -11.2 | -15.4 | **<0.0001** |
| Starvation | 10 | DJ694 | 9.1 | ♀ | 3 | 40 | 31.3 | 40 | 40 | 23.0 | 27 | 41 | 24.0 | 40 | 0.0 | 0.0 | |
| Starvation | 10 | DJ694 | 9.1 | ♀ | 4 | 40 | 27.0 | 34 | 41 | 24.7 | 30 | 40 | 20.6 | 22 | -16.7 | -26.7 | **<0.0001** |
| Starvation | 10 | DJ694 | 9.1 | ♀ | 5 | 40 | 27.6 | 30 | 40 | 26.0 | 30 | 40 | 19.3 | 26 | -25.8 | -13.3 | **<0.0001** |
| Starvation | 10 | DJ694 | 9.1 | ♂ | 1 | 39 | 11.3 | 15 | 38 | 10.0 | 10 | 39 | 10.0 | 10 | 0.0 | 0.0 | |
| Starvation | 10 | DJ694 | 9.1 | ♂ | 2 | 40 | 16.2 | 18 | 40 | 11.8 | 14 | 40 | 10.9 | 14 | -7.6 | 0.0 | **0.0345** |
| Starvation | 10 | DJ694 | 9.1 | ♂ | 3 | 40 | 11.8 | 15 | 40 | 10.6 | 15 | 40 | 10.1 | 10 | -4.7 | -33.3 | **0.0004** |
| Starvation | 10 | DJ694 | 9.1 | ♂ | 4 | 40 | 15.9 | 18 | 40 | 15.7 | 18 | 40 | 13.1 | 14 | -16.6 | -22.2 | **<0.0001** |
| Starvation | 10 | DJ694 | 9.1 | ♂ | 5 | 40 | 16.7 | 18 | 40 | 12.9 | 14 | 40 | 12.1 | 14 | -6.2 | 0.0 | 0.0805 |
| Starvation | 10 | DJ694 | 3.2 | ♀ | 1 | 35 | 39.2 | 48 | 38 | 44.4 | 53 | 40 | 47.0 | 53 | 5.9 | 0.0 | 0.3138 |
| Starvation | 10 | DJ694 | 3.2 | ♀ | 2 | 40 | 28.8 | 33 | 40 | 20.6 | 33 | 40 | 25.9 | 33 | 0.0 | 0.0 | |
| Starvation | 10 | DJ694 | 3.2 | ♀ | 3 | 40 | 26.4 | 34 | 40 | 22.4 | 26 | 40 | 23.6 | 34 | 0.0 | 0.0 | |
| Starvation | 10 | DJ694 | 3.2 | ♀ | 4 | 40 | 24.7 | 33 | 40 | 21.6 | 33 | 40 | 25.4 | 33 | 2.8 | 0.0 | **0.0147** |
| Starvation | 10 | DJ694 | 3.2 | ♀ | 5 | 40 | 26.1 | 30 | 41 | 24.7 | 30 | 40 | 24.7 | 30 | 0.0 | 0.0 | |
| Starvation | 10 | DJ694 | 3.2 | ♀ | 6 | 40 | 27.6 | 30 | 40 | 26.0 | 30 | 40 | 25.9 | 38 | -0.4 | 26.7 | 0.1205 |
| Starvation | 10 | DJ694 | 3.2 | ♂ | 1 | 40 | 20.9 | 29 | 36 | 19.6 | 29 | 40 | 24.1 | 38 | 15.3 | 31.0 | **0.0015** |
| Starvation | 10 | DJ694 | 3.2 | ♂ | 2 | 40 | 11.7 | 16 | 40 | 11.0 | 16 | 40 | 11.1 | 25 | 0.0 | 56.3 | |
| Starvation | 10 | DJ694 | 3.2 | ♂ | 3 | 40 | 18.5 | 22 | 40 | 11.8 | 14 | 40 | 16.5 | 18 | 0.0 | 0.0 | |
| Starvation | 10 | DJ694 | 3.2 | ♂ | 4 | 40 | 13.1 | 20 | 40 | 9.4 | 16 | 40 | 10.8 | 20 | 0.0 | 0.0 | |
| Starvation | 10 | DJ694 | 3.2 | ♂ | 5 | 40 | 18.1 | 22 | 40 | 15.7 | 18 | 40 | 17.0 | 22 | 0.0 | 0.0 | |
| Starvation | 10 | DJ694 | 3.2 | ♂ | 6 | 40 | 19.3 | 22 | 40 | 12.9 | 14 | 40 | 16.5 | 22 | 0.0 | 0.0 | |
| Starvation | 10 | DJ694 | 10.1 | ♀ | 1 | 36 | 26.9 | 47 | 32 | 26.3 | 47 | 40 | 29.3 | 47 | 8.7 | 0.0 | 0.0586 |
| Starvation | 10 | DJ694 | 10.1 | ♀ | 2 | 40 | 33.6 | 40 | 39 | 26.3 | 40 | 40 | 23.4 | 27 | -11.1 | -32.5 | **0.0032** |
| Starvation | 10 | DJ694 | 10.1 | ♀ | 3 | 40 | 24.5 | 30 | 40 | 22.4 | 26 | 40 | 22.3 | 26 | -0.4 | 0.0 | **0.0017** |
| Starvation | 10 | DJ694 | 10.1 | ♀ | 4 | 39 | 28.2 | 40 | 40 | 23.0 | 27 | 40 | 21.5 | 27 | -6.6 | 0.0 | 0.0272 |
| Starvation | 10 | DJ694 | 10.1 | ♀ | 5 | 40 | 27.6 | 38 | 41 | 24.7 | 30 | 39 | 24.6 | 30 | -0.7 | 0.0 | 0.7832 |
| Starvation | 10 | DJ694 | 10.1 | ♀ | 6 | 40 | 27.1 | 34 | 40 | 26.0 | 30 | 40 | 24.0 | 30 | -7.7 | 0.0 | **0.0002** |
| Starvation | 10 | DJ694 | 10.1 | ♂ | 1 | 36 | 19.8 | 22 | 32 | 20.4 | 22 | 38 | 22.7 | 29 | 11.6 | 31.8 | **0.0015** |
| Starvation | 10 | DJ694 | 10.1 | ♂ | 2 | 40 | 11.6 | 15 | 38 | 10.0 | 10 | 40 | 10.0 | 10 | 0.0 | 0.0 | |
| Starvation | 10 | DJ694 | 10.1 | ♂ | 3 | 40 | 17.5 | 22 | 40 | 11.8 | 14 | 40 | 15.1 | 18 | 0.0 | 0.0 | |
| Starvation | 10 | DJ694 | 10.1 | ♂ | 4 | 40 | 13.3 | 21 | 40 | 10.6 | 15 | 40 | 11.8 | 21 | 0.0 | 0.0 | |
| Starvation | 10 | DJ694 | 10.1 | ♂ | 5 | 40 | 18.8 | 22 | 40 | 15.7 | 18 | 40 | 17.1 | 18 | 0.0 | 0.0 | |
| Starvation | 10 | DJ694 | 10.1 | ♂ | 6 | 40 | 18.5 | 22 | 40 | 12.9 | 14 | 40 | 17.5 | 22 | 0.0 | 0.0 | |
| Starvation | 10 | DJ694 | 3.1 | ♀ | 1 | 37 | 32.7 | 46 | 40 | 41.1 | 54 | 39 | 34.2 | 50 | 0.0 | 0.0 | |
| Starvation | 10 | DJ694 | 3.1 | ♀ | 2 | 40 | 21.7 | 26 | 40 | 22.4 | 26 | 40 | 22.8 | 30 | 1.8 | 15.4 | 0.0746 |
| Starvation | 10 | DJ694 | 3.1 | ♀ | 3 | 40 | 23.6 | 26 | 41 | 24.7 | 30 | 40 | 22.5 | 30 | -4.7 | 0.0 | **0.0004** |
| Starvation | 10 | DJ694 | 3.1 | ♀ | 4 | 40 | 24.2 | 30 | 40 | 26.0 | 30 | 40 | 24.8 | 34 | 0.0 | 13.3 | |
| Starvation | 10 | DJ694 | 3.1 | ♂ | 1 | 40 | 18.6 | 26 | 40 | 18.2 | 26 | 40 | 16.0 | 22 | -12.2 | -15.4 | **0.0029** |

### Table S2



| Stress | Age | Gal4 | UAS | Sex | Rep[1] | UAS | | | Gal4 | | | Gal4 + UAS | | | % Extension | | *P* |
|---|---|---|---|---|---|---|---|---|---|---|---|---|---|---|---|---|---|
| | | | | | | n | Mean | Max[2] | n | Mean | Max[2] | n | Mean | Max[2] | Mean | Max[2] | |
| Starvation | 10 | DJ694 | 3.1 | ♂ | 2 | 40 | 17.5 | 18 | 40 | 11.8 | 14 | 20 | 13.0 | 14 | 0.0 | 0.0 | |
| Starvation | 10 | DJ694 | 3.1 | ♂ | 3 | 40 | 17.6 | 18 | 40 | 15.7 | 18 | 40 | 15.7 | 18 | 0.0 | 0.0 | |
| Starvation | 10 | DJ694 | 3.1 | ♂ | 4 | 40 | 16.0 | 18 | 40 | 12.9 | 14 | 40 | 16.4 | 18 | 2.5 | 0.0 | 0.5390 |
| Starvation | 10 | DJ694 | 4.4 | ♀ | 1 | 40 | 29.8 | 35 | 40 | 24.8 | 35 | 40 | 21.8 | 30 | -12.1 | -14.3 | **0.0025** |
| Starvation | 10 | DJ694 | 4.4 | ♀ | 2 | 40 | 30.5 | 40 | 39 | 26.3 | 40 | 39 | 22.4 | 27 | -15.0 | -32.5 | **0.0002** |
| Starvation | 10 | DJ694 | 4.4 | ♀ | 3 | 36 | 53.6 | 59 | 38 | 50.3 | 72 | 40 | 54.8 | 72 | 2.1 | 0.0 | **0.0406** |
| Starvation | 10 | DJ694 | 4.4 | ♀ | 4 | 40 | 23.9 | 26 | 40 | 22.4 | 26 | 40 | 23.1 | 30 | 0.0 | 15.4 | |
| Starvation | 10 | DJ694 | 4.4 | ♀ | 5 | 41 | 32.9 | 40 | 40 | 23.0 | 27 | 40 | 24.3 | 40 | 0.0 | 0.0 | |
| Starvation | 10 | DJ694 | 4.4 | ♀ | 6 | 40 | 25.9 | 30 | 41 | 24.7 | 30 | 40 | 25.5 | 30 | 0.0 | 0.0 | |
| Starvation | 10 | DJ694 | 4.4 | ♀ | 7 | 40 | 26.8 | 30 | 40 | 26.0 | 30 | 40 | 24.8 | 30 | -4.6 | 0.0 | 0.0578 |
| Starvation | 10 | DJ694 | 4.4 | ♂ | 1 | 40 | 19.5 | 25 | 40 | 19.8 | 25 | 40 | 18.6 | 25 | -4.5 | 0.0 | 0.1191 |
| Starvation | 10 | DJ694 | 4.4 | ♂ | 2 | 40 | 11.6 | 15 | 38 | 10.0 | 10 | 39 | 10.1 | 10 | 0.0 | 0.0 | |
| Starvation | 10 | DJ694 | 4.4 | ♂ | 3 | 37 | 27.2 | 34 | 40 | 25.9 | 34 | 38 | 24.4 | 34 | -5.8 | 0.0 | 0.0516 |
| Starvation | 10 | DJ694 | 4.4 | ♂ | 4 | 40 | 15.6 | 18 | 40 | 11.8 | 14 | 40 | 15.5 | 18 | 0.0 | 0.0 | |
| Starvation | 10 | DJ694 | 4.4 | ♂ | 5 | 40 | 12.4 | 15 | 40 | 10.6 | 15 | 40 | 10.3 | 15 | -3.5 | 0.0 | 0.2382 |
| Starvation | 10 | DJ694 | 4.4 | ♂ | 6 | 30 | 16.5 | 18 | 40 | 15.7 | 18 | 40 | 16.6 | 22 | 0.4 | 22.2 | 0.0956 |
| Starvation | 10 | DJ694 | 4.4 | ♂ | 7 | 40 | 16.2 | 18 | 40 | 12.9 | 14 | 40 | 14.3 | 18 | 0.0 | 0.0 | |
| Starvation | 10 | D42 | 2.1 | ♀ | 1 | 40 | 49.5 | 63 | 40 | 46.6 | 58 | 40 | 49.8 | 58 | 0.6 | 0.0 | 0.1613 |
| Starvation | 10 | D42 | 2.1 | ♀ | 2 | 40 | 25.8 | 32 | 40 | 25.1 | 32 | 39 | 25.3 | 28 | 0.0 | -12.5 | |
| Starvation | 10 | D42 | 2.1 | ♀ | 3 | 40 | 26.1 | 32 | 40 | 29.4 | 36 | 40 | 26.8 | 32 | 0.0 | 0.0 | |
| Starvation | 10 | D42 | 2.1 | ♀ | 4 | 40 | 27.5 | 32 | 40 | 29.8 | 36 | 40 | 29.4 | 32 | 0.0 | 0.0 | |
| Starvation | 10 | D42 | 2.1 | ♂ | 1 | 39 | 27.9 | 38 | 36 | 30.5 | 38 | 40 | 27.8 | 33 | -0.7 | -13.2 | **0.0013** |
| Starvation | 10 | D42 | 2.1 | ♂ | 2 | 37 | 15.2 | 16 | 40 | 17.4 | 20 | 39 | 14.7 | 16 | -3.3 | 0.0 | 0.4107 |
| Starvation | 10 | D42 | 2.1 | ♂ | 3 | 40 | 16.5 | 24 | 39 | 15.7 | 20 | 40 | 14.8 | 16 | -5.9 | -20.0 | **0.0023** |
| Starvation | 10 | D42 | 2.1 | ♂ | 4 | 40 | 15.6 | 16 | 40 | 16.9 | 20 | 40 | 15.8 | 16 | 0.0 | 0.0 | |
| Starvation | 10 | D42 | 4.3 | ♀ | 1 | 40 | 48.6 | 59 | 39 | 51.3 | 72 | 39 | 54.8 | 72 | 6.8 | 0.0 | **0.0457** |
| Starvation | 10 | D42 | 4.3 | ♀ | 2 | 40 | 29.6 | 38 | 40 | 24.9 | 43 | 40 | 23.0 | 33 | -7.9 | -13.2 | 0.1441 |
| Starvation | 10 | D42 | 4.3 | ♀ | 3 | 40 | 27.1 | 32 | 40 | 25.1 | 32 | 38 | 25.7 | 32 | 0.0 | 0.0 | |
| Starvation | 10 | D42 | 4.3 | ♀ | 4 | 40 | 32.6 | 43 | 40 | 23.5 | 33 | 40 | 23.7 | 33 | 0.0 | 0.0 | |
| Starvation | 10 | D42 | 4.3 | ♀ | 5 | 40 | 28.8 | 32 | 40 | 29.4 | 36 | 40 | 29.2 | 32 | 0.0 | 0.0 | |
| Starvation | 10 | D42 | 4.3 | ♀ | 6 | 40 | 29.9 | 32 | 40 | 29.8 | 36 | 40 | 27.0 | 32 | -9.4 | 0.0 | **<0.0001** |
| Starvation | 10 | D42 | 4.3 | ♂ | 1 | 37 | 26.6 | 34 | 36 | 25.5 | 34 | 40 | 24.1 | 34 | -5.5 | 0.0 | **0.0405** |
| Starvation | 10 | D42 | 4.3 | ♂ | 2 | 40 | 11.6 | 16 | 40 | 9.9 | 16 | 40 | 7.9 | 16 | -19.5 | 0.0 | **0.0042** |
| Starvation | 10 | D42 | 4.3 | ♂ | 3 | 40 | 14.5 | 16 | 40 | 17.4 | 20 | 39 | 14.3 | 16 | -1.8 | 0.0 | 0.6028 |
| Starvation | 10 | D42 | 4.3 | ♂ | 4 | 40 | 10.5 | 16 | 40 | 10.6 | 16 | 40 | 9.6 | 16 | -8.6 | 0.0 | 0.2443 |
| Starvation | 10 | D42 | 4.3 | ♂ | 5 | 39 | 15.2 | 16 | 39 | 15.7 | 20 | 40 | 13.6 | 16 | -10.4 | 0.0 | **0.0003** |
| Starvation | 10 | D42 | 4.3 | ♂ | 6 | 40 | 14.8 | 16 | 40 | 16.9 | 20 | 40 | 13.7 | 20 | -7.8 | 0.0 | 0.1176 |
| Starvation | 10 | D42 | 9.1 | ♀ | 1 | 39 | 32.7 | 40 | 40 | 36.1 | 40 | 40 | 29.2 | 40 | -10.8 | 0.0 | **0.0299** |
| Starvation | 10 | D42 | 9.1 | ♀ | 2 | 40 | 27.6 | 36 | 40 | 25.1 | 32 | 40 | 23.2 | 28 | -7.6 | -12.5 | **0.0058** |
| Starvation | 10 | D42 | 9.1 | ♀ | 3 | 40 | 31.3 | 40 | 40 | 28.9 | 40 | 40 | 26.2 | 40 | -9.6 | 0.0 | **0.0033** |
| Starvation | 10 | D42 | 9.1 | ♀ | 4 | 40 | 29.5 | 36 | 40 | 29.4 | 36 | 40 | 24.3 | 28 | -17.3 | -22.2 | **<0.0001** |
| Starvation | 10 | D42 | 9.1 | ♀ | 5 | 40 | 30.4 | 36 | 40 | 29.8 | 36 | 38 | 25.3 | 32 | -15.2 | -11.1 | **<0.0001** |
| Starvation | 10 | D42 | 9.1 | ♂ | 1 | 39 | 11.3 | 15 | 40 | 10.6 | 15 | 40 | 10.0 | 10 | -5.9 | -33.3 | **0.0013** |
| Starvation | 10 | D42 | 9.1 | ♂ | 2 | 40 | 15.4 | 16 | 40 | 17.4 | 20 | 40 | 12.3 | 16 | -19.9 | 0.0 | **<0.0001** |
| Starvation | 10 | D42 | 9.1 | ♂ | 3 | 40 | 11.8 | 15 | 40 | 11.9 | 21 | 40 | 10.5 | 15 | -10.8 | 0.0 | **0.0100** |
| Starvation | 10 | D42 | 9.1 | ♂ | 4 | 40 | 14.8 | 16 | 39 | 15.7 | 20 | 38 | 13.5 | 16 | -8.5 | 0.0 | **0.0002** |
| Starvation | 10 | D42 | 9.1 | ♂ | 5 | 40 | 15.4 | 16 | 40 | 16.9 | 20 | 40 | 16.0 | 16 | 0.0 | 0.0 | |
| Starvation | 10 | D42 | 3.2 | ♀ | 1 | 35 | 39.2 | 48 | 40 | 44.5 | 63 | 40 | 44.1 | 58 | 0.0 | 0.0 | |
| Starvation | 10 | D42 | 3.2 | ♀ | 2 | 40 | 28.8 | 33 | 40 | 24.9 | 43 | 40 | 31.9 | 38 | 10.7 | 0.0 | **0.0053** |

| Stress | Age | Gal4 | UAS | Sex | Rep[1] | UAS | | | Gal4 | | | Gal4 + UAS | | | % Extension | | P |
|---|---|---|---|---|---|---|---|---|---|---|---|---|---|---|---|---|---|
| | | | | | | n | Mean | Max[2] | n | Mean | Max[2] | n | Mean | Max[2] | Mean | Max[2] | |
| Starvation | 10 | D42 | 3.2 | ♀ | 3 | 40 | 26.3 | 32 | 40 | 25.1 | 32 | 40 | 26.1 | 36 | 0.0 | 12.5 | |
| Starvation | 10 | D42 | 3.2 | ♀ | 4 | 40 | 24.7 | 33 | 40 | 23.5 | 33 | 40 | 30.1 | 38 | 21.8 | 15.2 | **0.0004** |
| Starvation | 10 | D42 | 3.2 | ♀ | 5 | 40 | 26.7 | 32 | 40 | 29.4 | 36 | 40 | 26.9 | 32 | 0.0 | 0.0 | |
| Starvation | 10 | D42 | 3.2 | ♀ | 6 | 40 | 30.6 | 36 | 40 | 29.8 | 36 | 40 | 28.4 | 32 | -4.7 | -11.1 | **0.0026** |
| Starvation | 10 | D42 | 3.2 | ♂ | 1 | 40 | 20.9 | 29 | 39 | 22.9 | 33 | 40 | 26.1 | 38 | 13.8 | 15.2 | **0.0487** |
| Starvation | 10 | D42 | 3.2 | ♂ | 2 | 40 | 11.7 | 16 | 40 | 9.9 | 16 | 40 | 12.4 | 16 | 5.8 | 0.0 | **0.0006** |
| Starvation | 10 | D42 | 3.2 | ♂ | 3 | 40 | 17.5 | 20 | 40 | 17.4 | 20 | 40 | 16.8 | 20 | -3.5 | 0.0 | 0.0887 |
| Starvation | 10 | D42 | 3.2 | ♂ | 4 | 40 | 13.1 | 20 | 40 | 10.6 | 16 | 40 | 11.7 | 16 | 0.0 | 0.0 | |
| Starvation | 10 | D42 | 3.2 | ♂ | 5 | 40 | 18.1 | 20 | 39 | 15.7 | 20 | 40 | 17.9 | 20 | 0.0 | 0.0 | |
| Starvation | 10 | D42 | 3.2 | ♂ | 6 | 39 | 19.6 | 24 | 40 | 16.9 | 20 | 40 | 20.2 | 24 | 3.1 | 0.0 | 0.2151 |
| Starvation | 10 | D42 | 10.1 | ♀ | 1 | 36 | 26.9 | 47 | 33 | 26.7 | 47 | 40 | 25.8 | 33 | -3.4 | -29.8 | 0.5199 |
| Starvation | 10 | D42 | 10.1 | ♀ | 2 | 40 | 33.6 | 40 | 40 | 36.1 | 40 | 40 | 27.4 | 40 | -18.3 | 0.0 | **0.0002** |
| Starvation | 10 | D42 | 10.1 | ♀ | 3 | 39 | 24.0 | 32 | 40 | 25.1 | 32 | 40 | 25.8 | 32 | 2.8 | 0.0 | **0.0406** |
| Starvation | 10 | D42 | 10.1 | ♀ | 4 | 39 | 28.2 | 40 | 40 | 28.9 | 40 | 40 | 23.0 | 27 | -18.6 | -32.5 | **<0.0001** |
| Starvation | 10 | D42 | 10.1 | ♀ | 5 | 40 | 28.9 | 36 | 40 | 29.4 | 36 | 39 | 25.8 | 32 | -10.6 | -11.1 | **0.0004** |
| Starvation | 10 | D42 | 10.1 | ♀ | 6 | 40 | 30.6 | 40 | 40 | 29.8 | 36 | 40 | 29.9 | 40 | 0.0 | 0.0 | |
| Starvation | 10 | D42 | 10.1 | ♂ | 1 | 36 | 19.8 | 22 | 39 | 21.0 | 22 | 39 | 24.5 | 33 | 16.5 | 50.0 | **<0.0001** |
| Starvation | 10 | D42 | 10.1 | ♂ | 2 | 40 | 11.6 | 15 | 40 | 10.6 | 15 | 40 | 11.3 | 15 | 0.0 | 0.0 | |
| Starvation | 10 | D42 | 10.1 | ♂ | 3 | 40 | 18.5 | 24 | 40 | 17.4 | 20 | 40 | 19.9 | 24 | 7.6 | 0.0 | 0.0559 |
| Starvation | 10 | D42 | 10.1 | ♂ | 4 | 40 | 13.3 | 21 | 40 | 11.9 | 21 | 40 | 15.6 | 21 | 17.5 | 0.0 | **0.0031** |
| Starvation | 10 | D42 | 10.1 | ♂ | 5 | 27 | 16.7 | 24 | 39 | 15.7 | 20 | 40 | 22.2 | 28 | 32.9 | 16.7 | **<0.0001** |
| Starvation | 10 | D42 | 10.1 | ♂ | 6 | 41 | 20.0 | 28 | 40 | 16.9 | 20 | 40 | 24.4 | 32 | 22.0 | 14.3 | **<0.0001** |
| Starvation | 10 | D42 | 3.1 | ♀ | 1 | 37 | 32.7 | 46 | 35 | 37.7 | 50 | 40 | 37.9 | 58 | 0.6 | 16.0 | **0.0082** |
| Starvation | 10 | D42 | 3.1 | ♀ | 2 | 39 | 22.8 | 28 | 40 | 25.1 | 32 | 38 | 22.5 | 28 | -1.1 | 0.0 | **0.0008** |
| Starvation | 10 | D42 | 3.1 | ♀ | 3 | 40 | 26.2 | 32 | 40 | 29.4 | 36 | 39 | 24.4 | 28 | -6.8 | -12.5 | **0.0323** |
| Starvation | 10 | D42 | 3.1 | ♀ | 4 | 40 | 25.8 | 32 | 40 | 29.8 | 36 | 40 | 28.1 | 32 | 0.0 | 0.0 | |
| Starvation | 10 | D42 | 3.1 | ♂ | 1 | 40 | 18.6 | 26 | 40 | 18.3 | 26 | 40 | 21.1 | 30 | 13.7 | 15.4 | **0.0003** |
| Starvation | 10 | D42 | 3.1 | ♂ | 2 | 40 | 16.5 | 20 | 40 | 17.4 | 20 | 40 | 15.4 | 16 | -6.5 | -20.0 | **0.0028** |
| Starvation | 10 | D42 | 3.1 | ♂ | 3 | 40 | 16.9 | 20 | 39 | 15.7 | 20 | 40 | 16.3 | 20 | 0.0 | 0.0 | |
| Starvation | 10 | D42 | 3.1 | ♂ | 4 | 40 | 16.0 | 20 | 40 | 16.9 | 20 | 40 | 18.2 | 20 | 7.9 | 0.0 | 0.0572 |
| Starvation | 10 | D42 | 4.4 | ♀ | 1 | 40 | 29.8 | 35 | 40 | 24.3 | 35 | 40 | 19.9 | 25 | -18.0 | -28.6 | **<0.0001** |
| Starvation | 10 | D42 | 4.4 | ♀ | 2 | 40 | 30.5 | 40 | 40 | 36.1 | 40 | 40 | 25.4 | 27 | -16.8 | -32.5 | **0.0006** |
| Starvation | 10 | D42 | 4.4 | ♀ | 3 | 36 | 53.6 | 59 | 39 | 51.3 | 72 | 40 | 55.5 | 72 | 3.5 | 0.0 | 0.0973 |
| Starvation | 10 | D42 | 4.4 | ♀ | 4 | 40 | 26.5 | 32 | 40 | 25.1 | 32 | 40 | 24.8 | 32 | -1.3 | 0.0 | 0.0686 |
| Starvation | 10 | D42 | 4.4 | ♀ | 5 | 41 | 32.9 | 40 | 40 | 28.9 | 40 | 39 | 23.7 | 40 | -18.2 | 0.0 | **<0.0001** |
| Starvation | 10 | D42 | 4.4 | ♀ | 6 | 40 | 28.5 | 36 | 40 | 29.4 | 36 | 40 | 27.8 | 32 | -2.5 | -11.1 | **0.0206** |
| Starvation | 10 | D42 | 4.4 | ♀ | 7 | 40 | 31.2 | 40 | 40 | 29.8 | 36 | 30 | 28.9 | 32 | -2.9 | -11.1 | **0.0065** |
| Starvation | 10 | D42 | 4.4 | ♂ | 1 | 40 | 19.5 | 25 | 40 | 19.9 | 25 | 40 | 19.8 | 25 | 0.0 | 0.0 | |
| Starvation | 10 | D42 | 4.4 | ♂ | 2 | 40 | 11.6 | 15 | 40 | 10.6 | 15 | 40 | 10.0 | 10 | -5.9 | -33.3 | **0.0217** |
| Starvation | 10 | D42 | 4.4 | ♂ | 3 | 37 | 27.2 | 34 | 36 | 25.5 | 34 | 39 | 27.8 | 34 | 2.3 | 0.0 | 0.0617 |
| Starvation | 10 | D42 | 4.4 | ♂ | 4 | 40 | 14.5 | 16 | 40 | 17.4 | 20 | 40 | 15.3 | 20 | 0.0 | 0.0 | |
| Starvation | 10 | D42 | 4.4 | ♂ | 5 | 40 | 12.4 | 15 | 40 | 11.9 | 21 | 39 | 10.9 | 15 | -8.6 | 0.0 | **0.0082** |
| Starvation | 10 | D42 | 4.4 | ♂ | 6 | 40 | 14.6 | 16 | 39 | 15.7 | 20 | 41 | 16.1 | 20 | 2.6 | 0.0 | **0.0007** |
| Starvation | 10 | D42 | 4.4 | ♂ | 7 | 40 | 15.7 | 16 | 40 | 16.9 | 20 | 40 | 16.4 | 20 | 0.0 | 0.0 | |

**Table S2: Stress resistance data and analysis of Hsp70 overexpression.** Mean: average survival in hours, n: number of flies in the replicate, P: highest Log-rank test value, 1: replicate number, 2: 95% mortality (maximum lifespan)
