## Supplementary material for "Expression of heat shock protein 70 is insufficient to extend *Drosophila melanogaster* longevity": Table S3



| Gal4 | UAS | Sex | Rep[1] | UAS | | | Gal4 | | | Gal4 + UAS | | | % Extension | | *P* |
|---|---|---|---|---|---|---|---|---|---|---|---|---|---|---|---|
| | | | | n | Mean | Max[2] | n | Mean | Max[2] | n | Mean | Max[2] | Mean | Max[2] | |
| daGal4 | 2.1 | ♀ | 1 | 115 | 55.4 | 78 | 110 | 66.3 | 95 | 120 | 66.7 | 81 | 0.6 | 0.0 | **0.0013** |
| daGal4 | 2.1 | ♀ | 2 | 114 | 66.2 | 94 | 95 | 70.8 | 90 | 90 | 75.7 | 90 | 6.9 | 0.0 | 0.7069 |
| daGal4 | 2.1 | ♀ | 3 | 112 | 58.9 | 81 | 119 | 71.5 | 88 | 117 | 62.1 | 74 | 0.0 | -8.6 | |
| daGal4 | 2.1 | ♀ | 4 | 118 | 60.2 | 87 | 113 | 70.6 | 87 | 118 | 65.9 | 80 | 0.0 | -8.0 | |
| daGal4 | 2.1 | ♀ | 5 | 90 | 63.1 | 76 | 102 | 69.3 | 87 | 103 | 65.9 | 80 | 0.0 | 0.0 | |
| daGal4 | 2.1 | ♀ | 6 | 111 | 58.8 | 72 | 109 | 71.9 | 86 | 111 | 64.3 | 79 | 0.0 | 0.0 | |
| daGal4 | 2.1 | ♂ | 1 | 117 | 49.4 | 75 | 113 | 50.6 | 75 | 116 | 45.6 | 51 | -7.7 | -32.0 | **<0.0001** |
| daGal4 | 2.1 | ♂ | 2 | 113 | 52.3 | 63 | 107 | 53.9 | 70 | 118 | 47.1 | 60 | -9.9 | -4.8 | **<0.0001** |
| daGal4 | 2.1 | ♂ | 3 | 113 | 56.0 | 67 | 118 | 64.2 | 81 | 117 | 50.5 | 57 | -9.7 | -14.9 | **<0.0001** |
| daGal4 | 2.1 | ♂ | 4 | 112 | 57.0 | 69 | 117 | 60.5 | 80 | 116 | 52.0 | 66 | -8.9 | -4.3 | **<0.0001** |
| daGal4 | 2.1 | ♂ | 5 | 102 | 55.5 | 69 | 100 | 60.5 | 73 | 117 | 52.9 | 66 | -4.7 | -4.3 | **0.0035** |
| daGal4 | 2.1 | ♂ | 6 | 117 | 54.0 | 68 | 107 | 59.7 | 72 | 112 | 54.5 | 68 | 0.0 | 0.0 | |
| daGal4 | 4.3 | ♀ | 1 | 121 | 53.9 | 66 | 102 | 71.8 | 94 | 122 | 66.0 | 85 | 0.0 | 0.0 | |
| daGal4 | 4.3 | ♀ | 2 | 111 | 54.2 | 69 | 107 | 70.3 | 97 | 100 | 68.0 | 87 | 0.0 | 0.0 | |
| daGal4 | 4.3 | ♀ | 3 | 112 | 54.8 | 70 | 119 | 71.5 | 88 | 118 | 56.7 | 74 | 0.0 | 0.0 | |
| daGal4 | 4.3 | ♀ | 4 | 119 | 51.0 | 66 | 113 | 70.6 | 87 | 117 | 62.5 | 76 | 0.0 | 0.0 | |
| daGal4 | 4.3 | ♀ | 5 | 119 | 48.2 | 62 | 102 | 69.3 | 87 | 109 | 54.4 | 66 | 0.0 | 0.0 | |
| daGal4 | 4.3 | ♀ | 6 | 122 | 51.7 | 68 | 109 | 71.9 | 86 | 117 | 61.2 | 79 | 0.0 | 0.0 | |
| daGal4 | 4.3 | ♂ | 1 | 120 | 48.0 | 59 | 114 | 59.1 | 85 | 118 | 41.9 | 46 | -12.7 | -22.0 | **<0.0001** |
| daGal4 | 4.3 | ♂ | 2 | 120 | 48.2 | 63 | 115 | 57.7 | 76 | 115 | 41.5 | 45 | -13.9 | -28.6 | **<0.0001** |
| daGal4 | 4.3 | ♂ | 3 | 118 | 46.8 | 53 | 118 | 64.2 | 81 | 119 | 40.3 | 49 | -14.0 | -7.5 | **<0.0001** |
| daGal4 | 4.3 | ♂ | 4 | 113 | 43.0 | 52 | 117 | 60.5 | 80 | 112 | 42.0 | 52 | -2.3 | 0.0 | 0.6680 |
| daGal4 | 4.3 | ♂ | 5 | 118 | 48.1 | 56 | 100 | 60.5 | 73 | 107 | 38.7 | 52 | -19.5 | -7.1 | **<0.0001** |
| daGal4 | 4.3 | ♂ | 6 | 120 | 46.2 | 55 | 107 | 59.7 | 72 | 119 | 38.0 | 47 | -17.7 | -14.5 | **<0.0001** |
| daGal4 | 9.1 | ♀ | 1 | 119 | 54.6 | 74 | 119 | 71.5 | 88 | 115 | 53.9 | 67 | -1.2 | -9.5 | 0.9776 |
| daGal4 | 9.1 | ♀ | 2 | 118 | 56.2 | 73 | 113 | 70.6 | 87 | 120 | 55.3 | 69 | -1.6 | -5.5 | 0.5707 |
| daGal4 | 9.1 | ♀ | 3 | 126 | 52.0 | 66 | 102 | 69.3 | 87 | 120 | 52.5 | 66 | 0.0 | 0.0 | |
| daGal4 | 9.1 | ♀ | 4 | 118 | 54.8 | 75 | 109 | 71.9 | 86 | 116 | 57.0 | 72 | 0.0 | -4.0 | |
| daGal4 | 9.1 | ♂ | 1 | 116 | 47.0 | 60 | 118 | 64.2 | 81 | 116 | 46.4 | 53 | -1.2 | -11.7 | 0.2053 |
| daGal4 | 9.1 | ♂ | 2 | 117 | 48.6 | 59 | 117 | 60.5 | 80 | 120 | 44.9 | 56 | -7.8 | -5.1 | **<0.0001** |
| daGal4 | 9.1 | ♂ | 3 | 116 | 44.6 | 56 | 100 | 60.5 | 73 | 126 | 41.5 | 52 | -7.1 | -7.1 | **0.0019** |
| daGal4 | 9.1 | ♂ | 4 | 115 | 50.5 | 61 | 107 | 59.7 | 72 | 116 | 44.1 | 55 | -12.6 | -9.8 | **<0.0001** |
| daGal4 | 3.2 | ♀ | 1 | 118 | 70.5 | 93 | 116 | 55.8 | 79 | 117 | 58.0 | 62 | 0.0 | -21.5 | |
| daGal4 | 3.2 | ♀ | 2 | 117 | 74.2 | 101 | 116 | 61.7 | 88 | 115 | 58.8 | 66 | -4.7 | -25.0 | **<0.0001** |
| daGal4 | 3.2 | ♀ | 3 | 113 | 69.5 | 81 | 119 | 71.5 | 88 | 119 | 50.5 | 57 | -27.2 | -29.6 | **<0.0001** |
| daGal4 | 3.2 | ♀ | 4 | 118 | 74.0 | 87 | 113 | 70.6 | 87 | 122 | 51.3 | 56 | -27.4 | -35.6 | **<0.0001** |
| daGal4 | 3.2 | ♀ | 5 | 110 | 67.2 | 87 | 102 | 69.3 | 87 | 115 | 52.4 | 56 | -22.0 | -35.6 | **<0.0001** |
| daGal4 | 3.2 | ♀ | 6 | 115 | 71.0 | 86 | 109 | 71.9 | 86 | 118 | 50.6 | 55 | -28.7 | -36.0 | **<0.0001** |
| daGal4 | 3.2 | ♂ | 1 | 116 | 62.4 | 84 | 118 | 48.1 | 62 | 118 | 42.6 | 51 | -11.4 | -17.7 | **<0.0001** |
| daGal4 | 3.2 | ♂ | 2 | 103 | 63.5 | 88 | 121 | 50.0 | 66 | 119 | 43.4 | 52 | -13.2 | -21.2 | **<0.0001** |
| daGal4 | 3.2 | ♂ | 3 | 116 | 62.3 | 81 | 118 | 64.2 | 81 | 116 | 39.9 | 53 | -35.9 | -34.6 | **<0.0001** |
| daGal4 | 3.2 | ♂ | 4 | 118 | 65.0 | 83 | 117 | 60.5 | 80 | 120 | 42.4 | 52 | -29.9 | -35.0 | **<0.0001** |
| daGal4 | 3.2 | ♂ | 5 | 110 | 60.5 | 73 | 100 | 60.5 | 73 | 115 | 35.9 | 41 | -40.7 | -43.8 | **<0.0001** |
| daGal4 | 3.2 | ♂ | 6 | 116 | 64.1 | 79 | 107 | 59.7 | 72 | 117 | 39.9 | 51 | -33.2 | -29.2 | **<0.0001** |
| daGal4 | 10.1 | ♀ | 1 | 117 | 68.4 | 88 | 119 | 71.5 | 88 | 121 | 62.3 | 81 | -8.9 | -8.0 | **<0.0001** |
| daGal4 | 10.1 | ♀ | 2 | 110 | 63.2 | 87 | 113 | 70.6 | 87 | 115 | 65.2 | 80 | 0.0 | -8.0 | |
| daGal4 | 10.1 | ♀ | 3 | 111 | 63.0 | 80 | 102 | 69.3 | 87 | 97 | 61.0 | 80 | -3.2 | 0.0 | 0.3652 |
| daGal4 | 10.1 | ♀ | 4 | 116 | 64.1 | 86 | 109 | 71.9 | 86 | 115 | 55.9 | 72 | -12.8 | -16.3 | **<0.0001** |
| daGal4 | 10.1 | ♂ | 1 | 113 | 69.3 | 84 | 118 | 64.2 | 81 | 119 | 58.1 | 67 | -9.6 | -17.3 | **<0.0001** |
| daGal4 | 10.1 | ♂ | 2 | 121 | 67.3 | 83 | 117 | 60.5 | 80 | 118 | 57.7 | 69 | -4.5 | -13.8 | 0.0720 |
| daGal4 | 10.1 | ♂ | 3 | 96 | 64.9 | 80 | 100 | 60.5 | 73 | 109 | 57.5 | 66 | -5.0 | -9.6 | **<0.0001** |
| daGal4 | 10.1 | ♂ | 4 | 122 | 63.0 | 82 | 107 | 59.7 | 72 | 119 | 58.6 | 68 | -1.8 | -5.6 | **0.0248** |

### Table S3



| Gal4 | UAS | Sex | Rep[1] | UAS | | | Gal4 | | | Gal4 + UAS | | | % Extension | | *P* |
|---|---|---|---|---|---|---|---|---|---|---|---|---|---|---|---|
| | | | | n | Mean | Max[2] | n | Mean | Max[2] | n | Mean | Max[2] | Mean | Max[2] | |
| daGal4 | 3.1 | ♀ | 1 | 117 | 60.5 | 87 | 117 | 57.0 | 84 | 118 | 73.5 | 91 | 21.6 | 4.6 | **<0.0001** |
| daGal4 | 3.1 | ♀ | 2 | 113 | 63.8 | 87 | 116 | 69.1 | 94 | 119 | 75.2 | 94 | 8.8 | 0.0 | 0.2671 |
| daGal4 | 3.1 | ♀ | 3 | 118 | 83.7 | 106 | 102 | 71.8 | 94 | 108 | 73.5 | 91 | 0.0 | -3.2 | |
| daGal4 | 3.1 | ♀ | 4 | 120 | 78.2 | 105 | 107 | 70.3 | 97 | 121 | 69.3 | 84 | -1.4 | -13.4 | 0.5209 |
| daGal4 | 3.1 | ♀ | 5 | 110 | 68.0 | 84 | 119 | 71.5 | 88 | 115 | 60.5 | 81 | -11.0 | -3.6 | **0.0003** |
| daGal4 | 3.1 | ♀ | 6 | 118 | 58.4 | 80 | 113 | 70.6 | 87 | 110 | 61.3 | 80 | 0.0 | 0.0 | |
| daGal4 | 3.1 | ♀ | 7 | 94 | 62.1 | 73 | 102 | 69.3 | 87 | 101 | 64.3 | 83 | 0.0 | 0.0 | |
| daGal4 | 3.1 | ♀ | 8 | 102 | 54.9 | 75 | 109 | 71.9 | 86 | 118 | 57.3 | 75 | 0.0 | 0.0 | |
| daGal4 | 3.1 | ♂ | 1 | 116 | 48.6 | 78 | 122 | 48.6 | 68 | 121 | 46.6 | 63 | -4.0 | -7.4 | **0.0275** |
| daGal4 | 3.1 | ♂ | 2 | 115 | 45.6 | 57 | 116 | 56.7 | 80 | 119 | 47.6 | 60 | 0.0 | 0.0 | |
| daGal4 | 3.1 | ♂ | 3 | 116 | 72.8 | 98 | 114 | 59.1 | 85 | 122 | 62.3 | 77 | 0.0 | -9.4 | |
| daGal4 | 3.1 | ♂ | 4 | 121 | 67.8 | 87 | 115 | 57.7 | 76 | 118 | 56.9 | 72 | -1.4 | -5.3 | 0.9662 |
| daGal4 | 3.1 | ♂ | 5 | 115 | 66.3 | 81 | 118 | 64.2 | 81 | 116 | 53.8 | 67 | -16.1 | -17.3 | **<0.0001** |
| daGal4 | 3.1 | ♂ | 6 | 119 | 61.2 | 80 | 117 | 60.5 | 80 | 119 | 54.0 | 66 | -10.6 | -17.5 | **<0.0001** |
| daGal4 | 3.1 | ♂ | 7 | 102 | 59.3 | 80 | 100 | 60.5 | 73 | 117 | 51.5 | 62 | -13.1 | -15.1 | **<0.0001** |
| daGal4 | 3.1 | ♂ | 8 | 117 | 59.9 | 79 | 107 | 59.7 | 72 | 119 | 51.6 | 65 | -13.5 | -9.7 | **<0.0001** |
| daGal4 | 4.4 | ♀ | 1 | 118 | 63.7 | 81 | 119 | 71.5 | 88 | 119 | 56.8 | 74 | -10.8 | -8.6 | **<0.0001** |
| daGal4 | 4.4 | ♀ | 2 | 120 | 55.7 | 80 | 113 | 70.6 | 87 | 86 | 62.1 | 76 | 0.0 | -5.0 | |
| daGal4 | 4.4 | ♀ | 3 | 97 | 62.9 | 80 | 102 | 69.3 | 87 | 112 | 54.3 | 73 | -13.6 | -8.8 | **<0.0001** |
| daGal4 | 4.4 | ♀ | 4 | 119 | 52.7 | 79 | 109 | 71.9 | 86 | 113 | 59.8 | 79 | 0.0 | 0.0 | |
| daGal4 | 4.4 | ♂ | 1 | 117 | 61.2 | 74 | 118 | 64.2 | 81 | 117 | 50.5 | 63 | -17.4 | -14.9 | **<0.0001** |
| daGal4 | 4.4 | ♂ | 2 | 119 | 53.6 | 69 | 117 | 60.5 | 80 | 114 | 52.3 | 66 | -2.3 | -4.3 | 0.3705 |
| daGal4 | 4.4 | ♂ | 3 | 108 | 56.1 | 66 | 100 | 60.5 | 73 | 122 | 51.0 | 56 | -9.1 | -15.2 | **<0.0001** |
| daGal4 | 4.4 | ♂ | 4 | 117 | 54.1 | 68 | 107 | 59.7 | 72 | 117 | 53.4 | 65 | -1.3 | -4.4 | **0.0029** |
| DJ694 | 2.1 | ♀ | 1 | 115 | 55.4 | 78 | 112 | 72.7 | 88 | 120 | 79.2 | 91 | 9.0 | 3.4 | **<0.0001** |
| DJ694 | 2.1 | ♀ | 2 | 114 | 66.2 | 94 | 114 | 76.3 | 94 | 120 | 85.8 | 94 | 12.4 | 0.0 | **<0.0001** |
| DJ694 | 2.1 | ♀ | 3 | 120 | 65.1 | 75 | 119 | 74.0 | 83 | 119 | 74.0 | 83 | 0.1 | 0.0 | 0.2063 |
| DJ694 | 2.1 | ♀ | 4 | 118 | 58.0 | 71 | 117 | 68.3 | 82 | 120 | 68.0 | 82 | 0.0 | 0.0 | |
| DJ694 | 2.1 | ♀ | 5 | 119 | 62.2 | 82 | 118 | 72.3 | 90 | 120 | 68.5 | 82 | 0.0 | 0.0 | |
| DJ694 | 2.1 | ♀ | 6 | 121 | 57.2 | 70 | 115 | 69.3 | 81 | 120 | 67.4 | 77 | 0.0 | 0.0 | |
| DJ694 | 2.1 | ♂ | 1 | 117 | 49.4 | 75 | 111 | 51.3 | 64 | 119 | 61.0 | 75 | 19.0 | 0.0 | **<0.0001** |
| DJ694 | 2.1 | ♂ | 2 | 113 | 52.3 | 63 | 113 | 66.0 | 74 | 115 | 60.6 | 74 | 0.0 | 0.0 | |
| DJ694 | 2.1 | ♂ | 3 | 121 | 53.5 | 68 | 121 | 58.2 | 68 | 120 | 54.8 | 72 | 0.0 | 5.9 | |
| DJ694 | 2.1 | ♂ | 4 | 119 | 51.5 | 64 | 122 | 60.9 | 78 | 119 | 57.0 | 71 | 0.0 | 0.0 | |
| DJ694 | 2.1 | ♂ | 5 | 119 | 47.9 | 61 | 119 | 60.0 | 74 | 118 | 57.2 | 71 | 0.0 | 0.0 | |
| DJ694 | 2.1 | ♂ | 6 | 122 | 51.8 | 70 | 121 | 62.6 | 77 | 121 | 56.0 | 63 | 0.0 | -10.0 | |
| DJ694 | 4.3 | ♀ | 1 | 121 | 53.9 | 66 | 118 | 80.7 | 101 | 123 | 71.3 | 85 | 0.0 | 0.0 | |
| DJ694 | 4.3 | ♀ | 2 | 111 | 54.2 | 69 | 90 | 75.2 | 97 | 109 | 72.3 | 84 | 0.0 | 0.0 | |
| DJ694 | 4.3 | ♀ | 3 | 119 | 47.4 | 58 | 119 | 74.0 | 83 | 116 | 58.6 | 68 | 0.0 | 0.0 | |
| DJ694 | 4.3 | ♀ | 4 | 114 | 42.4 | 57 | 117 | 68.3 | 82 | 120 | 52.8 | 71 | 0.0 | 0.0 | |
| DJ694 | 4.3 | ♀ | 5 | 120 | 44.4 | 57 | 118 | 72.3 | 90 | 118 | 59.0 | 71 | 0.0 | 0.0 | |
| DJ694 | 4.3 | ♀ | 6 | 115 | 41.5 | 59 | 115 | 69.3 | 81 | 118 | 55.0 | 77 | 0.0 | 0.0 | |
| DJ694 | 4.3 | ♂ | 1 | 120 | 48.0 | 59 | 114 | 75.5 | 91 | 114 | 49.1 | 63 | 0.0 | 0.0 | |
| DJ694 | 4.3 | ♂ | 2 | 120 | 48.2 | 63 | 105 | 74.6 | 97 | 116 | 51.7 | 65 | 0.0 | 0.0 | |
| DJ694 | 4.3 | ♂ | 3 | 122 | 34.0 | 44 | 121 | 58.2 | 68 | 120 | 42.2 | 51 | 0.0 | 0.0 | |
| DJ694 | 4.3 | ♂ | 4 | 111 | 36.6 | 50 | 122 | 60.9 | 78 | 120 | 39.9 | 50 | 0.0 | 0.0 | |
| DJ694 | 4.3 | ♂ | 5 | 113 | 32.1 | 40 | 119 | 60.0 | 74 | 123 | 39.4 | 43 | 0.0 | 0.0 | |
| DJ694 | 4.3 | ♂ | 6 | 125 | 35.5 | 49 | 121 | 62.6 | 77 | 120 | 41.8 | 49 | 0.0 | 0.0 | |
| DJ694 | 9.1 | ♀ | 1 | 118 | 52.5 | 68 | 119 | 74.0 | 83 | 127 | 63.6 | 83 | 0.0 | 0.0 | |
| DJ694 | 9.1 | ♀ | 2 | 113 | 47.6 | 67 | 117 | 68.3 | 82 | 123 | 59.4 | 78 | 0.0 | 0.0 | |
| DJ694 | 9.1 | ♀ | 3 | 120 | 49.8 | 67 | 118 | 72.3 | 90 | 118 | 56.5 | 82 | 0.0 | 0.0 | |
| DJ694 | 9.1 | ♀ | 4 | 111 | 47.5 | 66 | 115 | 69.3 | 81 | 115 | 60.9 | 77 | 0.0 | 0.0 | |

| Gal4 | UAS | Sex | Rep[1] | UAS | | | Gal4 | | | Gal4 + UAS | | | % Extension | | *P* |
|---|---|---|---|---|---|---|---|---|---|---|---|---|---|---|---|
| | | | | n | Mean | Max[2] | n | Mean | Max[2] | n | Mean | Max[2] | Mean | Max[2] | |
| DJ694 | 9.1 | ♂ | 1 | 118 | 46.8 | 58 | 121 | 58.2 | 68 | 118 | 53.4 | 65 | 0.0 | 0.0 | |
| DJ694 | 9.1 | ♂ | 2 | 120 | 45.3 | 57 | 122 | 60.9 | 78 | 121 | 52.0 | 61 | 0.0 | 0.0 | |
| DJ694 | 9.1 | ♂ | 3 | 119 | 39.1 | 50 | 119 | 60.0 | 74 | 122 | 51.0 | 57 | 0.0 | 0.0 | |
| DJ694 | 9.1 | ♂ | 4 | 123 | 44.9 | 56 | 121 | 62.6 | 77 | 120 | 49.0 | 59 | 0.0 | 0.0 | |
| DJ694 | 3.2 | ♀ | 1 | 118 | 70.5 | 93 | 118 | 64.3 | 84 | 120 | 82.6 | 97 | 17.2 | 4.3 | **<0.0001** |
| DJ694 | 3.2 | ♀ | 2 | 117 | 74.2 | 101 | 118 | 61.8 | 81 | 119 | 87.6 | 101 | 18.1 | 0.0 | **0.0001** |
| DJ694 | 3.2 | ♀ | 3 | 128 | 67.6 | 83 | 119 | 74.0 | 83 | 117 | 75.4 | 83 | 1.9 | 0.0 | 0.8979 |
| DJ694 | 3.2 | ♀ | 4 | 124 | 62.9 | 78 | 117 | 68.3 | 82 | 118 | 69.1 | 82 | 1.2 | 0.0 | 0.1114 |
| DJ694 | 3.2 | ♀ | 5 | 119 | 60.7 | 74 | 118 | 72.3 | 90 | 118 | 69.7 | 82 | 0.0 | 0.0 | |
| DJ694 | 3.2 | ♀ | 6 | 120 | 60.3 | 81 | 115 | 69.3 | 81 | 121 | 68.1 | 85 | 0.0 | 4.9 | |
| DJ694 | 3.2 | ♂ | 1 | 116 | 62.4 | 84 | 118 | 54.4 | 69 | 117 | 74.1 | 91 | 18.8 | 8.3 | **<0.0001** |
| DJ694 | 3.2 | ♂ | 2 | 103 | 63.5 | 88 | 119 | 54.5 | 66 | 117 | 85.4 | 94 | 34.5 | 6.8 | **<0.0001** |
| DJ694 | 3.2 | ♂ | 3 | 120 | 53.0 | 68 | 121 | 58.2 | 68 | 119 | 64.9 | 83 | 11.5 | 22.1 | **<0.0001** |
| DJ694 | 3.2 | ♂ | 4 | 123 | 58.6 | 74 | 122 | 60.9 | 78 | 124 | 64.2 | 74 | 5.5 | 0.0 | 0.0851 |
| DJ694 | 3.2 | ♂ | 5 | 120 | 53.6 | 61 | 119 | 60.0 | 74 | 121 | 61.9 | 82 | 3.1 | 10.8 | 0.0987 |
| DJ694 | 3.2 | ♂ | 6 | 126 | 64.4 | 81 | 121 | 62.6 | 77 | 121 | 66.2 | 85 | 2.7 | 4.9 | 0.5922 |
| DJ694 | 10.1 | ♀ | 1 | 119 | 70.0 | 83 | 119 | 74.0 | 83 | 119 | 79.6 | 91 | 7.6 | 9.6 | **<0.0001** |
| DJ694 | 10.1 | ♀ | 2 | 121 | 65.9 | 82 | 117 | 68.3 | 82 | 121 | 72.8 | 82 | 6.5 | 0.0 | 0.1117 |
| DJ694 | 10.1 | ♀ | 3 | 118 | 62.7 | 82 | 118 | 72.3 | 90 | 118 | 75.0 | 82 | 3.7 | 0.0 | 0.7199 |
| DJ694 | 10.1 | ♀ | 4 | 120 | 67.8 | 85 | 115 | 69.3 | 81 | 123 | 72.8 | 81 | 5.0 | 0.0 | **0.0290** |
| DJ694 | 10.1 | ♂ | 1 | 120 | 58.2 | 72 | 121 | 58.2 | 68 | 119 | 70.3 | 83 | 20.8 | 15.3 | **<0.0001** |
| DJ694 | 10.1 | ♂ | 2 | 118 | 59.6 | 78 | 122 | 60.9 | 78 | 121 | 70.7 | 82 | 16.2 | 5.1 | **<0.0001** |
| DJ694 | 10.1 | ♂ | 3 | 125 | 58.4 | 71 | 119 | 60.0 | 74 | 113 | 69.9 | 82 | 16.4 | 10.8 | **<0.0001** |
| DJ694 | 10.1 | ♂ | 4 | 124 | 64.9 | 81 | 121 | 62.6 | 77 | 122 | 71.2 | 85 | 9.6 | 4.9 | **<0.0001** |
| DJ694 | 3.1 | ♀ | 1 | 117 | 60.5 | 87 | 116 | 59.7 | 84 | 119 | 68.5 | 91 | 13.3 | 4.6 | **0.0029** |
| DJ694 | 3.1 | ♀ | 2 | 113 | 63.8 | 87 | 119 | 72.2 | 91 | 120 | 71.4 | 94 | 0.0 | 3.3 | |
| DJ694 | 3.1 | ♀ | 3 | 118 | 83.7 | 106 | 118 | 80.7 | 101 | 117 | 77.7 | 98 | -3.7 | -3.0 | 0.1472 |
| DJ694 | 3.1 | ♀ | 4 | 120 | 78.2 | 105 | 90 | 75.2 | 97 | 121 | 76.0 | 93 | 0.0 | -4.1 | |
| DJ694 | 3.1 | ♀ | 5 | 119 | 61.2 | 72 | 119 | 74.0 | 83 | 116 | 67.4 | 83 | 0.0 | 0.0 | |
| DJ694 | 3.1 | ♀ | 6 | 107 | 50.4 | 67 | 117 | 68.3 | 82 | 120 | 55.8 | 71 | 0.0 | 0.0 | |
| DJ694 | 3.1 | ♀ | 7 | 112 | 51.7 | 67 | 118 | 72.3 | 90 | 121 | 60.8 | 82 | 0.0 | 0.0 | |
| DJ694 | 3.1 | ♀ | 8 | 119 | 57.2 | 77 | 115 | 69.3 | 81 | 121 | 58.3 | 70 | 0.0 | -9.1 | |
| DJ694 | 3.1 | ♂ | 1 | 116 | 48.6 | 78 | 119 | 54.7 | 71 | 118 | 59.1 | 84 | 8.1 | 7.7 | **0.0004** |
| DJ694 | 3.1 | ♂ | 2 | 115 | 45.6 | 57 | 116 | 63.3 | 84 | 119 | 70.5 | 100 | 11.4 | 19.0 | **<0.0001** |
| DJ694 | 3.1 | ♂ | 3 | 116 | 72.8 | 98 | 114 | 75.5 | 91 | 111 | 70.7 | 85 | -2.9 | -6.6 | **0.0170** |
| DJ694 | 3.1 | ♂ | 4 | 121 | 67.8 | 87 | 105 | 74.6 | 97 | 122 | 77.0 | 97 | 3.3 | 0.0 | **0.0711** |
| DJ694 | 3.1 | ♂ | 5 | 120 | 62.7 | 72 | 121 | 58.2 | 68 | 118 | 61.5 | 75 | 0.0 | 4.2 | |
| DJ694 | 3.1 | ♂ | 6 | 123 | 52.7 | 67 | 122 | 60.9 | 78 | 121 | 56.9 | 78 | 0.0 | 0.0 | |
| DJ694 | 3.1 | ♂ | 7 | 119 | 50.6 | 64 | 119 | 60.0 | 74 | 119 | 61.0 | 71 | 1.6 | 0.0 | 0.2226 |
| DJ694 | 3.1 | ♂ | 8 | 126 | 50.9 | 70 | 121 | 62.6 | 77 | 118 | 61.3 | 81 | 0.0 | 5.2 | |
| DJ694 | 4.4 | ♀ | 1 | 119 | 51.8 | 75 | 119 | 74.0 | 83 | 122 | 65.4 | 83 | 0.0 | 0.0 | |
| DJ694 | 4.4 | ♀ | 2 | 117 | 48.4 | 71 | 117 | 68.3 | 82 | 122 | 60.5 | 82 | 0.0 | 0.0 | |
| DJ694 | 4.4 | ♀ | 3 | 118 | 45.5 | 64 | 118 | 72.3 | 90 | 118 | 62.0 | 82 | 0.0 | 0.0 | |
| DJ694 | 4.4 | ♀ | 4 | 113 | 46.1 | 66 | 115 | 69.3 | 81 | 120 | 61.7 | 81 | 0.0 | 0.0 | |
| DJ694 | 4.4 | ♂ | 1 | 121 | 44.1 | 61 | 121 | 58.2 | 68 | 124 | 53.2 | 65 | 0.0 | 0.0 | |
| DJ694 | 4.4 | ♂ | 2 | 121 | 49.7 | 61 | 122 | 60.9 | 78 | 120 | 55.8 | 71 | 0.0 | 0.0 | |
| DJ694 | 4.4 | ♂ | 3 | 116 | 45.1 | 57 | 119 | 60.0 | 74 | 123 | 53.0 | 61 | 0.0 | 0.0 | |
| DJ694 | 4.4 | ♂ | 4 | 118 | 49.9 | 63 | 121 | 62.6 | 77 | 120 | 59.5 | 77 | 0.0 | 0.0 | |
| D42 | 2.1 | ♀ | 1 | 115 | 55.4 | 78 | 103 | 73.0 | 95 | 120 | 75.7 | 88 | 3.7 | 0.0 | 0.0934 |
| D42 | 2.1 | ♀ | 2 | 114 | 66.2 | 94 | 100 | 76.2 | 97 | 119 | 85.5 | 97 | 12.2 | 0.0 | 0.3860 |
| D42 | 2.1 | ♀ | 3 | 116 | 71.2 | 92 | 104 | 59.1 | 92 | 107 | 76.4 | 92 | 7.4 | 0.0 | **<0.0001** |
| D42 | 2.1 | ♀ | 4 | 121 | 73.9 | 90 | 101 | 61.1 | 94 | 118 | 84.5 | 104 | 14.4 | 10.6 | **<0.0001** |

**Table S3** 

| Gal4 | UAS | Sex | Rep[1] | UAS | | | Gal4 | | | Gal4 + UAS | | | % Extension | | *P* |
|---|---|---|---|---|---|---|---|---|---|---|---|---|---|---|---|
| | | | | n | Mean | Max[2] | n | Mean | Max[2] | n | Mean | Max[2] | Mean | Max[2] | |
| D42 | 2.1 | ♀ | 5 | 118 | 71.4 | 87 | 117 | 61.9 | 84 | 110 | 64.4 | 84 | 0.0 | 0.0 | |
| D42 | 2.1 | ♀ | 6 | 117 | 64.4 | 72 | 114 | 67.1 | 90 | 116 | 64.2 | 83 | -0.3 | 0.0 | 0.0644 |
| D42 | 2.1 | ♀ | 7 | 116 | 62.2 | 76 | 125 | 63.7 | 83 | 129 | 67.8 | 83 | 6.5 | 0.0 | 0.3398 |
| D42 | 2.1 | ♀ | 8 | 116 | 66.4 | 82 | 117 | 64.7 | 75 | 117 | 65.5 | 82 | 0.0 | 0.0 | |
| D42 | 2.1 | ♂ | 1 | 117 | 49.4 | 75 | 103 | 50.3 | 68 | 112 | 53.9 | 71 | 7.1 | 0.0 | 0.1028 |
| D42 | 2.1 | ♂ | 2 | 113 | 52.3 | 63 | 109 | 53.4 | 94 | 117 | 58.0 | 74 | 8.6 | 0.0 | 0.1147 |
| D42 | 2.1 | ♂ | 3 | 117 | 56.0 | 71 | 108 | 54.2 | 78 | 125 | 60.4 | 78 | 7.7 | 0.0 | **0.0038** |
| D42 | 2.1 | ♂ | 4 | 115 | 67.0 | 87 | 111 | 63.2 | 84 | 122 | 66.3 | 84 | 0.0 | 0.0 | |
| D42 | 2.1 | ♂ | 5 | 116 | 61.3 | 77 | 124 | 61.5 | 70 | 108 | 57.1 | 70 | -7.0 | 0.0 | **<0.0001** |
| D42 | 2.1 | ♂ | 6 | 125 | 61.3 | 76 | 117 | 61.9 | 76 | 102 | 55.7 | 72 | -9.1 | -5.3 | **<0.0001** |
| D42 | 2.1 | ♂ | 7 | 114 | 61.1 | 76 | 120 | 58.7 | 76 | 114 | 57.6 | 72 | -1.9 | -5.3 | 0.2165 |
| D42 | 2.1 | ♂ | 8 | 126 | 55.7 | 71 | 120 | 59.8 | 68 | 112 | 56.1 | 68 | 0.0 | 0.0 | |
| D42 | 4.3 | ♀ | 1 | 121 | 53.9 | 66 | 120 | 82.1 | 98 | 116 | 80.1 | 98 | 0.0 | 0.0 | |
| D42 | 4.3 | ♀ | 2 | 111 | 54.2 | 69 | 99 | 78.7 | 97 | 120 | 77.7 | 97 | 0.0 | 0.0 | |
| D42 | 4.3 | ♀ | 3 | 119 | 51.2 | 74 | 104 | 59.1 | 92 | 123 | 66.8 | 88 | 13.0 | 0.0 | 0.5597 |
| D42 | 4.3 | ♀ | 4 | 112 | 57.1 | 80 | 101 | 61.1 | 94 | 93 | 67.7 | 90 | 10.7 | 0.0 | **0.0002** |
| D42 | 4.3 | ♀ | 5 | 119 | 51.4 | 63 | 117 | 61.9 | 84 | 110 | 58.3 | 70 | 0.0 | 0.0 | |
| D42 | 4.3 | ♀ | 6 | 120 | 50.1 | 65 | 114 | 67.1 | 90 | 119 | 58.3 | 72 | 0.0 | 0.0 | |
| D42 | 4.3 | ♀ | 7 | 113 | 47.8 | 58 | 125 | 63.7 | 83 | 116 | 58.6 | 76 | 0.0 | 0.0 | |
| D42 | 4.3 | ♀ | 8 | 120 | 49.1 | 64 | 117 | 64.7 | 75 | 116 | 57.1 | 75 | 0.0 | 0.0 | |
| D42 | 4.3 | ♂ | 1 | 120 | 48.0 | 59 | 113 | 67.6 | 98 | 113 | 38.3 | 50 | -20.1 | -15.3 | **<0.0001** |
| D42 | 4.3 | ♂ | 2 | 120 | 48.2 | 63 | 118 | 66.4 | 93 | 106 | 35.9 | 45 | -25.4 | -28.6 | **<0.0001** |
| D42 | 4.3 | ♂ | 3 | 121 | 46.3 | 57 | 108 | 54.2 | 78 | 122 | 45.3 | 60 | -2.2 | 0.0 | 0.9349 |
| D42 | 4.3 | ♂ | 4 | 95 | 49.5 | 59 | 111 | 63.2 | 84 | 106 | 41.1 | 66 | -17.0 | 0.0 | **0.0001** |
| D42 | 4.3 | ♂ | 5 | 124 | 47.8 | 56 | 124 | 61.5 | 70 | 114 | 51.6 | 70 | 0.0 | 0.0 | |
| D42 | 4.3 | ♂ | 6 | 122 | 41.7 | 55 | 117 | 61.9 | 76 | 117 | 42.7 | 55 | 0.0 | 0.0 | |
| D42 | 4.3 | ♂ | 7 | 119 | 40.5 | 55 | 120 | 58.7 | 76 | 109 | 43.5 | 55 | 0.0 | 0.0 | |
| D42 | 4.3 | ♂ | 8 | 119 | 34.4 | 44 | 120 | 59.8 | 68 | 110 | 38.9 | 54 | 0.0 | 0.0 | |
| D42 | 9.1 | ♀ | 1 | 129 | 57.1 | 81 | 104 | 59.1 | 92 | 116 | 62.8 | 95 | 6.2 | 3.3 | 0.592 |
| D42 | 9.1 | ♀ | 2 | 115 | 70.6 | 90 | 101 | 61.1 | 94 | 117 | 84.2 | 101 | 19.3 | 7.4 | **<0.0001** |
| D42 | 9.1 | ♀ | 3 | 120 | 54.8 | 77 | 117 | 61.9 | 84 | 115 | 64.3 | 77 | 3.9 | 0.0 | 0.1598 |
| D42 | 9.1 | ♀ | 4 | 115 | 49.7 | 62 | 114 | 67.1 | 90 | 118 | 58.7 | 72 | 0.0 | 0.0 | |
| D42 | 9.1 | ♀ | 5 | 118 | 55.9 | 69 | 125 | 63.7 | 83 | 122 | 60.3 | 76 | 0.0 | 0.0 | |
| D42 | 9.1 | ♀ | 6 | 118 | 51.5 | 68 | 117 | 64.7 | 75 | 119 | 55.7 | 75 | 0.0 | 0.0 | |
| D42 | 9.1 | ♂ | 1 | 124 | 43.1 | 57 | 108 | 54.2 | 78 | 120 | 51.8 | 71 | 0.0 | 0.0 | |
| D42 | 9.1 | ♂ | 2 | 123 | 53.9 | 73 | 111 | 63.2 | 84 | 121 | 69.2 | 90 | 9.5 | 7.1 | 0.1051 |
| D42 | 9.1 | ♂ | 3 | 120 | 48.5 | 59 | 124 | 61.5 | 70 | 115 | 55.6 | 66 | 0.0 | 0.0 | |
| D42 | 9.1 | ♂ | 4 | 109 | 45.4 | 58 | 117 | 61.9 | 76 | 111 | 51.3 | 65 | 0.0 | 0.0 | |
| D42 | 9.1 | ♂ | 5 | 116 | 44.2 | 55 | 120 | 58.7 | 76 | 120 | 51.9 | 62 | 0.0 | 0.0 | |
| D42 | 9.1 | ♂ | 6 | 121 | 47.8 | 61 | 120 | 59.8 | 68 | 120 | 48.5 | 61 | 0.0 | 0.0 | |
| D42 | 3.2 | ♀ | 1 | 118 | 70.5 | 93 | 119 | 63.4 | 84 | 112 | 81.2 | 91 | 15.2 | 0.0 | **0.0013** |
| D42 | 3.2 | ♀ | 2 | 117 | 74.2 | 101 | 120 | 66.0 | 88 | 118 | 83.0 | 94 | 11.9 | 0.0 | 0.7125 |
| D42 | 3.2 | ♀ | 3 | 122 | 70.3 | 84 | 117 | 61.9 | 84 | 100 | 69.4 | 84 | 0.0 | 0.0 | |
| D42 | 3.2 | ♀ | 4 | 116 | 60.1 | 72 | 114 | 67.1 | 90 | 119 | 70.4 | 83 | 5.0 | 0.0 | 0.4230 |
| D42 | 3.2 | ♀ | 5 | 116 | 62.0 | 83 | 125 | 63.7 | 83 | 122 | 68.1 | 83 | 7.0 | 0.0 | 0.8400 |
| D42 | 3.2 | ♀ | 6 | 119 | 62.0 | 75 | 117 | 64.7 | 75 | 119 | 62.6 | 75 | 0.0 | 0.0 | |
| D42 | 3.2 | ♂ | 1 | 116 | 62.4 | 84 | 120 | 56.0 | 69 | 116 | 77.3 | 91 | 24.0 | 8.3 | **<0.0001** |
| D42 | 3.2 | ♂ | 2 | 103 | 63.5 | 88 | 115 | 57.7 | 73 | 119 | 79.4 | 94 | 25.0 | 6.8 | **<0.0001** |
| D42 | 3.2 | ♂ | 3 | 120 | 58.0 | 70 | 124 | 61.5 | 70 | 101 | 61.7 | 77 | 0.4 | 10.0 | 0.4165 |
| D42 | 3.2 | ♂ | 4 | 120 | 55.5 | 69 | 117 | 61.9 | 76 | 125 | 65.2 | 83 | 5.2 | 9.2 | **<0.0001** |
| D42 | 3.2 | ♂ | 5 | 125 | 53.7 | 69 | 120 | 58.7 | 76 | 114 | 57.0 | 76 | 0.0 | 0.0 | |
| D42 | 3.2 | ♂ | 6 | 117 | 57.1 | 71 | 120 | 59.8 | 68 | 126 | 63.9 | 75 | 6.9 | 5.6 | **<0.0001** |

### Table S3

| Gal4 | UAS | Sex | Rep[1] | UAS | | | Gal4 | | | Gal4 + UAS | | | % Extension | | *P* |
|---|---|---|---|---|---|---|---|---|---|---|---|---|---|---|---|
| | | | | n | Mean | Max[2] | n | Mean | Max[2] | n | Mean | Max[2] | Mean | Max[2] | |
| D42 | 10.1 | ♀ | 1 | 120 | 57.6 | 77 | 106 | 77.9 | 95 | 116 | 78.9 | 88 | 1.2 | 0.0 | **0.0114** |
| D42 | 10.1 | ♀ | 2 | 109 | 66.6 | 84 | 106 | 74.0 | 91 | 110 | 71.6 | 88 | 0.0 | 0.0 | |
| D42 | 10.1 | ♀ | 3 | 119 | 68.5 | 84 | 117 | 61.9 | 84 | 115 | 67.8 | 84 | 0.0 | 0.0 | |
| D42 | 10.1 | ♀ | 4 | 105 | 66.8 | 86 | 114 | 67.1 | 90 | 117 | 70.2 | 90 | 4.6 | 0.0 | **0.0329** |
| D42 | 10.1 | ♀ | 5 | 123 | 63.9 | 83 | 125 | 63.7 | 83 | 114 | 59.4 | 76 | -6.8 | -8.4 | **0.0016** |
| D42 | 10.1 | ♀ | 6 | 124 | 59.5 | 75 | 117 | 64.7 | 75 | 110 | 65.6 | 89 | 1.3 | 18.7 | **0.0262** |
| D42 | 10.1 | ♂ | 1 | 121 | 57.2 | 74 | 119 | 59.2 | 81 | 115 | 67.9 | 81 | 14.6 | 0.0 | **0.0053** |
| D42 | 10.1 | ♂ | 2 | 124 | 56.8 | 67 | 116 | 60.4 | 88 | 123 | 68.7 | 81 | 13.8 | 0.0 | 0.0531 |
| D42 | 10.1 | ♂ | 3 | 120 | 66.4 | 77 | 124 | 61.5 | 70 | 117 | 71.1 | 84 | 7.1 | 9.1 | **<0.0001** |
| D42 | 10.1 | ♂ | 4 | 122 | 65.1 | 86 | 117 | 61.9 | 76 | 109 | 75.7 | 98 | 16.4 | 14.0 | **<0.0001** |
| D42 | 10.1 | ♂ | 5 | 115 | 59.2 | 76 | 120 | 58.7 | 76 | 117 | 61.3 | 76 | 3.5 | 0.0 | 0.3541 |
| D42 | 10.1 | ♂ | 6 | 127 | 62.7 | 89 | 120 | 59.8 | 68 | 115 | 68.3 | 82 | 8.9 | 0.0 | **0.0116** |
| D42 | 3.1 | ♀ | 1 | 117 | 60.5 | 87 | 117 | 66.9 | 87 | 119 | 74.4 | 91 | 11.2 | 4.6 | **<0.0001** |
| D42 | 3.1 | ♀ | 2 | 113 | 63.8 | 87 | 112 | 74.0 | 97 | 120 | 79.4 | 100 | 7.3 | 3.1 | 0.0642 |
| D42 | 3.1 | ♀ | 3 | 118 | 83.7 | 106 | 120 | 82.1 | 98 | 119 | 89.0 | 106 | 6.4 | 0.0 | 0.6531 |
| D42 | 3.1 | ♀ | 4 | 120 | 78.2 | 105 | 99 | 78.7 | 97 | 114 | 84.7 | 97 | 7.5 | 0.0 | **0.0105** |
| D42 | 3.1 | ♀ | 5 | 118 | 67.5 | 80 | 117 | 61.9 | 84 | 128 | 67.2 | 87 | 0.0 | 3.6 | |
| D42 | 3.1 | ♀ | 6 | 127 | 62.0 | 72 | 114 | 67.1 | 90 | 122 | 70.6 | 98 | 5.2 | 8.9 | **0.0095** |
| D42 | 3.1 | ♀ | 7 | 125 | 61.8 | 76 | 125 | 63.7 | 83 | 119 | 71.6 | 83 | 12.5 | 0.0 | **0.0225** |
| D42 | 3.1 | ♀ | 8 | 110 | 59.2 | 75 | 117 | 64.7 | 75 | 119 | 69.3 | 89 | 7.1 | 18.7 | **0.0003** |
| D42 | 3.1 | ♂ | 1 | 116 | 48.6 | 78 | 120 | 55.2 | 74 | 118 | 54.2 | 68 | 0.0 | -8.1 | |
| D42 | 3.1 | ♂ | 2 | 115 | 45.6 | 57 | 115 | 60.7 | 73 | 118 | 66.0 | 91 | 8.7 | 24.7 | **<0.0001** |
| D42 | 3.1 | ♂ | 3 | 116 | 72.8 | 98 | 113 | 67.6 | 98 | 113 | 66.5 | 77 | -1.6 | -21.4 | 0.1023 |
| D42 | 3.1 | ♂ | 4 | 121 | 67.8 | 87 | 118 | 66.4 | 93 | 104 | 69.0 | 84 | 1.6 | -3.4 | 0.8434 |
| D42 | 3.1 | ♂ | 5 | 108 | 60.4 | 70 | 124 | 61.5 | 70 | 120 | 58.1 | 73 | -3.9 | 4.3 | **0.0055** |
| D42 | 3.1 | ♂ | 6 | 125 | 61.2 | 76 | 117 | 61.9 | 76 | 117 | 62.4 | 76 | 0.7 | 0.0 | 0.2374 |
| D42 | 3.1 | ♂ | 7 | 117 | 59.7 | 76 | 120 | 58.7 | 76 | 120 | 58.6 | 72 | -0.1 | -5.3 | 0.3712 |
| D42 | 3.1 | ♂ | 8 | 118 | 56.1 | 75 | 120 | 59.8 | 68 | 113 | 60.1 | 75 | 0.5 | 0.0 | **0.0246** |
| D42 | 4.4 | ♀ | 1 | 117 | 59.8 | 88 | 104 | 59.1 | 92 | 115 | 75.0 | 99 | 25.4 | 7.6 | **<0.0001** |
| D42 | 4.4 | ♀ | 2 | 115 | 67.8 | 97 | 101 | 61.1 | 94 | 105 | 80.6 | 104 | 18.9 | 7.2 | **<0.0001** |
| D42 | 4.4 | ♀ | 3 | 137 | 61.8 | 77 | 117 | 61.9 | 84 | 117 | 60.4 | 87 | -2.2 | 3.6 | 0.9710 |
| D42 | 4.4 | ♀ | 4 | 119 | 53.9 | 69 | 114 | 67.1 | 90 | 117 | 63.1 | 83 | 0.0 | 0.0 | |
| D42 | 4.4 | ♀ | 5 | 127 | 56.3 | 76 | 125 | 63.7 | 83 | 118 | 58.2 | 76 | 0.0 | 0.0 | |
| D42 | 4.4 | ♀ | 6 | 123 | 56.6 | 75 | 117 | 64.7 | 75 | 109 | 62.0 | 82 | 0.0 | 9.3 | |
| D42 | 4.4 | ♂ | 1 | 123 | 54.7 | 78 | 108 | 54.2 | 78 | 125 | 65.2 | 85 | 19.1 | 9.0 | **<0.0001** |
| D42 | 4.4 | ♂ | 2 | 120 | 68.5 | 84 | 111 | 63.2 | 84 | 123 | 70.8 | 90 | 3.4 | 7.1 | **0.0278** |
| D42 | 4.4 | ♂ | 3 | 113 | 55.0 | 70 | 124 | 61.5 | 70 | 112 | 54.7 | 70 | -0.6 | 0.0 | 0.5016 |
| D42 | 4.4 | ♂ | 4 | 109 | 52.3 | 69 | 117 | 61.9 | 76 | 119 | 50.8 | 69 | -2.9 | 0.0 | 0.2690 |
| D42 | 4.4 | ♂ | 5 | 117 | 52.6 | 69 | 120 | 58.7 | 76 | 116 | 56.5 | 76 | 0.0 | 0.0 | |
| D42 | 4.4 | ♂ | 6 | 121 | 53.0 | 68 | 120 | 59.8 | 68 | 109 | 53.3 | 68 | 0.0 | 0.0 | |

**Table S3: Longevity data and analysis of Hsp70 overexpression.** Mean: average survival in days, n: number of flies in the replicate, P: highest Log-rank test value, 1: replicate number, 2: 95% mortality (maximum lifespan)
