## Supplementary material for "Expression of heat shock protein 70 is insufficient to extend *Drosophila melanogaster* longevity": Table S4

| Gal4 | UAS | Sex | Rep[1] | UAS | | | Gal4 | | | Gal4 + UAS | | | % Extension | | *P* |
|---|---|---|---|---|---|---|---|---|---|---|---|---|---|---|---|
| | | | | n | Mean | Max[2] | n | Mean | Max[2] | n | Mean | Max[2] | Mean | Max[2] | |
| daGal4 | R1.26 | ♀ | 1 | 120 | 50.0 | 72 | 117 | 76.4 | 92 | 118 | 56.0 | 81 | 0.0 | 0.0 | |
| daGal4 | R1.26 | ♀ | 2 | 116 | 52.9 | 84 | 119 | 79.6 | 98 | 118 | 55.0 | 91 | 0.0 | 0.0 | |
| daGal4 | R1.26 | ♀ | 3 | 116 | 52.9 | 84 | 119 | 79.6 | 98 | 118 | 55.0 | 91 | 0.0 | 0.0 | |
| daGal4 | R1.26 | ♂ | 1 | 120 | 48.8 | 75 | 107 | 62.1 | 83 | 120 | 50.6 | 75 | 0.0 | -0.5 | |
| daGal4 | R1.26 | ♂ | 2 | 122 | 48.9 | 67 | 115 | 58.0 | 67 | 119 | 51.3 | 67 | 0.0 | 0.0 | |
| daGal4 | R1.26 | ♂ | 3 | 122 | 48.9 | 67 | 115 | 58.0 | 67 | 119 | 51.3 | 67 | 0.0 | 0.0 | |
| daGal4 | R3.12 | ♀ | 1 | 119 | 55.8 | 79 | 116 | 55.8 | 79 | 121 | 67.9 | 91 | 21.5 | 15.2 | **<0.0001** |
| daGal4 | R3.12 | ♀ | 2 | 118 | 52.5 | 73 | 116 | 61.7 | 88 | 118 | 71.4 | 94 | 15.8 | 6.8 | **<0.0001** |
| daGal4 | R3.12 | ♂ | 1 | 120 | 53.2 | 69 | 118 | 48.1 | 62 | 120 | 60.5 | 76 | 13.9 | 10.1 | **<0.0001** |
| daGal4 | R3.12 | ♂ | 2 | 120 | 50.1 | 66 | 121 | 50.0 | 66 | 118 | 63.1 | 81 | 26.1 | 23.0 | **<0.0001** |
| daGal4 | R5.14 | ♀ | 1 | 118 | 50.5 | 72 | 117 | 76.4 | 92 | 119 | 57.9 | 92 | 0.0 | 0.0 | |
| daGal4 | R5.14 | ♀ | 2 | 119 | 53.3 | 87 | 119 | 79.6 | 98 | 120 | 55.4 | 98 | 0.0 | 0.0 | |
| daGal4 | R5.14 | ♀ | 3 | 29 | 59.0 | 73 | 32 | 75.2 | 88 | 29 | 57.1 | 77 | -3.2 | 0.0 | 0.7638 |
| daGal4 | R5.14 | ♂ | 1 | 120 | 36.9 | 57 | 107 | 62.1 | 83 | 123 | 48.6 | 68 | 0.0 | 0.0 | |
| daGal4 | R5.14 | ♂ | 2 | 117 | 42.8 | 56 | 115 | 58.0 | 67 | 121 | 51.3 | 67 | 0.0 | 0.0 | |
| daGal4 | R5.14 | ♂ | 3 | 30 | 38.7 | 55 | 29 | 73.0 | 88 | 30 | 50.6 | 60 | 0.0 | 0.0 | |
| DJ694 | R1.26 | ♀ | 1 | 120 | 50.0 | 72 | 120 | 71.4 | 85 | 116 | 63.9 | 85 | 0.0 | 0.5 | |
| DJ694 | R1.26 | ♀ | 2 | 116 | 52.9 | 84 | 122 | 81.5 | 98 | 115 | 64.8 | 92 | 0.0 | 0.0 | |
| DJ694 | R1.26 | ♀ | 3 | 116 | 52.9 | 84 | 122 | 81.5 | 98 | 115 | 64.8 | 92 | 0.0 | 0.0 | |
| DJ694 | R1.26 | ♂ | 1 | 120 | 48.8 | 75 | 116 | 62.6 | 68 | 122 | 54.4 | 71 | 0.0 | 0.0 | |
| DJ694 | R1.26 | ♂ | 2 | 122 | 48.9 | 67 | 116 | 62.6 | 67 | 116 | 56.7 | 67 | 0.0 | 0.0 | |
| DJ694 | R1.26 | ♂ | 3 | 122 | 48.9 | 67 | 116 | 62.6 | 67 | 116 | 56.7 | 67 | 0.0 | 0.0 | |
| DJ694 | R3.12 | ♀ | 1 | 119 | 55.8 | 79 | 118 | 64.3 | 84 | 120 | 83.1 | 97 | 29.4 | 15.5 | **<0.0001** |
| DJ694 | R3.12 | ♀ | 2 | 118 | 52.5 | 73 | 118 | 61.8 | 81 | 120 | 76.1 | 97 | 23.0 | 19.8 | **<0.0001** |
| DJ694 | R3.12 | ♂ | 1 | 120 | 53.2 | 69 | 118 | 54.4 | 69 | 119 | 73.9 | 97 | 35.9 | 40.6 | **<0.0001** |
| DJ694 | R3.12 | ♂ | 2 | 120 | 50.1 | 66 | 119 | 54.5 | 66 | 119 | 75.4 | 94 | 38.3 | 42.4 | <0.0001 |
| DJ694 | R5.14 | ♀ | 1 | 118 | 50.5 | 72 | 120 | 71.4 | 85 | 118 | 65.0 | 92 | 0.0 | 8.2 | |
| DJ694 | R5.14 | ♀ | 2 | 119 | 53.3 | 87 | 122 | 81.5 | 98 | 112 | 64.4 | 95 | 0.0 | 0.0 | |
| DJ694 | R5.14 | ♀ | 3 | 29 | 59.0 | 73 | 30 | 76.9 | 93 | 28 | 77.6 | 88 | 0.9 | 0.0 | |
| DJ694 | R5.14 | ♂ | 1 | 120 | 36.9 | 57 | 116 | 62.6 | 68 | 117 | 49.9 | 68 | 0.0 | 0.0 | |
| DJ694 | R5.14 | ♂ | 2 | 117 | 42.8 | 56 | 116 | 62.6 | 67 | 119 | 53.0 | 67 | 0.0 | 0.0 | |
| DJ694 | R5.14 | ♂ | 3 | 30 | 38.7 | 55 | 29 | 58.0 | 60 | 30 | 47.7 | 60 | 0.0 | 0.0 | |
| D42 | R1.17 | ♀ | 1 | 28 | 73.2 | 88 | 30 | 81.2 | 95 | 30 | 84.6 | 90 | 4.2 | 0.0 | 0.7681 |
| D42 | R1.17 | ♂ | 1 | 29 | 59.7 | 76 | 30 | 78.3 | 91 | 29 | 64.8 | 74 | 0.0 | -2.6 | |
| D42 | R1.26 | ♀ | 1 | 120 | 50.0 | 72 | 119 | 75.9 | 92 | 115 | 61.1 | 76 | 0.0 | 0.0 | |
| D42 | R1.26 | ♀ | 2 | 116 | 52.9 | 84 | 119 | 84.7 | 98 | 116 | 72.5 | 98 | 0.0 | 0.0 | |
| D42 | R1.26 | ♀ | 3 | 116 | 52.9 | 84 | 119 | 84.7 | 98 | 116 | 72.5 | 98 | 0.0 | 0.0 | |
| D42 | R1.26 | ♂ | 1 | 120 | 48.8 | 75 | 118 | 59.4 | 68 | 115 | 52.9 | 68 | 0.0 | 0.0 | |
| D42 | R1.26 | ♂ | 2 | 122 | 48.9 | 67 | 119 | 60.2 | 67 | 117 | 54.5 | 67 | 0.0 | 0.0 | |
| D42 | R1.26 | ♂ | 3 | 122 | 48.9 | 67 | 119 | 60.2 | 67 | 117 | 54.5 | 67 | 0.0 | 0.0 | |
| D42 | R3.12 | ♀ | 1 | 119 | 55.8 | 79 | 119 | 63.4 | 84 | 118 | 69.9 | 91 | 10.4 | 8.3 | **0.0003** |
| D42 | R3.12 | ♀ | 2 | 118 | 52.5 | 73 | 120 | 66.0 | 88 | 119 | 71.0 | 94 | 7.6 | 6.8 | **0.0145** |
| D42 | R3.12 | ♀ | 3 | 27 | 69.4 | 89 | 30 | 81.2 | 95 | 30 | 65.9 | 90 | -5.0 | 0.0 | **0.0171** |
| D42 | R3.12 | ♂ | 1 | 120 | 53.2 | 69 | 120 | 56.0 | 69 | 118 | 61.8 | 76 | 10.4 | 10.1 | **<0.0001** |
| D42 | R3.12 | ♂ | 2 | 120 | 50.1 | 66 | 115 | 57.7 | 73 | 118 | 65.5 | 88 | 13.6 | 20.5 | **<0.0001** |
| D42 | R3.12 | ♂ | 3 | 30 | 70.1 | 88 | 30 | 78.3 | 91 | 30 | 62.4 | 83 | -10.9 | -5.5 | **0.031** |
| D42 | R5.2 | ♀ | 1 | 30 | 65.0 | 86 | 30 | 81.2 | 95 | 27 | 59.9 | 83 | -7.9 | -3.5 | 0.2044 |
| D42 | R5.2 | ♂ | 1 | 30 | 73.9 | 93 | 30 | 78.3 | 91 | 29 | 58.7 | 78 | -20.6 | -15.2 | **<0.0001** |
| D42 | R5.12e | ♀ | 1 | 30 | 60.9 | 83 | 30 | 81.2 | 95 | 25 | 55.8 | 93 | -8.3 | 0.0 | 0.7819 |
| D42 | R5.12e | ♂ | 1 | 31 | 59.0 | 76 | 30 | 78.3 | 91 | 30 | 60.1 | 82 | 0.0 | 0.0 | |

### Table S4



| Gal4 | UAS | Sex | Rep[1] | UAS | | | Gal4 | | | Gal4 + UAS | | | % Extension | | *P* |
|---|---|---|---|---|---|---|---|---|---|---|---|---|---|---|---|
| | | | | n | Mean | Max[2] | n | Mean | Max[2] | n | Mean | Max[2] | Mean | Max[2] | |
| D42 | R5.14 | ♀ | 1 | 118 | 50.5 | 72 | 119 | 75.9 | 92 | 114 | 65.4 | 82 | 0.0 | 0.0 | |
| D42 | R5.14 | ♀ | 2 | 119 | 53.3 | 87 | 119 | 84.7 | 98 | 117 | 64.3 | 91 | 0.0 | 0.0 | |
| D42 | R5.14 | ♀ | 3 | 29 | 59.0 | 73 | 32 | 79.0 | 88 | 29 | 58.9 | 77 | -0.1 | 0.0 | |
| D42 | R5.14 | ♂ | 1 | 120 | 36.9 | 57 | 118 | 59.4 | 68 | 119 | 46.8 | 63 | 0.0 | 0.0 | |
| D42 | R5.14 | ♂ | 2 | 117 | 42.8 | 56 | 119 | 60.2 | 67 | 119 | 53.9 | 63 | 0.0 | 0.0 | |
| D42 | R5.14 | ♂ | 3 | 30 | 38.7 | 55 | 29 | 58.6 | 60 | 29 | 44.6 | 50 | 0.0 | -8.3 | |

**Table S4: Longevity data and analysis of Hsp70 RNAi.** Mean: average survival in days, n: number of flies in the replicate, P: highest Log-rank test value, 1: replicate number, 2: 95% mortality (maximum lifespan)
