## Supplementary material for "Expression of heat shock protein 70 is insufficient to extend *Drosophila melanogaster* longevity": Reagents table

| Data type | Experimental species | Symbol/name used in publication | Source – public | Source – published | Source – unpublished | Identifiers | New reagent | Comments |
|---|---|---|---|---|---|---|---|---|
| gene (source not applicable) | D. melanogaster | hsp70 | NA | NA | NA | Flybase:FBgn0286924 | | |
| genetic reagent (in whole organism) | D. melanogaster | da-GAL4 | Bloomington Drosophila Stock Center | PMID:7606787 | | BDSC:55851 | | |
| genetic reagent (in whole organism) | D. melanogaster | D42 | Bloomington Drosophila Stock Center | PMID:7624365 | | BDSC:8816 | | |
| genetic reagent (in whole organism) | D. melanogaster | DJ694 | Bloomington Drosophila Stock Center | PMID:12882353 | | BDSC:8176 | | |
| genetic reagent (in whole organism) | D. melanogaster | UAS-lacZ | Bloomington Drosophila Stock Center | | | BDSC:1777 | | |
| genetic reagent (in whole organism) | D. melanogaster | UAS-hsp70 #2.1 | Bloomington Drosophila Stock Center | PMID:17443800 | | | | |
| genetic reagent (in whole organism) | D. melanogaster | UAS-hsp70 #3.1 | Bloomington Drosophila Stock Center | PMID:17443800 | | | | |
| genetic reagent (in whole organism) | D. melanogaster | UAS-hsp70 #3.2 | Bloomington Drosophila Stock Center | PMID:17443800 | | | | |
| genetic reagent (in whole organism) | D. melanogaster | UAS-hsp70 #4.3 | Bloomington Drosophila Stock Center | PMID:17443800 | | | | |
| genetic reagent (in whole organism) | D. melanogaster | UAS-hsp70 #4.4 | Bloomington Drosophila Stock Center | PMID:17443800 | | | | |
| genetic reagent (in whole organism) | D. melanogaster | UAS-hsp70 #9.1 | Bloomington Drosophila Stock Center | PMID:17443800 | | | | |
| genetic reagent (in whole organism) | D. melanogaster | UAS-hsp70 #10.1 | Bloomington Drosophila Stock Center | PMID:17443800 | | | | |
| genetic reagent (in whole organism) | D. melanogaster | UAS-hsp70-RNAi R1.17 | | this paper | | | pCX9 3rd chromosome transformant | |
| genetic reagent (in whole organism) | D. melanogaster | UAS-hsp70-RNAi R1.26 | | this paper | | | pCX9 2nd chromosome transformant | |
| genetic reagent (in whole organism) | D. melanogaster | UAS-hsp70-RNAi R2.17 | | this paper | | | pCX9 3rd chromosome transformant | |
| genetic reagent (in whole organism) | D. melanogaster | UAS-hsp70-RNAi R3.12 | | this paper | | | pCX9 X chromosome transformant | |
| genetic reagent (in whole organism) | D. melanogaster | UAS-hsp70-RNAi R4.15 | | this paper | | | pCX9 3rd chromosome transformant | |
| genetic reagent (in whole organism) | D. melanogaster | UAS-hsp70-RNAi R5.2 | | this paper | | | pCX9 3rd chromosome transformant | |
| genetic reagent (in whole organism) | D. melanogaster | UAS-hsp70-RNAi R5.12 | | this paper | | | pCX9 2nd chromosome transformant | |
| genetic reagent (in whole organism) | D. melanogaster | UAS-hsp70-RNAi R5.12e | | this paper | | | pCX9 3rd chromosome transformant | |
| genetic reagent (in whole organism) | D. melanogaster | UAS-hsp70-RNAi R5.14 | | this paper | | | pCX9 3rd chromosome transformant | |
| antibody | M. musculus | myc 9E10 | Roche | | | 11667203001 | | |
| antibody | M. musculus | hsp70 5A5 | ThermoFisher Scientific | | | MA3-007 | | |
| antibody | M. musculus | a-Tubulin DM1A | Abcam | | | Ab7291 | | |
| antibody | C. hircus | anti-mouse HRP | Bio-Rad | | | 170-6516 | | |
| recombinant DNA reagent | E. coli | pCX1 (pINDY5-hsp70) | | PMID:17443800 | | | | |
| recombinant DNA reagent | E. coli | Sym-pUASTw | | PMID:11861567 | | | | |
| recombinant DNA reagent | E. coli | pCX9.1 | | this paper | | | Progenitors: pCX1; Sym-pUASTw | |
| recombinant DNA reagent | E. coli | pCX9 | | this paper | | | Progenitors: pCX9.1 | |
